# Genome based Evolutionary study of SARS-CoV-2 towards the Prediction of Epitope Based Chimeric Vaccine

**DOI:** 10.1101/2020.04.15.036285

**Authors:** Mst Rubaiat Nazneen Akhand, Kazi Faizul Azim, Syeda Farjana Hoque, Mahmuda Akther Moli, Bijit Das Joy, Hafsa Akter, Ibrahim Khalil Afif, Nadim Ahmed, Mahmudul Hasan

## Abstract

SARS-CoV-2 is known to infect the neurological, respiratory, enteric, and hepatic systems of human and has already become an unprecedented threat to global healthcare system. COVID-19, the most serious public condition caused by SARS-CoV-2 leads the world to an uncertainty alongside thousands of regular death scenes. Unavailability of specific therapeutics or approved vaccine has made the recovery of COVI-19 more troublesome and challenging. The present *in silico* study aimed to predict a novel chimeric vaccines by simultaneously targeting four major structural proteins via the establishment of ancestral relationship among different strains of coronaviruses. Conserved regions from the homologous protein sets of spike glycoprotein (S), membrane protein (M), envelope protein and nucleocapsid protein (N) were identified through multiple sequence alignment. The phylogeny analyses of whole genome stated that four proteins (S, E, M and N) reflected the close ancestral relation of SARS-CoV-2 to SARS-COV-1 and bat coronavirus. Numerous immunogenic epitopes (both T cell and B cell) were generated from the common fragments which were further ranked on the basis of antigenicity, transmembrane topology, conservancy level, toxicity and allergenicity pattern and population coverage analysis. Top putative epitopes were combined with appropriate adjuvants and linkers to construct a novel multiepitope subunit vaccine against COVID-19. The designed constructs were characterized based on physicochemical properties, allergenicity, antigenicity and solubility which revealed the superiority of construct V3 in terms safety and efficacy. Essential molecular dynamics and Normal Mode analysis confirmed minimal deformability of the refined model at molecular level. In addition, disulfide engineering was investigated to accelerate the stability of the protein. Molecular docking study ensured high binding affinity between construct V3 and HLA cells, as well as with different host receptors. Microbial expression and translational efficacy of the constructs were checked using pET28a(+) vector of *E. coli* strain K12. The development of preventive measures to combat COVID-19 infections might be aided the present study. However, the *in vivo* and *in vitro* validation might be ensured with wet lab trials using model animals for the implementation of the presented data.

## 1. Introduction

Novel coronavirus named SARS-CoV-2/ 2019-nCoV was identified at the end of 2019 in Wuhan, a city in the Hubei province of China, causing severe pneumonia that leads to huge death cases (Wang et al., 2020). Gradually this virus emerged as a new threat to the whole world and affecting almost all parts of the world. To date, the pathogen has affected 198 countries, and thus becoming a global public health emergency. Global public health concern with pandemic notion of COVID-19 was declared on January 30th, 2020 by the World health organization (WHO, 2020). Agin, an adverse situation has also been announced on 13 March 2020 for increasing the infections of COVID-19 (Kunz and Minder, 2020). Till April 10, 2020, total virus affected people around the world exceeded 1,633,272 and more than 97,601 committed death, while 366,610 people fully recovered from the infection (WHO, 2020). The alarming situation is that the number of confirmed cases worldwide has exceeded one million by this time. It took more than three months to reach the first 10000 confirmed cases, while required only 12 days to detect the next 100000 cases. The situation is getting worse in European region. Total death cases in Italy, Spain, USA, France, United Kingdom was 14681,11744,7847,6507 and 4313 respectively (till April 4, 2020) and this number is exacerbating day by day (WHO, 2020).

Some common clinical manifestations of COVID-19 is fever, sputum production, shortness of breath, cough, fatigue, sore throat and headache which leads to severe cases of pneumonia. A few patients also have gastrointestinal symptoms with diarrhea and vomiting (Guan et al., 2019). Though several early studies showed that the mortality rate for SARS-CoV-2 is not as high (2- 3%), the latest global death rate for COVID-19 is 3.4% which indicates the increasing trends (Wu et al., 2020). The investigation of Chinese Center for Diseases Control and Prevention (2020) revealed that the prevalence of COVID-19 is more apparent in the people ages 50 years rather than the lower age groups (Jeong-ho et al., 2020). High fever and Lymphocytopenia were found more common in Covid-19, though the frequency of the patient without fever condition is also higher than in the earlier outbreaks caused by SARS-CoV (1%) and MERS-CoV (2%) (Huang et al., 2020; Chen et al., 2020).

SARS-CoV-2 is a betacoronavirus that has a positive sense, 26-32 kb in length, single stranded RNA molecule as its genetic material and belongs to the family Coronaviridae, order Nidovirales (Hui et al., 2019). It shares genome similarity with SARS-CoV (79.5%) and bat coronavirus (96%) (Zhou et al., 2020; Zhu et al., 2020). However, there are still obscured hypothesis regarding the vector or carrier of SARS-CoV-2, though its detection was primarily linked to Wuhan’s Huanan Seafood wholesale market (Lu et al., 2020; WHO, 2020). Though the species of SARS-CoV-1 and bat coronavirus shares sufficient sequence similarities with the COVID-19, the known way mechanism of infection to the host, and the death rate is quite different in case of the novel coronavirus. In addition, there is an evolutionary distance between SARS-CoV-1 and bat coronavirus as well as the COVID-19 (Hu et al., 2018; Wu et al., 2020; Wu, 2020b). Because of high sequence variability of the pathogen, many of the efforts that have been undertaken to develop vaccine against SARS-CoV2, remain unsuccessful (Graham et al., 2013). Therefore, there is an urgent need to develop vaccines for treatment of SARS-CoV-2 based on the understanding of actual evolutionary ancestral relationship. While some natural metabolites and traditional medication may come up with comfort and take the edge off few symptoms of COVID-19, there is no proof that existing treatment procedures can effectively combat against the diseased condition (WHO, 2020). However, inactivated or live-attenuated forms of pathogenic organisms are usually recommended for the initiation of antigen-specific responses that alleviate or reduce the possibility of host experience with secondary infections (Thompson & Staats, 2011). Moreover, all of the proteins are not usually targeted for protective immunity, whereas only a few numbers of proteins are necessary depending on the microbes (Tesh et al., 200, Li et al., 2014). Depending on sufficient antigen expression from experimental assays, traditional vaccine could take 15 years to develop, while sometimes can lead to undesirable consequences (Purcell et al., 2007, Petrovsky & Aguilar, 2004).

Reverse vaccinology approach, on the other hand, is an effective way to develop vaccine against COVID-19. In this method, computation analysis towards genomic architecture of pathogenic candidate could predict the antigens of pathogens without the prerequisite to culture the pathogens in lab condition. Although, few pathogens that challenge to develop effective vaccines so far may become possible through such approach (Rappuoli, 2000) which initiates a huge move in the development of vaccine against the deadly pathogens. The strategy included the comprehensive utilization of bioinformatics algorithm or tools to develop epitope based vaccine molecules, though further validation and experimental procedures are also needed (Moxon et al., 2019). In addition, peptide based subunit vaccines are biologically safer due to the absence of continuous *in vitro* culture during the production period, and also implies an appropriate activation of immune responses (Purcell et al., 2007; Dudek et al., 2010). Such immunoinformatic approaches have already been employed by the researchers to design vaccines against a number of deadly pathogens including Ebola virus (Khan et al., 2015), HIV (Pandey et al., 2018), Areanaviruses (Azim et al., 2019a), Marburgvirus (Hasan et al., 2019a), Norwalk virus (Azim et al., 2019b), Nipah virus (Saha et al., 2017), influenza virus (Hasan et al., 2019b) and so on. At present, a suitable peptide vaccine against SARS-CoV-2 is urgently necessary that could efficiently generate enough immune response to destroy the virus. Hence, the study was designed to develop a chimeric recombinant vaccine against COVID-19 by targeting four major structural proteins of the pathogen, while revealing the evolutionary history of different species of coronavirus based on whole genome and protein domain-based phylogeny.

## 2. Materials and Methods

### 2.1. Data Acquisition

Complete Genomes of the COVID-19 and other coronaviruses were retrieved from the NCBI (https://www.ncbi.nlm.nih.gov/), using the keyword ‘coronavirus’ and the search option ‘nucleotide’. A total 61 complete genomes were retrieved, with unique identity (Supplementary File 1). Protein sequence of the spike, envelope, membrane and nucleocapsid were also retrieved from the corresponding genome sequences found in NCBI (Supplementary File 1).

### 2.2. Phylogeny construction and visualization

Tthe complete genome sequences of coronaviruses and the proteins of envelope, envelope, membrane and nucleocapsid were employed to construct different phylogenetic trees. Multiple sequence alignment (MSA) of the complete genome and protein sequences were performed using MAFFT v7.310 (Katoh & Standley, 2013) tool. For the whole genome alignment, we used MAFFT Auto algorithm, while for the protein sequences alignment, MAFFT G-INS-I algorithm was used using default parameters. Next, alignment was visualized using the JalView-2.11 (Waterhouse et al., 2009). Alignment position with more than 50% gaps was pruned from coronavirus genome using Phyutility 2.2.6 program (Smith & Dunn, 2008). Again, more than 20% gaps from the spike protein alignment was removed. PartitionFinder-2.1.1 (Lanfear et al., 2017) indicated the best fit substitution model of the completed genome sequences and the protein sequences. The phylogeny of the whole genome sequences of coronavirus was constructed using both the Maximum Likelihood Method and Bayesian Method. RAxML version

8.2.11 (Stamatakis, 2014) with the substitution model GTRGAMMAI was used using 1000 rapid bootstrap replicates. MrBayes version 3.2.6 (Ronquist et al., 2012) with INVGAMMA model was used for the corona virus genomes. Phylogenetic analyses of four different protein sequences were performed by using RAxML-8.2.11 tool. For spike and nucleocapsid proteins, we found PROTGAMMAIWAG and PROTGAMMAIWAG as the best fil model, respectively. Again, PROTGAMMAWAG was the best fit model of evolution for both the membrane and envelope proteins. For the retrieval of the domain sequences of the stated protein sequences, InterPro database (https://www.ebi.ac.uk/interpro/) was utilized. Finally, the Interactive Tree of Life (iTOL; EMBL, Heidelberg, Germany) was used for the visualization of the phylogenetic trees. All the trees were rooted in the midpoint.

### 2.3. Identification of conserved regions as vaccine target

In the present study, reverse vaccinology technique was utilized to model a novel multiepitope subunit vaccine against 2019-nCoV. The scheme in Figure 1 represents the complete methodology that has been adopted to develop the final vaccine construct. Among 496 proteins (available in the NCBI database) from different strains of novel corona virus, four structural proteins, i.e. spike glycoprotein, membrane glycoprotein, envelope protein and nucleocapsid protein, were prioritized for further investigation (Supplementary File 2). After sequence retrieval from NCBI, the sequences were subjected to BLASTp analysis to find out the homologous protein sequences. Multiple sequence alignment was done by using Clustal Omega to identify the conserved regions (Sievers and Higgins, 2014). The topology of each conserved regions were predicted by TMHMM Server v.2.0 (http://www.cbs.dtu.dk/services/TMHMM/), while the antigenicity of the conserved regions was determined by VaxiJen v2.0 (Doytchinova and Flower, 2007a).

**Figure 1:**
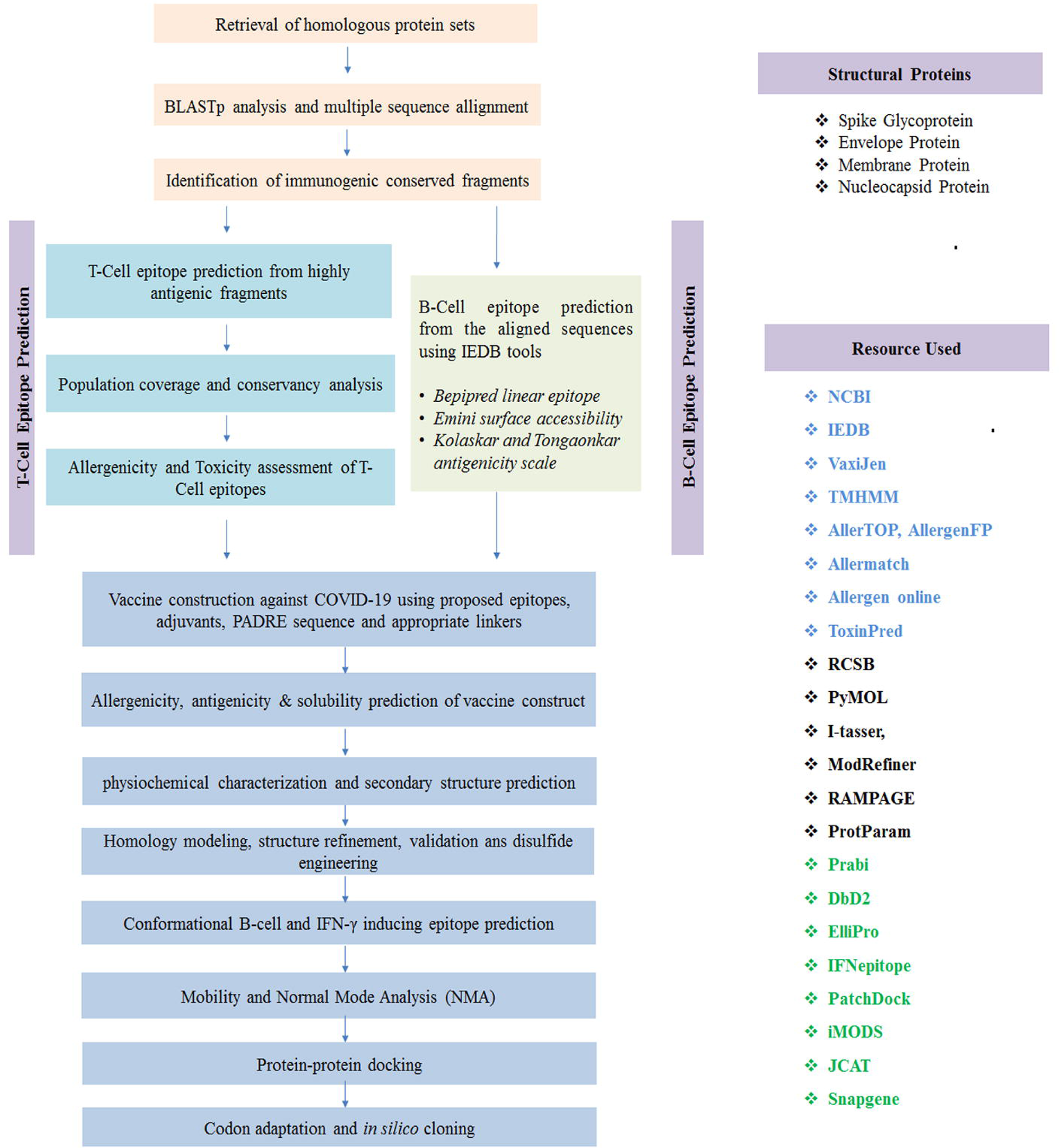
Flow chart summarizing the protocols over multi-epitope subunit vaccine design against SARS-CoV-2 through reverse vaccinology approach.

### 2.4. T-cell epitope prediction, transmembrane topology screening and antigenicity analysis

Only the common fragments were used for T-Cell epitopes enumeration via T-Cell epitope prediction server of IEDB (http://tools.iedb.org/main/tcell/) (Vita et al., 2014). Again, TMHMM server was utilized for the prediction of transmembrane topology of predicted MHC-I and MHC- II binding peptides followed by antigenicity scoring via VaxiJen v2.0 server (Krogh et al., 2001; Doytchinova and Flower, 2007b). The epitopes which have antigenic potency were picked and used for preceding analysis.

### 2.5. Conservancy analysis and toxicity profiling of the predicted epitopes

The level of conservancy scrutinizes the ability of epitope candidates to impart capacious spectrum immunity. Homologous sequence sets of the chosen antigenic proteins were retrieved form the NCBI database by utilizing BLASTp tool. Later, conservancy analysis tool (http://tools.iedb.org/conservancy/) in IEDB was used to demonstrate the conservancy level of the predicted epitopes among different viral starins. The toxicity of non-allergenic epitopes was enumerated by using ToxinPred server (Gupta et al., 2013).

### 2.6. Population coverage and allergenicity pattern of putative epitopes

Among different ethnic societies and geographic spaces, the HLA distribution varies around the world. Population coverage study was conducted by using IEDB population coverage calculation server (Vita et al., 2014). To check the allergenicity of the proposed epitopes, four distinct servers i.e. AllergenFP (Dimitrov et al., 2014), AllerTOP (Dimitrov et al., 2013), Allermatch (Fiers et al., and Allergen Online (http://www.allergenonline.org/) servers were utilized.

### 2.7. Identification of B-Cell epitopes

Three different algorithms i.e. Bepipred Linear Epitope Prediction 2.0 (Jespersen et al., 2017), Emini surface accessibility prediction (Emini et al., 1985) and Kolaskar and Tongaonkar antigenicity scale (Kolaskar and Tongaonkar, 1990) from IEDB predicted the potential B-Cell epitopes within conserved fragments of the chosen viral proteins.

### 2.8. Construction of vaccine molecules and prediction of allergenicity, antigenicity and solubility of the constructs

Top CTL, HTL and B cell epitopes were compiled to design the final vaccine constructs in the study. Each vaccine constructs commenced with an adjuvant followed by top CTL epitopes, HTL epitopes and BCL epitopes respectively. For construction of novel corona vaccine, the chosen adjuvants i.e. L7/L12 ribosomal protein, beta defensin (a 45 mer peptide) and HABA protein (*M. tuberculosis*, accession number: AGV15514.1) were used (Rana and Akhter, 2016). Several linkers such as EAAAK, GGGS, GPGPG and KK in association with PADRE sequence were incorporated to construct fruitful vaccine sequences against COVID-19. The constructed vaccines were then analyzed whether they are non-allergenic by utilizing the following tool named Algpred (Azim et al., 2019). The most potential vaccine among the three constructs was then determined by assessing the antigenicity and solubility of the vaccines via VaxiJen v2.0 (Doytchinova and Flower, 2007b) and Proso II server (Smialowski et al., 2006), respectively.

### 2.9. Physicochemical characterization and secondary structure analysis

ProtParam tool (https://web.expasy.org/protparam/), provided by ExPASy server (Hasan et al., 2019c) was used to functionally characterize (Gasteiger et al., 2003) the vaccine constructs. The studied functional properties were isoelectric pH, molecular weight, aliphatic index, instability index, hydropathicity, estimated half-life, GRAVY values and other physicochemical characteristics. Alpha helix, beta sheet and coil structures of the vaccine constructs were analyzed through GOR4 secondary structure prediction method using Prabi (https://npsa-prabi.ibcp.fr/). In addition, Espript 3.0 (Robert & Gouet, 2014) was also used to predict the secondary structure of the stated protein sequences.

### 2.10. Homology modeling, structure refinement, validation and disulfide engineering

Vaccine 3D model was generated on the basis of percentage similarity between target protein and available template structures from PDB by using I-TASSER (Peng and Xu, 2011). The modeled structures were further refined via FG-MD refinement server. Structure validation was performed by Ramachandran plot assessment in RAMPAGE (Hasan et al., 2019b). By utilizing DbD2 server, probable disulfide bonds were designed for the anticipated vaccine constructs (Craig and Dombkowski, 2013). The value of energy was considered < 2.5, while the chi3 value for the residue screening was chosen between –87 to +97 for the operation (Hasan et al., 2019b).

### 2.11. Conformational B-cell and IFN-α inducing epitopes prediction

The B-cell epitopes of putative vaccine molecules were predicted via ElliPro server (http://tools.iedb.org/ellipro/) with minimum score 0.5 and maximum distance of 7 Å (Ponomarenko et al., 2004). Moreover, IFN- inducing epitopes within the vaccine were predicted using IFNepitope with motif and SVM hybrid detection strategy (Hajighahramani et al., 2017).

### 2.12. Molecular dynamics and normal mode analysis (NMA)

Normal mode analysis (NMA) was performed to predict the stability and large scale mobility of the vaccine protein. The iMod server determined the stability of construct V3 by comparing the essential dynamics to the normal modes of protein (Aalten et al, 1997; Wuthrich et al., 1980). It is a recommended alternative to costly atomistic simulation (Tama and Brooks, 2006; Cui and Bahar, 2007) and shows much quicker and efficient assessments than the typical molecular dynamics (MD) simulations tools (Prabhakar et al., 2016; Awan et al., 2017). The main-chain deformability was also predicted by measuring the efficacy of target molecule to deform at each of its residues. The motion stiffness was represented via eigenvalue, while the covariance matrix and elastic network model was also analyzed.

### 2.13. Protein-protein interaction study

Patchdock server was prioritized for docking between different HLA alleles and the putative vaccine molecules. In addition, the superior construct was also docked with different human immune receptors such as, ACE 3, APN, DPP4 and TLR-8.The 3D structure of these receptors were retrieved from RCSB protein data bank. Detection of highest binding affinity between the putative vaccine molecules and the receptor was experimented based on the lowest interaction energy of the docked structure.

### 2.14. Codon adaptation and in silico cloning

JCAT tool was utilized for codon adaptation in order to fasten the expression of vaccine construct V3 in E. coli strain K12. For this, some restriction enzymes (i.e. BglI and BglII), Rho independent transcription termination and prokaryote ribosome-binding site were put away from the work (Grote et al., 2005). After that, the mRNA sequence of constructed V3 vaccine was ligated within BglI (401) and BglII (2187) restriction site at the C-terminal and N-terminal sites respectively. SnapGene tool was utilized for in silico restriction cloning (Solanki and Tiwari, 2018).

## 3. Results

### 3.1. COVID-19 exhibits close ancestral relation to SARS-CoV-1 and bat-coronavirus

In the phylogenetic analysis, we introduced different coronavirus from three different genera: Alpha coronavirus, Beta coronavirus and Gamma coronavirus. Total 61 species of the corona virus covered 21 sub-genera (Supplementary Table 1 and Figure 2). These 61 species of corona viruses included 7 pathogenic species (Figure 2), which are: COVID-19 or SARS-CoV-2, SARS-CoV-1 virus, MERS virus, HCoV-HKU1, HCoV-OC43, HCoV-NL63, HCoV-229E (Forni et al., 2017; Zhou et al., 2020; Zumla et al., 2016). Among these, the first five species belong to the beta coronavirus genera, while the last two belongs to the alpha genera. Apart from the human coronaviruses, we introduced other coronaviruses which choose different species of bats, whale, turkey, rat, mink, ferret, swine, camel, rabbit, cow and others as host (Supplementary Table-1).

**Table 1:**
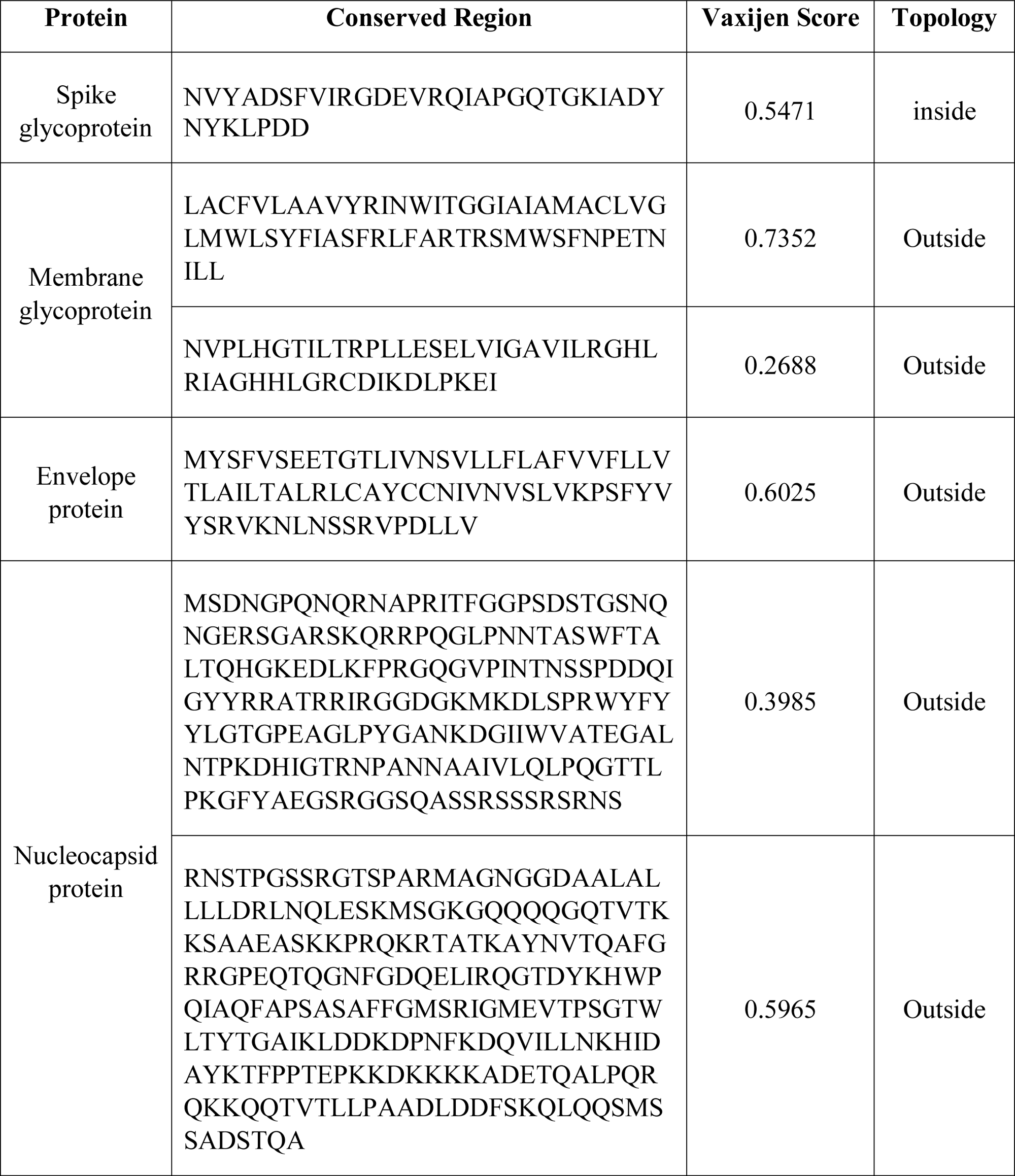
Identified conserved regions among different homologous protein sets.

**Figure 2:**
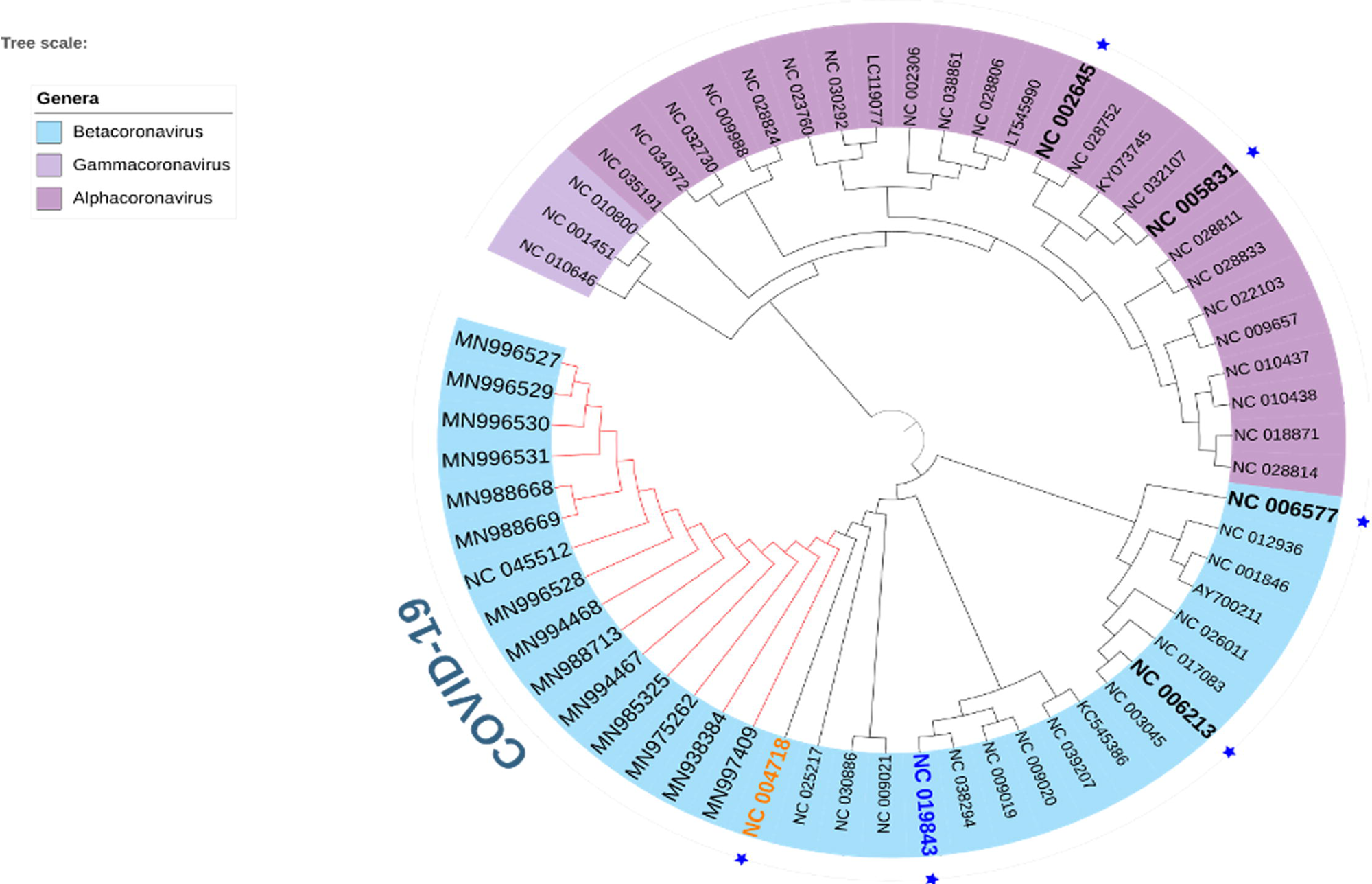
Phylogeny of 61 species of coronaviruses. Seven pathogenic human coronaviruses have been represented by blue star and the IDs have been made bold. COVID-19 clade has been shown with red color. SARS-CoV-1 and MERS virus have been represented by orange and blue colors, respectively. The genera have been represented on the left colored labels.

The phylogeny of these species clearly revealed two broad clades (Figure 2), where first large clade contains Gammacoronavirus and Alphacoronavirus genera, while the other belongs to the Betacoronavirus. Within the Betacoronavirus clade, we found three clear divisions. HCoV- HKU1 and HCoV-OC43 have been placed in the first clade, while in the second clade we found MERS coronavirus. COVID-19 or SARS-COV-2 formed clade with SARS-COV-1 and bat betacoronaviruses (Figure 2), which is consistent with the previous finding (Ceraolo & Giorgi, 2020). Though SARS-CoV-1 belongs to the same sub-genus as the COVID-19, the bat coronaviruses belong to two different sub-genera including Hibecovirus and Nobecovirus (Supplementary File 1).

### 3.2. Evolution of spike proteins based on domain

Domain analysis of spike protein of coronaviruses reveals that they contain mainly one signature domains namely, coronavirus S2 glycoprotein (IPR002552), which is present in all the candidates. All other betacoronavirus contains spike receptor binding protein (IPR018548), coronavirus spike glycoprotein hapted receptor 2 domain (IPR027400) and spike receptor binding domain superfamily (IPR036326). SARS-CoV-1 contains an extra domain, namely spike glycoprotein N-terminal domain (IPR032500), which is also present in some the sub-genera (Embecovirus) of Betacoronavirus, but not in COVID-19. One important finding in our study is that the COVID-19 candidates do not contain the domain spike glycoprotein (IPR042578), which is present in the SARS-CoV-1 (Figure 3). The secondary structure prediction study shows a large numbers of cysteine residues which contribute to the formation of disulfide bonds within the spike protein. Most of them fall within the S1 spike protein, which is 654 amino acid long in SARS-CoV-1, while 672 amino acids long in COVID-19. The RGD motif which is conserved within the COVID-19 is present in the vicinity of the S1 protein. It exists as KGD that clearly demonstrates the mutation over the short time period. Again, the receptor binding domain and receptor binding motif analyses disclose variations within several region between the COVID19 and SARS-CoV-1 (Supplementary File 2). The domain-based phylogenetic analysis reflects two main divisions, where the all the novel betacoronavirus i.e., COVID19 form clade with the SARS-CoV-1; while other betacoronavirus fall in another clade which further divide to give rise different sub-genera. This clearly shows that the COVID-19 exerts specific ancestral connection to the SARS-CoV-1 in terms of spike glycoproteins. Interestingly, our study also revealed close relatedness of both the SARS-CoV-1 and COVID-19 to the bat betacoronavirus that belongs to the Hibecovirus sub-genus. However, in our study, the bat coronaviruses of Nobecovirus sub-genus did not fall into the same clade of novel coronaviruses. The phylogenetic study and MSA also revealed that, the functional portion of the spike glycoprotein domain and spike glycoprotein N-terminal domain might be lost from the COVID-19 during the course of evolution.

**Figure 3:**
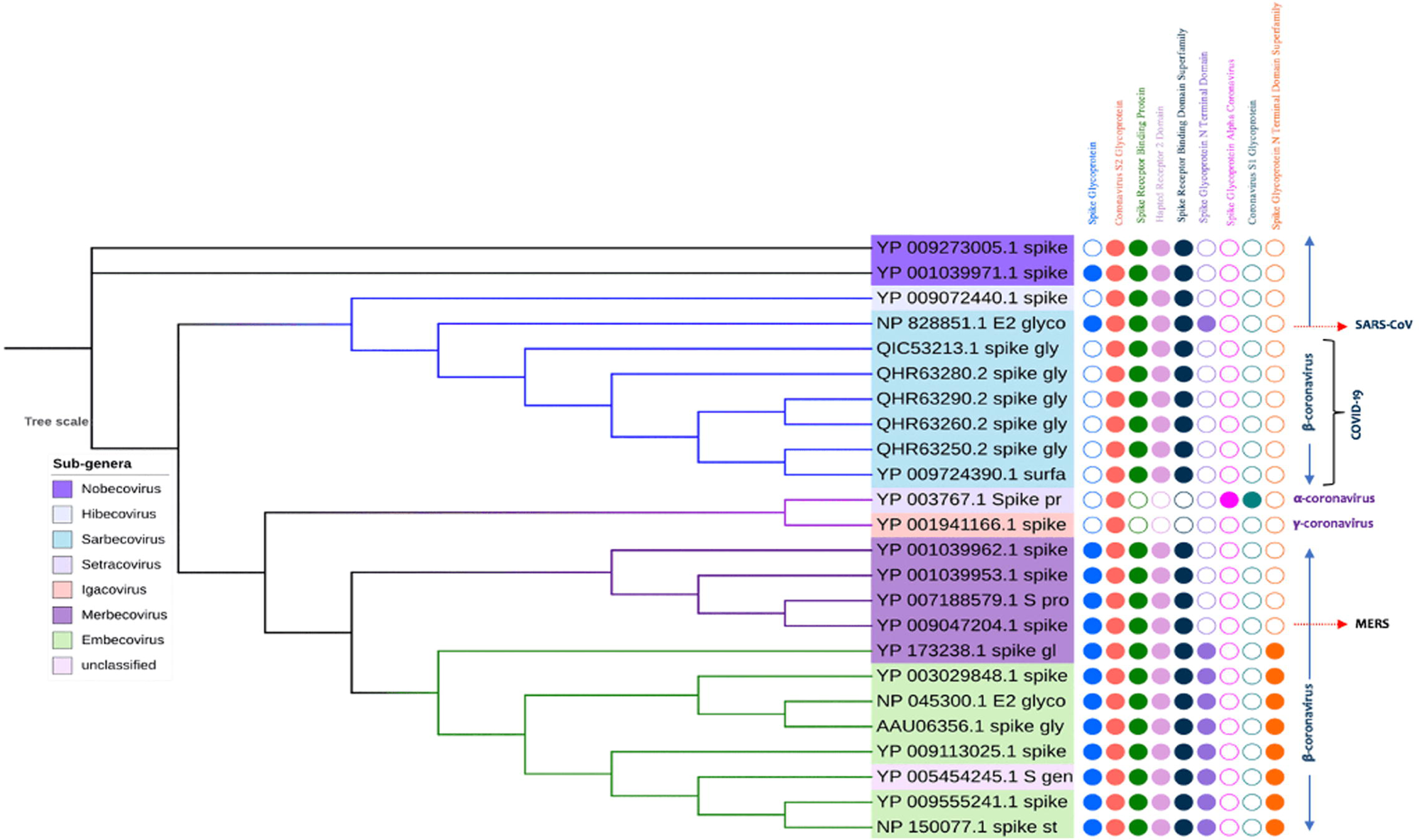
Phylogeny of spike protein of coronavirus. The sub-genera have been labeled in the left table. The filled and unfilled circles show the presence and absence of the domains labeled on the top.

### 3.3. Domain architecture and ancestral state of envelope proteins

The envelope proteins of both Betacoronavirus and Alphacoronavirus contain only one protein domain (IPR003873) namely, Nonstructural protein NS3 or small envelope protein E (NS3/E). This domain is well conserved in coronavirus and also found in murine hepatitis virus. On the other hands, the gamma coronavirus shows the exception, which possess (IPR005296) IBV3C protein domain, which thought to be expressed from the *ORF3C* gene of infectious bronchitis virus (Jia & Naqi, 1997) (Supplementary File 1 and 2). The length of the domains for the COVID-19, SARS-CoV and MARS virus are 75, 76 and 82 amino acids, respectively. While, the length of the gamma coronavirus candidate in our study, which utilizes turkey as host is 99 amino acids.

The average length of the COVID-19 envelope proteins is 75 amino acid long. The NS3/E protein domains spans the whole length of the protein and possess mainly one transmembrane domain, one non-cytoplasmic domain and a cytoplasmic domain. However, some species from the sub-genera of Embecovirus shows two transmembrane domains (Figure 4, Supplementary File 1 and 2). While in our *in-silico* study, we found 2 transmembrane domains in SARS-CoV-1 and 2-3 transmembrane domains in MERS virus, previous experiments proved that both contains only one α-helical transmembrane domains (Nieto-Torres et al., 2011; Surya et al., 2015). Though the computational analysis of CoVID-19 envelope protein secondary structure shows 2 transmembrane domains, our domain analysis shows only one such domain in their structures. Again, the Turkey corona virus also possess the same transmembrane, non-cytoplasmic domain and cytoplasmic domain, with a variation in the orientation. The domain-based phylogeny of the envelope proteins of novel corona viruses reveals close ancestral relationship with the SARS-CoV-1 and bat coronaviruses (Figure 4). In spite to the previous findings, where it was found that the envelope proteins of the MERS virus and SARS-CoV-1 exerted close proximity in terms of secondary structure and functions (Surya et al., 2015). Unlike to earlier finding, we got that gamma corona virus candidate in our study shows close connection with both SARS-CoV-1 and COVID-19 in terms of envelope proteins.

**Figure 4:**
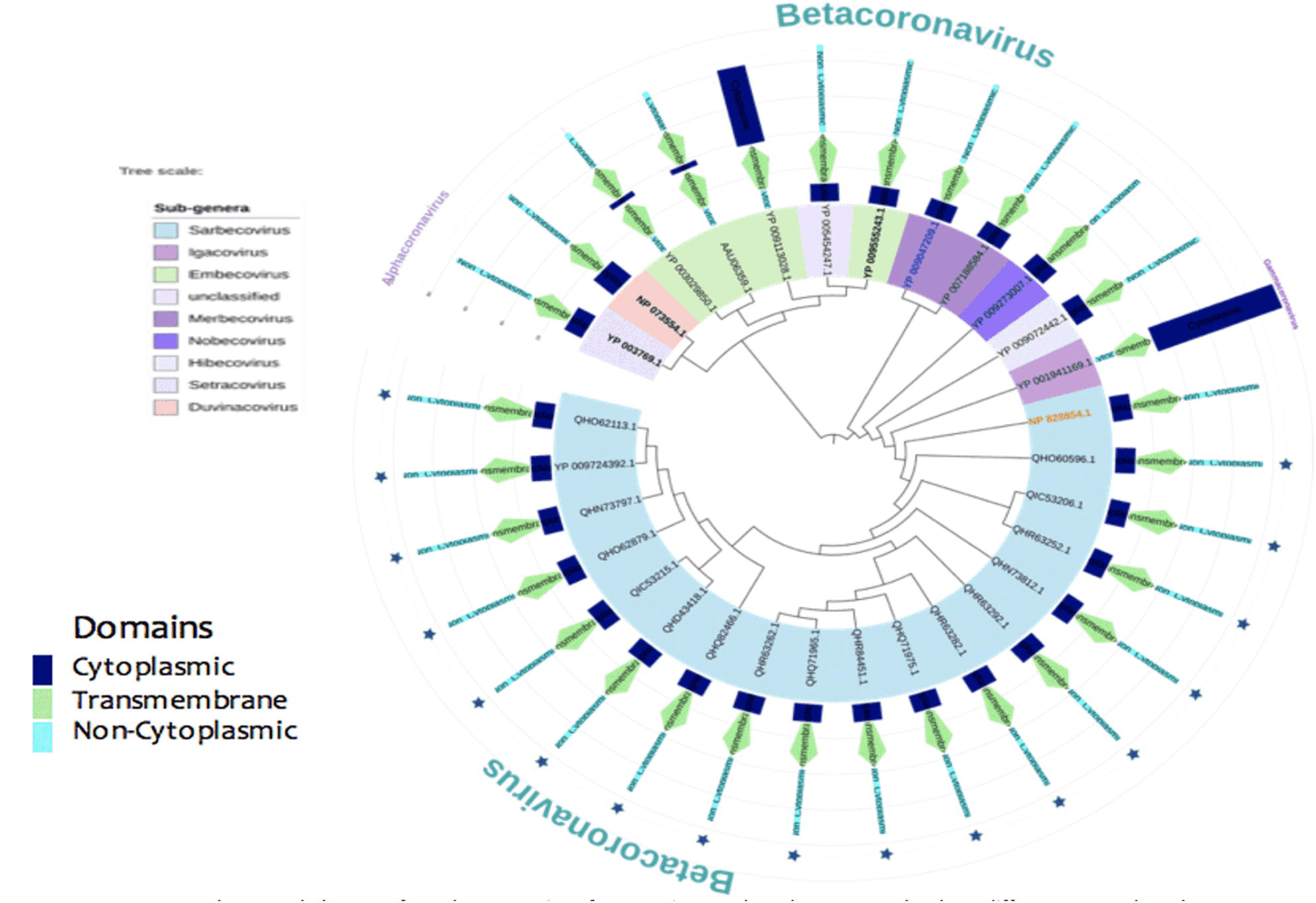
Phylogeny of envelope proteins of coronaviruses. The sub-genera under three different genera have been shown on the left labels. The star signs represent the COVID-19 virus. 5 different pathogenic human corona viruses have been shown in bold form, including the MERS virus (blue) and SARS-CoV-1 (yellow).

### 3.4. Domain architecture and phylogeny of membrane proteins

Membrane proteins of all the coronavirus mainly contain coronavirus M matrix/glycoprotein (IPR002574) domain family. However, the candidates of alpha and gamma coronaviruses contain M matrix/glycoprotein: Alpha coronavirus (IPR042551) and M matrix/glycoprotein: Gamma coronavirus (IPR042550) domains, respectively. The next two domains belong to the M matrix/glycoprotein domain family. The membrane proteins of coronaviruses ranges from 221 to 230 amino acid long. Computational analysis of secondary structures shows some variations of COVID-19 with SARS-CoV-1 and MERS virus (Supplementary File 3). COVID-19 possess alpha helical structure in their structure, while the other two completely devoid of this structure. Again, both SARS-CoV-1 and MARS contain parallel beta sheets, while it is completely absent in the novel corona virus (Supplementary File 3). The phylogenetic analysis of the membrane supports that the novel coronavirus is closely connected to the SARS-CoV-1 virus membrane protein. As well as it produced connections with bat coronaviruses of Hibecovirus and Nobecovirus sub-genus (Figure 5).

**Figure 5:**
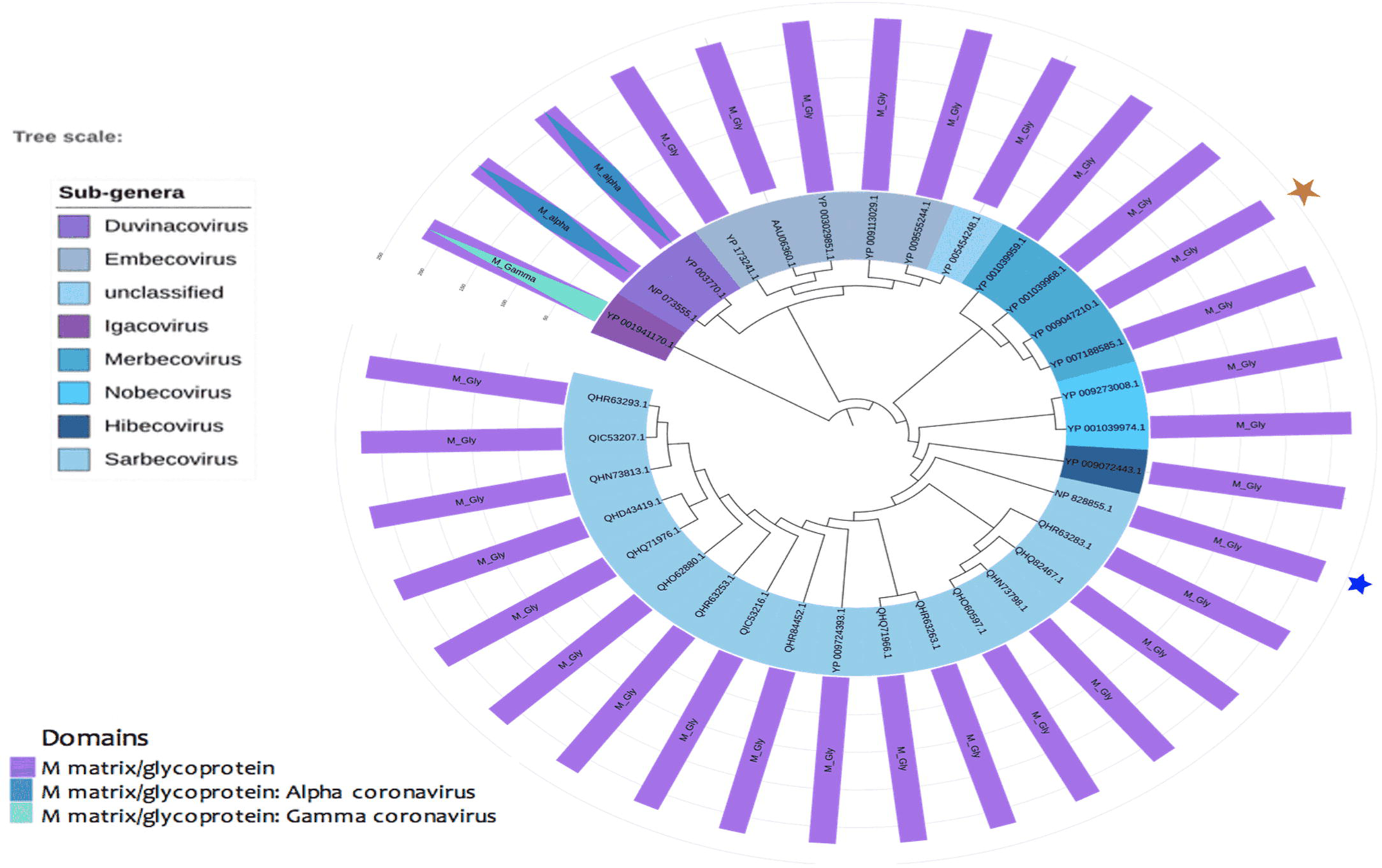
Phylogeny of membrane proteins of coronavirus. The sub-genera under three different genera have been shown on the left labels. The blue and orange star represent the SARS-CoV1 and MERS virus, respectively.

### 3.5. Domain-based phylogeny of nucleocapsid proteins

The length of nucleocapsid proteins of betacoronavirus genus ranges from 410 to 450 amino acids. Three signature domains are mainly present in the nucleocapsid proteins, which are: Coronavirus Nucleocapsid protein (IPR001218), Nucleocapsid Proteins C-terminal (IPR037179) and Nucleocapsid Proteins N-terminal (IPR037195). However, in our experiment, we didn’t find these domains in HCoV-HKU1 (Figure 6); this virus belongs to Embecovirus sub-genus and contains only Coronavirus Nucleocapsid I (IPR004876) domain. The candidates of alpha and gamma coronavirus have their special domains. According to our domain-based phylogeny study of nucleocapsid proteins, we found the close approximation of the COVID-19 with the SARS-CoV-1, which is consistent with the findings of phylogeny of whole genome. Strikingly, unlike spike and membrane proteins, the closest homologs of COVID-19 are not only the SARS-COV-1 and bat coronaviruses (Sub-genus: Hibecovirus and Nobecovirus), but it also includes the MERS viruses and other proteins which is from the Merbecovirus sub-genus.

**Figure 6:**
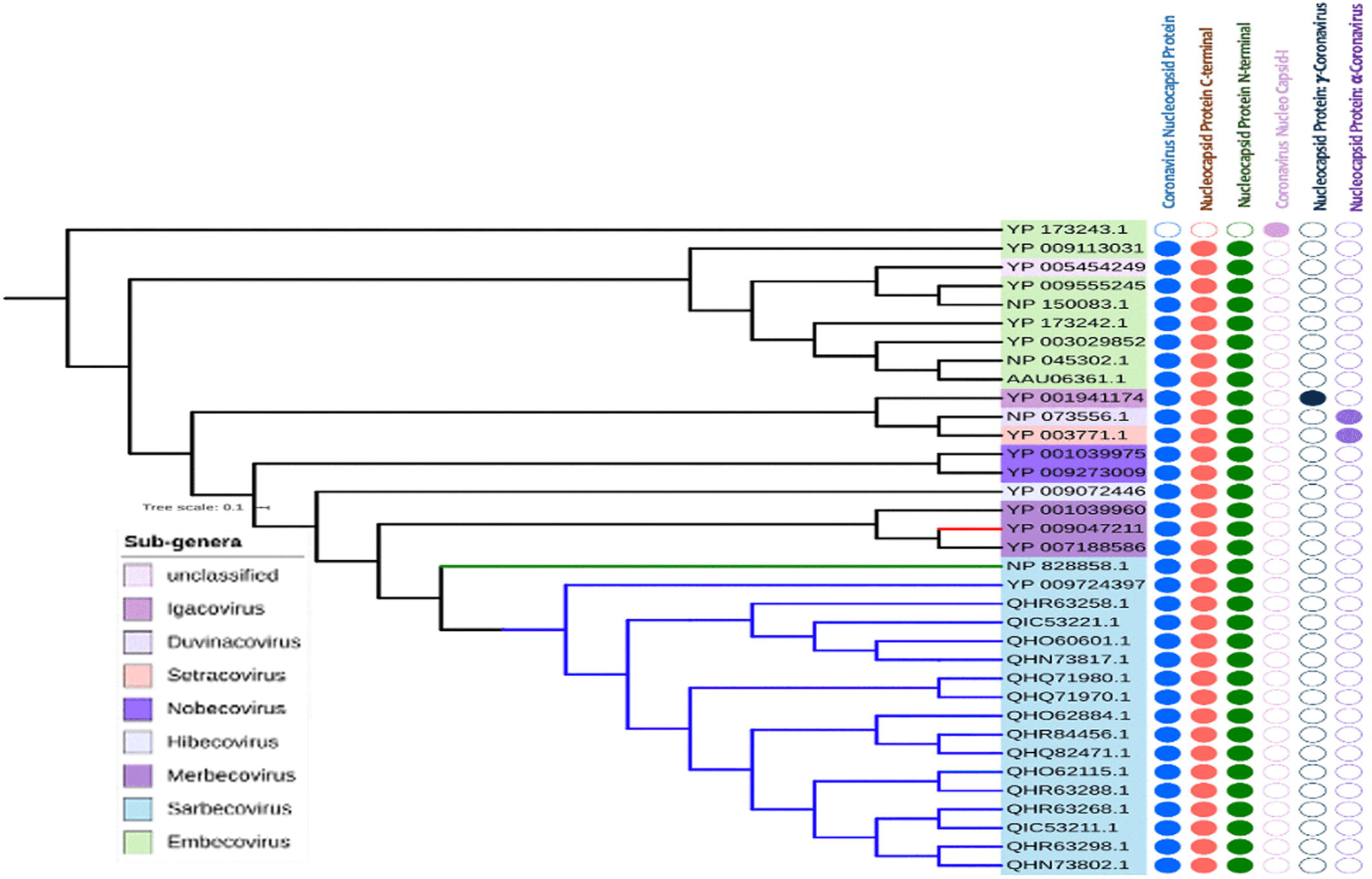
Phylogeny of nucleocapsid proteins of coronavirus. The sub-genera under three different genera have been shown on the left labels. The filled and unfilled circles show the presence and absence of the domains labeled on the top. COVID-19, SARS-CoV1 and MERS viruses are clades are labeled with blue, green and red colors, respectively.

### 3.6. Identification of conserved regions as vaccine target

A total 31, 24, 29 and 29 sequences of spike glycoprotein, membrane glycoprotein, envelope protein and nucleocapsid protein were retrieved from the NCBI database belonging to different strains of SARS-CoV-2. Following by BLASTp analysis and Multiple Sequence Alignment, two conserved regions were detected for membrane glycoprotein and nucleocapsid, while single fragments were identified for both spike glycoprotein and envelope protein (Table 1). Results showed that, all the conserved sequences except one from membrane glycoprotein met the criteria of default threshold standard in VaxiJen. Again, transmembrane topology scrutinizing showed that among the immunogenic conserved sequences from the corresponding proteins except spike glycoprotein met the criteria of desired exomembrane characteristics (Table 1).

### 3.7. T-cell epitope prediction, transmembrane topology screening and antigenicity analysis

A plethora of immunogenic epitopes were generated from the conserved sequences that were able to bind with most noteworthy number of HLA cells (Supplementary Table 1, Supplementary Table 2, Supplementary Table 3 and Supplementary Table 4). Top epitopes with exomembrane characteristics were ranked for each individual protein after investigating their antigenicity score and transmembrane topology (Table 2).

**Table 2:**
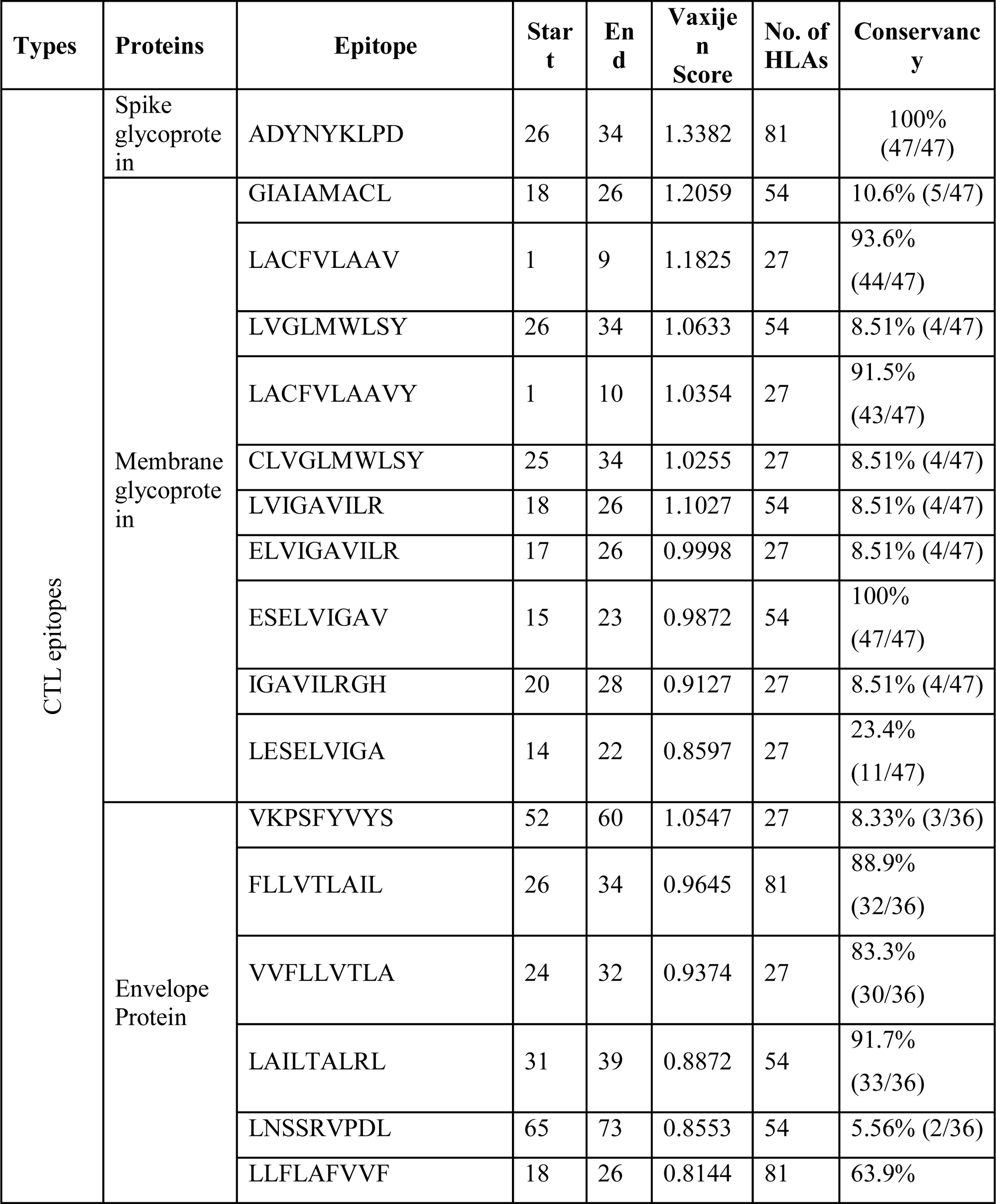
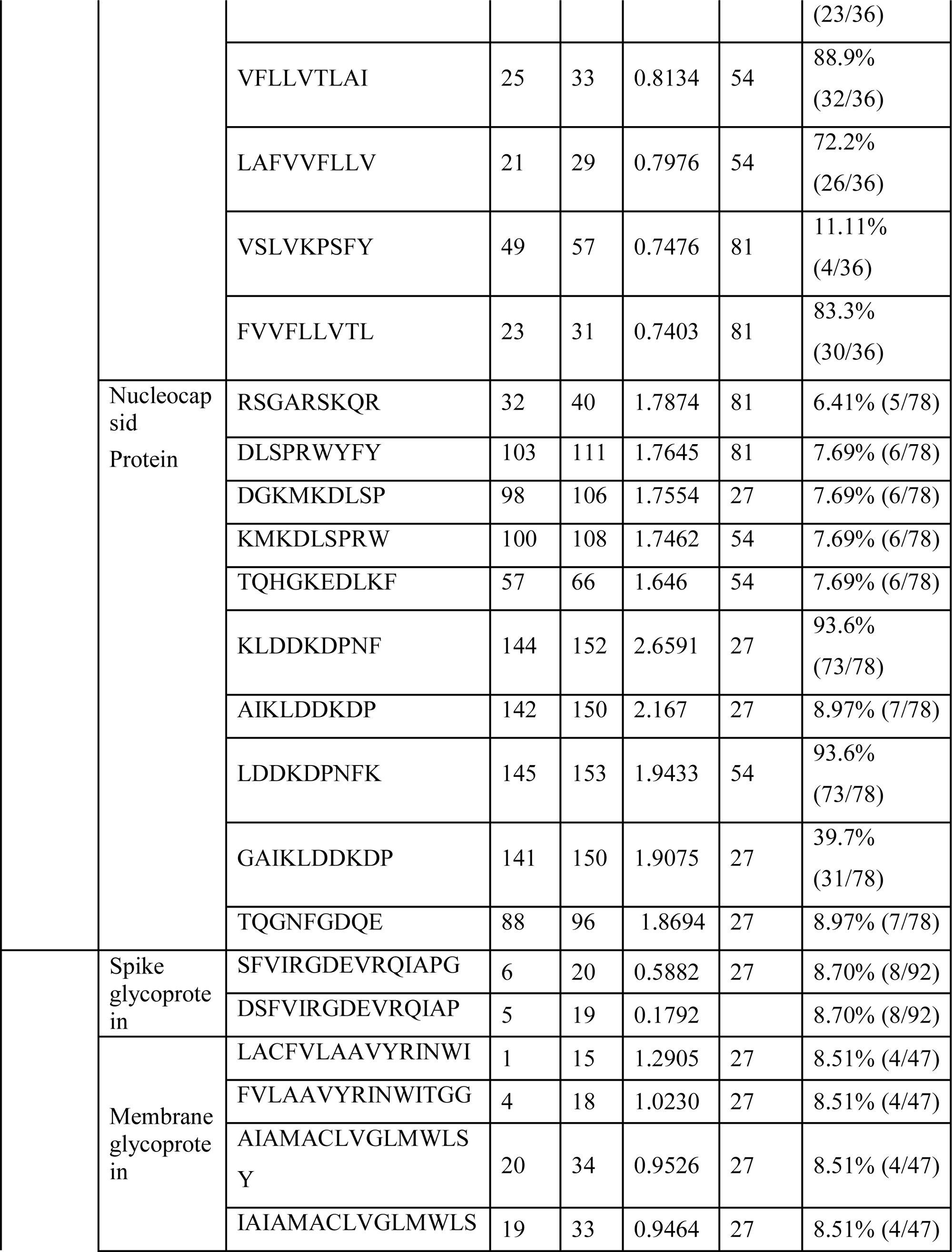
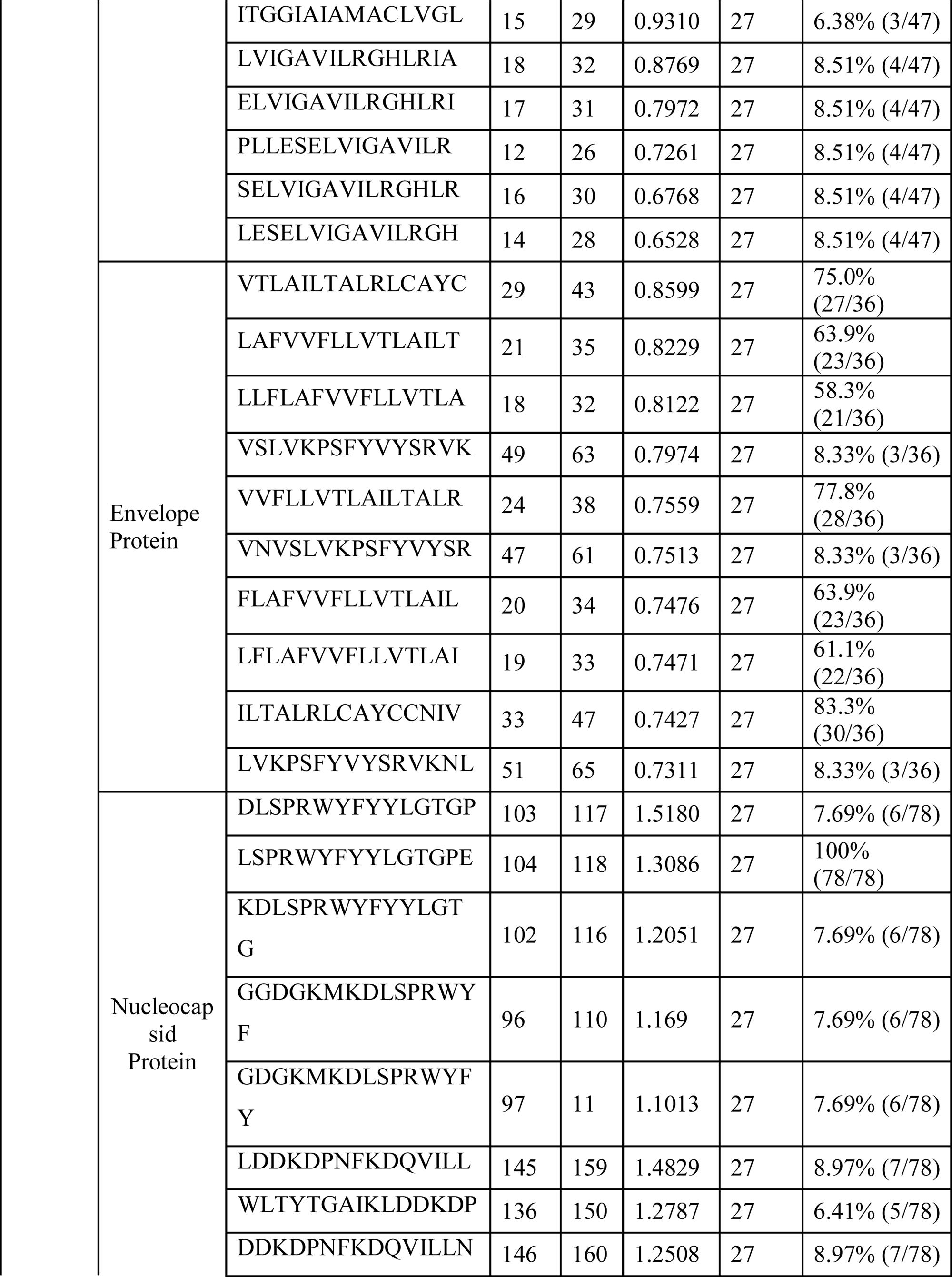
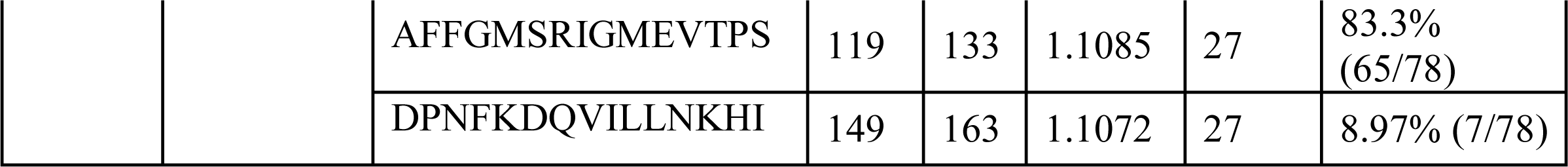
Predicted T-cell (CTL & HTL) epitopes of Spike glycoprotein, membrane glycoprotein, envelope protein and nucleocapsid protein

**Table 3:**
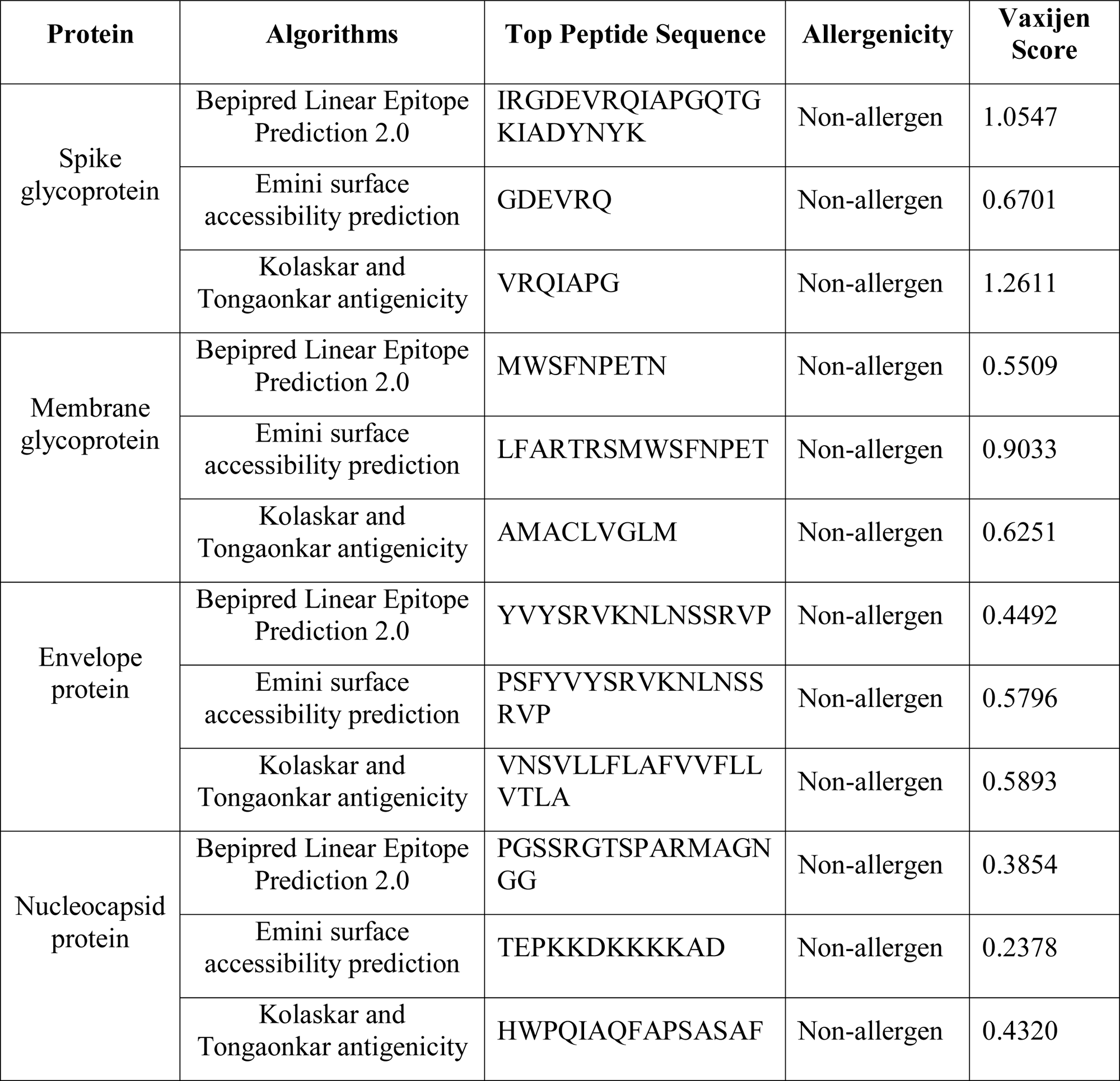
Allergenicity and antigenicity assessment of predicted B-cell epitopes.

**Table 4:**
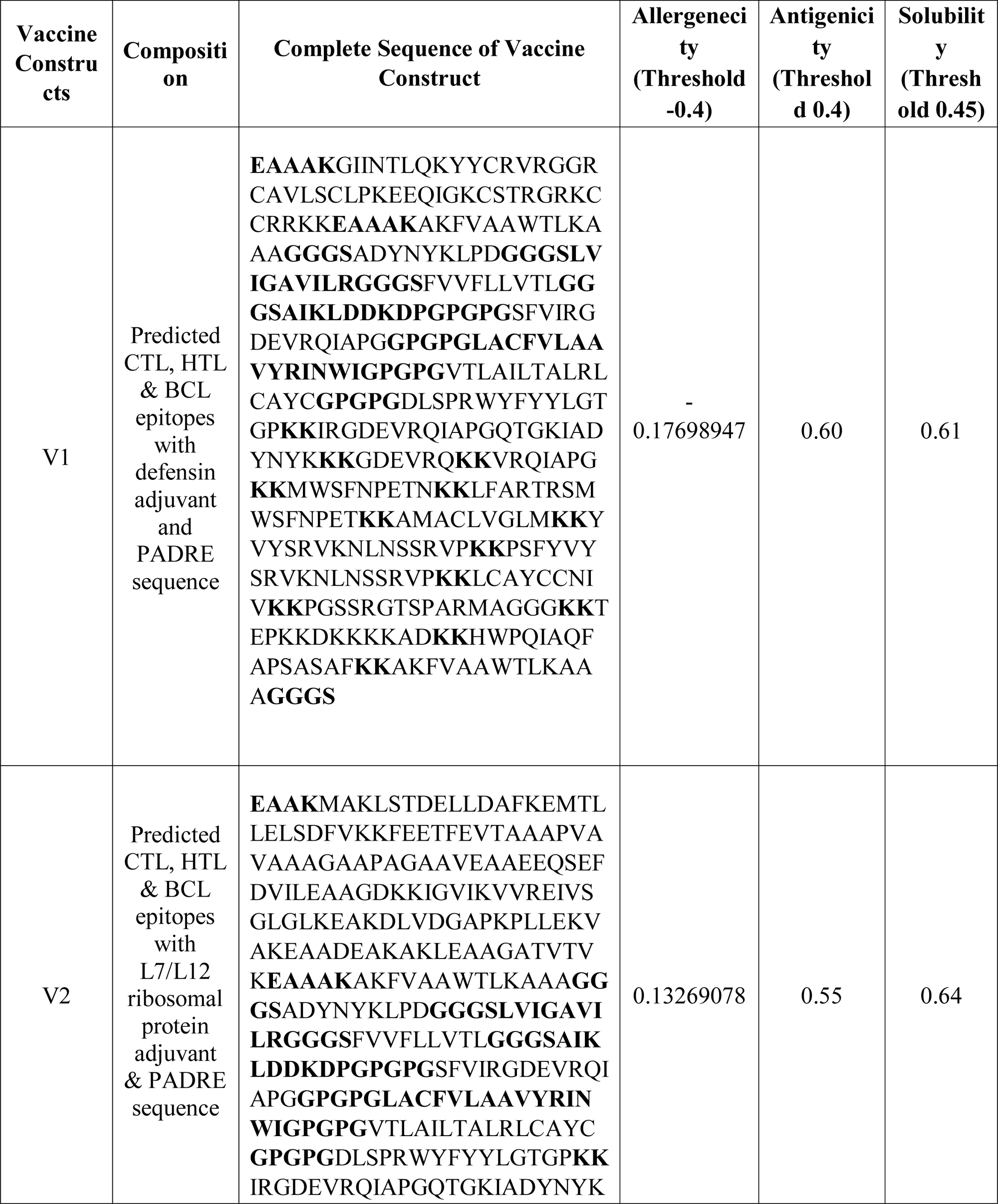
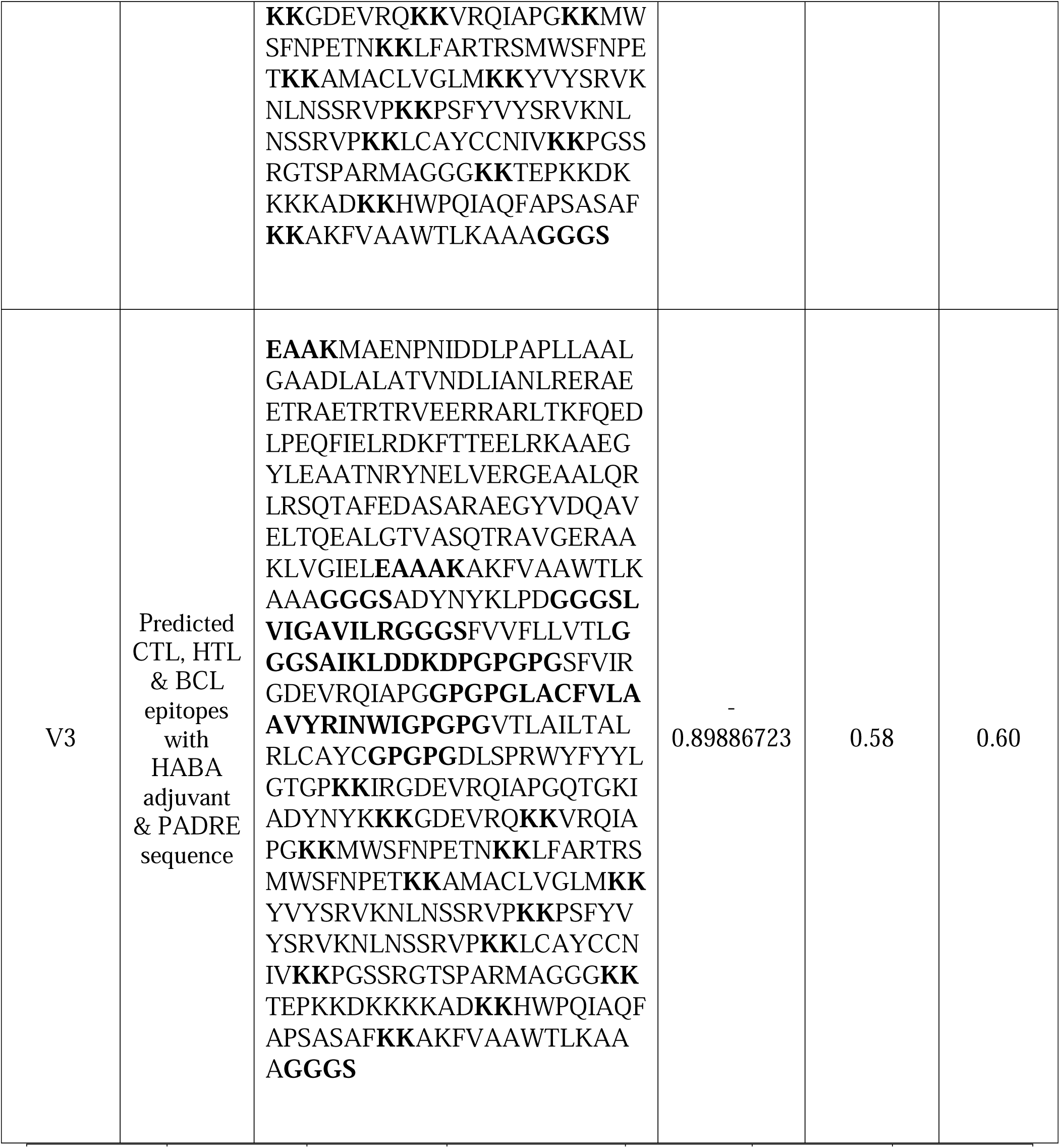
Allergenicity, antigenicity and solubility prediction of designed vaccine constructs.

### 3.8. Conservancy analysis, toxicity profiling, population coverage and allergenicity pattern of the predicted epitopes

Epitopes from each protein showed high level of conservancy up to 100% (Table 2). ToxinPred server predicted the relative toxicity of each epitope which indicated that the top epitopes were non-toxin in nature (Supplementary Table 5). Population coverage of four structures proteins were also done for the predicted CTL and HTL epitopes. From the screening, results showed that population of the various geographic regions could be covered by the predicted T-cell epitopes (Figure 7). Finally, the allergenic epitopes were excluded from the list based on the evaluation of four allergenicity prediction server (Supplementary Table 5).

**Table 5:**
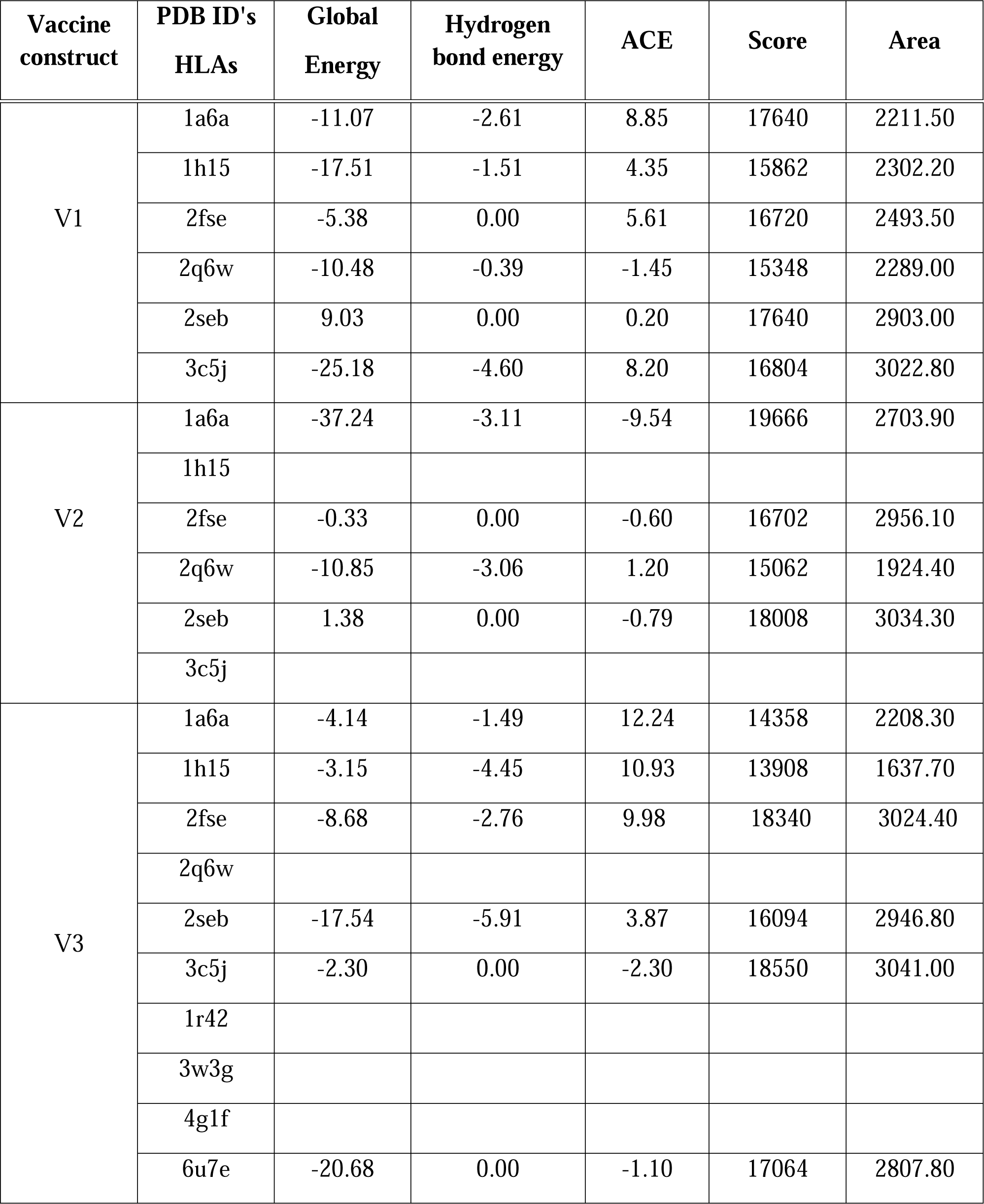
Docking score of vaccine construct V3 with different HLA alleles and human immune receptors

**Figure 7:**
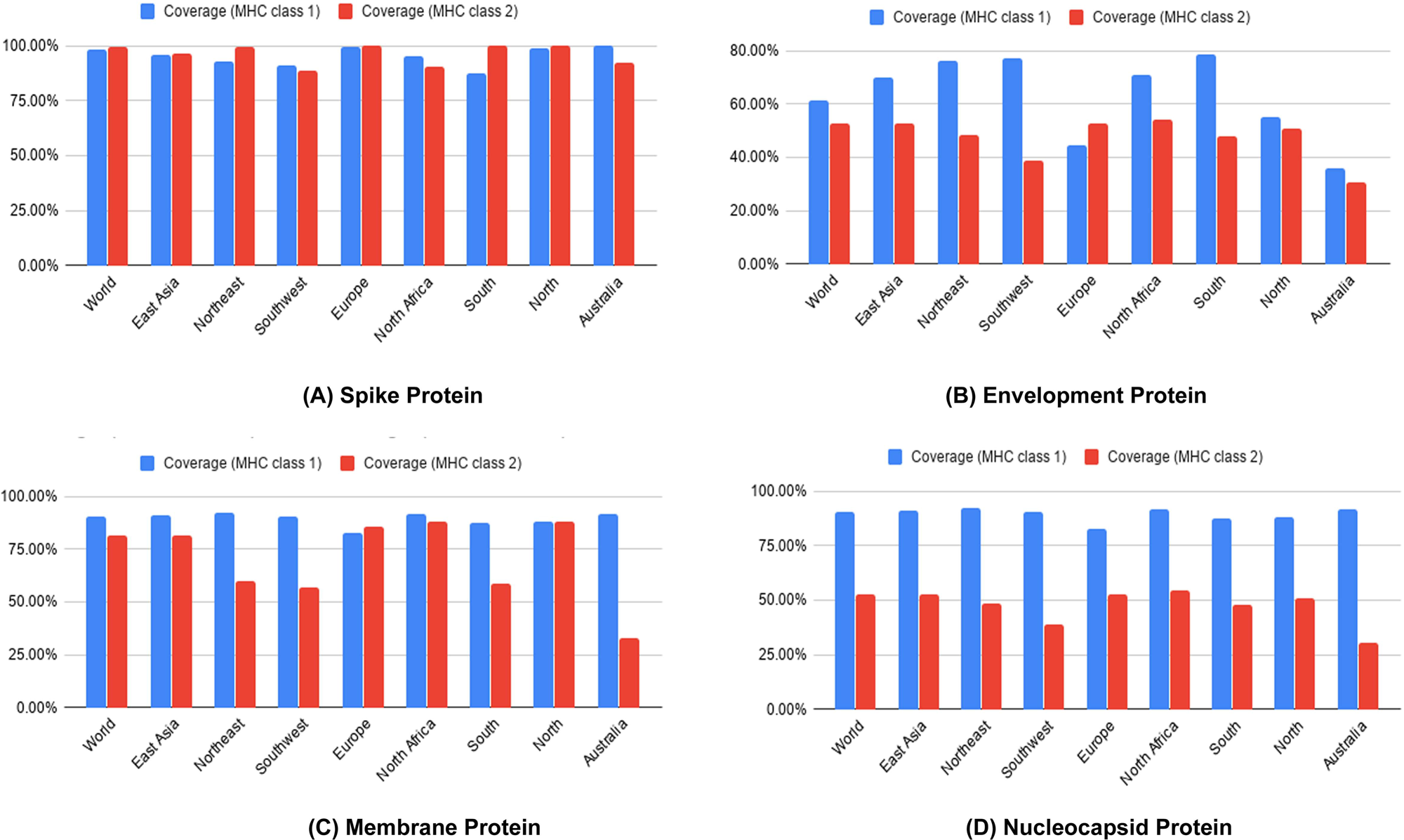
Population coverage analysis of spike protein (A), envelope protein (B), membrane protein (C) and nucleocapsid proteins (D).

### 3.9. Identification of B cell epitopes

Top B-cell epitopes were predicted for Spike glycoprotein, membrane glycoprotein, envelope protein and nucleocapsid protein using 3 distinct algorithms (i.e. Bepipred Linear Epitope prediction, Emini Surface Accessibility, Kolaskar & Tongaonkar Antigenicity prediction) from IEDB. Epitopes were also allowed to analyze their vaxijen scoring and allergenicity (Table 3).

### 3.10 Construction of vaccine molecules and prediction of allergenicity, antigenicity and solubility of the constructs

Three putative vaccine molecules (i.e. V1, V2 and V3) were constructed, each comprising a protein adjuvant, eight T-cell epitopes, twelve B-cell epitopes and respective linkers (Supplementary Table 6). PADRE sequence was included to extend the efficacy and potency of the constructed vaccine. The putative vaccine constructs, V1, V2 and V3 were 397, 481 and 510 residues long respectively. However, allergenicity score of V3 (−0.89886723) revealed that it was superior among the three constructs in terms safety and efficacy. V3 also had a solubility score (0.60) (Figure 8E) and antigenicity (0.58) over threshold value (Table 4).

**Figure 8:**
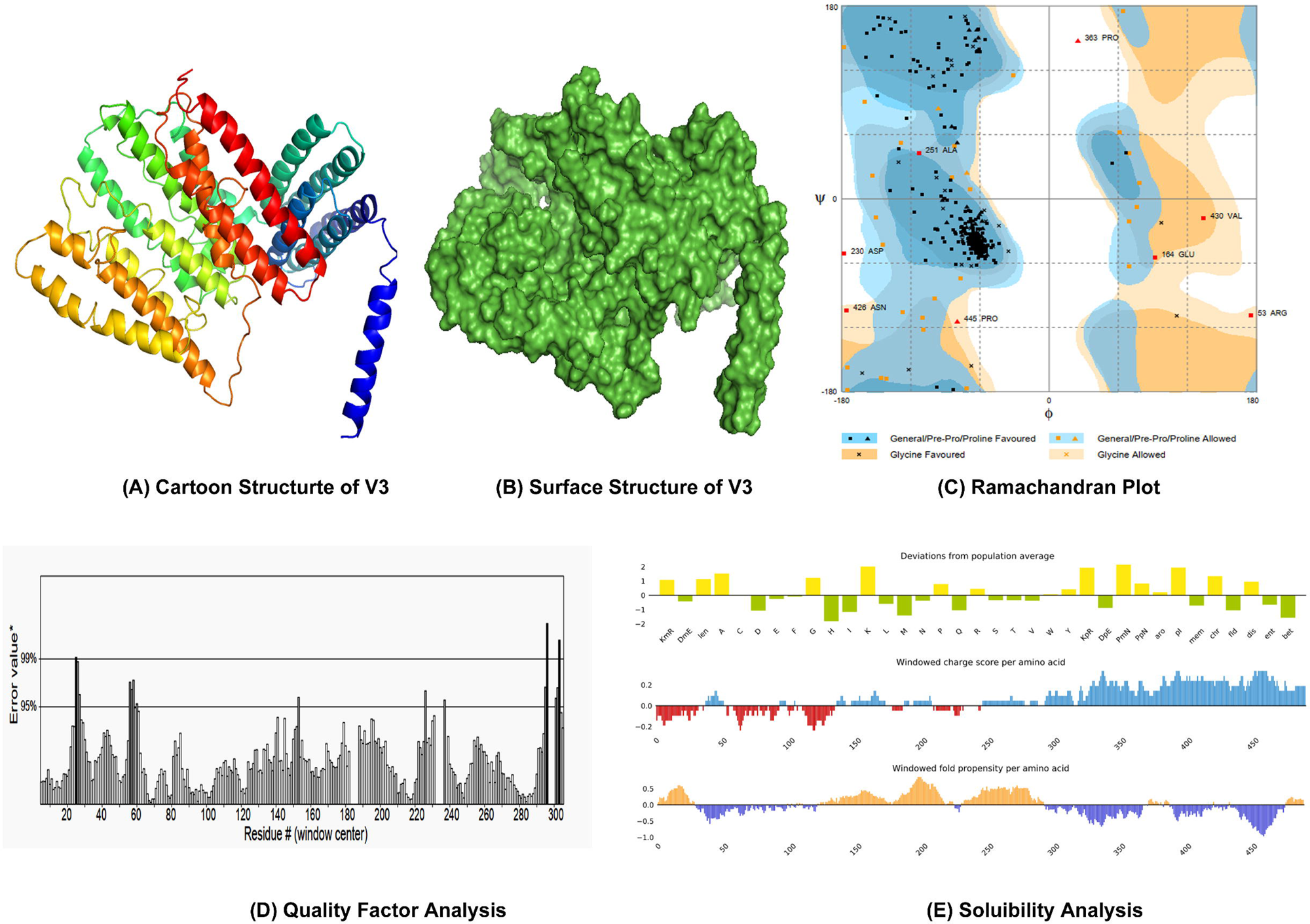
Homology modelling, structure validation and solubility prediction of construct V3 (A: Cartoon structure, B: Surface structure, C: Ramachandran Plot analysis, D: Quality factor analysis, E: Solubility analysis)

### 3.11 Physicochemical characterization and secondary structure analysis of the construct

ProtParam tool was employed to analyze the physicochemical properties of V3. Molecular weight of 3 was scored as 55.181 kDa. The extinction coefficient of V3 was calculated as 63830 at 0.1% absorption. It had been found that the protein would have net negative charge which was higher than the recommended pI 9.81. Aliphatic index and GRAVY value were found 77.80 and −0.383 respectively, which could express the thermostability and hydrophilic status of the V3 vaccine construct. Around sixty minutes *in vitro* half-life stability in mammalian reticulocytes was predicted for V3. The computed instability index (II) 36.98 classified the protein as a stable one. In contrast, Secondary structure of V3 exhibited to have 46.47% alpha helix, 15.00% sheet and 38.63% coil structure (Supplementary Figure 1).

### 3.12. Homology modeling, structure refinement, validation and disulfide engineering

Tertiary structure of the putative vaccine construct V3 was generated using I-TASSER server (Figure 8A and 8B). The server used 10 best templates with highest significant (measured via Z-score) from the LOMETS threading program to model the 3D structure. After refinement, Ramachandran plot analysis revealed that 92.7% and 5.7% residues were in the favored and allowed regions respectively, while only 8 residues (1.6%) occupied in the outlier region (Figure 8C). The overall quality factor determined by ERRAT server was 91.56% (Figure 8D). 3D modelled structure of V1 and V2 are shown in Supplementary Figure 2. DbD2 server recognized 33 pairs of amino acid residue with the potentiality to create disulfide bond between them. After analysis chi3 and B-factor parameter of residue pairs on the basis of energy, only 2 pairs (PRO 277-THR 329 and LEU 425- CYS 435) met the criteria for disulfide bond formation which were changed with cysteine (Supplementary Figure 3).

### 3.13. Conformational B-cell and IFN- inducing epitopes prediction

Ellipro server predicted a total 6 conformational B-cell epitopes from the 3D structure of the construct V3. Epitopes No. 1 were considered as the broadest conformational B cell epitopes with 25 amino acid residues (Figure 9 and Supplementary Table 7). Results also revealed that predicted linear epitopes from 76-101, 131-143 and 48-55 were included in the conformational B-cell epitopes. Moreover, the sequence of the final vaccine was scanned for 15-mer IFN-inducing epitopes. Results showed that there were 292 positive IFN- inducing epitopes from which 20 had a score ≥5 (Supplementary Table 8). Residues of 194-209 regions (GGGSLVIGAVILRGG) in the vaccine showed highest score of 17. (Supplementary Table 8).

**Figure 9:**
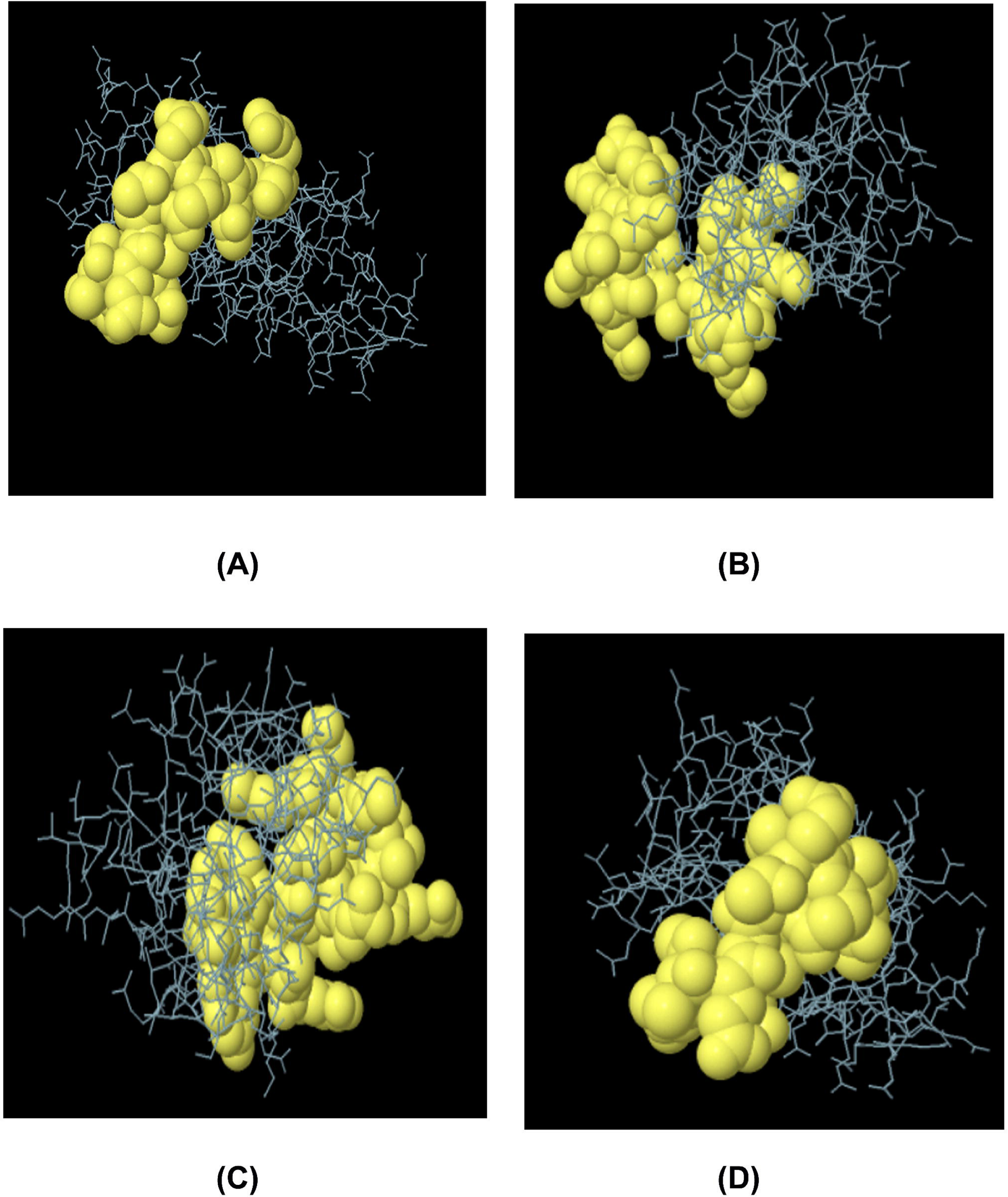
Predicted conformational epitopes (A and B) and linear epitopes (C and D) within construct V3.

### 3.14. Molecular dynamics and normal mode analysis

Stability of the vaccine construct V3 was investigated through mobility analysis (Figure 10A and 10B), B-factor, eigenvalue & deformability analysis, covariance map and recommended elastic network model. Results revealed that the placements of hinges in the chain was insignificant (Figure 10C) and the B-factor column gave an averaged RMS (Figure 10D). The estimated higher eigenvalue 6.341333e-06 (Figure 10E) indicated low chance of deformation of vaccine protein V3. The correlation matrix and elasticity of the construct have been shown in Figure 10G and Figure 10H, respectively.

**Figure 10:**
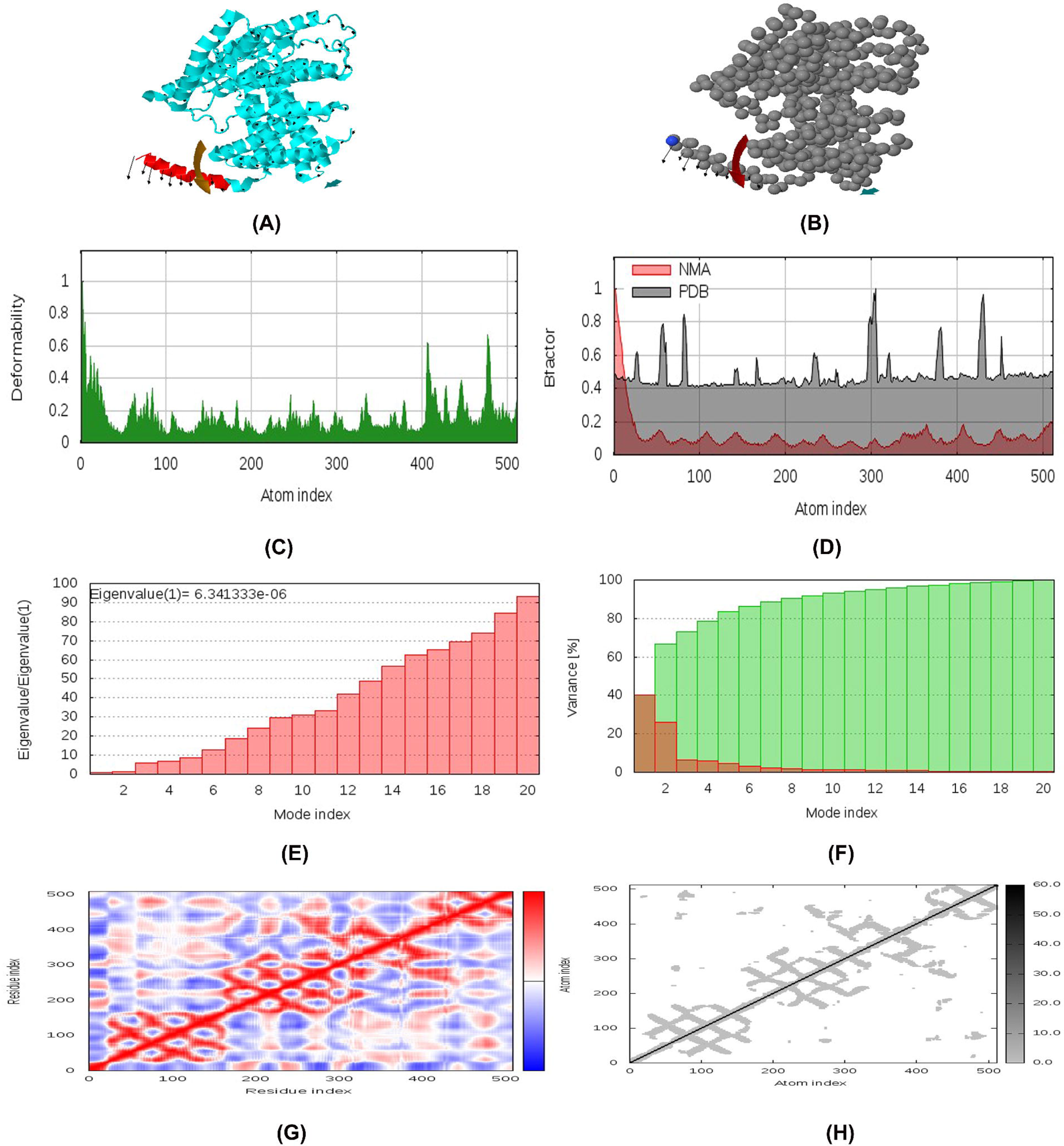
Normal Mode Analysis (NMA) of vaccine protein V3. The directions of each residues are given by arrows and the length of the line represented the degree of mobility in the 3D model (A and B). The main-chain deformability derived from high deformability regions indicated by hinges in the chain which are negligible (C). The experimental B-factor was taken from the corresponding PDB field and calculated from NMA (D). The eigenvalue represents the motion stiffness and directly related to the energy required to deform the structure (E). The variance associated to each normal mode is inversely related to the eigenvalue. Coloured bars show the individual (red) and cummulative (green) variances (F). The covariance matrix indicates coupling between pairs of residues, where they may be associated with correlated, uncorrelated or anti-correlated motions, indicated by red, white and blue colours respectively (G). The elastic network model identifies the pairs of atoms connected via springs. Each dot in the diagram is coloured based on extent of stiffness between the corresponding pair of atoms. The darker the greys, the stiffer the springs (H).

### 3.15. Protein-protein docking

The structural interaction between HLA alleles and the designed vaccines were investigated by molecular docking approach. The server detected the complexed structure by focusing on complementarity score, ACE (Atomic Contact Energy) and estimated interface area of the compound (Table 5). The molecular affinity between the putative vaccine molecules V3 and several immune receptors were also experimented. The result showed that construct V3 interacted with each receptor with significantly lower binding energy (Figure 11).

**Figure 11:**
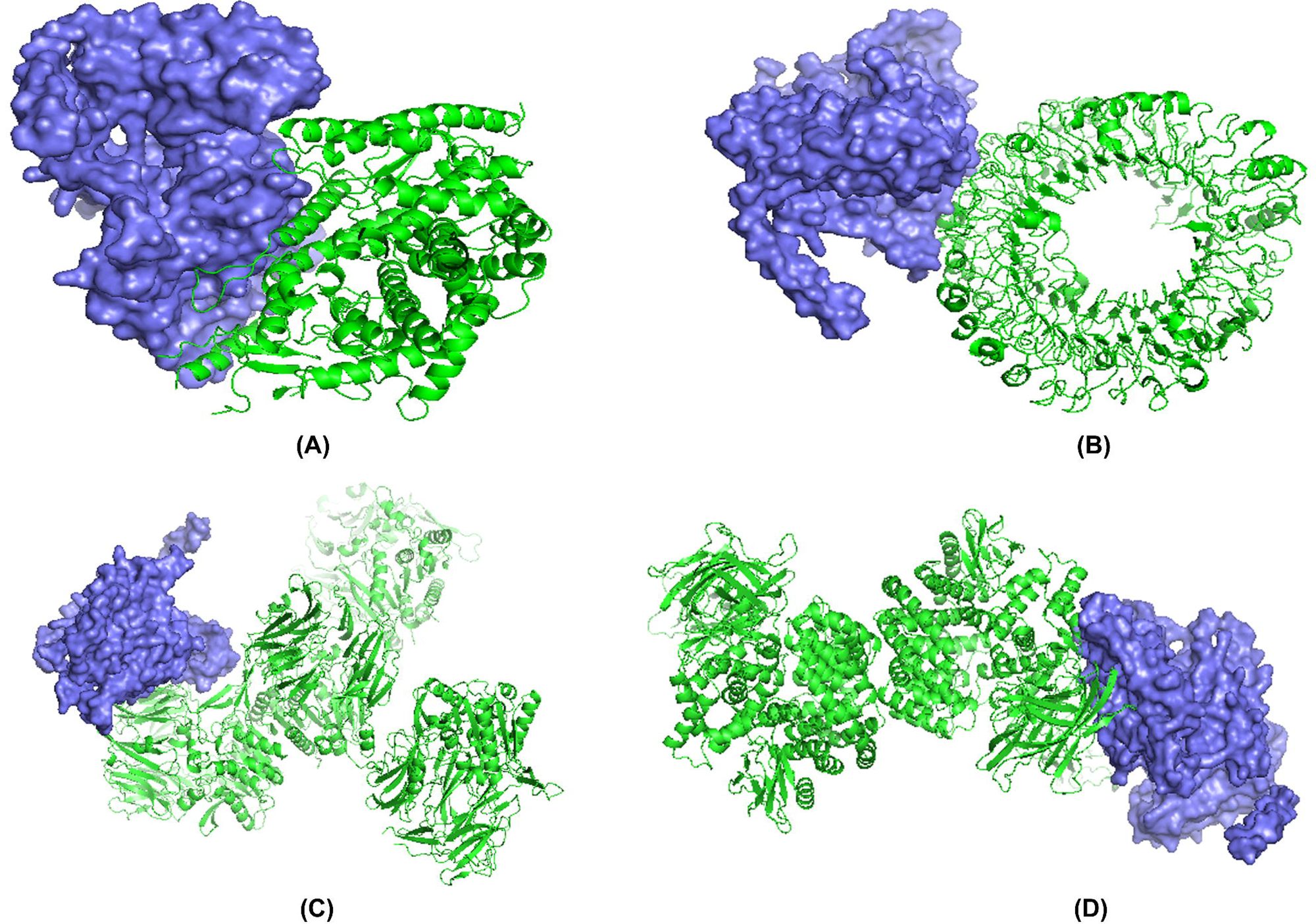
Docked complex of vaccine construct V3 with human ACE 2 (A), TLR- (B), DPP4 (C) and APN (D).

### 3.16. Codon adaptation and in silico cloning

The Codon Adaptation Index (CAI) and GC content for the predicted codons of the putative vaccine constructs V1 were demonstrated as 1.0 and 51.56% respectively. An insert of 1542 bp was found which lacked the restriction sites for BglI and BglII, thus providing comfort zone for cloning. The codons were inserted into pET28a(+) vector alongside two restriction sites (BglI and BglII) and a clone of 5125 base pair was generated (Figure 12).

**Figure 12:**
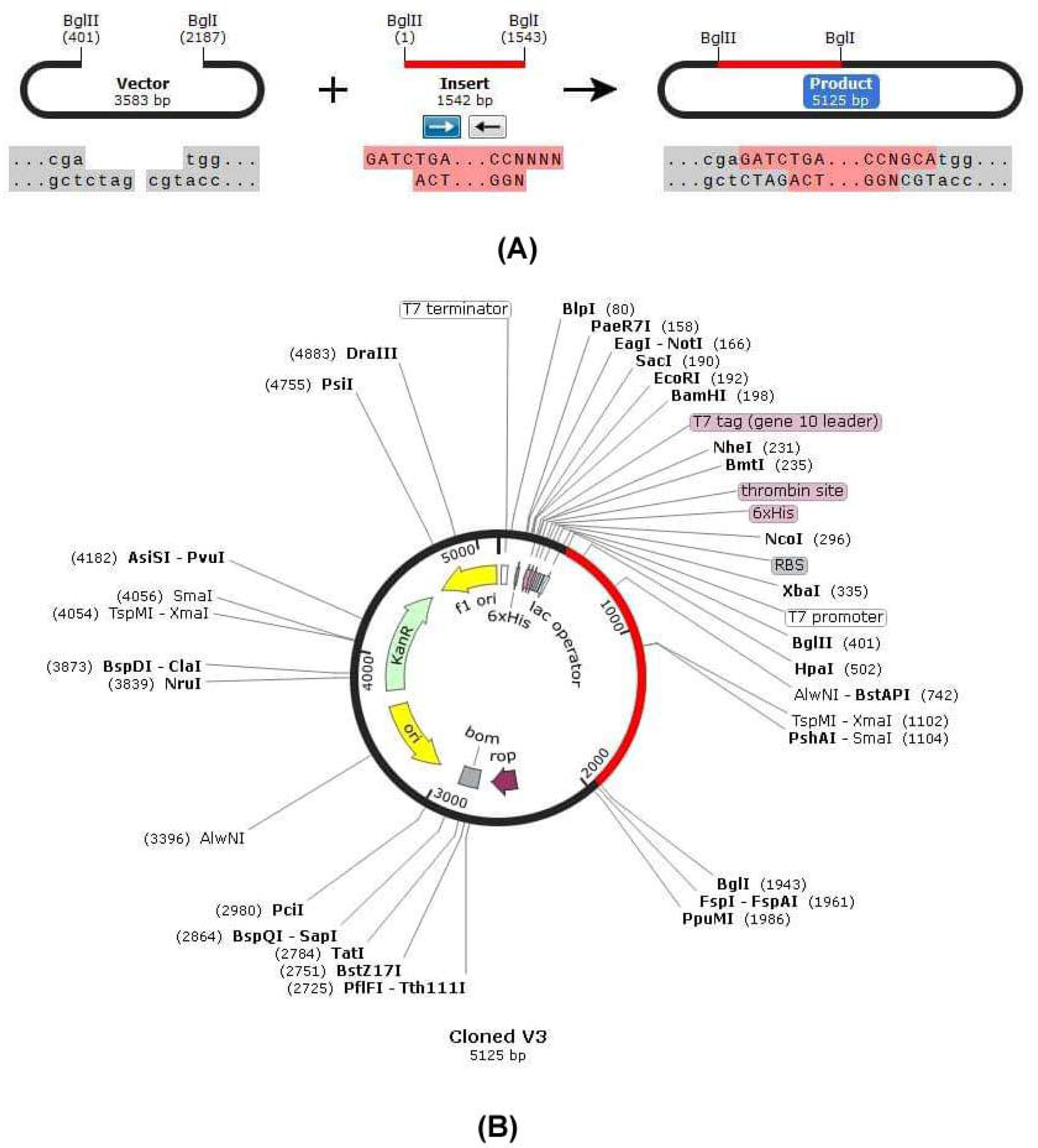
*In silico* restriction cloning of the gene sequence of final vaccine construct V3 into pET28a(+) expression vector (A: Restriction digestion of the vector pET28a(+) and construct V3 with BglII and BglI, B: Inserted desired fragment (V3 Construct) between BglII (401) and BglI (1943) indicated in red color.

## 4. Discussion

In December 2019, a new coronavirus prevalence flourished in Wuhan, China, causing clutter among the medical community, as well as to the rest of the world (Sun et al., 2020). The new species has been renamed as 2019-nCoV or, SARS-CoV-2, already causing considerable number infections and deaths in China, Italy, Spain, Iran, USA and to a growing degree throughout the world. The major outbreak and spread of SARS-CoV-2 in 2020 forced the scientific community to make considerable investment and research activity for developing a vaccine against the pathogen. However, owing to high infectivity and pathogenicity, the culture of SARS-CoV-2 needs biosafety level 3 conditions, which may obstructed the rapid development of any vaccine or therapeutics. It had been found that about 35 companies and academic institutions are engaged in such works (Spinney et al., 2020, Ziady et al., 2020). Among the potential SARS-CoV-2 vaccines in the pipeline, four have nucleic acid based designs, four involve non-replicating viruses or protein constructs, two contain live attenuated virus and one involves a viral vector (Pang et al., 2020), while only one, called mRNA-1273 (developed by NIAID collaboration with Moderna, Inc.), has confirmed to start phase-1 trial (NIH, 2020). However, in this study we emphasized on a different approaches by prioritizing the advantages of different genome and proteome database using the immunoinformatic approach. Computational vaccine predictions were adopted by the researchers to design vaccines against both MERS-CoV (Sudhakar et al., 2013; Fernando et al., 2013) and SARS-CoV-1 (Yang et al., 2003; Oany et al., 2014), targeting the outer membrane or functional proteins (Sharmin and Islam, 2014). Several *in silico* strategies have also been employed to predict potential T cell and B cell epitopes against SARS-CoV-2, either emphasizing on spike glycoprotein or envelope proteins (Behbahani, 2020; Rasheed et al., 2020). None of the studies, however, focused on other structural proteins. Moreover, random genetic changes and mutations in the protein sequences (Yin, 2020) may obstruct the development of effective vaccines and therapeutics against human coronavirus in the future. Hence, the present study was employed to identify the similarity and divergence among the close relatives of the target pathogen and develop a novel chimeric recombinant vaccine considering all major structural proteins i.e. spike glycoprotein, membrane glycoprotein, envelope protein and nucleocapsid protein simultaneously.

The topology of the phylogenetic trees of the whole genome and the stated four proteins sequences from different species of coronaviruses reveal that SARS-CoV-1 and bat coronaviruses are the closest homologs of the novel coronaviruses. Our results infer a significant level of similarities within the COVID-19 and SARS-CoV-1 which was also aligned with the previous findings (Jaimes et al., 2020; Sun et al., 2020; Wu, 2020a). The sequence similarities between the SARS-CoV, bat coronaviruses and the COVID-19 from the reported studies (Hu et al., 2018; Wu et al., 2020; Wu, 2020b) suggests that those are distantly related, in spite those are capable of infecting the humans and therefore possess the adaptive convergent evolution. Interestingly, the COVID-19 envelope proteins form clade with the Turkey coronavirus which belongs to Gamma coronavirus genus. So, in terms of envelope proteins, the envelope gene of turkey coronavirus might contribute to the convergence process, which need further analysis. In addition, from the domain-based phylogeny of nucleocapsid proteins, it can be deduced that this protein might have originated in bats and was transmitted to camels and then later on choose human as the potential host. Overall, the COVID-19 might go through complex adaptation strategies in order to be transmitted into the human via different animals.

The homologous protein sets for four structural proteins of Coronavirus were sorted to identify conserved regions through BLASTp analysis and MSA. Only the conserved sequences were utilized to identify potential B-cell and T-cell epitopes for each individual protein (Table 1). Thus, our constructs are expected to stimulate a broad-spectrum immunity in host upon administration. Cytotoxic CD8+T lymphocytes (CTL) play a crucial role to control the spread of pathogens by recognizing and killing diseased cells or by means of antiviral cytokine secretion (Garcia et al., 1999). Thus, T cell epitope-based vaccination is a unique process to confer defensive response against pathogenic candidates (Shrestha, 2004). Approximately 800 MHC-I peptides (CTL epitopes) and 600 MHC-II peptides (HTL epitopes) were predicted via IEDB server, from which we screened the top ones through analyzing the antigenicity score, transmembrane topology, conservancy level and other important physiochemical parameters employing a number of bioinformatics tools (Table 2). The top 10 epitopes from each protein was further assessed by investigating the toxicity profile and allergenicity pattern. Different servers rely on different parameters to predict the allergenic nature of small peptides. Therefore, we used 4 distinct servers for such assessment and the epitopes predicted as non-allergen at least via 3 servers were retained for further analysis (Supplementary Table 5). Vaccine initiates the generation of effective antibodies that are usually produced by B cells and plays effector functions by targeting specifically to a foreign particles (Cooper & Nemerow, 1984). The potential B cell epitopes were generated by three different algorithms (Bepipred linear epitope prediction 2.0, Kolaskar and Tongaonkar antigenicity prediction and Emini surface accessibility prediction) from IEDB database (Table 3).

Suitable linkers and adjuvants were used to combine top finalized epitopes from each protein that led to develop a multi epitope vaccine molecules (Supplementary Table 6). As PADRE sequence was usually recommended to lessen the polymorphism of HLA molecules in the population (Ghaffari-Nazari et al., 2015), it was also considered to construct the final vaccine molecule. Here, adjuvants would enhance the immunogenicity of the vaccine constructs and appropriate separation of epitopes in the host environment would be ensured by the linker (Yang et al., 2015). Allergenicity, physiochemical properties, antigenicity and three-dimensional structure of vaccine constructs were characterized, and it had been concluded that V3 was superior to V1 and V2 vaccine constr. The final construct also occupied by several interferon-α producing epitopes (Supplementary Table 8). The vaccine protein (V3) was subjected to disulfide engineering to enhance its stability. Analysis of the normal modes in internal coordinates by iMODS was employed to investigate the collective motion of vaccine molecules (Lopez-Blanco et al., 2014). Negligible chance of deformability at molecular level was analyzed for the putative vaccine construct V3, thereby strengthening our prediction. Moreover, molecular docking was investigated to analyze the molecular affinity of the vaccine with different HLA molecules i.e. DRB1*0101, DRB5*0101, DRB3*0202, DRB1*0401, DRB3*0101 and DRB1*0301 (Table 5). It had been reported that a specific receptor-binding domain of CoV spike protein usually recognizes its host receptor ACE2 (angiotensin-converting enzyme 2) (Li. et al., 2003; Li, 2015). Previous studies also identified dipeptidyl peptidase 4 (DPP4) as a functional receptor for human coronavirus (Raj et al., 2013). Therefore, we performed another docking study prioritizing these immune receptors to strengthen our prediction (Figure 11). Results showed that the designed construct bound with the selected receptors with minimum binding energy which was biologically significant. Finally, *in-silico* restriction cloning was adopted to check the suitability of construct V3 for entry into pET28a (+) vector and expression in *E. coli* strain K12 (Figure 12).

Traditional ways to vaccine development are time consuming and laborious. Moreover, the result may not be always as expected or fruitful (Stratton et al., 2003; Hasan et al., 2019). *In silico* prediction and prescreening methods, on the contrary, offer some advantages while saving time and cost for production. Therefore, the present study may aid in the development of preventive strategies and novel vaccines to combat infections caused by 2019-nCoV. However, further wet lab trials involving model organism needs to be experimented for validating our findings.

## Acknowledgements

Authors would like to acknowledge the Department of Biochemistry and Chemistry, Department of Microbial Biotechnology and Department of Pharmaceuticals and Industrial Biotechnology of Sylhet Agricultural University for the technical support of the project.

## Funding information

This research did not receive any specific grant from funding agencies in the public, commercial, or not-for-profit sectors.

## Conflict of interest

Authors declare that they have no conflict of interests.

## Supplementary Figures

**Supplementary Figure 1:** Secondary structure prediction of constructed vaccine protein V3.

**Supplementary Figure 2:** 3D modelled structure of vaccine protein V1 and V2.

**Supplementary Figure 3:** Disulfide engineering of vaccine protein V3 (A: Initial form, B: Mutant form).

## Supplementary Tables

**Supplementary Table 1**: Predicted CTL and HTL epitopes of spike glycoprotein.

**Supplementary Table 2:** Predicted CTL and HTL epitopes of membrane.

**Supplementary Table 3:** Predicted CTL and HTL epitopes of envelope protein.

**Supplementary Table 4:** Predicted CTL and HTL epitopes of nucleocapsid protein.

**Supplementary Table 5:** Allergenicity pattern and toxicity profile of top T cell epitopes.

**Supplementary Table 6**: Proposed CTL and HTL epitopes for vaccine construction.

**Supplementary Table 7:** Predicted conformational epitopes within construct V3.

**Supplementary Table 8:** Predicted IFN alpha producing epitopes in the vaccine construct V3.

## Supplementary Files

**Supplementary File 1:** NCBI IDs of the complete genome, spike glycoprotein, envelope protein, membrane protein and nucleocapsid protein of coronavirus with Genera and Sub- Genera.

**Supplementary File 2:** Retrieved protein sequences of major structural proteins of COVID-19.

**Supplementary File 3:** Secondary structure and domain analysis of spike glycoprotein, envelope protein, membrane protein and nucleocapsid proteins.

## Notes

### Competing Interest Statement

The authors have declared no competing interest.

