## Supplementary Table 1 for "Genome based Evolutionary study of SARS-CoV-2 towards the Prediction of Epitope Based Chimeric Vaccine"

**Supplementary Table 1**: Predicted CTL and HTL epitopes of spike glycoprotein.

| **Type** | **Epitope** | **start** | **end** | **length** | | **No. of HLAs** | | **topology** | **Vexigen score** |
| --- | --- | --- | --- | --- | --- | --- | --- | --- | --- |
| CTL epitopes | NVYADSFVIR | 1 | 10 | 10 | | 27 | | inside | -0.3210 |
|  | FVIRGDEVR | 7 | 15 | 9 | | 81 | | inside | 0.0083 |
|  | KIADYNYKL | 24 | 32 | 9 | | 81 | | inside | 2.0574 |
|  | RQIAPGQTG | 15 | 23 | 9 | | 81 | | inside | 1.7890 |
|  | QTGKIADYNY | 21 | 30 | 10 | | 27 | | inside | 1.5116 |
|  | QIAPGQTGK | 15 | 24 | 10 | | 81 | | inside | 1.8297 |
|  | GQTGKIADY | 20 | 28 | 9 | | 81 | | inside | 1.4019 |
|  | VYADSFVIR | 2 | 10 | | 9 | | 81 | inside | -0.2347 |
|  | TGKIADYNY | 22 | 30 | | 9 | | 81 | inside | 1.5305 |
|  | NVYADSFVI | 1 | 9 | | 9 | | 54 | inside | -0.5617 |
|  | IAPGQTGKI | 17 | 25 | | 9 | | 81 | Inside | 1.6527 |
|  | APGQTGKIA | 18 | 26 | | 9 | | 81 | Inside | 1.2002 |
|  | SFVIRGDEVR | 6 | 15 | | 10 | | 27 | Inside | 0.1752 |
|  | RGDEVRQIA | 10 | 18 | | 9 | | 81 | Inside | -0.2528 |
|  | IRGDEVRQIA | 9 | 18 | | 10 | | 27 | Inside | -0.364 |
|  | GQTGKIADY | 20 | 28 | | 9 | | 81 | Inside | 1.4019 |
|  | ADSFVIRGDE | 4 | 13 | | 10 | | 27 | Inside | 0.0298 |
|  | ADYNYKLPD | 26 | 34 | | 9 | | 81 | Outside | 1.3382 |
|  | VIRGDEVRQI | 8 | 17 | | 10 | | 27 | Inside | -0.3175 |
|  | DSFVIRGDEV | 5 | 14 | | 10 | | 27 | Inside | 0.3245 |
|  | EVRQIAPGQ | 13 | 21 | | 9 | | 81 | Inside | 1.1205 |
|  | PGQTGKIADY | 19 | 28 | | 10 | | 27 | Inside | 1.5583 |
|  | YADSFVIRG | 3 | 11 | | 9 | | 81 | Inside | 0.2816 |
|  | GKIADYNYKL | 23 | 32 | | 10 | | 27 | inside | 1.6079 |
|  | DEVRQIAPGQ | 12 | 21 | | 10 | | 27 | inside | 0.5825 |
|  | DSFVIRGDE | 5 | 13 | | 9 | | 81 | inside | 0.1674 |
|  | EVRQIAPGQT | 13 | 22 | | 10 | | 27 | inside | 1.0655 |
|  | DYNYKLPDD | 27 | 35 | | 9 | | 54 | inside | 0.5946 |
|  | IADYNYKLPD | 25 | 34 | | 10 | | 27 | inside | 1.2589 |
|  | GDEVRQIAPG | 11 | 20 | | 10 | | 27 | inside | 0.8536 |
|  | VRQIAPGQTG | 14 | 23 | | 10 | | 27 | inside | 1.3856 |
| HTL epitopes | ADSFVIRGDEVRQIA | 4 | 18 | | 15 | | 27 | inside | -0.0663 |
|  | YADSFVIRGDEVRQI | 3 | 17 | | 15 | | 27 | inside | -0.0699 |
|  | DSFVIRGDEVRQIAP | 5 | 9 | | 15 | | 27 | outside | 0.1792 |
|  | SFVIRGDEVRQIAPG | 6 | 20 | | 15 | | 27 | outside | 0.5882 |
|  | FVIRGDEVRQIAPGQ | 7 | 21 | | 15 | | 27 | inside | 0.4940 |
|  | VIRGDEVRQIAPGQT | 8 | 22 | | 15 | | 27 | inside | 0.3090 |
|  | IRGDEVRQIAPGQTG | 9 | 23 | | 15 | | 27 | inside | 0.6544 |
|  | VYADSFVIRGDEVRQ | 2 | 16 | | 15 | | 27 | inside | -0.2063 |
|  | NVYADSFVIRGDEVR | 1 | 15 | | 15 | | 27 | inside | -0.2814 |
|  | GDEVRQIAPGQTGKI | 11 | 25 | | 15 | | 27 | inside | 0.9741 |
|  | RGDEVRQIAPGQTGK | 10 | 24 | | 15 | | 27 | inside | 0.9034 |
|  | DEVRQIAPGQTGKIA | 12 | 26 | | 15 | | 27 | inside | 0.9862 |
|  | QTGKIADYNYKLPDD | 21 | 35 | | 15 | | 27 | inside | 0.9278 |
|  | EVRQIAPGQTGKIAD | 13 | 27 | | 15 | | 27 | inside | 1.3487 |
|  | RQIAPGQTGKIADYN | 15 | 29 | | 15 | | 27 | inside | 1.5209 |
|  | VRQIAPGQTGKIADY | 14 | 28 | | 15 | | 27 | inside | 1.3048 |
|  | GQTGKIADYNYKLPD | 20 | 34 | | 15 | | 27 | inside | 1.3474 |
|  | APGQTGKIADYNYKL | 18 | 32 | | 15 | | 27 | inside | 1.4441 |
|  | PGQTGKIADYNYKLP | 19 | 33 | | 15 | | 27 | inside | 1.3465 |
|  | QIAPGQTGKIADYNY | 16 | 30 | | 15 | | 27 | inside | 1.6088 |
|  | IAPGQTGKIADYNYK | 17 | 31 | | 15 | | 27 | inside | 1.8362 |
