## Supplementary Table 2 for "Genome based Evolutionary study of SARS-CoV-2 towards the Prediction of Epitope Based Chimeric Vaccine"

**Supplementary Table 2:** Predicted CTL and HTL epitopes of membrane glycoprotein.

| **Types** | Epitope | Start | End | Length | Antigenic Value | Topology | No. of HLAs |
| --- | --- | --- | --- | --- | --- | --- | --- |
| **CTL epitopes** | SELVIGAVIL | 16 | 25 | 10 | 0.6521 | Outside | 27 |
|  | ELVIGAVILR | 17 | 26 | 10 | 0.9998 | Outside | 27 |
|  | LVIGAVILR | 18 | 26 | 9 | 1.1027 | Outside | 54 |
|  | SELVIGAVI | 16 | 24 | 9 | 0.6409 | Outside | 54 |
|  | AVILRGHLR | 22 | 30 | 9 | 0.3371 | Inside | 81 |
|  | RIAGHHLGR | 30 | 38 | 9 | -0.4947 | Inside | 81 |
|  | RPLLESELVI | 11 | 20 | 10 | 0.2055 | Outside | 27 |
|  | ESELVIGAV | 15 | 23 | 9 | 0.9872 | Outside | 54 |
|  | HLRIAGHHL | 28 | 36 | 9 | 0.2446 | Outside | 81 |
|  | ILRGHLRIA | 24 | 32 | 9 | 0.5057 | Inside | 81 |
|  | RPLLESELV | 11 | 19 | 9 | 0.1504 | Outside | 54 |
|  | HGTILTRPLL | 5 | 14 | 10 | 0.0776 | Outside | 27 |
|  | LLESELVIGA | 13 | 22 | 10 | 0.6692 | Outside | 27 |
|  | LESELVIGA | 14 | 22 | 9 | 0.8597 | Outside | 27 |
|  | VILRGHLRI | 23 | 31 | 9 | 0.4298 | Inside | 27 |
|  | LTRPLLESEL | 9 | 18 | 10 | -0.3509 | Outside | 27 |
|  | VPLHGTILT | 2 | 10 | 9 | -0.016 | Outside | 81 |
|  | HGTILTRPL | 5 | 13 | 9 | 0.2144 | Outside | 54 |
|  | GTILTRPLL | 6 | 14 | 9 | 0.1403 | Outside | 54 |
|  | IGAVILRGHL | 20 | 29 | 10 | 0.681 | Outside | 27 |
|  | AGHHLGRCDI | 32 | 41 | 10 | -0.0222 | Inside | 27 |
|  | RGHLRIAGHH | 26 | 35 | 10 | 0.1013 | Inside | 27 |
|  | ELVIGAVIL | 17 | 25 | 9 | 0.7969 | Outside | 27 |
|  | HLGRCDIKDL | 35 | 44 | 10 | 1.1136 | Inside | 27 |
|  | PLHGTILTR | 3 | 11 | 9 | 0.7584 | Outside | 54 |
|  | CDIKDLPKEI | 39 | 48 | 10 | 0.3695 | Inside | 27 |
|  | PLLESELVI54 | 12 | 20 | 9 | 0.5354 | Outside | 54 |
|  | NVPLHGTIL | 1 | 9 | 9 | -0.1458 | Outside | 27 |
|  | LGRCDIKDL | 36 | 44 | 9 | 1.2129 | Inside | 54 |
|  | RGHLRIAGH | 26 | 34 | 9 | -0.0258 | Inside | 54 |
|  | RCDIKDLPK | 38 | 46 | 9 | 0.5923 | Inside | 81 |
|  | GAVILRGHL | 21 | 29 | 9 | 0.434 | Outside | 27 |
|  | HHLGRCDIK | 34 | 42 | 9 | 0.4984 | Inside | 81 |
|  | TILTRPLLES | 7 | 16 | 10 | -0.2168 | Outside | 27 |
|  | TILTRPLLE | 7 | 15 | 9 | -0.4576 | Outside | 27 |
|  | VIGAVILRGH | 19 | 28 | 10 | 0.5844 | Outside | 27 |
|  | DIKDLPKEI | 40 | 48 | 9 | 0.5528 | Inside | 27 |
|  | LTRPLLESE | 9 | 17 | 9 | -0.3434 | Outside | 54 |
|  | GHLRIAGHH | 27 | 35 | 9 | -0.0013 | Inside | 54 |
|  | IAGHHLGRC | 31 | 39 | 9 | -0.3274 | Inside | 54 |
|  | ILTRPLLES | 8 | 16 | 9 | -0.4392 | Outside | 27 |
|  | IGAVILRGH | 20 | 28 | 9 | 0.9127 | Outside | 27 |
|  | VIGAVILRG | 19 | 27 | 9 | 0.6184 | Outside | 27 |
|  | LLESELVIG | 13 | 21 | 9 | 0.6747 | Outside | 27 |
|  | MWLSYFIASF | 30 | 39 | 10 | -0.1104 | Outside | 28 |
|  | SYFIASFRLF | 33 | 42 | 10 | -0.0998 | Outside | 27 |
|  | YFIASFRLF | 34 | 42 | 9 | -0.1142 | Outside | 54 |
|  | LSYFIASFR | 32 | 40 | 9 | 0.3283 | Outside | 81 |
|  | FIASFRLFAR | 35 | 44 | 10 | 0.0220 | Outside | 27 |
|  | MACLVGLMW | 23 | 31 | 9 | 0.7889 | Outside | 81 |
|  | GLMWLSYFI | 28 | 36 | 9 | 0.2537 | Outside | 81 |
|  | LAAVYRINW | 6 | 14 | 9 | 1.4322 | Inside | 82 |
|  | SYFIASFRL | 33 | 41 | 9 | 0.4821 | Outside | 27 |
|  | GHHLGRCDI | 33 | 41 | 9 | 0.015 | Inside | 27 |
|  | LRIAGHHLG | 29 | 37 | 9 | -0.238 | Outside | 27 |
|  | AGHHLGRCD | 32 | 40 | 9 | -0.0538 | Inside | 27 |
|  | TRPLLESEL | 10 | 18 | 9 | -0.1264 | Outside | 27 |
|  | CDIKDLPKE | 39 | 47 | 9 | 0.5074 | Inside | 27 |
|  | LHGTILTRP | 4 | 12 | 9 | 0.4759 | Outside | 27 |
|  | LRGHLRIAG | 25 | 33 | 9 | 0.1788 | Outside | 27 |
|  | HLGRCDIKD | 35 | 43 | 9 | 1.186 | Inside | 27 |
|  | GRCDIKDLP | 37 | 45 | 9 | 1.4662 | Inside | 27 |
|  | IASFRLFAR | 36 | 44 | 9 | 0.0646 | Outside | 54 |
|  | ASFRLFARTR | 37 | 46 | 10 | 0.6237 | Inside | 27 |
|  | LACFVLAAVY | 1 | 10 | 10 | 1.0354 | Outside | 27 |
|  | WLSYFIASF | 31 | 39 | 9 | -0.1644 | Outside | 27 |
|  | RTRSMWSFN | 44 | 52 | 9 | 1.4147 | Inside | 81 |
|  | RLFARTRSM | 40 | 48 | 9 | 0.1998 | Inside | 82 |
|  | FVLAAVYRI | 4 | 12 | 9 | 0.5136 | Inside | 81 |
|  | CFVLAAVYR | 3 | 11 | 9 | 1.0909 | Inside | 54 |
|  | CLVGLMWLSY | 25 | 34 | 10 | 1.0255 | Outside | 27 |
|  | IAIAMACLV | 19 | 27 | 9 | 1.1704 | Inside | 82 |
|  | ACFVLAAVY | 2 | 10 | 9 | 1.1400 | Inside | 27 |
|  | RINWITGGI | 11 | 19 | 9 | 1.4580 | Inside | 81 |
|  | FARTRSMWSF | 42 | 51 | 10 | 0.9202 | Inside | 27 |
|  | LVGLMWLSY | 26 | 34 | 9 | 1.0633 | Outside | 54 |
|  | SFRLFARTR | 38 | 46 | 9 | 0.7038 | Inside | 54 |
|  | LMWLSYFIA | 29 | 37 | 9 | 0.3601 | Outside | 54 |
|  | VGLMWLSYF | 27 | 35 | 9 | 0.7686 | Outside | 27 |
|  | AMACLVGLM | 22 | 30 | 9 | 0.6251 | Outside | 54 |
|  | FARTRSMWS | 42 | 50 | 9 | 0.6000 | Inside | 54 |
|  | SMWSFNPET | 47 | 55 | 9 | 0.7633 | Outside | 81 |
|  | FIASFRLFA | 35 | 43 | 9 | -0.2267 | Outside | 27 |
|  | MWSFNPETNI | 48 | 57 | 10 | 0.4936 | Inside | 27 |
|  | IAMACLVGL | 21 | 29 | 9 | 1.1306 | Inside | 55 |
|  | LFARTRSMW | 41 | 49 | 9 | 0.8560 | Inside | 27 |
|  | LACFVLAAV | 1 | 9 | 9 | 1.1825 | Outside | 27 |
|  | RSMWSFNPE | 46 | 54 | 9 | 1.1476 | Inside | 54 |
|  | MWLSYFIAS | 30 | 38 | 9 | -0.0642 | Outside | 27 |
|  | VYRINWITG | 9 | 17 | 9 | 0.5362 | Inside | 81 |
|  | AAVYRINWI | 7 | 15 | 9 | 1.1454 | Inside | 54 |
|  | WSFNPETNI | 49 | 57 | 9 | 0.7550 | Outside | 54 |
|  | MWSFNPETN | 48 | 56 | 9 | 0.5509 | Outside | 27 |
|  | ITGGIAIAM | 15 | 23 | 9 | 0.7715 | Inside | 81 |
|  | WITGGIAIA | 14 | 22 | 9 | 0.8209 | Inside | 54 |
|  | NWITGGIAI | 13 | 21 | 9 | 0.9690 | Inside | 54 |
|  | ASFRLFART | 37 | 45 | 9 | 0.2319 | Inside | 27 |
|  | GIAIAMACL | 18 | 26 | 9 | 1.2059 | Outside | 54 |
|  | ACLVGLMWL | 24 | 32 | 9 | 0.7321 | Outside | 54 |
|  | ARTRSMWSF | 43 | 51 | 9 | 1.2394 | Inside | 27 |
|  | CLVGLMWLS | 25 | 33 | 9 | 0.8369 | Outside | 27 |
|  | SFNPETNIL | 1 | 50 | 58 | 0.3864 | Outside | 54 |
|  | AVYRINWIT | 8 | 16 | 9 | 0.7987 | Inside | 27 |
|  | FNPETNILL | 51 | 59 | 9 | 0.2578 | Outside | 28 |
|  | VLAAVYRIN | 5 | 13 | 9 | 0.5773 | Inside | 27 |
|  | INWITGGIA | 12 | 20 | 9 | 1.1906 | Inside | 27 |
|  | TGGIAIAMAC | 16 | 25 | 10 | 0.8528 | Inside | 27 |
|  | GGIAIAMAC | 17 | 25 | 9 | 0.9415 | Inside | 27 |
|  | TRSMWSFNP | 45 | 53 | 9 | 1.3775 | Inside | 27 |
|  | YRINWITGG | 10 | 18 | 9 | 1.4003 | Inside | 27 |
|  | AIAMACLVG | 20 | 28 | 9 | 0.9160 | Inside | 27 |
|  | TGGIAIAMA | 16 | 24 | 9 | 0.8190 | Inside | 27 |
|  | FRLFARTRS | 39 | 47 | 9 | 0.4849 | Inside | 27 |
| **HTL epitopes** | GLMWLSYFIASFRLF | 28 | 42 | 15 | 0.0766 | Outside | 27 |
|  | LMWLSYFIASFRLFA | 29 | 43 | 15 | 0.1022 | Outside | 27 |
|  | VGLMWLSYFIASFRL | 27 | 41 | 15 | 0.6658 | Outside | 27 |
|  | MWLSYFIASFRLFAR | 30 | 44 | 15 | 0.0043 | Outside | 27 |
|  | WLSYFIASFRLFART | 31 | 45 | 15 | 0.0462 | Outside | 27 |
|  | LVGLMWLSYFIASFR | 26 | 40 | 15 | 0.5535 | Outside | 27 |
|  | CLVGLMWLSYFIASF | 25 | 39 | 15 | 0.4280 | Outside | 27 |
|  | LSYFIASFRLFARTR | 32 | 46 | 15 | 0.3427 | Outside | 27 |
|  | SYFIASFRLFARTRS | 33 | 47 | 15 | 0.3751 | Outside | 27 |
|  | YFIASFRLFARTRSM | 34 | 48 | 15 | 0.3975 | Inside | 27 |
|  | ASFRLFARTRSMWSF | 37 | 51 | 15 | 0.7304 | Inside | 27 |
|  | FIASFRLFARTRSMW | 35 | 49 | 15 | 0.4072 | Outside | 27 |
|  | IASFRLFARTRSMWS | 36 | 50 | 15 | 0.4424 | Outside | 27 |
|  | NWITGGIAIAMACLV | 13 | 27 | 15 | 0.8998 | Inside | 27 |
|  | MACLVGLMWLSYFIA | 23 | 37 | 15 | 0.5584 | Outside | 27 |
|  | INWITGGIAIAMACL | 12 | 26 | 15 | 1.1352 | Inside | 27 |
|  | RINWITGGIAIAMAC | 11 | 25 | 15 | 1.1629 | Inside | 27 |
|  | ACLVGLMWLSYFIAS | 24 | 38 | 15 | 0.4585 | Inside | 27 |
|  | YRINWITGGIAIAMA | 10 | 24 | 15 | 1.1274 | Inside | 27 |
|  | WITGGIAIAMACLVG | 14 | 28 | 15 | 0.8261 | Outside | 27 |
|  | LACFVLAAVYRINWI | 1 | 15 | 15 | 1.2905 | Outside | 27 |
|  | SFRLFARTRSMWSFN | 38 | 52 | 15 | 0.7955 | Inside | 27 |
|  | VYRINWITGGIAIAM | 9 | 23 | 15 | 0.9236 | Inside | 27 |
|  | ACFVLAAVYRINWIT | 2 | 16 | 15 | 1.1115 | Inside | 27 |
|  | AAVYRINWITGGIAI | 7 | 21 | 15 | 0.9197 | Inside | 27 |
|  | AVYRINWITGGIAIA | 8 | 22 | 15 | 0.9497 | Inside | 27 |
|  | FRLFARTRSMWSFNP | 39 | 53 | 15 | 0.8873 | Inside | 27 |
|  | LAAVYRINWITGGIA | 6 | 20 | 15 | 1.0581 | Inside | 27 |
|  | LFARTRSMWSFNPET | 41 | 55 | 15 | 1.0581 | Inside | 27 |
|  | RLFARTRSMWSFNPE | 40 | 54 | 15 | 0.7744 | Inside | 27 |
|  | VLAAVYRINWITGGI | 5 | 19 | 15 | 0.9478 | Inside | 27 |
|  | ITGGIAIAMACLVGL | 15 | 29 | 15 | 0.9310 | Outside | 27 |
|  | CFVLAAVYRINWITG | 3 | 17 | 15 | 1.0062 | Inside | 27 |
|  | FVLAAVYRINWITGG | 4 | 18 | 15 | 1.0230 | Outside | 27 |
|  | TRSMWSFNPETNILL | 45 | 59 | 15 | 0.6067 | Inside | 27 |
|  | AMACLVGLMWLSYFI | 22 | 36 | 15 | 0.5388 | Outside | 27 |
|  | FARTRSMWSFNPETN | 42 | 56 | 15 | 0.7761 | Inside | 27 |
|  | TGGIAIAMACLVGLM | 16 | 30 | 15 | 0.8302 | Inside | 27 |
|  | RTRSMWSFNPETNIL | 44 | 58 | 15 | 0.7782 | Inside | 27 |
|  | IAMACLVGLMWLSYF | 21 | 35 | 15 | 0.7552 | Outside | 27 |
|  | GGIAIAMACLVGLMW | 17 | 31 | 15 | 0.9049 | Outside | 27 |
|  | AIAMACLVGLMWLSY | 20 | 34 | 15 | 0.9526 | Outside | 27 |
|  | IAIAMACLVGLMWLS | 19 | 33 | 15 | 0.9464 | Outside | 27 |
|  | GIAIAMACLVGLMWL | 18 | 32 | 15 | 0.9155 | Outside | 27 |
|  | ARTRSMWSFNPETNI | 43 | 57 | 15 | 0.8632 | Inside | 27 |
|  | ESELVIGAVILRGHL | 15 | 29 | 15 | 0.5735 | Outside | 27 |
|  | AVILRGHLRIAGHHL | 22 | 36 | 15 | 0.1069 | Outside | 27 |
|  | ILRGHLRIAGHHLGR | 24 | 38 | 15 | -0.0989 | Outside | 27 |
|  | LRGHLRIAGHHLGRC | 25 | 39 | 15 | -0.1484 | Outside | 27 |
|  | VILRGHLRIAGHHLG | 23 | 37 | 15 | 0.0809 | Outside | 27 |
|  | SELVIGAVILRGHLR | 16 | 30 | 15 | 0.6768 | Outside | 27 |
|  | ELVIGAVILRGHLRI | 17 | 31 | 15 | 0.7972 | Outside | 27 |
|  | LESELVIGAVILRGH | 14 | 28 | 15 | 0.6528 | Outside | 27 |
|  | LLESELVIGAVILRG | 13 | 27 | 15 | 0.5636 | Outside | 27 |
|  | LVIGAVILRGHLRIA | 18 | 32 | 15 | 0.8769 | Outside | 27 |
|  | GTILTRPLLESELVI | 6 | 20 | 15 | 0.2554 | Outside | 27 |
|  | PLHGTILTRPLLESE | 3 | 17 | 15 | 0.2336 | Outside | 27 |
|  | VPLHGTILTRPLLES | 2 | 16 | 15 | 0.0197 | Outside | 27 |
|  | HGTILTRPLLESELV | 5 | 19 | 15 | 0.1887 | Outside | 27 |
|  | PLLESELVIGAVILR | 12 | 26 | 15 | 0.7261 | Outside | 27 |
|  | VIGAVILRGHLRIAG | 19 | 33 | 15 | 0.4903 | Outside | 27 |
|  | RGHLRIAGHHLGRCD | 26 | 40 | 15 | 0.0223 | Inside | 27 |
|  | IGAVILRGHLRIAGH | 20 | 34 | 15 | 0.4539 | Outside | 27 |
|  | LHGTILTRPLLESEL | 4 | 18 | 15 | 0.1462 | Inside | 27 |
|  | GAVILRGHLRIAGHH | 21 | 35 | 15 | 0.3053 | Outside | 27 |
|  | RPLLESELVIGAVIL | 11 | 25 | 15 | 0.4332 | Outside | 27 |
|  | TILTRPLLESELVIG | 7 | 21 | 15 | 0.2970 | Outside | 27 |
|  | GHLRIAGHHLGRCDI | 27 | 41 | 15 | 0.0062 | Outside | 27 |
|  | LTRPLLESELVIGAV | 9 | 23 | 15 | 0.2715 | Outside | 27 |
|  | TRPLLESELVIGAVI | 10 | 24 | 15 | 0.4473 | Outside | 27 |
|  | HLRIAGHHLGRCDIK | 28 | 42 | 15 | 0.4291 | Outside | 27 |
|  | ILTRPLLESELVIGA | 8 | 22 | 15 | 0.2447 | Outside | 27 |
|  | NVPLHGTILTRPLLE | 1 | 15 | 15 | -0.1534 | Outside | 27 |
|  | LRIAGHHLGRCDIKD | 29 | 43 | 15 | 0.4163 | Outside | 27 |
|  | HHLGRCDIKDLPKEI | 34 | 48 | 15 | 0.3167 | Inside | 27 |
|  | AGHHLGRCDIKDLPK | 32 | 46 | 15 | 0.3469 | Inside | 27 |
|  | GHHLGRCDIKDLPKE | 33 | 47 | 15 | 0.3828 | Inside | 27 |
|  | IAGHHLGRCDIKDLP | 31 | 45 | 15 | 0.5721 | Outside | 27 |
|  | RIAGHHLGRCDIKDL | 30 | 44 | 15 | 0.3979 | Inside | 27 |
