## Supplementary Table 3 for "Genome based Evolutionary study of SARS-CoV-2 towards the Prediction of Epitope Based Chimeric Vaccine"

**Supplementary Table 3:** Predicted CTL and HTL epitopes of envelope protein.

| **Type** | **Epitope** | **Start** | **End** | **Length** | **Antigenic Value** | **Topology** | **No. of HLAs** |
| --- | --- | --- | --- | --- | --- | --- | --- |
| CTL Epitopes | LLFLAFVVF | 18 | 26 | 9 | 0.8144 | Outside | 81 |
|  | LTALRLCAY | 34 | 42 | 9 | 0.2825 | Inside | 81 |
|  | FLAFVVFLLV | 20 | 29 | 10 | 0.5651 | Outside | 27 |
|  | RVKNLNSSR | 61 | 69 | 9 | 0.8998 | Inside | 81 |
|  | FLAFVVFLL | 20 | 28 | 9 | 0.5308 | Outside | 54 |
|  | LVKPSFYVY | 51 | 59 | 9 | 0.4213 | Outside | 81 |
|  | LAFVVFLLV | 21 | 29 | 9 | 0.7976 | Outside | 54 |
|  | TLAILTALR | 30 | 38 | 9 | 0.7223 | Inside | 81 |
|  | VLLFLAFVV | 17 | 25 | 9 | 0.5677 | Outside | 54 |
|  | VSLVKPSFY | 49 | 57 | 9 | 0.7476 | Outside | 81 |
|  | SVLLFLAFV | 16 | 24 | 9 | 0.4765 | Outside | 54 |
|  | SLVKPSFYV | 50 | 58 | 9 | 0.4140 | Outside | 27 |
|  | FLLVTLAIL | 26 | 34 | 9 | 0.9645 | Outside | 81 |
|  | SEETGTLIV | 6 | 14 | 9 | 0.3052 | Outside | 81 |
|  | FVVFLLVTL | 23 | 31 | 9 | 0.7403 | Outside | 81 |
|  | VFLLVTLAI | 25 | 33 | 9 | 0.8134 | Outside | 54 |
|  | ETGTLIVNSV | 8 | 17 | 10 | 0.3003 | Inside | 27 |
|  | IVNSVLLFL | 13 | 21 | 9 | -0.0239 | Outside | 81 |
|  | TLIVNSVLLF | 11 | 20 | 10 | -0.0561 | Outside | 27 |
|  | LIVNSVLLF | 12 | 20 | 9 | -0.0953 | Outside | 27 |
|  | RLCAYCCNI | 38 | 46 | 9 | 1.1243 | Inside | 81 |
|  | NSVLLFLAF | 15 | 23 | 9 | 0.4134 | Outside | 54 |
|  | FVSEETGTLI | 4 | 13 | 10 | 0.3044 | Outside | 27 |
|  | NIVNVSLVK | 45 | 53 | 9 | 0.9310 | Inside | 81 |
|  | VTLAILTAL | 29 | 37 | 9 | 0.6140 | Outside | 54 |
|  | KPSFYVYSR | 53 | 61 | 9 | 0.9740 | Inside | 81 |
|  | LAILTALRL | 31 | 39 | 9 | 0.8872 | Outside | 54 |
|  | FVSEETGTL | 4 | 12 | 9 | 0.3864 | Outside | 54 |
|  | FYVYSRVKNL | 56 | 65 | 10 | 0.8286 | Inside | 27 |
|  | YVYSRVKNL | 57 | 65 | 9 | 0.7020 | Inside | 54 |
|  | LLVTLAILTA | 27 | 36 | 10 | 0.5774 | Outside | 27 |
|  | LFLAFVVFL | 19 | 27 | 9 | 0.4568 | Outside | 27 |
|  | TGTLIVNSV | 9 | 17 | 9 | 0.2573 | Inside | 54 |
|  | SFYVYSRVK | 55 | 63 | 9 | 0.8251 | Inside | 81 |
|  | SSRVPDLLV | 67 | 75 | 9 | 0.5455 | Outside | 54 |
|  | AYCCNIVNV | 41 | 49 | 9 | 1.0856 | Inside | 81 |
|  | VSEETGTLI | 5 | 13 | 9 | 0.2570 | Outside | 27 |
|  | AILTALRLCA | 32 | 41 | 10 | 0.1882 | Inside | 27 |
|  | MYSFVSEETG | 1 | 10 | 10 | 0.3885 | Outside | 27 |
|  | TLIVNSVLL | 11 | 19 | 9 | -0.0406 | Outside | 54 |
|  | VYSRVKNLN | 58 | 66 | 9 | 1.0946 | Inside | 54 |
|  | EETGTLIVN | 7 | 15 | 9 | 0.3631 | Inside | 54 |
|  | LCAYCCNIV | 39 | 47 | 9 | 0.7349 | Inside | 54 |
|  | NSSRVPDLL | 66 | 74 | 9 | 0.6086 | Outside | 27 |
|  | ETGTLIVNS | 8 | 16 | 9 | 0.2951 | Inside | 27 |
|  | CAYCCNIVN | 40 | 48 | 9 | 1.0526 | Inside | 27 |
|  | VVFLLVTLA | 24 | 32 | 9 | 0.9374 | Outside | 27 |
|  | VNVSLVKPSF | 47 | 56 | 10 | 0.7346 | Outside | 27 |
|  | PSFYVYSRV | 54 | 62 | 9 | 0.6060 | Outside | 27 |
|  | FYVYSRVKN | 56 | 64 | 9 | 0.8145 | Inside | 27 |
|  | YCCNIVNVSL | 42 | 51 | 10 | 1.3643 | Inside | 27 |
|  | LLVTLAILT | 27 | 35 | 9 | 0.5583 | Outside | 27 |
|  | MYSFVSEET | 1 | 9 | 9 | 0.3428 | Outside | 27 |
|  | LNSSRVPDL | 65 | 73 | 9 | 0.8553 | Outside | 54 |
|  | YSFVSEETGT | 2 | 11 | 10 | 0.5995 | Outside | 27 |
|  | GTLIVNSVL | 10 | 18 | 9 | 0.0475 | Outside | 27 |
|  | CNIVNVSLV | 44 | 52 | 9 | 1.4201 | Inside | 54 |
|  | NVSLVKPSF | 48 | 56 | 9 | 0.6836 | Outside | 27 |
|  | CCNIVNVSL | 43 | 51 | 9 | 1.3710 | Inside | 27 |
|  | YSFVSEETG | 2 | 10 | 9 | 0.6840 | Outside | 27 |
|  | LVTLAILTA | 28 | 36 | 9 | 0.5720 | Outside | 27 |
|  | YSRVKNLNS | 59 | 67 | 9 | 0.5961 | Inside | 54 |
|  | TALRLCAYC | 35 | 43 | 9 | 0.9356 | Inside | 54 |
|  | ILTALRLCA | 33 | 41 | 9 | 0.1234 | Inside | 27 |
|  | IVNVSLVKPS | 46 | 55 | 10 | 0.5729 | Outside | 27 |
|  | KNLNSSRVPD | 63 | 72 | 10 | 0.4746 | Inside | 27 |
|  | VNSVLLFLA | 14 | 22 | 9 | 0.1915 | Outside | 27 |
|  | AILTALRLC | 32 | 40 | 9 | 0.3823 | Inside | 27 |
|  | KNLNSSRVP | 63 | 71 | 9 | 0.2669 | Inside | 54 |
|  | ALRLCAYCC | 36 | 44 | 9 | 0.7465 | Inside | 54 |
|  | NLNSSRVPD | 64 | 72 | 9 | 0.4406 | Outside | 27 |
|  | VKNLNSSRV | 62 | 70 | 9 | 0.4432 | Inside | 27 |
|  | AFVVFLLVT | 22 | 30 | 9 | 0.6178 | Outside | 27 |
|  | LRLCAYCCN | 37 | 45 | 9 | 0.7603 | Inside | 27 |
|  | IVNVSLVKP | 46 | 54 | 9 | 0.4312 | Outside | 27 |
|  | YCCNIVNVS | 42 | 50 | 9 | 1.3929 | Inside | 27 |
|  | SRVKNLNSS | 60 | 68 | 9 | 0.9490 | Inside | 27 |
|  | VKPSFYVYS | 52 | 60 | 9 | 1.0547 | Outside | 27 |
|  | VNVSLVKPS | 47 | 55 | 9 | 0.5145 | Outside | 27 |
|  | SFVSEETGT | 3 | 11 | 9 | 0.5596 | Outside | 27 |
| HTL Epitopes | LLFLAFVVFLLVTLA | 18 | 32 | 15 | 0.8122 | Outside | 27 |
|  | VLLFLAFVVFLLVTL | 17 | 31 | 15 | 0.6386 | Outside | 27 |
|  | LFLAFVVFLLVTLAI | 19 | 33 | 15 | 0.7471 | Outside | 27 |
|  | VYSRVKNLNSSRVPD | 58 | 72 | 15 | 0.5993 | Inside | 27 |
|  | YVYSRVKNLNSSRVP | 57 | 71 | 15 | 0.4492 | Inside | 27 |
|  | SFYVYSRVKNLNSSR | 55 | 69 | 15 | 0.6291 | Inside | 27 |
|  | FYVYSRVKNLNSSRV | 56 | 70 | 15 | 0.6103 | Inside | 27 |
|  | VSEETGTLIVNSVLL | 5 | 19 | 15 | 0.1951 | Outside | 27 |
|  | KPSFYVYSRVKNLNS | 53 | 67 | 15 | 0.8229 | Inside | 27 |
|  | LTALRLCAYCCNIVN | 34 | 48 | 15 | 0.8649 | Inside | 27 |
|  | VKPSFYVYSRVKNLN | 52 | 66 | 15 | 1.2319 | Inside | 27 |
|  | IVNSVLLFLAFVVFL | 13 | 27 | 15 | 0.2731 | Outside | 27 |
|  | SVLLFLAFVVFLLVT | 16 | 30 | 15 | 0.5446 | Outside | 27 |
|  | TALRLCAYCCNIVNV | 35 | 49 | 15 | 0.8876 | Inside | 27 |
|  | LVKPSFYVYSRVKNL | 51 | 65 | 15 | 0.7311 | Outside | 27 |
|  | SLVKPSFYVYSRVKN | 50 | 64 | 15 | 0.6514 | Outside | 27 |
|  | SFVSEETGTLIVNSV | 3 | 17 | 15 | 0.3658 | Outside | 27 |
|  | YSFVSEETGTLIVNS | 2 | 16 | 15 | 0.3987 | Outside | 27 |
|  | RVKNLNSSRVPDLLV | 61 | 75 | 15 | 0.7925 | Inside | 27 |
|  | NSVLLFLAFVVFLLV | 15 | 29 | 15 | 0.4220 | Outside | 27 |
|  | PSFYVYSRVKNLNSS | 54 | 68 | 15 | 0.7986 | Inside | 27 |
|  | VNSVLLFLAFVVFLL | 14 | 28 | 15 | 0.3985 | Outside | 27 |
|  | FLAFVVFLLVTLAIL | 20 | 34 | 15 | 0.7476 | Outside | 27 |
|  | MYSFVSEETGTLIVN | 1 | 15 | 15 | 0.2951 | Outside | 27 |
|  | LRLCAYCCNIVNVSL | 37 | 51 | 15 | 1.0896 | Inside | 27 |
|  | VSLVKPSFYVYSRVK | 49 | 63 | 15 | 0.7974 | Outside | 27 |
|  | VNVSLVKPSFYVYSR | 47 | 61 | 15 | 0.7513 | Outside | 27 |
|  | NVSLVKPSFYVYSRV | 48 | 62 | 15 | 0.6449 | Outside | 27 |
|  | RLCAYCCNIVNVSLV | 38 | 52 | 15 | 1.2823 | Inside | 27 |
|  | ALRLCAYCCNIVNVS | 36 | 50 | 15 | 0.9857 | Inside | 27 |
|  | ILTALRLCAYCCNIV | 33 | 47 | 15 | 0.7427 | Outside | 27 |
|  | FVSEETGTLIVNSVL | 4 | 18 | 15 | 0.1704 | Outside | 27 |
|  | YSRVKNLNSSRVPDL | 59 | 73 | 15 | 0.7705 | Inside | 27 |
|  | LIVNSVLLFLAFVVF | 12 | 26 | 15 | 0.2925 | Outside | 27 |
|  | LAFVVFLLVTLAILT | 21 | 35 | 15 | 0.8229 | Outside | 27 |
|  | FVVFLLVTLAILTAL | 23 | 37 | 15 | 0.5738 | Outside | 27 |
|  | AILTALRLCAYCCNI | 32 | 46 | 15 | 0.7040 | Outside | 27 |
|  | CNIVNVSLVKPSFYV | 44 | 58 | 15 | 0.8081 | Inside | 27 |
|  | SRVKNLNSSRVPDLL | 60 | 74 | 15 | 0.7404 | Inside | 27 |
|  | LCAYCCNIVNVSLVK | 39 | 53 | 15 | 0.8552 | Inside | 27 |
|  | CCNIVNVSLVKPSFY | 43 | 57 | 15 | 0.8879 | Inside | 27 |
|  | IVNVSLVKPSFYVYS | 46 | 60 | 15 | 0.6373 | Outside | 27 |
|  | NIVNVSLVKPSFYVY | 45 | 59 | 15 | 0.7680 | Inside | 27 |
|  | CAYCCNIVNVSLVKP | 40 | 54 | 15 | 0.8337 | Inside | 27 |
|  | SEETGTLIVNSVLLF | 6 | 20 | 15 | 0.1882 | Outside | 27 |
|  | GTLIVNSVLLFLAFV | 10 | 24 | 15 | 0.3383 | Outside | 27 |
|  | AYCCNIVNVSLVKPS | 41 | 55 | 15 | 0.8053 | Inside | 27 |
|  | LVTLAILTALRLCAY | 28 | 42 | 15 | 0.4070 | Outside | 27 |
|  | TLAILTALRLCAYCC | 30 | 44 | 15 | 0.7304 | Outside | 27 |
|  | VVFLLVTLAILTALR | 24 | 38 | 15 | 0.7559 | Outside | 27 |
|  | LAILTALRLCAYCCN | 31 | 45 | 15 | 0.7009 | Outside | 27 |
|  | VTLAILTALRLCAYC | 29 | 43 | 15 | 0.8599 | Outside | 27 |
|  | TGTLIVNSVLLFLAF | 9 | 23 | 15 | 0.3354 | Outside | 27 |
|  | VFLLVTLAILTALRL | 25 | 39 | 15 | 0.7218 | Outside | 27 |
|  | TLIVNSVLLFLAFVV | 11 | 25 | 15 | 0.2903 | Outside | 27 |
|  | YCCNIVNVSLVKPSF | 42 | 56 | 15 | 0.9396 | Inside | 27 |
|  | FLLVTLAILTALRLC | 26 | 40 | 15 | 0.6311 | Outside | 27 |
|  | LLVTLAILTALRLCA | 27 | 41 | 15 | 0.3840 | Outside | 27 |
