## Supplementary Table 4 for "Genome based Evolutionary study of SARS-CoV-2 towards the Prediction of Epitope Based Chimeric Vaccine"

**Supplementary Table 4:** Predicted CTL and HTL epitopes of nucleocapsid protein.

| **Type** | **Epitope** | **start** | **end** | **length** | **topology** | **Vaxijen score** | **No of HLAs** |
| --- | --- | --- | --- | --- | --- | --- | --- |
| CTL epitopes | FPRGQGVPI | 66 | 74 | 9 | Outside | 0.7585 | 81 |
|  | FTALTQHGK | 53 | 61 | 9 | Inside | 0.8510 | 81 |
|  | SPRWYFYYL | 105 | 113 | 9 | Outside | 0.7340 | 81 |
|  | SSPDDQIGYY | 78 | 87 | 10 | Outside | 0.4533 | 27 |
|  | LSPRWYFYY | 104 | 112 | 9 | Outside | 1.2832 | 54 |
|  | LPYGANKDGI | 121 | 130 | 10 | Outside | 0.0009 | 27 |
|  | YGANKDGIIW | 123 | 132 | 9 | Inside | -0.5336 | 27 |
|  | YYRRATRRI | 86 | 94 | 9 | Inside | -0.6565 | 81 |
|  | LPNNTASWF | 45 | 53 | 9 | Outside | -0.0835 | 81 |
|  | KMKDLSPRWY | 100 | 109 | 10 | Inside | 1.5032 | 27 |
|  | ATEGALNTPK | 134 | 143 | 10 | Inside | -0.3602 | 27 |
|  | IGYYRRATR | 84 | 92 | 9 | Inside | 0.8880 | 81 |
|  | APRITFGGPS | 12 | 21 | 9 | Outside | 0.6584 | 27 |
|  | NTASWFTAL | 48 | 56 | 9 | Outside | -0.0708 | 81 |
|  | RSKQRRPQG | 36 | 44 | 9 | Inside | -0.3861 | 81 |
|  | GYYRRATRR | 85 | 93 | 9 | Inside | -0.4627 | 27 |
|  | RPQGLPNNTA | 41 | 50 | 10 | Inside | 0.5551 | 27 |
|  | KDLSPRWYFY | 102 | 111 | 10 | Inside | 0.9981 | 27 |
|  | NQRNAPRITF | 8 | 17 | 10 | Inside | 0.8505 | 27 |
|  | GANKDGIIW | 124 | 132 | 9 | Inside | -0.6134 | 54 |
|  | DGKMKDLSPR | 98 | 107 | 10 | Inside | 1.2869 | 27 |
|  | NSSPDDQIGY | 77 | 86 | 10 | Outside | 0.6630 | 27 |
|  | KMKDLSPRW | 100 | 108 | 9 | Inside | 1.7462 | 54 |
|  | RSGARSKQR | 32 | 40 | 9 | Inside | 1.7874 | 81 |
|  | QASSRSSSR | 181 | 189 | 9 | Inside | 0.8294 | 81 |
|  | QLPQGTTLPK | 160 | 169 | 10 | Outside | 0.1299 | 81 |
|  | GTTLPKGFY | 164 | 172 | 9 | Outside | 0.0225 | 81 |
|  | RIRGGDGKMK | 93 | 102 | 10 | Inside | 0.6300 | 27 |
|  | NPANNAAIV | 150 | 158 | 9 | Inside | 0.5483 | 81 |
|  | GTGPEAGLPY | 114 | 123 | 10 | Outside | 0.0380 | 27 |
|  | LQLPQGTTL | 159 | 167 | 9 | Outside | 0.0480 | 81 |
|  | DLSPRWYFY | 103 | 111 | 9 | Outside | 1.7645 | 27 |
|  | GTRNPANNA | 147 | 155 | 9 | Inside | -0.0002 | 81 |
|  | HGKEDLKFPR | 59 | 68 | 10 | Inside | 0.6273 | 27 |
|  | DDQIGYYRR | 81 | 89 | 9 | Inside | 0.8473 | 81 |
|  | RWYFYYLGTG | 107 | 116 | 10 | Inside | 1.1548 | 27 |
|  | YLGTGPEAGL | 112 | 121 | 10 | Outside | 0.6066 | 27 |
|  | SSRSSSRSR | 183 | 191 | 9 | Inside | 1.2286 | 81 |
|  | LPQGTTLPK | 161 | 169 | 9 | Outside | 0.4172 | 81 |
|  | NTPKDHIGTR | 140 | 149 | 10 | Inside | -0.5461 | 27 |
|  | SPDDQIGYY | 79 | 87 | 9 | Outside | 0.4863 | 54 |
|  | MKDLSPRWYF | 101 | 110 | 10 | Inside | 0.8708 | 27 |
|  | MSDNGPQNQR | 1 | 10 | 10 | Inside | 0.4968 | 27 |
|  | KQRRPQGLP | 38 | 46 | 9 | Inside | 0.9734 | 81 |
|  | KDLSPRWYF | 102 | 110 | 9 | Outside | 0.8745 | 27 |
|  | SSPDDQIGY | 78 | 86 | 9 | Outside | 0.5260 | 27 |
|  | KDHIGTRNPA | 143 | 152 | 10 | Inside | 0.1216 | 27 |
|  | TPKDHIGTR | 141 | 149 | 9 | Inside | -0.1284 | 81 |
|  | TGPEAGLPY | 115 | 123 | 9 | Outside | -0.0349 | 54 |
|  | SKQRRPQGL | 37 | 45 | 9 | Inside | 0.5547 | 27 |
|  | AGLPYGANK | 119 | 127 | 9 | Outside | 0.2631 | 81 |
|  | RIRGGDGKM | 93 | 101 | 9 | Inside | 0.1803 | 81 |
|  | YYLGTGPEA | 111 | 119 | 9 | Outside | 0.7969 | 81 |
|  | PQNQRNAPR | 6 | 14 | 9 | Inside | 0.1594 | 81 |
|  | RPQGLPNNT | 41 | 49 | 9 | Inside | 0.5758 | 54 |
|  | RSSSRSRNS | 185 | 193 | 9 | Inside | 1.2101 | 54 |
|  | STGSNQNGER | 23 | 32 | 10 | Inside | 0.0839 | 27 |
|  | TQHGKEDLKF | 57 | 66 | 10 | Inside | 1.6460 | 27 |
|  | ATRRIRGGD | 90 | 98 | 9 | Inside | 1.1408 | 81 |
|  | APRITFGGP | 12 | 20 | 9 | Outside | 0.4775 | 54 |
|  | GLPNNTASW | 44 | 52 | 9 | Outside | 0.1359 | 54 |
|  | PYGANKDGI | 122 | 130 | 9 | Outside | -0.2279 | 54 |
|  | AEGSRGGSQA | 173 | 182 | 10 | Inside | 0.7171 | 27 |
|  | TTLPKGFYA | 165 | 173 | 9 | Outside | -0.3102 | 54 |
|  | LTQHGKEDLK | 56 | 65 | 10 | Inside | 1.0003 | 27 |
|  | WYFYYLGTGP | 108 | 117 | 10 | Outside | 1.3578 | 27 |
|  | RGGSQASSR | 177 | 185 | 9 | Inside | 0.8611 | 81 |
|  | KGFYAEGSR | 169 | 177 | 9 | Inside | -0.3722 | 81 |
|  | RNPANNAAI | 149 | 157 | 9 | Inside | -0.0357 | 54 |
|  | YRRATRRIR | 87 | 95 | 9 | Inside | -0.2931 | 54 |
|  | TQHGKEDLK | 57 | 65 | 9 | Inside | 1.2574 | 27 |
|  | ASWFTALTQ | 50 | 58 | 9 | Inside | 0.1624 | 81 |
|  | GERSGARSK | 30 | 38 | 9 | Inside | 0.8862 | 81 |
|  | DGKMKDLSP | 98 | 106 | 9 | Outside | 1.7554 | 54 |
|  | RATRRIRGG | 89 | 97 | 9 | Inside | 0.8588 | 54 |
|  | TEGALNTPK | 135 | 143 | 9 | Inside | -0.5316 | 54 |
|  | RWYFYYLGT | 107 | 115 | 9 | Inside | 0.8640 | 54 |
|  | NTNSSPDDQI | 75 | 84 | 10 | Outside | 0.4913 | 27 |
|  | VPINTNSSPD | 72 | 81 | 10 | Outside | 0.4588 | 27 |
|  | GKEDLKFPR | 60 | 68 | 9 | Inside | 0.7725 | 27 |
|  | SGARSKQRR | 33 | 41 | 9 | Inside | 0.7581 | 54 |
|  | QNGERSGAR | 28 | 36 | 9 | Inside | 0.1301 | 54 |
|  | TGSNQNGER | 24 | 32 | 9 | Inside | -0.0721 | 54 |
|  | QHGKEDLKF | 58 | 66 | 9 | Inside | 1.5980 | 54 |
|  | LPYGANKDG | 121 | 129 | 9 | Outside | 0.0639 | 54 |
|  | TASWFTALT | 49 | 57 | 9 | Outside | 0.2451 | 27 |
|  | VPINTNSSP | 72 | 80 | 9 | Outside | 0.4439 | 54 |
|  | IWVATEGAL | 131 | 139 | 9 | Outside | 0.0457 | 81 |
|  | KEDLKFPRGQ | 61 | 70 | 10 | Inside | 0.0110 | 27 |
|  | GSRGGSQAS | 175 | 183 | 9 | Inside | 0.9518 | 81 |
|  | LPKGFYAEG | 167 | 175 | 9 | Outside | -0.5997 | 81 |
|  | SQASSRSSS | 180 | 188 | 9 | Inside | 0.8519 | 54 |
|  | AAIVLQLPQG | 155 | 164 | 10 | Outside | 0.0620 | 27 |
|  | NNAAIVLQL | 153 | 161 | 9 | Inside | 0.8662 | 81 |
|  | QRNAPRITF | 9 | 17 | 9 | Inside | 0.4654 | 54 |
|  | SWFTALTQHG | 51 | 60 | 10 | Outside | 0.1054 | 27 |
|  | GIIWVATEGA | 129 | 138 | 10 | Inside | 0.3353 | 27 |
|  | WYFYYLGTG | 108 | 116 | 9 | Outside | 1.4829 | 27 |
|  | ALNTPKDHI | 138 | 146 | 9 | Inside | -0.8423 | 81 |
|  | LKFPRGQGV | 64 | 72 | 9 | Outside | -0.3703 | 54 |
|  | FYYLGTGPE | 110 | 118 | 9 | Outside | 1.1904 | 54 |
|  | GKMKDLSPR | 99 | 107 | 9 | Inside | 1.3435 | 27 |
|  | MKDLSPRWY | 101 | 109 | 9 | Inside | 0.8441 | 27 |
|  | RNAPRITFG | 10 | 18 | 9 | Inside | 0.8029 | 54 |
|  | NAPRITFGG | 11 | 19 | 9 | Inside | 0.5110 | 27 |
|  | IIWVATEGA | 130 | 138 | 9 | Inside | 0.5150 | 27 |
|  | RITFGGPSD | 14 | 22 | 9 | Outside | 0.8689 | 81 |
|  | GARSKQRRP | 34 | 42 | 9 | Inside | 0.8316 | 81 |
|  | SDNGPQNQR | 2 | 10 | 9 | Inside | 0.6164 | 54 |
|  | MSDNGPQNQ | 1 | 9 | 9 | Inside | 0.0243 | 27 |
|  | GPSDSTGSNQ | 19 | 28 | 10 | Outside | 0.1739 | 27 |
|  | VLQLPQGTT | 158 | 166 | 9 | Outside | -0.1246 | 54 |
|  | PDDQIGYYR | 80 | 88 | 9 | Inside | 0.0087 | 27 |
|  | ITFGGPSDST | 15 | 24 | 9 | Outside | 0.5142 | 27 |
|  | AEGSRGGSQ | 173 | 181 | 9 | Inside | 0.5985 | 27 |
|  | GPQNQRNAP | 5 | 13 | 9 | Inside | 1.2141 | 54 |
|  | KDGIIWVAT | 127 | 135 | 9 | Inside | -0.0387 | 81 |
|  | NAAIVLQLPQ | 154 | 163 | 10 | Outside | 0.1736 | 27 |
|  | NAAIVLQLP | 154 | 162 | 9 | Outside | 0.2931 | 27 |
|  | TLPKGFYAE | 166 | 174 | 9 | Outside | -0.5507 | 27 |
|  | AAIVLQLPQ | 155 | 163 | 9 | Outside | 0.0848 | 27 |
|  | TNSSPDDQIG | 76 | 85 | 10 | Outside | 0.7517 | 27 |
|  | DQIGYYRRA | 82 | 90 | 9 | Inside | 0.4629 | 54 |
|  | GPEAGLPYGA | 116 | 125 | 10 | Outside | -0.2130 | 27 |
|  | YLGTGPEAG | 112 | 120 | 9 | Outside | 0.7971 | 27 |
|  | PQGTTLPKGF | 162 | 171 | 10 | Outside | 0.3012 | 27 |
|  | ITFGGPSDS | 15 | 23 | 9 | Outside | 0.3449 | 27 |
|  | YGANKDGII | 123 | 131 | 9 | Inside | -0.3386 | 27 |
|  | GPSDSTGSN | 19 | 27 | 9 | Outside | 0.0921 | 27 |
|  | NQRNAPRIT | 8 | 16 | 9 | Inside | 0.7931 | 54 |
|  | HIGTRNPAN | 145 | 153 | 9 | Inside | 0.5680 | 81 |
|  | QGLPNNTAS | 43 | 51 | 9 | Outside | 0.0743 | 54 |
|  | NNTASWFTA | 47 | 55 | 9 | Inside | 0.0936 | 54 |
|  | PANNAAIVL | 151 | 159 | 9 | Inside | 0.5440 | 54 |
|  | GTGPEAGLP | 114 | 122 | 9 | Outside | -0.0917 | 54 |
|  | RRIRGGDGK | 92 | 100 | 9 | Inside | 0.2078 | 54 |
|  | NQNGERSGA | 27 | 35 | 9 | Inside | 0.0267 | 54 |
|  | KFPRGQGVP | 65 | 73 | 9 | Outside | -0.1527 | 54 |
|  | PSDSTGSNQ | 20 | 28 | 9 | Outside | 0.1566 | 54 |
|  | WVATEGALN | 132 | 140 | 9 | Outside | 0.3479 | 54 |
|  | RGQGVPINTN | 68 | 77 | 10 | Inside | 0.7769 | 27 |
|  | PEAGLPYGA | 117 | 125 | 9 | Outside | -0.0745 | 54 |
|  | FYAEGSRGG | 171 | 179 | 9 | Outside | 0.4322 | 81 |
|  | ASSRSSSRS | 182 | 190 | 9 | Inside | 0.9269 | 27 |
|  | VATEGALNT | 133 | 141 | 9 | Inside | 0.0276 | 54 |
|  | NTPKDHIGT | 140 | 148 | 9 | Inside | -1.0849 | 54 |
|  | STGSNQNGE | 23 | 31 | 9 | Outside | 0.3895 | 54 |
|  | SWFTALTQH | 51 | 59 | 9 | Outside | 0.1846 | 27 |
|  | LTQHGKEDL | 56 | 64 | 9 | Outside | 0.6157 | 54 |
|  | GPEAGLPYG | 116 | 124 | 9 | Outside | -0.2381 | 27 |
|  | GGSQASSRSS | 178 | 187 | 10 | Inside | 0.6931 | 27 |
|  | DLKFPRGQG | 63 | 71 | 9 | Outside | -0.0028 | 54 |
|  | RGGDGKMKDL | 95 | 104 | 10 | Inside | 0.9272 | 27 |
|  | YFYYLGTGP | 109 | 117 | 9 | Outside | 1.2234 | 27 |
|  | ANKDGIIWV | 125 | 133 | 9 | Inside | -0.2097 | 27 |
|  | QGTTLPKGF | 163 | 171 | 9 | Outside | 0.3392 | 27 |
|  | KEDLKFPRG | 61 | 69 | 9 | Inside | -0.0113 | 54 |
|  | SDSTGSNQNG | 21 | 30 | 10 | Outside | 0.1200 | 27 |
|  | QNQRNAPRI | 7 | 15 | 9 | Inside | 0.5023 | 27 |
|  | TNSSPDDQI | 76 | 84 | 9 | Outside | 0.4559 | 27 |
|  | FGGPSDSTG | 17 | 25 | 9 | Outside | -0.3122 | 81 |
|  | HGKEDLKFP | 59 | 67 | 9 | Outside | 1.1473 | 27 |
|  | DHIGTRNPA | 144 | 152 | 9 | Inside | 0.4313 | 27 |
|  | WFTALTQHG | 52 | 60 | 9 | Outside | 0.1115 | 27 |
|  | RGQGVPINT | 68 | 76 | 9 | inside | 0.7201 | 54 |
|  | GIIWVATEG | 129 | 137 | 9 | Inside | 0.3055 | 54 |
|  | GSNQNGERSG | 25 | 34 | 10 | inside | 0.0715 | 27 |
|  | NKDGIIWVA | 126 | 134 | 9 | inside | -0.0710 | 27 |
|  | QIGYYRRAT | 83 | 91 | 9 | inside | 0.9982 | 27 |
|  | EAGLPYGAN | 118 | 126 | 9 | outside | 0.2692 | 54 |
|  | RRATRRIRG | 88 | 96 | 9 | inside | -0.6094 | 27 |
|  | KDHIGTRNP | 143 | 151 | 9 | inside | 0.2308 | 54 |
|  | QLPQGTTLP | 160 | 168 | 9 | outside | 0.1515 | 54 |
|  | NTNSSPDDQ | 75 | 83 | 9 | outside | 0.8162 | 54 |
|  | DGIIWVATE | 128 | 136 | 9 | inside | -0.1207 | 27 |
|  | EGSRGGSQA | 174 | 182 | 9 | Inside | 0.7712 | 27 |
|  | PRWYFYYLG | 106 | 114 | 9 | outside | 0.9227 | 27 |
|  | NSSPDDQIG | 77 | 85 | 9 | Outside | 0.6147 | 27 |
|  | IVLQLPQGT | 157 | 165 | 9 | outside | -0.1184 | 54 |
|  | ATEGALNTP | 134 | 142 | 9 | Outside | -0.3427 | 27 |
|  | GQGVPINTN | 69 | 77 | 9 | outside | 0.3805 | 27 |
|  | GSQASSRSS | 179 | 187 | 9 | Inside | 0.8687 | 27 |
|  | LGTGPEAGL | 113 | 121 | 9 | outside | 0.5587 | 27 |
|  | GALNTPKDH | 137 | 145 | 9 | outside | -0.5689 | 54 |
|  | TRNPANNAA | 148 | 156 | 9 | Inside | -0.0637 | 27 |
|  | SDSTGSNQN | 21 | 29 | 9 | outside | 0.5273 | 27 |
|  | ARSKQRRPQ | 35 | 43 | 9 | Inside | 0.2003 | 27 |
|  | ANNAAIVLQ | 152 | 160 | 9 | inside | 0.8095 | 27 |
|  | TRRIRGGDG | 91 | 99 | 9 | Inside | 0.6693 | 27 |
|  | QRRPQGLPN | 39 | 47 | 9 | inside | 0.8277 | 54 |
|  | AIVLQLPQG | 156 | 164 | 9 | outside | -0.1279 | 27 |
|  | PNNTASWFT | 46 | 54 | 9 | outside | 0.1113 | 27 |
|  | TALTQHGKE | 54 | 62 | 9 | Inside | 0.6454 | 27 |
|  | QGVPINTNS | 70 | 78 | 9 | inside | 0.3642 | 54 |
|  | YAEGSRGGS | 172 | 180 | 9 | outside | 0.4231 | 27 |
|  | GFYAEGSRG | 170 | 178 | 9 | outside | 0.2811 | 27 |
|  | GSNQNGERS | 25 | 33 | 9 | inside | 0.0657 | 27 |
|  | GVPINTNSS | 71 | 79 | 9 | Outside | 0.2429 | 27 |
|  | LNTPKDHIG | 139 | 147 | 9 | outside | -1.2393 | 27 |
|  | PINTNSSPD | 73 | 81 | 9 | outside | 0.5204 | 27 |
|  | SRGGSQASS | 176 | 184 | 9 | inside | 0.7639 | 27 |
|  | ALTQHGKED | 55 | 63 | 9 | inside | 0.6773 | 27 |
|  | DSTGSNQNG | 22 | 30 | 9 | outside | 0.1687 | 27 |
|  | ERSGARSKQ | 31 | 39 | 9 | inside | 1.2889 | 27 |
|  | IGTRNPANN | 146 | 154 | 9 | Inside | 0.3555 | 27 |
|  | GGSQASSRS | 178 | 186 | 9 | inside | 0.6667 | 27 |
|  | GGDGKMKDL | 96 | 104 | 9 | outside | 1.3319 | 54 |
|  | EGALNTPKD | 136 | 144 | 9 | outside | -0.8109 | 27 |
|  | TFGGPSDST | 16 | 24 | 9 | outside | 0.0722 | 27 |
|  | PQGLPNNTA | 42 | 50 | 9 | outside | 0.1067 | 27 |
|  | RRPQGLPNN | 40 | 48 | 9 | inside | 0.2580 | 27 |
|  | SRSSSRSRN | 184 | 192 | 9 | inside | 1.4625 | 27 |
|  | INTNSSPDD | 74 | 82 | 9 | outside | 0.4458 | 27 |
|  | GLPYGANKD | 120 | 128 | 9 | outside | 0.2865 | 27 |
|  | RGGDGKMKD | 95 | 103 | 9 | inside | 0.8805 | 54 |
|  | IRGGDGKMK | 94 | 102 | 9 | inside | 0.7307 | 27 |
|  | EDLKFPRGQ | 62 | 70 | 9 | outside | -0.1378 | 27 |
|  | SNQNGERSG | 26 | 34 | 9 | inside | 0.2165 | 27 |
|  | PRITFGGPS | 13 | 21 | 9 | outside | 0.7925 | 27 |
|  | PQGTTLPKG | 162 | 171 | 9 | Outside | 0.4960 | 27 |
|  | NGPQNQRNA | 4 | 12 | 9 | Inside | 0.4649 | 54 |
|  | NGERSGARS | 29 | 37 | 9 | inside | 0.2020 | 27 |
|  | GDGKMKDLS | 97 | 105 | 9 | Outside | 1.4231 | 27 |
|  | GGPSDSTGS | 18 | 26 | 9 | outside | 0.1074 | 27 |
|  | DNGPQNQRN | 3 | 11 | 9 | inside | 0.6849 | 27 |
|  | PRGQGVPIN | 67 | 75 | 9 | outside | 1.1707 | 27 |
|  | PKDHIGTRN | 142 | 150 | 9 | inside | 0.1000 | 27 |
|  | PKGFYAEGS | 168 | 176 | 9 | outside | -0.3486 | 27 |
|  | MEVTPSGTW | 128 | 136 | 9 | outside | 0.7550 | 81 |
|  | KPRQKRTAT | 63 | 71 | 9 | Inside | 0.2029 | 81 |
|  | KTFPPTEPK | 167 | 175 | 9 | outside | 0.7571 | 81 |
|  | AQFAPSASAF | 111 | 120 | 10 | outside | 0.5986 | 27 |
|  | RQKRTATKAY | 65 | 74 | 10 | inside | 0.2928 | 27 |
|  | KAYNVTQAF | 72 | 80 | 9 | inside | 0.5669 | 81 |
|  | TPSGTWLTY | 131 | 139 | 9 | Outside | 0.1452 | 81 |
|  | QELIRQGTDY | 95 | 104 | 10 | inside | -0.3589 | 27 |
|  | LLNKHIDAY | 158 | 166 | 9 | inside | -0.3003 | 81 |
|  | SASAFFGMSR | 116 | 125 | 10 | outside | 0.3129 | 27 |
|  | ASAFFGMSR | 117 | 125 | 9 | outside | 0.3153 | 54 |
|  | ELIRQGTDY | 96 | 104 | 9 | inside | -0.2497 | 54 |
|  | FAPSASAFF | 113 | 121 | 9 | outside | 0.2642 | 81 |
|  | LLLDRLNQL | 28 | 36 | 9 | outside | 0.1566 | 81 |
|  | KSAAEASKK | 55 | 63 | 9 | inside | 0.7679 | 81 |
|  | GMSRIGMEV | 122 | 130 | 9 | inside | 0.6287 | 81 |
|  | LPAADLDDF | 201 | 209 | 9 | outside | 0.1864 | 81 |
|  | KQQTVTLLPA | 194 | 203 | 10 | inside | 0.7056 | 27 |
|  | DPNFKDQVI | 149 | 157 | 9 | inside | 1.7367 | 81 |
|  | ASKKPRQKR | 60 | 68 | 9 | inside | -0.1717 | 81 |
|  | LIRQGTDYK | 97 | 105 | 9 | inside | 0.0836 | 81 |
|  | WPQIAQFAP | 107 | 115 | 9 | outside | 0.6110 | 81 |
|  | AYNVTQAFGR | 73 | 82 | 10 | inside | 0.0315 | 27 |
|  | NVTQAFGRR | 75 | 83 | 9 | inside | 0.0331 | 81 |
|  | AQFAPSASA | 111 | 119 | 9 | outside | 0.7468 | 54 |
|  | QQTVTLLPAA | 195 | 204 | 10 | inside | 0.6369 | 27 |
|  | APSASAFFGM | 114 | 123 | 9 | outside | 0.3290 | 27 |
|  | YNVTQAFGR | 74 | 82 | 9 | inside | -0.1100 | 27 |
|  | FSKQLQQSM | 209 | 217 | 9 | inside | 0.1526 | 81 |
|  | RTATKAYNV | 68 | 76 | 9 | inside | 0.6395 | 81 |
|  | KLDDKDPNFK | 144 | 153 | 10 | inside | 2.1298 | 27 |
|  | KHIDAYKTF | 181 | 189 | 9 | inside | -0.3081 | 81 |
|  | SSRGTSPAR | 7 | 15 | 9 | inside | 1.2455 | 81 |
|  | RQKRTATKA | 65 | 73 | 9 | inside | 0.3937 | 54 |
|  | TWLTYTGAI | 135 | 143 | 9 | inside | 0.5439 | 81 |
|  | QFAPSASAF | 112 | 120 | 9 | outside | 0.5495 | 27 |
|  | QLESKMSGK | 35 | 43 | 9 | inside | 1.2535 | 81 |
|  | MAGNGGDAAL | 16 | 25 | 10 | outside | 0.3845 | 27 |
|  | LALLLLDRL | 25 | 33 | 9 | outside | 0.5933 | 81 |
|  | SAFFGMSRI | 118 | 126 | 9 | outside | 0.4244 | 54 |
|  | SAAEASKKPR | 56 | 65 | 10 | inside | 0.8859 | 27 |
|  | MAGNGGDAA | 16 | 24 | 9 | outside | 0.4562 | 54 |
|  | MSSADSTQA | 217 | 225 | 9 | inside | 0.6556 | 54 |
|  | EVTPSGTWL | 129 | 137 | 9 | outside | 0.4548 | 54 |
|  | VTKKSAAEA | 52 | 60 | 9 | inside | 0.5016 | 81 |
|  | RMAGNGGDA | 15 | 23 | 9 | inside | 0.3914 | 54 |
|  | SPARMAGNGG | 12 | 21 | 10 | inside | 0.2807 | 27 |
|  | QLQQSMSSA | 212 | 220 | 9 | inside | 0.3180 | 81 |
|  | GTDYKHWPQI | 101 | 110 | 10 | inside | 0.8032 | 27 |
|  | YKHWPQIAQF | 104 | 113 | 10 | inside | 0.3388 | 27 |
|  | RQGTDYKHW | 99 | 107 | 9 | inside | 0.5721 | 81 |
|  | KHWPQIAQF | 105 | 113 | 9 | inside | 0.2970 | 54 |
|  | RQKKQQTVTL | 191 | 200 | 10 | inside | 0.7542 | 27 |
|  | AALALLLLDR | 23 | 32 | 10 | outside | 0.7534 | 27 |
|  | RNSTPGSSR | 1 | 9 | 9 | inside | 0.7535 | 54 |
|  | KKSAAEASK | 54 | 62 | 9 | inside | 0.4284 | 54 |
|  | QQTVTLLPA | 195 | 203 | 9 | inside | 0.6671 | 27 |
|  | AADLDDFSK | 203 | 211 | 9 | outside | 0.0062 | 81 |
|  | LNKHIDAYK | 159 | 167 | 9 | inside | -0.9872 | 54 |
|  | AEASKKPRQK | 58 | 67 | 10 | inside | 0.5889 | 27 |
|  | QQQGQTVTK | 46 | 54 | 9 | inside | 0.6709 | 81 |
|  | KDQVILLNK | 153 | 161 | 9 | inside | 0.6619 | 81 |
|  | DAALALLLL | 22 | 30 | 9 | outside | 0.6571 | 81 |
|  | KQLQQSMSS | 211 | 219 | 9 | inside | 0.2082 | 54 |
|  | TQALPQRQK | 185 | 193 | 9 | inside | 0.7432 | 81 |
|  | AAEASKKPR | 57 | 65 | 9 | inside | 1.0129 | 54 |
|  | NFGDQELIR | 91 | 99 | 9 | inside | 0.3437 | 81 |
|  | GTSPARMAGN | 10 | 19 | 10 | inside | 0.2425 | 27 |
|  | LQQSMSSADS | 213 | 222 | 10 | outside | 0.3423 | 27 |
|  | KLDDKDPNF | 144 | 152 | 9 | outside | 2.6591 | 54 |
|  | YTGAIKLDDK | 139 | 148 | 10 | inside | 1.3094 | 27 |
|  | ATKAYNVTQA | 70 | 79 | 10 | inside | 0.7630 | 27 |
|  | QTVTLLPAA | 196 | 204 | 9 | outside | 0.6871 | 54 |
|  | GQQQQGQTV | 44 | 52 | 9 | inside | 0.3323 | 81 |
|  | GDAALALLL | 21 | 29 | 9 | outside | 0.4529 | 54 |
|  | WLTYTGAIK | 136 | 144 | 9 | inside | 0.4074 | 54 |
|  | ALALLLLDR | 24 | 32 | 9 | outside | 0.8120 | 27 |
|  | GPEQTQGNF | 84 | 92 | 9 | outside | 0.7349 | 81 |
|  | DAYKTFPPT | 164 | 172 | 9 | outside | 0.1836 | 81 |
|  | EASKKPRQK | 59 | 67 | 9 | inside | 0.3859 | 27 |
|  | QALPQRQKK | 186 | 194 | 9 | inside | 1.0859 | 54 |
|  | ILLNKHIDA | 157 | 165 | 9 | inside | 0.1121 | 54 |
|  | RLNQLESKM | 32 | 40 | 9 | inside | 0.5632 | 81 |
|  | IAQFAPSAS | 110 | 118 | 9 | outside | 0.1755 | 54 |
|  | PQIAQFAPSA | 108 | 117 | 10 | outside | 0.4549 | 27 |
|  | MSRIGMEVT | 123 | 131 | 9 | inside | 1.2800 | 54 |
|  | SMSSADSTQ | 216 | 224 | 9 | inside | 0.7364 | 54 |
|  | QTQGNFGDQ | 87 | 95 | 9 | inside | 1.5902 | 81 |
|  | DYKHWPQIA | 103 | 111 | 9 | inside | 0.9806 | 81 |
|  | LTYTGAIKL | 137 | 145 | 9 | inside | 0.6524 | 54 |
|  | LQQSMSSAD | 213 | 221 | 9 | inside | 0.5110 | 27 |
|  | KQQTVTLLP | 194 | 202 | 9 | inside | 0.8116 | 54 |
|  | RQKKQQTVT | 191 | 199 | 9 | inside | 0.6051 | 54 |
|  | AYKTFPPTEP | 165 | 174 | 9 | outside | 0.4470 | 27 |
|  | TPGSSRGTS | 4 | 12 | 9 | inside | -0.0319 | 81 |
|  | TYTGAIKLD | 138 | 146 | 9 | inside | 0.4307 | 54 |
|  | QKKQQTVTL | 192 | 200 | 9 | inside | 0.9285 | 54 |
|  | GSSRGTSPA | 6 | 14 | 9 | outside | 0.5500 | 54 |
|  | TDYKHWPQI | 102 | 110 | 9 | inside | 0.7426 | 27 |
|  | FPPTEPKKD | 169 | 177 | 9 | outside | 1.0650 | 81 |
|  | AEASKKPRQ | 58 | 66 | 9 | inside | 0.8511 | 27 |
|  | QSMSSADST | 215 | 223 | 9 | inside | 0.6446 | 54 |
|  | FFGMSRIGM | 120 | 128 | 9 | outside | 0.9199 | 81 |
|  | QQGQTVTKK | 47 | 55 | 9 | inside | 0.8224 | 54 |
|  | LLDRLNQLES | 29 | 38 | 10 | outside | 0.1577 | 27 |
|  | LLDRLNQLE | 29 | 37 | 9 | outside | 0.2914 | 27 |
|  | ADLDDFSKQL | 204 | 213 | 10 | outside | -0.0562 | 27 |
|  | PSASAFFGM | 115 | 123 | 9 | outside | 0.3416 | 54 |
|  | ATKAYNVTQ | 70 | 78 | 9 | inside | 0.6635 | 54 |
|  | RGTSPARMAG | 9 | 18 | 10 | inside | 0.7592 | 27 |
|  | AALALLLLD | 23 | 31 | 9 | outside | 0.6504 | 27 |
|  | AYKTFPPTE | 23 | 31 | 9 | outside | 0.5694 | 27 |
|  | DETQALPQR | 183 | 191 | 9 | inside | 0.4122 | 81 |
|  | APSASAFFG | 114 | 122 | 9 | outside | 0.1254 | 27 |
|  | KMSGKGQQQQ | 39 | 48 | 10 | inside | 1.1707 | 27 |
|  | KKQQTVTLL | 193 | 201 | 9 | inside | 0.7709 | 27 |
|  | QIAQFAPSA | 109 | 117 | 9 | inside | 0.4353 | 27 |
|  | KMSGKGQQQ | 39 | 47 | 9 | inside | 1.4257 | 54 |
|  | QTVTKKSAA | 50 | 58 | 9 | inside | 0.3891 | 81 |
|  | RGTSPARMA | 9 | 17 | 9 | inside | 1.2953 | 54 |
|  | QVILLNKHI | 155 | 163 | 9 | inside | 0.3492 | 81 |
|  | TGAIKLDDK | 140 | 148 | 9 | inside | 1.6407 | 54 |
|  | AYNVTQAFG | 73 | 81 | 9 | inside | 0.2807 | 27 |
|  | PSGTWLTYT | 132 | 140 | 9 | outside | 0.3799 | 54 |
|  | QQQQGQTVT | 45 | 53 | 9 | inside | 0.4252 | 27 |
|  | QAFGRRGPE | 78 | 86 | 9 | inside | 0.6906 | 81 |
|  | PQIAQFAPS | 108 | 116 | 9 | outside | 0.3861 | 27 |
|  | AFFGMSRIG | 109 | 117 | 9 | outside | 0.9208 | 27 |
|  | AYKTFPPTE | 165 | 173 | 9 | outside | 0.5694 | 27 |
|  | VTLLPAADL | 198 | 206 | 9 | outside | 0.6417 | 81 |
|  | HWPQIAQFA | 106 | 114 | 9 | inside | 0.4995 | 27 |
|  | VTPSGTWLT | 130 | 138 | 9 | outside | 0.0505 | 27 |
|  | FGMSRIGME | 121 | 129 | 9 | outside | 0.9467 | 27 |
|  | KGQQQQGQT | 43 | 51 | 9 | inside | 0.0803 | 54 |
|  | AGNGGDAAL | 17 | 25 | 9 | outside | 0.5577 | 54 |
|  | LESKMSGKG | 36 | 44 | 9 | inside | 1.5254 | 54 |
|  | SASAFFGMS | 116 | 124 | 9 | outside | 0.4967 | 27 |
|  | NFKDQVILL | 151 | 159 | 9 | outside | 1.1677 | 81 |
|  | HIDAYKTFPP | 162 | 171 | 10 | outside | 0.1578 | 27 |
|  | RIGMEVTPS | 125 | 133 | 9 | inside | 1.5314 | 81 |
|  | QKRTATKAY | 66 | 74 | 9 | inside | -0.0777 | 54 |
|  | KKADETQAL | 180 | 188 | 9 | inside | 0.4135 | 81 |
|  | FKDQVILLN | 152 | 160 | 9 | outside | 0.5640 | 27 |
|  | TFPPTEPKK | 168 | 176 | 9 | outside | 0.9945 | 27 |
|  | KKKADETQA | 179 | 187 | 9 | inside | 0.3995 | 54 |
|  | GTDYKHWPQ | 101 | 109 | 9 | outside | 0.5450 | 54 |
|  | PARMAGNGG | 13 | 21 | 9 | inside | 0.4091 | 54 |
|  | QRQKKQQTV | 190 | 198 | 9 | inside | 0.5602 | 54 |
|  | TQGNFGDQEL | 88 | 97 | 10 | outside | 1.7322 | 27 |
|  | LPQRQKKQQ | 188 | 196 | 9 | inside | 1.0046 | 81 |
|  | SPARMAGNG | 12 | 20 | 9 | inside | -0.0249 | 54 |
|  | MSGKGQQQQ | 40 | 48 | 9 | inside | 1.0288 | 54 |
|  | GGDAALALL | 20 | 28 | 9 | outside | 0.3724 | 54 |
|  | IGMEVTPSG | 126 | 134 | 9 | outside | 1.3812 | 54 |
|  | NGGDAALAL | 19 | 27 | 9 | outside | 0.6302 | 54 |
|  | SKKPRQKRTA | 61 | 70 | 10 | inside | 0.0387 | 27 |
|  | TVTLLPAAD | 197 | 205 | 9 | outside | 0.5896 | 27 |
|  | QGNFGDQELI | 89 | 98 | 10 | outside | 1.1386 | 27 |
|  | SKKPRQKRT | 61 | 69 | 9 | inside | -0.2668 | 27 |
|  | VTQAFGRRG | 76 | 84 | 9 | inside | 0.5301 | 54 |
|  | ETQALPQRQ | 184 | 192 | 10 | inside | 0.7178 | 27 |
|  | EPKKDKKKKA | 173 | 182 | 10 | inside | -0.2542 | 27 |
|  | GTWLTYTGA | 134 | 142 | 9 | outside | 0.6281 | 54 |
|  | GDQELIRQG | 93 | 101 | 9 | inside | -0.2476 | 81 |
|  | NSTPGSSRGT | 2 | 11 | 10 | inside | 0.3678 | 27 |
|  | PTEPKKDKK | 171 | 179 | 9 | inside | 0.7945 | 27 |
|  | GTSPARMAG | 10 | 18 | 9 | inside | 0.4822 | 27 |
|  | SGKGQQQQG | 41 | 49 | 9 | inside | 0.6482 | 54 |
|  | TVTKKSAAE | 51 | 59 | 9 | inside | 0.5090 | 27 |
|  | LDDKDPNFK | 145 | 153 | 9 | outside | 1.9433 | 54 |
|  | QELIRQGTD | 95 | 103 | 9 | inside | -0.4497 | 54 |
|  | QQSMSSADS | 214 | 222 | 9 | inside | 0.5061 | 27 |
|  | GNFGDQELI | 90 | 98 | 9 | outside | 1.1888 | 27 |
|  | TKAYNVTQA | 71 | 79 | 9 | inside | 0.7210 | 27 |
|  | PEQTQGNFG | 85 | 93 | 9 | outside | 1.2090 | 54 |
|  | DQVILLNKH | 154 | 162 | 9 | inside | 0.5069 | 27 |
|  | DRLNQLESK | 31 | 39 | 9 | inside | 0.4998 | 54 |
|  | KADETQALP | 181 | 189 | 9 | inside | 0.4487 | 54 |
|  | AFGRRGPEQ | 79 | 87 | 9 | inside | 0.8208 | 54 |
|  | NQLESKMSG | 34 | 42 | 9 | inside | 0.4574 | 54 |
|  | STPGSSRGT | 3 | 11 | 9 | outside | -0.0962 | 27 |
|  | TQGNFGDQE | 88 | 96 | 9 | outside | 1.8694 | 27 |
|  | SRIGMEVTP | 124 | 132 | 9 | inside | 1.6178 | 27 |
|  | NSTPGSSRG | 2 | 10 | 9 | outside | 0.1617 | 27 |
|  | TATKAYNVT | 69 | 77 | 9 | inside | 0.7877 | 27 |
|  | QGNFGDQEL | 89 | 97 | 9 | outside | 1.6879 | 27 |
|  | DDKDPNFKD | 146 | 154 | 9 | outside | 1.8484 | 54 |
|  | QGQTVTKKSA | 48 | 57 | 10 | inside | 0.7648 | 27 |
|  | GMEVTPSGT | 127 | 135 | 9 | outside | 1.0873 | 27 |
|  | GAIKLDDKDP | 141 | 150 | 10 | outside | 1.9075 | 27 |
|  | IRQGTDYKH | 98 | 106 | 9 | inside | 0.7166 | 27 |
|  | YTGAIKLDD | 139 | 147 | 9 | inside | 0.6986 | 27 |
|  | SKMSGKGQQ | 38 | 46 | 9 | inside | 1.3002 | 54 |
|  | SRGTSPARM | 8 | 16 | 9 | inside | 1.1235 | 27 |
|  | TSPARMAGN | 11 | 19 | 9 | inside | -0.1475 | 27 |
|  | ALLLLDRLN | 26 | 34 | 9 | outside | 0.7100 | 54 |
|  | HIDAYKTFP | 162 | 170 | 9 | inside | -0.2052 | 27 |
|  | KKPRQKRTA | 62 | 70 | 9 | inside | -0.3895 | 27 |
|  | TQAFGRRGP | 77 | 85 | 9 | inside | 0.3694 | 27 |
|  | TEPKKDKKK | 172 | 180 | 9 | inside | 0.1612 | 54 |
|  | DLDDFSKQ | 205 | 213 | 9 | outside | 0.2063 | 81 |
|  | ESKMSGKGQ | 37 | 45 | 9 | inside | 1.4412 | 27 |
|  | GAIKLDDKD | 141 | 149 | 9 | inside | 2.1285 | 27 |
|  | YKHWPQIAQ | 104 | 112 | 9 | inside | 0.3804 | 27 |
|  | IDAYKTFPP | 163 | 171 | 9 | outside | 0.2654 | 27 |
|  | SAAEASKKP | 56 | 64 | 9 | inside | 0.7465 | 27 |
|  | EPKKDKKKK | 173 | 181 | 9 | inside | -0.4226 | 27 |
|  | SGTWLTYTG | 133 | 141 | 9 | outside | 0.7233 | 27 |
|  | PRQKRTATK | 64 | 72 | 9 | inside | 0.1432 | 27 |
|  | EQTQGNFGD | 96 | 84 | 9 | outside | 1.4392 | 27 |
|  | TLLPAADLD | 199 | 207 | 9 | outside | 0.4015 | 54 |
|  | DFSKQLQQS | 208 | 216 | 9 | inside | 0.2660 | 54 |
|  | GQTVTKKSA | 49 | 57 | 9 | inside | 0.8152 | 27 |
|  | RRGPEQTQG | 82 | 90 | 9 | inside | 0.2451 | 81 |
|  | ARMAGNGGD | 14 | 22 | 9 | inside | 0.4506 | 27 |
|  | LLLLDRLNQ | 27 | 35 | 9 | outside | 0.4527 | 27 |
|  | ADETQALPQ | 182 | 190 | 9 | inside | 0.3670 | 27 |
|  | PNFKDQVIL | 150 | 158 | 9 | outside | 1.3108 | 27 |
|  | PAADLDDFS | 202 | 210 | 9 | outside | 0.1537 | 27 |
|  | RGPEQTQGN | 83 | 91 | 9 | inside | 0.1392 | 27 |
|  | KDPNFKDQV | 148 | 156 | 9 | inside | 1.3338 | 54 |
|  | VILLNKHID | 156 | 164 | 9 | inside | 0.5902 | 27 |
|  | LDRLNQLES | 30 | 38 | 9 | outside | 0.0833 | 27 |
|  | PQRQKKQQT | 189 | 197 | 9 | inside | 0.5997 | 27 |
|  | DDFSKQLQQ | 207 | 215 | 9 | inside | -0.3669 | 54 |
|  | SKQLQQSMS | 210 | 218 | 9 | inside | 0.3690 | 27 |
|  | KRTATKAYN | 67 | 75 | 9 | inside | 0.0948 | 27 |
|  | FGRRGPEQT | 80 | 88 | 9 | inside | 0.8738 | 54 |
|  | PPTEPKKDK | 170 | 178 | 9 | inside | 0.4913 | 27 |
|  | LDDFSKQLQ | 206 | 214 | 9 | outside | -0.1179 | 27 |
|  | KKKKADETQ | 178 | 186 | 9 | inside | 0.6561 | 54 |
|  | KDKKKKADE | 176 | 184 | 9 | inside | 0.2850 | 81 |
|  | FGDQELIRQ | 92 | 100 | 9 | inside | -0.2052 | 27 |
|  | YKTFPPTEP | 166 | 174 | 9 | outside | 0.4772 | 27 |
|  | GRRGPEQTQ | 81 | 89 | 9 | inside | 0.9455 | 27 |
|  | IKLDDKDPN | 143 | 151 | 9 | inside | 2.3118 | 54 |
|  | GNGGDAALA | 18 | 26 | 9 | outside | 0.5946 | 27 |
|  | QGTDYKHWP | 100 | 108 | 9 | outside | 0.9394 | 27 |
|  | LLPAADLDD | 200 | 208 | 9 | outside | 0.4791 | 27 |
|  | PKKDKKKKAD | 174 | 183 | 9 | inside | 0.1322 | 27 |
|  | TKKSAAEAS | 53 | 61 | 9 | inside | 0.6016 | 27 |
|  | PKKDKKKKA | 174 | 182 | 9 | inside | -0.2463 | 27 |
|  | NKHIDAYKT | 160 | 168 | 9 | inside | -0.1554 | 27 |
|  | LNQLESKMS | 33 | 41 | 9 | inside | 0.6213 | 27 |
|  | DKKKKADET | 177 | 185 | 9 | inside | 0.7019 | 27 |
|  | AIKLDDKDP | 142 | 150 | 9 | outside | 2.1670 | 27 |
|  | DQELIRQGT | 94 | 102 | 9 | inside | -0.0794 | 27 |
|  | PGSSRGTSP | 5 | 13 | 9 | outside | 0.0300 | 27 |
|  | QGQTVTKKS | 48 | 56 | 9 | inside | 0.9279 | 27 |
|  | KKDKKKKAD | 175 | 183 | 9 | inside | 0.0586 | 27 |
|  | ALPQRQKKQ | 187 | 195 | 9 | inside | 1.3717 | 27 |
|  | DKDPNFKDQ | 147 | 155 | 9 | inside | 1.8060 | 27 |
|  | GKGQQQQGQ | 42 | 50 | 9 | inside | 0.4438 | 27 |
| HTL Epitopes | NPANNAAIVLQLPQG | 150 | 164 | 15 | 0.2093 | outside | 27 |
|  | PANNAAIVLQLPQGT | 151 | 165 | 15 | 0.1847 | outside | 27 |
|  | RNPANNAAIVLQLPQ | 149 | 163 | 15 | 0.0580 | inside | 27 |
|  | TRNPANNAAIVLQLP | 148 | 162 | 15 | 0.1785 | inside | 27 |
|  | GTRNPANNAAIVLQL | 147 | 161 | 15 | 0.4463 | inside | 27 |
|  | ANNAAIVLQLPQGTT | 152 | 166 | 15 | 0.2745 | outside | 27 |
|  | QIGYYRRATRRIRGG | 83 | 97 | 15 | 0.4614 | inside | 27 |
|  | NNAAIVLQLPQGTTL | 153 | 167 | 15 | 0.3369 | outside | 27 |
|  | DDQIGYYRRATRRIR | 81 | 95 | 15 | 0.3334 | inside | 27 |
|  | DQIGYYRRATRRIRG | 82 | 96 | 15 | -0.0825 | inside | 27 |
|  | GYYRRATRRIRGGDG | 85 | 99 | 15 | 0.1805 | inside | 27 |
|  | IGYYRRATRRIRGGD | 84 | 98 | 15 | 0.6649 | inside | 27 |
|  | SPRWYFYYLGTGPEA | 105 | 119 | 15 | 0.8767 | Outside | 27 |
|  | PRWYFYYLGTGPEAG | 106 | 120 | 15 | 0.8083 | Outside | 27 |
|  | NAAIVLQLPQGTTLP | 154 | 168 | 15 | 0.3104 | Outside | 27 |
|  | RWYFYYLGTGPEAGL | 107 | 121 | 15 | 0.7505 | Outside | 27 |
|  | AAIVLQLPQGTTLPK | 155 | 169 | 15 | 0.2308 | Outside | 27 |
|  | YYRRATRRIRGGDGK | 86 | 100 | 15 | 0.1215 | Inside | 27 |
|  | GIIWVATEGALNTPK | 129 | 143 | 15 | -0.1361 | Outside | 27 |
|  | LSPRWYFYYLGTGPE | 104 | 118 | 15 | 1.3086 | outside | 27 |
|  | WYFYYLGTGPEAGLP | 108 | 122 | 15 | 0.7188 | Outside | 27 |
|  | NKDGIIWVATEGALN | 126 | 140 | 15 | 0.1381 | Inside | 27 |
|  | KDGIIWVATEGALNT | 127 | 141 | 15 | 0.0501 | inside | 27 |
|  | DGIIWVATEGALNTP | 128 | 142 | 15 | 0.3979 | Outside | 27 |
|  | ANKDGIIWVATEGAL | 125 | 139 | 15 | 0.0405 | Inside | 27 |
|  | GANKDGIIWVATEGA | 124 | 138 | 15 | -0.0951 | inside | 27 |
|  | KDLSPRWYFYYLGTG | 102 | 116 | 15 | 1.2051 | outside | 27 |
|  | DLSPRWYFYYLGTGP | 103 | 117 | 15 | 1.5180 | Outside | 27 |
|  | PYGANKDGIIWVATE | 122 | 136 | 15 | 0.0782 | Outside | 27 |
|  | IIWVATEGALNTPKD | 130 | 144 | 15 | -0.0712 | Outside | 27 |
|  | MKDLSPRWYFYYLGT | 101 | 115 | 15 | 1.0027 | Inside | 27 |
|  | YGANKDGIIWVATEG | 123 | 137 | 15 | -0.1378 | Inside | 27 |
|  | PDDQIGYYRRATRRI | 80 | 94 | 15 | -0.0050 | Inside | 27 |
|  | IGTRNPANNAAIVLQ | 146 | 160 | 15 | 0.6867 | Inside | 27 |
|  | YLGTGPEAGLPYGAN | 112 | 126 | 15 | 0.3639 | Outside | 27 |
|  | YYLGTGPEAGLPYGA | 111 | 125 | 15 | 0.3321 | Outside | 27 |
|  | YRRATRRIRGGDGKM | 87 | 101 | 15 | 0.0546 | Inside | 27 |
|  | LGTGPEAGLPYGANK | 113 | 127 | 15 | 0.3163 | Outside | 27 |
|  | APRITFGGPSDSTGS | 12 | 26 | 15 | 0.3620 | Outside | 27 |
|  | NAPRITFGGPSDSTG | 11 | 25 | 15 | 0.1207 | Outside | 27 |
|  | YFYYLGTGPEAGLPY | 109 | 123 | 15 | 0.5906 | Outside | 27 |
|  | PRITFGGPSDSTGSN | 13 | 27 | 15 | 0.4038 | Outside | 27 |
|  | GTGPEAGLPYGANKD | 114 | 128 | 15 | 0.1322 | Outside | 27 |
|  | HIGTRNPANNAAIVL | 145 | 159 | 15 | 0.1322 | Inside | 27 |
|  | TGPEAGLPYGANKDG | 115 | 129 | 15 | -0.1189 | Outside | 27 |
|  | FYYLGTGPEAGLPYG | 110 | 124 | 15 | 0.5263 | Outside | 27 |
|  | RNAPRITFGGPSDST | 10 | 24 | 15 | 0.4234 | Outside | 27 |
|  | KMKDLSPRWYFYYLG | 100 | 114 | 15 | 1.4297 | Inside | 27 |
|  | RITFGGPSDSTGSNQ | 14 | 28 | 15 | 0.5529 | Outside | 27 |
|  | ASWFTALTQHGKEDL | 50 | 64 | 15 | 0.4116 | Outside | 27 |
|  | SWFTALTQHGKEDLK | 51 | 65 | 15 | 0.6641 | outside | 27 |
|  | NTASWFTALTQHGKE | 48 | 62 | 15 | 0.3057 | inside | 27 |
|  | GKMKDLSPRWYFYYL | 99 | 113 | 15 | 1.1625 | inside | 27 |
|  | AIVLQLPQGTTLPKG | 156 | 170 | 15 | 0.1658 | outside | 27 |
|  | IVLQLPQGTTLPKGF | 157 | 171 | 15 | 0.0702 | outside | 27 |
|  | TASWFTALTQHGKED | 49 | 63 | 15 | 0.4491 | inside | 27 |
|  | NNTASWFTALTQHGK | 47 | 61 | 15 | 0.3547 | inside | 27 |
|  | LPYGANKDGIIWVAT | 121 | 135 | 15 | 0.0656 | outside | 27 |
|  | FYAEGSRGGSQASSR | 171 | 185 | 15 | 0.5305 | outside | 27 |
|  | ATRRIRGGDGKMKDL | 90 | 104 | 15 | 0.8359 | inside | 27 |
|  | TRRIRGGDGKMKDLS | 91 | 105 | 15 | 0.9654 | inside | 27 |
|  | WFTALTQHGKEDLKF | 27 | 66 | 15 | 0.9819 | inside | 27 |
|  | LPNNTASWFTALTQH | 45 | 59 | 15 | 0.0509 | outside | 27 |
|  | GLPNNTASWFTALTQ | 44 | 58 | 15 | 0.1365 | outside | 27 |
|  | PNNTASWFTALTQHG | 46 | 60 | 15 | 0.1595 | outside | 27 |
|  | IWVATEGALNTPKDH | 131 | 145 | 15 | -0.3371 | outside | 27 |
|  | RRIRGGDGKMKDLSP | 92 | 106 | 15 | 1.1408 | inside | 27 |
|  | QGLPNNTASWFTALT | 43 | 57 | 15 | 0.1406 | outside | 27 |
|  | GFYAEGSRGGSQASS | 170 | 184 | 15 | 0.4459 | outside | 27 |
|  | DGKMKDLSPRWYFYY | 98 | 112 | 15 | 1.0720 | inside | 27 |
|  | GDGKMKDLSPRWYFY | 97 | 111 | 15 | 1.1013 | outside | 27 |
|  | GGDGKMKDLSPRWYF | 96 | 110 | 15 | 1.1690 | outside | 27 |
|  | GLPYGANKDGIIWVA | 120 | 134 | 15 | 0.0313 | outside | 27 |
|  | RATRRIRGGDGKMKD | 89 | 103 | 15 | 1.1690 | inside | 27 |
|  | GPEAGLPYGANKDGI | 116 | 130 | 15 | -0.2346 | outside | 27 |
|  | PEAGLPYGANKDGII | 117 | 131 | 15 | -0.0574 | outside | 27 |
|  | YAEGSRGGSQASSRS | 172 | 186 | 15 | 0.6466 | inside | 27 |
|  | ITFGGPSDSTGSNQN | 15 | 29 | 15 | 0.5446 | outside | 27 |
|  | DHIGTRNPANNAAIV | 144 | 158 | 15 | 0.4064 | inside | 27 |
|  | IRGGDGKMKDLSPRW | 94 | 108 | 15 | 1.1802 | inside | 27 |
|  | RGGDGKMKDLSPRWY | 95 | 109 | 15 | 1.0419 | inside | 27 |
|  | AGLPYGANKDGIIWV | 119 | 133 | 15 | -0.0329 | outside | 27 |
|  | AEGSRGGSQASSRSS | 173 | 187 | 15 | 0.7822 | outside | 27 |
|  | KGFYAEGSRGGSQAS | 169 | 183 | 15 | 0.3373 | outside | 27 |
|  | EGSRGGSQASSRSSS | 174 | 188 | 15 | 0.7516 | outside | 27 |
|  | TFGGPSDSTGSNQNG | 16 | 30 | 15 | 0.0665 | outside | 27 |
|  | RRATRRIRGGDGKMK | 88 | 102 | 15 | 0.3546 | inside | 27 |
|  | DLKFPRGQGVPINTN | 63 | 77 | 15 | 0.4686 | outside | 27 |
|  | PKGFYAEGSRGGSQA | 168 | 182 | 15 | 0.2173 | outside | 27 |
|  | RPQGLPNNTASWFTA | 41 | 55 | 15 | 0.2735 | inside | 27 |
|  | GSRGGSQASSRSSSR | 175 | 189 | 15 | 0.8928 | inside | 27 |
|  | PQGLPNNTASWFTAL | 42 | 56 | 15 | 0.0349 | outside | 27 |
|  | FTALTQHGKEDLKFP | 53 | 67 | 15 | 0.9591 | inside | 27 |
|  | LKFPRGQGVPINTNS | 64 | 78 | 15 | 0.0207 | outside | 27 |
|  | LPKGFYAEGSRGGSQ | 167 | 181 | 15 | 0.0172 | outside | 27 |
|  | KEDLKFPRGQGVPIN | 61 | 75 | 15 | 0.4931 | outside | 27 |
|  | RRPQGLPNNTASWFT | 40 | 54 | 15 | 0.1417 | outside | 27 |
|  | EDLKFPRGQGVPINT | 62 | 76 | 15 | 0.2878 | outside | 27 |
|  | SPDDQIGYYRRATRR | 79 | 93 | 15 | 0.1636 | inside | 27 |
|  | KFPRGQGVPINTNSS | 65 | 79 | 15 | -0.0094 | outside | 27 |
|  | TLPKGFYAEGSRGGS | 166 | 180 | 15 | -0.0942 | outside | 27 |
|  | KDHIGTRNPANNAAI | 143 | 157 | 15 | 0.1458 | inside | 27 |
|  | GKEDLKFPRGQGVPI | 60 | 74 | 15 | 0.6770 | outside | 27 |
|  | QRRPQGLPNNTASWF | 39 | 53 | 15 | 0.4337 | inside | 27 |
|  | RIRGGDGKMKDLSPR | 93 | 107 | 15 | 1.0728 | inside | 27 |
|  | QRNAPRITFGGPSDS | 9 | 23 | 15 | 0.2555 | inside | 27 |
|  | SRGGSQASSRSSSRS | 176 | 190 | 15 | 0.9478 | inside | 27 |
|  | EAGLPYGANKDGIIW | 118 | 132 | 15 | -0.0813 | outside | 27 |
|  | NQNGERSGARSKQRR | 27 | 41 | 15 | 0.5150 | inside | 27 |
|  | TTLPKGFYAEGSRGG | 165 | 179 | 15 | 0.0131 | outside | 27 |
|  | QNGERSGARSKQRRP | 28 | 42 | 15 | 0.6413 | inside | 27 |
|  | DNGPQNQRNAPRITF | 3 | 17 | 15 | 0.5752 | inside | 27 |
|  | PKDHIGTRNPANNAA | 142 | 156 | 15 | -0.0741 | inside | 27 |
|  | NGPQNQRNAPRITFG | 4 | 18 | 15 | 0.5159 | inside | 27 |
|  | SSPDDQIGYYRRATR | 78 | 92 | 15 | 0.4099 | inside | 27 |
|  | NGERSGARSKQRRPQ | 29 | 43 | 15 | 0.4696 | inside | 27 |
|  | SNQNGERSGARSKQR | 26 | 40 | 15 | 0.7792 | inside | 27 |
|  | FPRGQGVPINTNSSP | 66 | 80 | 15 | 0.5170 | outside | 27 |
|  | GPQNQRNAPRITFGG | 5 | 19 | 15 | 0.8853 | inside | 27 |
|  | QGVPINTNSSPDDQI | 70 | 84 | 15 | 0.4461 | inside | 27 |
|  | SDNGPQNQRNAPRIT | 2 | 16 | 15 | 0.5092 | inside | 27 |
|  | RGGSQASSRSSSRSR | 177 | 191 | 15 | 1.0041 | inside | 27 |
|  | WVATEGALNTPKDHI | 132 | 146 | 15 | -0.3653 | inside | 27 |
|  | ERSGARSKQRRPQGL | 31 | 45 | 15 | 0.6421 | inside | 27 |
|  | GARSKQRRPQGLPNN | 34 | 48 | 15 | 0.5955 | inside | 27 |
|  | GERSGARSKQRRPQG | 30 | 44 | 15 | 0.3890 | inside | 27 |
|  | GQGVPINTNSSPDDQ | 69 | 83 | 15 | 0.5092 | outside | 27 |
|  | RSGARSKQRRPQGLP | 32 | 46 | 15 | 0.8571 | inside | 27 |
|  | SGARSKQRRPQGLPN | 33 | 47 | 15 | 0.5652 | inside | 27 |
|  | PQNQRNAPRITFGGP | 6 | 20 | 15 | 0.5137 | inside | 27 |
|  | QNQRNAPRITFGGPS | 7 | 21 | 15 | 0.6923 | inside | 27 |
|  | TPKDHIGTRNPANNA | 141 | 155 | 15 | -0.0445 | Inside | 27 |
|  | RGQGVPINTNSSPDD | 68 | 82 | 15 | 0.6559 | inside | 27 |
|  | ARSKQRRPQGLPNNT | 35 | 49 | 15 | 0.3825 | Inside | 27 |
|  | GGSQASSRSSSRSRN | 178 | 192 | 15 | 0.9881 | inside | 27 |
|  | PRGQGVPINTNSSPD | 67 | 81 | 15 | 0.7277 | outside | 27 |
|  | GVPINTNSSPDDQIG | 71 | 85 | 15 | 0.5606 | outside | 27 |
|  | INTNSSPDDQIGYYR | 74 | 88 | 15 | 0.3198 | Inside | 27 |
|  | NTNSSPDDQIGYYRR | 75 | 89 | 15 | 0.6832 | Inside | 27 |
|  | PINTNSSPDDQIGYY | 73 | 87 | 15 | 0.5475 | Outside | 27 |
|  | VPINTNSSPDDQIGY | 72 | 86 | 15 | 0.5684 | outside | 27 |
|  | GSQASSRSSSRSRNS | 179 | 193 | 15 | 1.0087 | Inside | 27 |
|  | HGKEDLKFPRGQGVP | 59 | 73 | 15 | 0.5750 | outside | 27 |
|  | PQGTTLPKGFYAEGS | 162 | 176 | 15 | 0.0491 | outside | 27 |
|  | KQRRPQGLPNNTASW | 38 | 52 | 15 | 0.5019 | Inside | 27 |
|  | GTTLPKGFYAEGSRG | 164 | 178 | 15 | 0.0529 | outside | 27 |
|  | VLQLPQGTTLPKGFY | 158 | 172 | 15 | -0.0584 | outside | 27 |
|  | NTPKDHIGTRNPANN | 140 | 154 | 15 | -0.4004 | Inside | 27 |
|  | QGTTLPKGFYAEGSR | 163 | 177 | 15 | -0.0402 | outside | 27 |
|  | TALTQHGKEDLKFPR | 54 | 68 | 15 | 0.6244 | Inside | 27 |
|  | NQRNAPRITFGGPSD | 8 | 22 | 15 | 0.6081 | Inside | 27 |
|  | QHGKEDLKFPRGQGV | 58 | 72 | 15 | 0.4777 | outside | 27 |
|  | GSNQNGERSGARSKQ | 25 | 39 | 15 | 0.5478 | Inside | 27 |
|  | VATEGALNTPKDHIG | 133 | 147 | 15 | -0.4356 | outside | 27 |
|  | MSDNGPQNQRNAPRI | 1 | 15 | 15 | 0.3356 | Inside | 27 |
|  | LPQGTTLPKGFYAEG | 27 | 175 | 15 | -0.0409 | outside | 27 |
|  | QLPQGTTLPKGFYAE | 160 | 174 | 15 | -0.0083 | outside | 27 |
|  | TGSNQNGERSGARSK | 24 | 38 | 15 | 0.5307 | Inside | 27 |
|  | LQLPQGTTLPKGFYA | 159 | 173 | 15 | 0.0056 | outside | 27 |
|  | RSKQRRPQGLPNNTA | 36 | 50 | 15 | 0.3191 | Inside | 27 |
|  | SKQRRPQGLPNNTAS | 37 | 51 | 15 | 0.4611 | Inside | 27 |
|  | DSTGSNQNGERSGAR | 22 | 36 | 15 | 0.1319 | Inside | 27 |
|  | STGSNQNGERSGARS | 23 | 37 | 15 | 0.3387 | Inside | 27 |
|  | ALTQHGKEDLKFPRG | 55 | 69 | 15 | 0.4340 | Inside | 27 |
|  | GGPSDSTGSNQNGER | 18 | 32 | 15 | 0.0269 | outside | 27 |
|  | ATEGALNTPKDHIGT | 134 | 148 | 15 | -0.4671 | Inside | 27 |
|  | LNTPKDHIGTRNPAN | 139 | 153 | 15 | -0.4639 | Inside | 27 |
|  | SDSTGSNQNGERSGA | 21 | 35 | 15 | 0.2041 | outside | 27 |
|  | FGGPSDSTGSNQNGE | 17 | 31 | 15 | 0.0470 | outside | 27 |
|  | LTQHGKEDLKFPRGQ | 56 | 70 | 15 | 0.4310 | Inside | 27 |
|  | NSSPDDQIGYYRRAT | 77 | 91 | 15 | 0.5765 | Inside | 27 |
|  | TQHGKEDLKFPRGQG | 57 | 71 | 15 | 0.4765 | Inside | 27 |
|  | GPSDSTGSNQNGERS | 19 | 33 | 15 | 0.1892 | outside | 27 |
|  | TEGALNTPKDHIGTR | 135 | 149 | 15 | -0.3176 | Inside | 27 |
|  | ALNTPKDHIGTRNPA | 138 | 152 | 15 | -0.3204 | Inside | 27 |
|  | EGALNTPKDHIGTRN | 136 | 150 | 15 | -0.3112 | Inside | 27 |
|  | GALNTPKDHIGTRNP | 137 | 151 | 15 | -0.2763 | Inside | 27 |
|  | PSDSTGSNQNGERSG | 20 | 34 | 15 | 0.1668 | outside | 27 |
|  | AQFAPSASAFFGMSR | 111 | 125 | 15 | 0.5266 | outside | 27 |
|  | IAQFAPSASAFFGMS | 110 | 124 | 15 | 0.3975 | outside | 27 |
|  | PQIAQFAPSASAFFG | 108 | 122 | 15 | 0.3094 | outside | 27 |
|  | QIAQFAPSASAFFGM | 109 | 123 | 15 | 0.4032 | outside | 27 |
|  | WPQIAQFAPSASAFF | 107 | 121 | 15 | 0.3028 | outside | 27 |
|  | GTWLTYTGAIKLDDK | 134 | 148 | 15 | 0.9934 | inside | 27 |
|  | PSGTWLTYTGAIKLD | 132 | 146 | 15 | 0.3466 | outside | 27 |
|  | SGTWLTYTGAIKLDD | 133 | 147 | 15 | 0.6215 | outside | 27 |
|  | TPSGTWLTYTGAIKL | 131 | 145 | 15 | 0.3271 | outside | 27 |
|  | TWLTYTGAIKLDDKD | 135 | 149 | 15 | 1.2416 | inside | 27 |
|  | AALALLLLDRLNQLE | 23 | 27 | 15 | 0.6031 | outside | 27 |
|  | DAALALLLLDRLNQL | 22 | 36 | 15 | 0.5531 | outside | 27 |
|  | ALALLLLDRLNQLES | 24 | 38 | 15 | 0.5057 | outside | 27 |
|  | ATKAYNVTQAFGRRG | 70 | 84 | 15 | 0.7146 | inside | 27 |
|  | LALLLLDRLNQLESK | 25 | 39 | 15 | 0.7357 | outside | 27 |
|  | TATKAYNVTQAFGRR | 69 | 83 | 15 | 0.3835 | inside | 27 |
|  | ALLLLDRLNQLESKM | 26 | 40 | 15 | 0.5669 | outside | 27 |
|  | GDAALALLLLDRLNQ | 21 | 35 | 15 | 0.4458 | outside | 27 |
|  | KAYNVTQAFGRRGPE | 72 | 86 | 15 | 0.6104 | inside | 27 |
|  | RTATKAYNVTQAFGR | 68 | 82 | 15 | 0.2424 | inside | 27 |
|  | TKAYNVTQAFGRRGP | 71 | 85 | 15 | 0.5975 | inside | 27 |
|  | KDQVILLNKHIDAYK | 153 | 167 | 15 | 0.0299 | inside | 27 |
|  | GGDAALALLLLDRLN | 20 | 34 | 15 | 0.4860 | outside | 27 |
|  | DQVILLNKHIDAYKT | 154 | 168 | 15 | 0.0226 | inside | 27 |
|  | MAGNGGDAALALLLL | 16 | 30 | 15 | 0.4852 | outside | 27 |
|  | AFFGMSRIGMEVTPS | 119 | 133 | 15 | 1.1085 | outside | 27 |
|  | APSASAFFGMSRIGM | 114 | 128 | 15 | 0.5987 | outside | 27 |
|  | ASAFFGMSRIGMEVT | 117 | 131 | 15 | 0.8620 | outside | 27 |
|  | PSASAFFGMSRIGME | 115 | 129 | 15 | 0.6408 | outside | 27 |
|  | SAFFGMSRIGMEVTP | 118 | 132 | 15 | 1.0398 | outside | 27 |
|  | SASAFFGMSRIGMEV | 116 | 130 | 15 | 0.6584 | outside | 27 |
|  | HWPQIAQFAPSASAF | 106 | 120 | 15 | 0.4320 | outside | 27 |
|  | QVILLNKHIDAYKTF | 155 | 169 | 15 | -0.0910 | inside | 27 |
|  | HIDAYKTFPPTEPKK | 162 | 176 | 15 | 0.4004 | inside | 27 |
|  | IDAYKTFPPTEPKKD | 163 | 177 | 15 | 0.4427 | inside | 27 |
|  | KHIDAYKTFPPTEPK | 161 | 175 | 15 | 0.3912 | inside | 27 |
|  | NKHIDAYKTFPPTEP | 160 | 174 | 15 | 0.1650 | inside | 27 |
|  | AGNGGDAALALLLLD | 17 | 31 | 15 | 0.5829 | outside | 27 |
|  | GNGGDAALALLLLDR | 18 | 32 | 15 | 0.5897 | outside | 27 |
|  | KKQQTVTLLPAADLD | 193 | 207 | 15 | 0.7325 | inside | 27 |
|  | QKKQQTVTLLPAADL | 192 | 206 | 15 | 0.7662 | inside | 27 |
|  | RQKKQQTVTLLPAAD | 191 | 205 | 15 | 0.6555 | inside | 27 |
|  | LTYTGAIKLDDKDPN | 137 | 151 | 15 | 1.4693 | inside | 27 |
|  | FKDQVILLNKHIDAY | 152 | 166 | 15 | 0.2669 | inside | 27 |
|  | NGGDAALALLLLDRL | 19 | 33 | 15 | 0.3946 | outside | 27 |
|  | QRQKKQQTVTLLPAA | 190 | 204 | 15 | 0.6824 | inside | 27 |
|  | WLTYTGAIKLDDKDP | 136 | 150 | 15 | 1.2787 | outside | 27 |
|  | KHWPQIAQFAPSASA | 105 | 119 | 15 | 0.4293 | inside | 27 |
|  | AYNVTQAFGRRGPEQ | 73 | 87 | 15 | 0.3870 | inside | 27 |
|  | GAIKLDDKDPNFKDQ | 141 | 155 | 15 | 1.9373 | inside | 27 |
|  | TGAIKLDDKDPNFKD | 140 | 154 | 15 | 1.8513 | inside | 27 |
|  | TYTGAIKLDDKDPNF | 138 | 152 | 15 | 1.6988 | inside | 27 |
|  | YTGAIKLDDKDPNFK | 139 | 153 | 15 | 1.6136 | inside | 27 |
|  | QQTVTLLPAADLDDF | 195 | 209 | 15 | 0.4614 | outside | 27 |
|  | DPNFKDQVILLNKHI | 149 | 163 | 15 | 1.1072 | outside | 27 |
|  | QTVTLLPAADLDDFS | 196 | 210 | 15 | 0.5213 | outside | 27 |
|  | KQQTVTLLPAADLDD | 194 | 208 | 15 | 0.6264 | inside | 27 |
|  | LLLDRLNQLESKMSG | 28 | 42 | 15 | 0.3146 | outside | 27 |
|  | LLLLDRLNQLESKMS | 27 | 41 | 15 | 0.6286 | outside | 27 |
|  | PNFKDQVILLNKHID | 150 | 164 | 15 | 0.9880 | outside | 27 |
|  | VILLNKHIDAYKTFP | 156 | 170 | 15 | 0.0060 | inside | 27 |
|  | DKDPNFKDQVILLNK | 147 | 161 | 15 | 1.1508 | inside | 27 |
|  | KDPNFKDQVILLNKH | 148 | 162 | 15 | 1.1376 | inside | 27 |
|  | QFAPSASAFFGMSRI | 112 | 126 | 15 | 0.4658 | outside | 27 |
|  | FAPSASAFFGMSRIG | 113 | 127 | 15 | 0.6236 | outside | 27 |
|  | YKHWPQIAQFAPSAS | 104 | 118 | 15 | 0.4340 | inside | 27 |
|  | YNVTQAFGRRGPEQT | 74 | 88 | 15 | 0.3806 | inside | 27 |
|  | ARMAGNGGDAALALL | 14 | 28 | 15 | 0.4955 | inside | 27 |
|  | RMAGNGGDAALALL | 15 | 29 | 15 | 0.4735 | inside | 27 |
|  | DDKDPNFKDQVILLN | 146 | 160 | 15 | 1.2508 | outside | 27 |
|  | PARMAGNGGDAALAL | 13 | 27 | 15 | 0.5182 | outside | 27 |
|  | NFKDQVILLNKHIDA | 151 | 165 | 15 | 0.7693 | outside | 27 |
|  | FSKQLQQSMSSADST | 209 | 223 | 15 | 0.3589 | inside | 27 |
|  | SKQLQQSMSSADSTQ | 210 | 224 | 15 | 0.4792 | inside | 27 |
|  | TVTLLPAADLDDFSK | 197 | 211 | 15 | 0.2514 | outside | 27 |
|  | VTLLPAADLDDFSKQ | 198 | 212 | 15 | 0.3389 | outside | 27 |
|  | AIKLDDKDPNFKDQV | 142 | 156 | 15 | 1.7616 | inside | 27 |
|  | LDDKDPNFKDQVILL | 145 | 159 | 15 | 1.4829 | outside | 27 |
|  | IKLDDKDPNFKDQVI | 143 | 157 | 15 | 1.8034 | inside | 27 |
|  | RQKRTATKAYNVTQA | 65 | 79 | 15 | 0.6318 | inside | 27 |
|  | FFGMSRIGMEVTPSG | 120 | 134 | 15 | 0.9397 | outside | 27 |
|  | KQLQQSMSSADSTQA | 211 | 225 | 15 | 0.4771 | inside | 27 |
|  | DFSKQLQQSMSSADS | 208 | 212 | 15 | 0.2999 | inside | 27 |
|  | SRIGMEVTPSGTWLT | 124 | 138 | 15 | 0.7912 | outside | 27 |
|  | ILLNKHIDAYKTFPP | 157 | 171 | 15 | 0.0320 | inside | 27 |
|  | PRQKRTATKAYNVTQ | 64 | 78 | 15 | 0.4981 | inside | 27 |
|  | YNVTQAFGRRGPEQT | 74 | 88 | 15 | 0.3806 | inside | 27 |
|  | GMSRIGMEVTPSGTW | 122 | 136 | 15 | 0.6835 | outside | 27 |
|  | MSRIGMEVTPSGTWL | 123 | 137 | 15 | 0.7397 | outside | 27 |
|  | LNQLESKMSGKGQQQ | 33 | 47 | 15 | 1.0116 | outside | 27 |
|  | LLNKHIDAYKTFPPT | 158 | 172 | 15 | 0.0019 | inside | 27 |
|  | LNKHIDAYKTFPPTE | 159 | 173 | 15 | -0.0811 | inside | 27 |
|  | ALPQRQKKQQTVTLL | 187 | 201 | 15 | 0.8033 | inside | 27 |
|  | DQELIRQGTDYKHWP | 94 | 108 | 15 | 0.2156 | inside | 27 |
|  | GDQELIRQGTDYKHW | 93 | 107 | 15 | 0.0488 | inside | 27 |
|  | RIGMEVTPSGTWLTY | 125 | 139 | 15 | 0.8819 | outside | 27 |
|  | QKRTATKAYNVTQAF | 66 | 80 | 15 | 0.3605 | inside | 27 |
|  | SPARMAGNGGDAALA | 12 | 26 | 15 | 0.3731 | outside | 27 |
|  | FGMSRIGMEVTPSGT | 121 | 135 | 15 | 1.0171 | outside | 27 |
|  | QELIRQGTDYKHWPQ | 95 | 109 | 15 | -0.0543 | inside | 27 |
|  | KKPRQKRTATKAYNV | 62 | 76 | 15 | -0.0382 | inside | 27 |
|  | KPRQKRTATKAYNVT | 63 | 77 | 15 | 0.4122 | inside | 27 |
|  | DRLNQLESKMSGKGQ | 31 | 45 | 15 | 0.8181 | inside | 27 |
|  | LDRLNQLESKMSGKG | 30 | 44 | 15 | 0.7029 | inside | 27 |
|  | LLDRLNQLESKMSGK | 29 | 43 | 15 | 0.5599 | outside | 27 |
|  | QALPQRQKKQQTVTL | 186 | 200 | 15 | 0.7256 | inside | 27 |
|  | TKKSAAEASKKPRQK | 53 | 67 | 15 | 0.6677 | inside | 27 |
|  | VTPSGTWLTYTGAIK | 130 | 144 | 15 | 0.4055 | outside | 27 |
|  | KKSAAEASKKPRQKR | 54 | 68 | 15 | 0.3818 | inside | 27 |
|  | KSAAEASKKPRQKRT | 55 | 59 | 15 | 0.3968 | inside | 27 |
|  | SAAEASKKPRQKRTA | 56 | 70 | 15 | 0.4551 | inside | 27 |
|  | STPGSSRGTSPARMA | 3 | 17 | 15 | 0.4457 | inside | 27 |
|  | DDFSKQLQQSMSSAD | 207 | 221 | 15 | 0.1468 | inside | 27 |
|  | TLLPAADLDDFSKQL | 199 | 213 | 15 | 0.0310 | outside | 27 |
|  | LPQRQKKQQTVTLLP | 188 | 210 | 15 | 0.8232 | inside | 27 |
|  | AAEASKKPRQKRTAT | 57 | 71 | 15 | 0.3565 | inside | 27 |
|  | PQRQKKQQTVTLLPA | 189 | 203 | 15 | 0.6955 | inside | 27 |
|  | EQTQGNFGDQELIRQ | 86 |  | 15 | 0.6565 | inside | 27 |
|  | PEQTQGNFGDQELIR | 85 | 99 | 15 | 0.6270 | outside | 27 |
|  | DAYKTFPPTEPKKDK | 164 | 178 | 15 | 0.3426 | inside | 27 |
|  | FGDQELIRQGTDYKH | 92 | 106 | 15 | 0.3788 | inside | 27 |
|  | QTQGNFGDQELIRQG | 87 | 101 | 15 | 0.8396 | inside | 27 |
|  | GQTVTKKSAAEASKK | 49 | 63 | 15 | 0.6905 | inside | 27 |
|  | NSTPGSSRGTSPARM | 2 | 16 | 15 | 0.5755 | inside | 27 |
|  | PGSSRGTSPARMAGN | 5 | 9 | 15 | 0.1543 | outside | 27 |
|  | QTVTKKSAAEASKKP | 50 | 64 | 15 | 0.5220 | inside | 27 |
|  | TSPARMAGNGGDAAL | 11 | 25 | 15 | 0.3716 | inside | 27 |
|  | TVTKKSAAEASKKPR | 51 | 65 | 15 | 0.8012 | inside | 27 |
|  | KRTATKAYNVTQAFG | 67 | 81 | 15 | 0.2574 | inside | 27 |
|  | RLNQLESKMSGKGQQ | 32 | 46 | 15 | 0.9603 | inside | 27 |
|  | TQGNFGDQELIRQGT | 88 | 102 | 15 | 0.8020 | outside | 27 |
|  | TPGSSRGTSPARMAG | 4 | 18 | 15 | 0.2581 | outside | 27 |
|  | ELIRQGTDYKHWPQI | 96 | 110 | 15 | 0.2209 | inside | 27 |
|  | VTKKSAAEASKKPRQ | 52 | 66 | 15 | 0.7317 | inside | 27 |
|  | QGQTVTKKSAAEASK | 48 | 62 | 15 | 0.5323 | inside | 27 |
|  | EVTPSGTWLTYTGAI | 129 | 143 | 15 | 0.6375 | outside | 27 |
|  | GSSRGTSPARMAGNG | 6 | 20 | 15 | 0.4182 | inside | 27 |
|  | SSRGTSPARMAGNGG | 7 | 21 | 15 | 0.7644 | inside | 27 |
|  | AYKTFPPTEPKKDKK | 165 | 179 | 15 | 0.7524 | inside | 27 |
|  | QGNFGDQELIRQGTD | 89 | 103 | 15 | 0.5806 | outside | 27 |
|  | DYKHWPQIAQFAPSA | 103 | 117 | 15 | 0.6956 | inside | 27 |
|  | QQGQTVTKKSAAEAS | 47 | 61 | 15 | 0.6399 | inside | 27 |
|  | GMEVTPSGTWLTYTG | 127 | 141 | 15 | 0.8311 | outside | 27 |
|  | IGMEVTPSGTWLTYT | 126 | 140 | 15 | 1.0147 | outside | 27 |
|  | SRGTSPARMAGNGGD | 8 | 22 | 15 | 0.6703 | inside | 27 |
|  | ETQALPQRQKKQQTV | 184 | 198 | 15 | 0.7720 | inside | 27 |
|  | TQALPQRQKKQQTVT | 185 | 199 | 15 | 0.6457 | inside | 27 |
|  | LPAADLDDFSKQLQQ | 201 | 215 | 15 | 0.0345 | outside | 27 |
|  | NFGDQELIRQGTDYK | 91 | 105 | 15 | 0.4988 | inside | 27 |
|  | AEASKKPRQKRTATK | 58 | 72 | 15 | 0.0755 | inside | 27 |
|  | MEVTPSGTWLTYTGA | 128 | 142 | 15 | 0.7886 | outside | 27 |
|  | KKKKADETQALPQRQ | 178 | 192 | 15 | 0.7223 | inside | 27 |
|  | LDDFSKQLQQSMSSA | 206 | 220 | 15 | 0.0710 | outside | 27 |
|  | EASKKPRQKRTATKA | 59 | 73 | 15 | 0.0417 | inside | 27 |
|  | GTDYKHWPQIAQFAP | 101 | 115 | 15 | 0.7631 | inside | 27 |
|  | RGPEQTQGNFGDQEL | 83 | 97 | 15 | 1.0160 | outside | 27 |
|  | RQGTDYKHWPQIAQ | 99 | 113 | 15 | 0.5146 | inside | 27 |
|  | KKKADETQALPQRQK | 179 | 193 | 15 | 0.7180 | inside | 27 |
|  | PAADLDDFSKQLQQS | 202 | 216 | 15 | 0.0636 | outside | 27 |
|  | QGTDYKHWPQIAQFA | 100 | 114 | 15 | 0.6625 | inside | 27 |
|  | RGTSPARMAGNGGDA | 9 | 23 | 15 | 0.6743 | inside | 27 |
|  | AADLDDFSKQLQQSM | 203 | 217 | 15 | 0.1905 | outside | 27 |
|  | LLPAADLDDFSKQLQ | 200 | 214 | 15 | 0.1729 | outside | 27 |
|  | TDYKHWPQIAQFAPS | 102 | 116 | 15 | 0.6354 | inside | 27 |
|  | ADLDDFSKQLQQSMS | 204 | 218 | 15 | 0.1923 | outside | 27 |
