## Supplementary Table 5 for "Genome based Evolutionary study of SARS-CoV-2 towards the Prediction of Epitope Based Chimeric Vaccine"

**Supplementary Table 5:** Allergenicity pattern and toxicity profile of top T cell epitopes.

| ***Types*** | ***Protein*** | ***Peptide*** | ***AllergenFP*** | ***AllerTop*** | ***Allermatch*** | ***Allergen Online*** | ***ToxinPred*** |
| --- | --- | --- | --- | --- | --- | --- | --- |
| CTL epitopes | Surface glycoprotein | ADYNYKLPD | Non-allergen | Allergen | Non-allergen | Non-allergen | Nontoxin |
|  | Membrane glycoprotein | GIAIAMACL | Allergen | Non-allergen | Non-allergen | Non-allergen | Nontoxin |
|  |  | LACFVLAAV | Allergen | Non-allergen | Non-allergen | Non-allergen | Nontoxin |
|  |  | LVGLMWLSY | Non-allergen | Non-allergen | Non-allergen | Non-allergen | Nontoxin |
|  |  | LACFVLAAVY | Allergen | Non-allergen | Non-allergen | Non-allergen | Nontoxin |
|  |  | CLVGLMWLSY | Non-allergen | Allergen | Non-allergen | Non-allergen | Nontoxin |
|  |  | CLVGLMWLS | Non-allergen | Allergen | Non-allergen | Non-allergen | Nontoxin |
|  |  | MACLVGLMW | Non-allergen | Non-allergen | Non-allergen | Non-allergen | Nontoxin |
|  |  | VGLMWLSYF | Non-allergen | Allergen | Non-allergen | Non-allergen | Nontoxin |
|  |  | SMWSFNPET | Non-allergen | Non-allergen | Allergen | Non-allergen | Nontoxin |
|  |  | WSFNPETNI | Non-allergen | Allergen | Allergen | Non-allergen | Nontoxin |
|  |  | LVIGAVILR | Non-allergen | Non-allergen | Non-allergen | Non-allergen | Nontoxin |
|  |  | ELVIGAVILR | Non-allergen | Non-allergen | Non-allergen | Non-allergen | Nontoxin |
|  |  | ESELVIGAV | Non-allergen | Non-allergen | Non-allergen | Non-allergen | Nontoxin |
|  |  | IGAVILRGH | Non-allergen | Allergen | Non-allergen | Non-allergen | Nontoxin |
|  |  | LESELVIGA | Non-allergen | Non-allergen | Non-allergen | Non-allergen | Nontoxin |
|  |  | ELVIGAVIL | Non-allergen | Allergen | Non-allergen | Non-allergen | Nontoxin |
|  |  | PLHGTILTR | Non-allergen | Allergen | Non-allergen | Non-allergen | Nontoxin |
|  |  | IGAVILRGHL | Non-allergen | Allergen | Non-allergen | Non-allergen | Nontoxin |
|  |  | LLESELVIG | Non-allergen | Allergen | Non-allergen | Non-allergen | Nontoxin |
|  | Envelope protein | VKPSFYVYS | Allergen | Allergen | Non Allergen | Non Allergen | Nontoxin |
|  |  | FLLVTLAIL | Allergen | Non Allergen | Non Allergen | Non Allergen | Nontoxin |
|  |  | VVFLLVTLA | Allergen | Non Allergen | Non Allergen | Non Allergen | Nontoxin |
|  |  | LAILTALRL | Allergen | Non Allergen | Non Allergen | Non Allergen | Nontoxin |
|  |  | LNSSRVPDL | Allergen | Non Allergen | Non Allergen | Non Allergen | Nontoxin |
|  |  | LLFLAFVVF | Allergen | Non Allergen | Non Allergen | Non Allergen | Nontoxin |
|  |  | VFLLVTLAI | Allergen | Non Allergen | Non Allergen | Non Allergen | Nontoxin |
|  |  | LAFVVFLLV | Allergen | Non Allergen | Non Allergen | Non Allergen | Nontoxin |
|  |  | VSLVKPSFY | Non Allergen | Non Allergen | Non Allergen | Non Allergen | Nontoxin |
|  |  | FVVFLLVTL | Allergen | Non Allergen | Non Allergen | Non Allergen | Nontoxin |
|  |  | VNVSLVKPSF | Allergen | Allergen | Non Allergen | Non Allergen | Nontoxin |
|  |  | YSFVSEETG | Non Allergen | Non Allergen | Non Allergen | Non Allergen | Nontoxin |
|  |  | NVSLVKPSF | Allergen | Non Allergen | Non Allergen | Non Allergen | Nontoxin |
|  |  | AFVVFLLVT | Allergen | Non Allergen | Non Allergen | Non Allergen | Nontoxin |
|  |  | VTLAILTAL | Allergen | Non Allergen | Non Allergen | Non Allergen | Nontoxin |
|  |  | NSSRVPDLL | Allergen | Allergen | Non Allergen | Non Allergen | Nontoxin |
|  |  | PSFYVYSRV | Non Allergen | Non Allergen | Non Allergen | Non Allergen | Nontoxin |
|  |  | YSFVSEETGT | Non Allergen | Allergen | Non Allergen | Non Allergen | Nontoxin |
|  |  | LLVTLAILTA | Allergen | Non Allergen | Non Allergen | Non Allergen | Nontoxin |
|  | Nucleocapsid protein | RSGARSKQR | Allergen | Allergen | Non-allergen | Non-allergen | Nontoxin |
|  |  | DLSPRWYFY | Allergen | Non-allergen | Non-allergen | Non-allergen | Nontoxin |
|  |  | DGKMKDLSP | Non-allergen | Non-allergen | Non-allergen | Non-allergen | Nontoxin |
|  |  | KMKDLSPRW | Allergen | Non-allergen | Non-allergen | Non-allergen | Nontoxin |
|  |  | TQHGKEDLKF | Non-allergen | Non-allergen | Non-allergen | Non-allergen | Nontoxin |
|  |  | QHGKEDLKF | Non-allergen | Non-allergen | Non-allergen | Non-allergen | Nontoxin |
|  |  | KMKDLSPRWY | Allergen | Non-allergen | Non-allergen | Non-allergen | Nontoxin |
|  |  | WYFYYLGTG | Allergen | Non-allergen | Non-allergen | Non-allergen | Nontoxin |
|  |  | SRSSSRSRN | Non-allergen | Allergen | Non-allergen | Non-allergen | Nontoxin |
|  |  | GDGKMKDLS | Non-allergen | Allergen | Non-allergen | Non-allergen | Nontoxin |
|  |  | KLDDKDPNF | Allergen | Allergen | Non-allergen | Non-allergen | Nontoxin |
|  |  | AIKLDDKDP | Allergen | Non-allergen | Non-allergen | Non-allergen | Nontoxin |
|  |  | LDDKDPNFK | Allergen | Non-allergen | Non-allergen | Non-allergen | Nontoxin |
|  |  | GAIKLDDKDP | Non-allergen | Allergen | Non-allergen | Non-allergen | Nontoxin |
|  |  | TQGNFGDQE | Non-allergen | Allergen | Non-allergen | Non-allergen | Nontoxin |
|  |  | DDKDPNFKD | Non-allergen | Non-allergen | Non-allergen | Non-allergen | Nontoxin |
|  |  | TQGNFGDQEL | Non-allergen | Allergen | Non-allergen | Non-allergen | Nontoxin |
|  |  | QGNFGDQEL | Non-allergen | Allergen | Non-allergen | Non-allergen | Nontoxin |
|  |  | EQTQGNFGD | Allergen | Allergen | Non-allergen | Non-allergen | Nontoxin |
|  |  | IGMEVTPSG | Allergen | Non-allergen | Non-allergen | Non-allergen | Nontoxin |
| HTL epitopes | Surface glycoprotein | DSFVIRGDEVRQIAP | Non-Allergen | Non-Allergen | Non-allergen | Non-allergen | Nontoxin |
|  |  | SFVIRGDEVRQIAPG | Allergen | Allergen | Non-allergen | Non-allergen | Nontoxin |
|  | Membrane glycoprotein | LACFVLAAVYRINWI | Non-Allergen | Non-Allergen | Non-Allergen | Non-Allergen | Nontoxin |
|  |  | FVLAAVYRINWITGG | Non-Allergen | Allergen | Non-Allergen | Non-Allergen | Nontoxin |
|  |  | AIAMACLVGLMWLSY | Allergen | Non-Allergen | Non-Allergen | Non-Allergen | Nontoxin |
|  |  | IAIAMACLVGLMWLS | Non-Allergen | Non-Allergen | Non-Allergen | Non-Allergen | Nontoxin |
|  |  | ITGGIAIAMACLVGL | Allergen | Non-Allergen | Non-Allergen | Non-Allergen | Nontoxin |
|  |  | GIAIAMACLVGLMWL | Non-Allergen | Non-Allergen | Non-Allergen | Non-Allergen | Nontoxin |
|  |  | GGIAIAMACLVGLMW | Non-Allergen | Non-Allergen | Non-Allergen | Non-Allergen | Nontoxin |
|  |  | WITGGIAIAMACLVG | Non-Allergen | Non-Allergen | Non-Allergen | Non-Allergen | Nontoxin |
|  |  | IAMACLVGLMWLSYF | Non-Allergen | Non-Allergen | Non-Allergen | Non-Allergen | Nontoxin |
|  |  | VGLMWLSYFIASFRL | Non-Allergen | Allergen | Non-Allergen | Non-Allergen | Nontoxin |
|  |  | LVIGAVILRGHLRIA | Non-Allergen | Non-Allergen | Non-Allergen | Non-Allergen | Nontoxin |
|  |  | ELVIGAVILRGHLRI | Non-Allergen | Allergen | Non-Allergen | Non-Allergen | Nontoxin |
|  |  | PLLESELVIGAVILR | Non-Allergen | Non-Allergen | Non-Allergen | Non-Allergen | Nontoxin |
|  |  | SELVIGAVILRGHLR | Non-Allergen | Non-Allergen | Non-Allergen | Non-Allergen | Nontoxin |
|  |  | LESELVIGAVILRGH | Non-Allergen | Allergen | Non-Allergen | Non-Allergen | Nontoxin |
|  |  | ESELVIGAVILRGHL | Allergen | Allergen | Non-Allergen | Non-Allergen | Nontoxin |
|  |  | IAGHHLGRCDIKDLP | Non-Allergen | Allergen | Non-Allergen | Non-Allergen | Nontoxin |
|  |  | LLESELVIGAVILRG | Non-Allergen | Non-Allergen | Non-Allergen | Non-Allergen | Nontoxin |
|  |  | VIGAVILRGHLRIAG | Non-Allergen | Non-Allergen | Non-Allergen | Non-Allergen | Nontoxin |
|  |  | IGAVILRGHLRIAGH | Non-Allergen | Allergen | Non-Allergen | Non-Allergen | Nontoxin |
|  | Envelope peotein | VTLAILTALRLCAYC | Non allergen | Non allergen | Non allergen | Non allergen | Nontoxin |
|  |  | LAFVVFLLVTLAILT | Non allergen | Non allergen | Non allergen | Non allergen | Nontoxin |
|  |  | LLFLAFVVFLLVTLA | Non allergen | Non allergen | Non allergen | Non allergen | Nontoxin |
|  |  | VSLVKPSFYVYSRVK | allergen | Non allergen | Non allergen | Non allergen | Nontoxin |
|  |  | VVFLLVTLAILTALR | Non allergen | Non allergen | Non allergen | Non allergen | Nontoxin |
|  |  | VNVSLVKPSFYVYSR | Non allergen | allergen | Non allergen | Non allergen | Nontoxin |
|  |  | FLAFVVFLLVTLAIL | Non allergen | Non allergen | Non allergen | Non allergen | Nontoxin |
|  |  | LFLAFVVFLLVTLAI | Non allergen | Non allergen | Non allergen | Non allergen | Nontoxin |
|  |  | ILTALRLCAYCCNIV | allergen | Non allergen | Non allergen | Non allergen | Nontoxin |
|  |  | LVKPSFYVYSRVKNL | Non allergen | Non allergen | Non allergen | Non allergen | Nontoxin |
|  |  | TLAILTALRLCAYCC | Non allergen | Non allergen | Non allergen | Non allergen | Nontoxin |
|  |  | VFLLVTLAILTALRL | allergen | Non allergen | Non allergen | Non allergen | Nontoxin |
|  |  | AILTALRLCAYCCNI | Non allergen | Non allergen | Non allergen | Non allergen | Nontoxin |
|  |  | LAILTALRLCAYCCN | Non allergen | Non allergen | Non allergen | Non allergen | Nontoxin |
|  |  | SLVKPSFYVYSRVKN | allergen | Non allergen | Non allergen | Non allergen | Nontoxin |
|  |  | NVSLVKPSFYVYSRV | Non allergen | Non allergen | Non allergen | Non allergen | Nontoxin |
|  |  | VLLFLAFVVFLLVTL | Non allergen | Non allergen | Non allergen | Non allergen | Nontoxin |
|  |  | IVNVSLVKPSFYVYS | Non allergen | allergen | Non allergen | Non allergen | Nontoxin |
|  |  | FLLVTLAILTALRLC | Non allergen | Non allergen | Non allergen | Non allergen | Nontoxin |
|  |  | FLLVTLAILTALRLC | Non allergen | Non allergen | Non allergen | Non allergen | Nontoxin |
|  | Nucleocapsid protein | DLSPRWYFYYLGTGP | Non-allergen | Non-allergen | Non-allergen | Non-allergen | Nontoxin |
|  |  | LSPRWYFYYLGTGPE | Allergen | Non-allergen | Non-allergen | Non-allergen | Nontoxin |
|  |  | KDLSPRWYFYYLGTG | Non-allergen | Non-allergen | Non-allergen | Non-allergen | Nontoxin |
|  |  | GGDGKMKDLSPRWYF | Non-allergen | Non-allergen | Non-allergen | Non-allergen | Nontoxin |
|  |  | GDGKMKDLSPRWYFY | Non-allergen | Non-allergen | Non-allergen | Non-allergen | Nontoxin |
|  |  | SPRWYFYYLGTGPEA | Non-allergen | Non-allergen | Non-allergen | Non-allergen | Nontoxin |
|  |  | PRWYFYYLGTGPEAG | Non-allergen | Non-allergen | Non-allergen | Non-allergen | Nontoxin |
|  |  | AEGSRGGSQASSRSS | Non-allergen | Non-allergen | Non-allergen | Non-allergen | Nontoxin |
|  |  | RWYFYYLGTGPEAGL | Allergen | Non-allergen | Non-allergen | Non-allergen | Nontoxin |
|  |  | PRGQGVPINTNSSPD | Non-allergen | Non-allergen | Non-allergen | Non-allergen | Nontoxin |
|  |  | LDDKDPNFKDQVILL | Non-allergen | Allergen | Non-allergen | Non-allergen | Nontoxin |
|  |  | WLTYTGAIKLDDKDP | Allergen | Non-allergen | Non-allergen | Non-allergen | Nontoxin |
|  |  | DDKDPNFKDQVILLN | Non-allergen | Non-allergen | Non-allergen | Non-allergen | Nontoxin |
|  |  | AFFGMSRIGMEVTPS | Non-allergen | Allergen | Non-allergen | Non-allergen | Nontoxin |
|  |  | DPNFKDQVILLNKHI | Non-allergen | Allergen | Non-allergen | Non-allergen | Nontoxin |
|  |  | SAFFGMSRIGMEVTP | Non-allergen | Allergen | Non-allergen | Non-allergen | Nontoxin |
|  |  | FGMSRIGMEVTPSGT | Allergen | Non-allergen | Non-allergen | Non-allergen | Nontoxin |
|  |  | RGPEQTQGNFGDQEL | Non-allergen | Allergen | Non-allergen | Non-allergen | Nontoxin |
|  |  | IGMEVTPSGTWLTYT | Non-allergen | Allergen | Non-allergen | Non-allergen | Nontoxin |
|  |  | LNQLESKMSGKGQQQ | Non-allergen | Non-allergen | Non-allergen | Non-allergen | Nontoxin |
