## Supplementary Table 6 for "Genome based Evolutionary study of SARS-CoV-2 towards the Prediction of Epitope Based Chimeric Vaccine"

**Supplementary Table 6**: Proposed CTL and HTL epitopes for vaccine construction.

| ***Protein*** | ***Conserved Sequence*** | ***Allergenicity*** | ***Toxicity*** | ***Vaxijen Score*** | ***Topology*** |
| --- | --- | --- | --- | --- | --- |
| Surface glycoprotein | ADYNYKLPD | Non-allergen | Non Toxin | 1.3382 | Outside |
|  | SFVIRGDEVRQIAPG | Non-allergen | Non Toxin | 0.5882 | Outside |
| Membrane glycoprotein | LVIGAVILR | Non-allergen | Non Toxin | 1.1027 | Outside |
|  | LACFVLAAVYRINWI | Non-allergen | Non Toxin | 1.2905 | Outside |
| Envelope protein | FVVFLLVTL | Non-allergen | Non Toxin | 0.7403 | Outside |
|  | VTLAILTALRLCAYC | Non-allergen | Non Toxin | 0.8599 | Outside |
| Nucleocapsid protein | AIKLDDKDP | Non-allergen | Non Toxin | 2.1670 | Outside |
|  | DLSPRWYFYYLGTGP | Non-allergen | Non Toxin | 1.5180 | Outside |
