## Supplementary Table 7 for "Genome based Evolutionary study of SARS-CoV-2 towards the Prediction of Epitope Based Chimeric Vaccine"

**Supplementary Table 7:** Predicted conformational epitopes within construct V3.

| **No.** | **Residues** | **No. of residues** | **Score** |
| --- | --- | --- | --- |
| 1 | A:D77, A:K78, A:F79, A:T80, A:T81, A:E82, A:E83, A:L84, A:R85, A:A87, A:A88, A:G90, A:Y91, A:L92, A:E93, A:A94, A:A95, A:T96, A:N97, A:R98, A:Y99, A:N100, A:E101, A:E104, A:R105 | 25 | 0.816 |
| 2 | A:A43, A:E44, A:E45 | 3 | 0.803 |
| 3 | A:D131, A:Q132, A:A133, A:V134, A:E135, A:L136, A:T137, A:Q138, A:E139, A:A140, A:G142, A:T143, A:S146, A:R149 | 14 | 0.775 |
| 4 | A:A48, A:E49, A:R51 | 3 | 0.617 |
| 5 | A:I73, A:E74, A:R76 | 3 | 0.592 |
| 6 | A:T52, A:R53, A:V54, A:E55, A:R58 | 5 | 0.575 |
