## Supplementary Table 8 for "Genome based Evolutionary study of SARS-CoV-2 towards the Prediction of Epitope Based Chimeric Vaccine"

**Supplementary Table 8:** Predicted IFN alpha producing epitopes in the vaccine construct V3.

| **Start-End** | **Sequence** | **Method** | **Result** | **Score** |
| --- | --- | --- | --- | --- |
| 194-209 | [GGGSLVIGAVILRGG](http://crdd.osdd.net/raghava/ifnepitope/pep_design.php?sequence=GGGSLVIGAVILRGG&method=hybrid&model=main) | MERCI | NEGATIVE | 17 |
| 195-210 | [GGSLVIGAVILRGGG](http://crdd.osdd.net/raghava/ifnepitope/pep_design.php?sequence=GGSLVIGAVILRGGG&method=hybrid&model=main) | MERCI | NEGATIVE | 17 |
| 197-212 | [SLVIGAVILRGGGSF](http://crdd.osdd.net/raghava/ifnepitope/pep_design.php?sequence=SLVIGAVILRGGGSF&method=hybrid&model=main) | MERCI | NEGATIVE | 17 |
| 196-211 | [GSLVIGAVILRGGGS](http://crdd.osdd.net/raghava/ifnepitope/pep_design.php?sequence=GSLVIGAVILRGGGS&method=hybrid&model=main) | MERCI | NEGATIVE | 16 |
| 198-213 | [LVIGAVILRGGGSFV](http://crdd.osdd.net/raghava/ifnepitope/pep_design.php?sequence=LVIGAVILRGGGSFV&method=hybrid&model=main) | MERCI | NEGATIVE | 15 |
| 193-208 | [DGGGSLVIGAVILRG](http://crdd.osdd.net/raghava/ifnepitope/pep_design.php?sequence=DGGGSLVIGAVILRG&method=hybrid&model=main) | MERCI | NEGATIVE | 13 |
| 191-206 | [LPDGGGSLVIGAVIL](http://crdd.osdd.net/raghava/ifnepitope/pep_design.php?sequence=LPDGGGSLVIGAVIL&method=hybrid&model=main) | MERCI | NEGATIVE | 9 |
| 192-207 | [PDGGGSLVIGAVILR](http://crdd.osdd.net/raghava/ifnepitope/pep_design.php?sequence=PDGGGSLVIGAVILR&method=hybrid&model=main) | MERCI | NEGATIVE | 9 |
| 272-287 | [IGPGPGVTLAILTAL](http://crdd.osdd.net/raghava/ifnepitope/pep_design.php?sequence=IGPGPGVTLAILTAL&method=hybrid&model=main) | MERCI | NEGATIVE | 9 |
| 273-288 | [GPGPGVTLAILTALR](http://crdd.osdd.net/raghava/ifnepitope/pep_design.php?sequence=GPGPGVTLAILTALR&method=hybrid&model=main) | MERCI | NEGATIVE | 8 |
| 274-289 | [PGPGVTLAILTALRL](http://crdd.osdd.net/raghava/ifnepitope/pep_design.php?sequence=PGPGVTLAILTALRL&method=hybrid&model=main) | MERCI | NEGATIVE | 8 |
| 275-290 | [GPGVTLAILTALRLC](http://crdd.osdd.net/raghava/ifnepitope/pep_design.php?sequence=GPGVTLAILTALRLC&method=hybrid&model=main) | MERCI | NEGATIVE | 8 |
| 199-214 | [VIGAVILRGGGSFVV](http://crdd.osdd.net/raghava/ifnepitope/pep_design.php?sequence=VIGAVILRGGGSFVV&method=hybrid&model=main) | MERCI | NEGATIVE | 7 |
| 276-291 | [PGVTLAILTALRLCA](http://crdd.osdd.net/raghava/ifnepitope/pep_design.php?sequence=PGVTLAILTALRLCA&method=hybrid&model=main) | MERCI | NEGATIVE | 7 |
| 269-284 | [INWIGPGPGVTLAIL](http://crdd.osdd.net/raghava/ifnepitope/pep_design.php?sequence=INWIGPGPGVTLAIL&method=hybrid&model=main) | MERCI | NEGATIVE | 6 |
| 271-286 | [WIGPGPGVTLAILTA](http://crdd.osdd.net/raghava/ifnepitope/pep_design.php?sequence=WIGPGPGVTLAILTA&method=hybrid&model=main) | MERCI | NEGATIVE | 6 |
| 277-292 | [GVTLAILTALRLCAY](http://crdd.osdd.net/raghava/ifnepitope/pep_design.php?sequence=GVTLAILTALRLCAY&method=hybrid&model=main) | MERCI | NEGATIVE | 6 |
| 176-191 | [LKAAAGGGSADYNYK](http://crdd.osdd.net/raghava/ifnepitope/pep_design.php?sequence=LKAAAGGGSADYNYK&method=hybrid&model=main) | MERCI | NEGATIVE | 5 |
| 177-192 | [KAAAGGGSADYNYKL](http://crdd.osdd.net/raghava/ifnepitope/pep_design.php?sequence=KAAAGGGSADYNYKL&method=hybrid&model=main) | MERCI | NEGATIVE | 5 |
| 270-285 | [NWIGPGPGVTLAILT](http://crdd.osdd.net/raghava/ifnepitope/pep_design.php?sequence=NWIGPGPGVTLAILT&method=hybrid&model=main) | MERCI | NEGATIVE | 5 |
| 173-188 | [AWTLKAAAGGGSADY](http://crdd.osdd.net/raghava/ifnepitope/pep_design.php?sequence=AWTLKAAAGGGSADY&method=hybrid&model=main) | MERCI | NEGATIVE | 4 |
| 174-189 | [WTLKAAAGGGSADYN](http://crdd.osdd.net/raghava/ifnepitope/pep_design.php?sequence=WTLKAAAGGGSADYN&method=hybrid&model=main) | MERCI | NEGATIVE | 4 |
| 175-190 | [TLKAAAGGGSADYNY](http://crdd.osdd.net/raghava/ifnepitope/pep_design.php?sequence=TLKAAAGGGSADYNY&method=hybrid&model=main) | MERCI | NEGATIVE | 4 |
| 190-205 | [KLPDGGGSLVIGAVI](http://crdd.osdd.net/raghava/ifnepitope/pep_design.php?sequence=KLPDGGGSLVIGAVI&method=hybrid&model=main) | MERCI | NEGATIVE | 4 |
| 172-187 | [AAWTLKAAAGGGSAD](http://crdd.osdd.net/raghava/ifnepitope/pep_design.php?sequence=AAWTLKAAAGGGSAD&method=hybrid&model=main) | MERCI | NEGATIVE | 3 |
| 200-215 | [IGAVILRGGGSFVVF](http://crdd.osdd.net/raghava/ifnepitope/pep_design.php?sequence=IGAVILRGGGSFVVF&method=hybrid&model=main) | MERCI | NEGATIVE | 3 |
| 154-169 | [AAKLVGIELEAAAKA](http://crdd.osdd.net/raghava/ifnepitope/pep_design.php?sequence=AAKLVGIELEAAAKA&method=hybrid&model=main) | MERCI | POSITIVE | 2 |
| 155-170 | [AKLVGIELEAAAKAK](http://crdd.osdd.net/raghava/ifnepitope/pep_design.php?sequence=AKLVGIELEAAAKAK&method=hybrid&model=main) | MERCI | POSITIVE | 2 |
| 156-171 | [KLVGIELEAAAKAKF](http://crdd.osdd.net/raghava/ifnepitope/pep_design.php?sequence=KLVGIELEAAAKAKF&method=hybrid&model=main) | MERCI | POSITIVE | 2 |
| 157-172 | [LVGIELEAAAKAKFV](http://crdd.osdd.net/raghava/ifnepitope/pep_design.php?sequence=LVGIELEAAAKAKFV&method=hybrid&model=main) | MERCI | POSITIVE | 2 |
| 158-173 | [VGIELEAAAKAKFVA](http://crdd.osdd.net/raghava/ifnepitope/pep_design.php?sequence=VGIELEAAAKAKFVA&method=hybrid&model=main) | MERCI | POSITIVE | 2 |
| 159-174 | [GIELEAAAKAKFVAA](http://crdd.osdd.net/raghava/ifnepitope/pep_design.php?sequence=GIELEAAAKAKFVAA&method=hybrid&model=main) | MERCI | POSITIVE | 2 |
| 160-175 | [IELEAAAKAKFVAAW](http://crdd.osdd.net/raghava/ifnepitope/pep_design.php?sequence=IELEAAAKAKFVAAW&method=hybrid&model=main) | MERCI | POSITIVE | 2 |
| 161-176 | [ELEAAAKAKFVAAWT](http://crdd.osdd.net/raghava/ifnepitope/pep_design.php?sequence=ELEAAAKAKFVAAWT&method=hybrid&model=main) | MERCI | POSITIVE | 2 |
| 162-177 | [LEAAAKAKFVAAWTL](http://crdd.osdd.net/raghava/ifnepitope/pep_design.php?sequence=LEAAAKAKFVAAWTL&method=hybrid&model=main) | MERCI | POSITIVE | 2 |
| 230-245 | [KDPGPGPGSFVIRGD](http://crdd.osdd.net/raghava/ifnepitope/pep_design.php?sequence=KDPGPGPGSFVIRGD&method=hybrid&model=main) | MERCI | POSITIVE | 2 |
| 231-246 | [DPGPGPGSFVIRGDE](http://crdd.osdd.net/raghava/ifnepitope/pep_design.php?sequence=DPGPGPGSFVIRGDE&method=hybrid&model=main) | MERCI | POSITIVE | 2 |
| 232-247 | [PGPGPGSFVIRGDEV](http://crdd.osdd.net/raghava/ifnepitope/pep_design.php?sequence=PGPGPGSFVIRGDEV&method=hybrid&model=main) | MERCI | POSITIVE | 2 |
| 233-248 | [GPGPGSFVIRGDEVR](http://crdd.osdd.net/raghava/ifnepitope/pep_design.php?sequence=GPGPGSFVIRGDEVR&method=hybrid&model=main) | MERCI | POSITIVE | 2 |
| 234-249 | [PGPGSFVIRGDEVRQ](http://crdd.osdd.net/raghava/ifnepitope/pep_design.php?sequence=PGPGSFVIRGDEVRQ&method=hybrid&model=main) | MERCI | POSITIVE | 2 |
| 235-250 | [GPGSFVIRGDEVRQI](http://crdd.osdd.net/raghava/ifnepitope/pep_design.php?sequence=GPGSFVIRGDEVRQI&method=hybrid&model=main) | MERCI | POSITIVE | 2 |
| 236-251 | [PGSFVIRGDEVRQIA](http://crdd.osdd.net/raghava/ifnepitope/pep_design.php?sequence=PGSFVIRGDEVRQIA&method=hybrid&model=main) | MERCI | POSITIVE | 2 |
| 237-252 | [GSFVIRGDEVRQIAP](http://crdd.osdd.net/raghava/ifnepitope/pep_design.php?sequence=GSFVIRGDEVRQIAP&method=hybrid&model=main) | MERCI | POSITIVE | 2 |
| 238-253 | [SFVIRGDEVRQIAPG](http://crdd.osdd.net/raghava/ifnepitope/pep_design.php?sequence=SFVIRGDEVRQIAPG&method=hybrid&model=main) | MERCI | POSITIVE | 2 |
| 239-254 | [FVIRGDEVRQIAPGG](http://crdd.osdd.net/raghava/ifnepitope/pep_design.php?sequence=FVIRGDEVRQIAPGG&method=hybrid&model=main) | MERCI | POSITIVE | 2 |
| 206-221 | [RGGGSFVVFLLVTLG](http://crdd.osdd.net/raghava/ifnepitope/pep_design.php?sequence=RGGGSFVVFLLVTLG&method=hybrid&model=main) | MERCI | NEGATIVE | 2 |
| 278-293 | [VTLAILTALRLCAYC](http://crdd.osdd.net/raghava/ifnepitope/pep_design.php?sequence=VTLAILTALRLCAYC&method=hybrid&model=main) | MERCI | NEGATIVE | 2 |
| 163-178 | [EAAAKAKFVAAWTLK](http://crdd.osdd.net/raghava/ifnepitope/pep_design.php?sequence=EAAAKAKFVAAWTLK&method=hybrid&model=main) | SVM | POSITIVE | 1.12401 |
| 164-179 | [AAAKAKFVAAWTLKA](http://crdd.osdd.net/raghava/ifnepitope/pep_design.php?sequence=AAAKAKFVAAWTLKA&method=hybrid&model=main) | SVM | POSITIVE | 1.11149 |
| 165-180 | [AAKAKFVAAWTLKAA](http://crdd.osdd.net/raghava/ifnepitope/pep_design.php?sequence=AAKAKFVAAWTLKAA&method=hybrid&model=main) | SVM | POSITIVE | 1.11149 |
| 166-181 | [AKAKFVAAWTLKAAA](http://crdd.osdd.net/raghava/ifnepitope/pep_design.php?sequence=AKAKFVAAWTLKAAA&method=hybrid&model=main) | SVM | POSITIVE | 1.11149 |
| 68-83 | [PEQFIELRDKFTTEE](http://crdd.osdd.net/raghava/ifnepitope/pep_design.php?sequence=PEQFIELRDKFTTEE&method=hybrid&model=main) | MERCI | POSITIVE | 1 |
| 69-84 | [EQFIELRDKFTTEEL](http://crdd.osdd.net/raghava/ifnepitope/pep_design.php?sequence=EQFIELRDKFTTEEL&method=hybrid&model=main) | MERCI | POSITIVE | 1 |
| 70-85 | [QFIELRDKFTTEELR](http://crdd.osdd.net/raghava/ifnepitope/pep_design.php?sequence=QFIELRDKFTTEELR&method=hybrid&model=main) | MERCI | POSITIVE | 1 |
| 71-86 | [FIELRDKFTTEELRK](http://crdd.osdd.net/raghava/ifnepitope/pep_design.php?sequence=FIELRDKFTTEELRK&method=hybrid&model=main) | MERCI | POSITIVE | 1 |
| 72-87 | [IELRDKFTTEELRKA](http://crdd.osdd.net/raghava/ifnepitope/pep_design.php?sequence=IELRDKFTTEELRKA&method=hybrid&model=main) | MERCI | POSITIVE | 1 |
| 73-88 | [ELRDKFTTEELRKAA](http://crdd.osdd.net/raghava/ifnepitope/pep_design.php?sequence=ELRDKFTTEELRKAA&method=hybrid&model=main) | MERCI | POSITIVE | 1 |
| 106-121 | [EAALQRLRSQTAFED](http://crdd.osdd.net/raghava/ifnepitope/pep_design.php?sequence=EAALQRLRSQTAFED&method=hybrid&model=main) | MERCI | POSITIVE | 1 |
| 107-122 | [AALQRLRSQTAFEDA](http://crdd.osdd.net/raghava/ifnepitope/pep_design.php?sequence=AALQRLRSQTAFEDA&method=hybrid&model=main) | MERCI | POSITIVE | 1 |
| 108-123 | [ALQRLRSQTAFEDAS](http://crdd.osdd.net/raghava/ifnepitope/pep_design.php?sequence=ALQRLRSQTAFEDAS&method=hybrid&model=main) | MERCI | POSITIVE | 1 |
| 109-124 | [LQRLRSQTAFEDASA](http://crdd.osdd.net/raghava/ifnepitope/pep_design.php?sequence=LQRLRSQTAFEDASA&method=hybrid&model=main) | MERCI | POSITIVE | 1 |
| 110-125 | [QRLRSQTAFEDASAR](http://crdd.osdd.net/raghava/ifnepitope/pep_design.php?sequence=QRLRSQTAFEDASAR&method=hybrid&model=main) | MERCI | POSITIVE | 1 |
| 111-126 | [RLRSQTAFEDASARA](http://crdd.osdd.net/raghava/ifnepitope/pep_design.php?sequence=RLRSQTAFEDASARA&method=hybrid&model=main) | MERCI | POSITIVE | 1 |
| 112-127 | [LRSQTAFEDASARAE](http://crdd.osdd.net/raghava/ifnepitope/pep_design.php?sequence=LRSQTAFEDASARAE&method=hybrid&model=main) | MERCI | POSITIVE | 1 |
| 385-400 | [AMACLVGLMKKYVYS](http://crdd.osdd.net/raghava/ifnepitope/pep_design.php?sequence=AMACLVGLMKKYVYS&method=hybrid&model=main) | MERCI | POSITIVE | 1 |
| 386-401 | [MACLVGLMKKYVYSR](http://crdd.osdd.net/raghava/ifnepitope/pep_design.php?sequence=MACLVGLMKKYVYSR&method=hybrid&model=main) | MERCI | POSITIVE | 1 |
| 387-402 | [ACLVGLMKKYVYSRV](http://crdd.osdd.net/raghava/ifnepitope/pep_design.php?sequence=ACLVGLMKKYVYSRV&method=hybrid&model=main) | MERCI | POSITIVE | 1 |
| 388-403 | [CLVGLMKKYVYSRVK](http://crdd.osdd.net/raghava/ifnepitope/pep_design.php?sequence=CLVGLMKKYVYSRVK&method=hybrid&model=main) | MERCI | POSITIVE | 1 |
| 429-444 | [VPKKLCAYCCNIVKK](http://crdd.osdd.net/raghava/ifnepitope/pep_design.php?sequence=VPKKLCAYCCNIVKK&method=hybrid&model=main) | MERCI | POSITIVE | 1 |
| 430-445 | [PKKLCAYCCNIVKKP](http://crdd.osdd.net/raghava/ifnepitope/pep_design.php?sequence=PKKLCAYCCNIVKKP&method=hybrid&model=main) | MERCI | POSITIVE | 1 |
| 431-446 | [KKLCAYCCNIVKKPG](http://crdd.osdd.net/raghava/ifnepitope/pep_design.php?sequence=KKLCAYCCNIVKKPG&method=hybrid&model=main) | MERCI | POSITIVE | 1 |
| 432-447 | [KLCAYCCNIVKKPGS](http://crdd.osdd.net/raghava/ifnepitope/pep_design.php?sequence=KLCAYCCNIVKKPGS&method=hybrid&model=main) | MERCI | POSITIVE | 1 |
| 433-448 | [LCAYCCNIVKKPGSS](http://crdd.osdd.net/raghava/ifnepitope/pep_design.php?sequence=LCAYCCNIVKKPGSS&method=hybrid&model=main) | MERCI | POSITIVE | 1 |
| 434-449 | [CAYCCNIVKKPGSSR](http://crdd.osdd.net/raghava/ifnepitope/pep_design.php?sequence=CAYCCNIVKKPGSSR&method=hybrid&model=main) | MERCI | POSITIVE | 1 |
| 435-450 | [AYCCNIVKKPGSSRG](http://crdd.osdd.net/raghava/ifnepitope/pep_design.php?sequence=AYCCNIVKKPGSSRG&method=hybrid&model=main) | MERCI | POSITIVE | 1 |
| 436-451 | [YCCNIVKKPGSSRGT](http://crdd.osdd.net/raghava/ifnepitope/pep_design.php?sequence=YCCNIVKKPGSSRGT&method=hybrid&model=main) | MERCI | POSITIVE | 1 |
| 474-489 | [KKHWPQIAQFAPSAS](http://crdd.osdd.net/raghava/ifnepitope/pep_design.php?sequence=KKHWPQIAQFAPSAS&method=hybrid&model=main) | MERCI | POSITIVE | 1 |
| 475-490 | [KHWPQIAQFAPSASA](http://crdd.osdd.net/raghava/ifnepitope/pep_design.php?sequence=KHWPQIAQFAPSASA&method=hybrid&model=main) | MERCI | POSITIVE | 1 |
| 476-491 | [HWPQIAQFAPSASAF](http://crdd.osdd.net/raghava/ifnepitope/pep_design.php?sequence=HWPQIAQFAPSASAF&method=hybrid&model=main) | MERCI | POSITIVE | 1 |
| 477-492 | [WPQIAQFAPSASAFK](http://crdd.osdd.net/raghava/ifnepitope/pep_design.php?sequence=WPQIAQFAPSASAFK&method=hybrid&model=main) | MERCI | POSITIVE | 1 |
| 478-493 | [PQIAQFAPSASAFKK](http://crdd.osdd.net/raghava/ifnepitope/pep_design.php?sequence=PQIAQFAPSASAFKK&method=hybrid&model=main) | MERCI | POSITIVE | 1 |
| 479-493 | [QIAQFAPSASAFKK](http://crdd.osdd.net/raghava/ifnepitope/pep_design.php?sequence=QIAQFAPSASAFKK&method=hybrid&model=main) | MERCI | POSITIVE | 1 |
| 19-34 | [AALGAADLALATVND](http://crdd.osdd.net/raghava/ifnepitope/pep_design.php?sequence=AALGAADLALATVND&method=hybrid&model=main) | MERCI | NEGATIVE | 1 |
| 20-35 | [ALGAADLALATVNDL](http://crdd.osdd.net/raghava/ifnepitope/pep_design.php?sequence=ALGAADLALATVNDL&method=hybrid&model=main) | MERCI | NEGATIVE | 1 |
| 21-36 | [LGAADLALATVNDLI](http://crdd.osdd.net/raghava/ifnepitope/pep_design.php?sequence=LGAADLALATVNDLI&method=hybrid&model=main) | MERCI | NEGATIVE | 1 |
| 22-37 | [GAADLALATVNDLIA](http://crdd.osdd.net/raghava/ifnepitope/pep_design.php?sequence=GAADLALATVNDLIA&method=hybrid&model=main) | MERCI | NEGATIVE | 1 |
| 23-38 | [AADLALATVNDLIAN](http://crdd.osdd.net/raghava/ifnepitope/pep_design.php?sequence=AADLALATVNDLIAN&method=hybrid&model=main) | MERCI | NEGATIVE | 1 |
| 24-39 | [ADLALATVNDLIANL](http://crdd.osdd.net/raghava/ifnepitope/pep_design.php?sequence=ADLALATVNDLIANL&method=hybrid&model=main) | MERCI | NEGATIVE | 1 |
| 25-40 | [DLALATVNDLIANLR](http://crdd.osdd.net/raghava/ifnepitope/pep_design.php?sequence=DLALATVNDLIANLR&method=hybrid&model=main) | MERCI | NEGATIVE | 1 |
| 143-158 | [VASQTRAVGERAAKL](http://crdd.osdd.net/raghava/ifnepitope/pep_design.php?sequence=VASQTRAVGERAAKL&method=hybrid&model=main) | MERCI | NEGATIVE | 1 |
| 144-159 | [ASQTRAVGERAAKLV](http://crdd.osdd.net/raghava/ifnepitope/pep_design.php?sequence=ASQTRAVGERAAKLV&method=hybrid&model=main) | MERCI | NEGATIVE | 1 |
| 145-160 | [SQTRAVGERAAKLVG](http://crdd.osdd.net/raghava/ifnepitope/pep_design.php?sequence=SQTRAVGERAAKLVG&method=hybrid&model=main) | MERCI | NEGATIVE | 1 |
| 146-161 | [QTRAVGERAAKLVGI](http://crdd.osdd.net/raghava/ifnepitope/pep_design.php?sequence=QTRAVGERAAKLVGI&method=hybrid&model=main) | MERCI | NEGATIVE | 1 |
| 147-162 | [TRAVGERAAKLVGIE](http://crdd.osdd.net/raghava/ifnepitope/pep_design.php?sequence=TRAVGERAAKLVGIE&method=hybrid&model=main) | MERCI | NEGATIVE | 1 |
| 171-186 | [VAAWTLKAAAGGGSA](http://crdd.osdd.net/raghava/ifnepitope/pep_design.php?sequence=VAAWTLKAAAGGGSA&method=hybrid&model=main) | MERCI | NEGATIVE | 1 |
| 178-193 | [AAAGGGSADYNYKLP](http://crdd.osdd.net/raghava/ifnepitope/pep_design.php?sequence=AAAGGGSADYNYKLP&method=hybrid&model=main) | MERCI | NEGATIVE | 1 |
| 179-194 | [AAGGGSADYNYKLPD](http://crdd.osdd.net/raghava/ifnepitope/pep_design.php?sequence=AAGGGSADYNYKLPD&method=hybrid&model=main) | MERCI | NEGATIVE | 1 |
| 180-195 | [AGGGSADYNYKLPDG](http://crdd.osdd.net/raghava/ifnepitope/pep_design.php?sequence=AGGGSADYNYKLPDG&method=hybrid&model=main) | MERCI | NEGATIVE | 1 |
| 181-196 | [GGGSADYNYKLPDGG](http://crdd.osdd.net/raghava/ifnepitope/pep_design.php?sequence=GGGSADYNYKLPDGG&method=hybrid&model=main) | MERCI | NEGATIVE | 1 |
| 182-197 | [GGSADYNYKLPDGGG](http://crdd.osdd.net/raghava/ifnepitope/pep_design.php?sequence=GGSADYNYKLPDGGG&method=hybrid&model=main) | MERCI | NEGATIVE | 1 |
| 183-198 | [GSADYNYKLPDGGGS](http://crdd.osdd.net/raghava/ifnepitope/pep_design.php?sequence=GSADYNYKLPDGGGS&method=hybrid&model=main) | MERCI | NEGATIVE | 1 |
| 189-204 | [YKLPDGGGSLVIGAV](http://crdd.osdd.net/raghava/ifnepitope/pep_design.php?sequence=YKLPDGGGSLVIGAV&method=hybrid&model=main) | MERCI | NEGATIVE | 1 |
| 205-220 | [LRGGGSFVVFLLVTL](http://crdd.osdd.net/raghava/ifnepitope/pep_design.php?sequence=LRGGGSFVVFLLVTL&method=hybrid&model=main) | MERCI | NEGATIVE | 1 |
| 213-228 | [VFLLVTLGGGSAIKL](http://crdd.osdd.net/raghava/ifnepitope/pep_design.php?sequence=VFLLVTLGGGSAIKL&method=hybrid&model=main) | MERCI | NEGATIVE | 1 |
| 214-229 | [FLLVTLGGGSAIKLD](http://crdd.osdd.net/raghava/ifnepitope/pep_design.php?sequence=FLLVTLGGGSAIKLD&method=hybrid&model=main) | MERCI | NEGATIVE | 1 |
| 215-230 | [LLVTLGGGSAIKLDD](http://crdd.osdd.net/raghava/ifnepitope/pep_design.php?sequence=LLVTLGGGSAIKLDD&method=hybrid&model=main) | MERCI | NEGATIVE | 1 |
| 216-231 | [LVTLGGGSAIKLDDK](http://crdd.osdd.net/raghava/ifnepitope/pep_design.php?sequence=LVTLGGGSAIKLDDK&method=hybrid&model=main) | MERCI | NEGATIVE | 1 |
| 217-232 | [VTLGGGSAIKLDDKD](http://crdd.osdd.net/raghava/ifnepitope/pep_design.php?sequence=VTLGGGSAIKLDDKD&method=hybrid&model=main) | MERCI | NEGATIVE | 1 |
| 279-294 | [TLAILTALRLCAYCG](http://crdd.osdd.net/raghava/ifnepitope/pep_design.php?sequence=TLAILTALRLCAYCG&method=hybrid&model=main) | MERCI | NEGATIVE | 1 |
| 320-335 | [VRQIAPGQTGKIADY](http://crdd.osdd.net/raghava/ifnepitope/pep_design.php?sequence=VRQIAPGQTGKIADY&method=hybrid&model=main) | MERCI | NEGATIVE | 1 |
| 321-336 | [RQIAPGQTGKIADYN](http://crdd.osdd.net/raghava/ifnepitope/pep_design.php?sequence=RQIAPGQTGKIADYN&method=hybrid&model=main) | MERCI | NEGATIVE | 1 |
| 322-337 | [QIAPGQTGKIADYNY](http://crdd.osdd.net/raghava/ifnepitope/pep_design.php?sequence=QIAPGQTGKIADYNY&method=hybrid&model=main) | MERCI | NEGATIVE | 1 |
| 323-338 | [IAPGQTGKIADYNYK](http://crdd.osdd.net/raghava/ifnepitope/pep_design.php?sequence=IAPGQTGKIADYNYK&method=hybrid&model=main) | MERCI | NEGATIVE | 1 |
| 324-339 | [APGQTGKIADYNYKK](http://crdd.osdd.net/raghava/ifnepitope/pep_design.php?sequence=APGQTGKIADYNYKK&method=hybrid&model=main) | MERCI | NEGATIVE | 1 |
| 325-340 | [PGQTGKIADYNYKKK](http://crdd.osdd.net/raghava/ifnepitope/pep_design.php?sequence=PGQTGKIADYNYKKK&method=hybrid&model=main) | MERCI | NEGATIVE | 1 |
| 326-341 | [GQTGKIADYNYKKKG](http://crdd.osdd.net/raghava/ifnepitope/pep_design.php?sequence=GQTGKIADYNYKKKG&method=hybrid&model=main) | MERCI | NEGATIVE | 1 |
| 327-342 | [QTGKIADYNYKKKGD](http://crdd.osdd.net/raghava/ifnepitope/pep_design.php?sequence=QTGKIADYNYKKKGD&method=hybrid&model=main) | MERCI | NEGATIVE | 1 |
| 356-371 | [KMWSFNPETNKKLFA](http://crdd.osdd.net/raghava/ifnepitope/pep_design.php?sequence=KMWSFNPETNKKLFA&method=hybrid&model=main) | MERCI | NEGATIVE | 1 |
| 357-372 | [MWSFNPETNKKLFAR](http://crdd.osdd.net/raghava/ifnepitope/pep_design.php?sequence=MWSFNPETNKKLFAR&method=hybrid&model=main) | MERCI | NEGATIVE | 1 |
| 358-373 | [WSFNPETNKKLFART](http://crdd.osdd.net/raghava/ifnepitope/pep_design.php?sequence=WSFNPETNKKLFART&method=hybrid&model=main) | MERCI | NEGATIVE | 1 |
| 359-374 | [SFNPETNKKLFARTR](http://crdd.osdd.net/raghava/ifnepitope/pep_design.php?sequence=SFNPETNKKLFARTR&method=hybrid&model=main) | MERCI | NEGATIVE | 1 |
| 360-375 | [FNPETNKKLFARTRS](http://crdd.osdd.net/raghava/ifnepitope/pep_design.php?sequence=FNPETNKKLFARTRS&method=hybrid&model=main) | MERCI | NEGATIVE | 1 |
| 361-376 | [NPETNKKLFARTRSM](http://crdd.osdd.net/raghava/ifnepitope/pep_design.php?sequence=NPETNKKLFARTRSM&method=hybrid&model=main) | MERCI | NEGATIVE | 1 |
| 362-377 | [PETNKKLFARTRSMW](http://crdd.osdd.net/raghava/ifnepitope/pep_design.php?sequence=PETNKKLFARTRSMW&method=hybrid&model=main) | MERCI | NEGATIVE | 1 |
| 363-378 | [ETNKKLFARTRSMWS](http://crdd.osdd.net/raghava/ifnepitope/pep_design.php?sequence=ETNKKLFARTRSMWS&method=hybrid&model=main) | MERCI | NEGATIVE | 1 |
| 364-379 | [TNKKLFARTRSMWSF](http://crdd.osdd.net/raghava/ifnepitope/pep_design.php?sequence=TNKKLFARTRSMWSF&method=hybrid&model=main) | MERCI | NEGATIVE | 1 |
| 379-394 | [NPETKKAMACLVGLM](http://crdd.osdd.net/raghava/ifnepitope/pep_design.php?sequence=NPETKKAMACLVGLM&method=hybrid&model=main) | MERCI | NEGATIVE | 1 |
| 380-395 | [PETKKAMACLVGLMK](http://crdd.osdd.net/raghava/ifnepitope/pep_design.php?sequence=PETKKAMACLVGLMK&method=hybrid&model=main) | MERCI | NEGATIVE | 1 |
| 381-396 | [ETKKAMACLVGLMKK](http://crdd.osdd.net/raghava/ifnepitope/pep_design.php?sequence=ETKKAMACLVGLMKK&method=hybrid&model=main) | MERCI | NEGATIVE | 1 |
| 382-397 | [TKKAMACLVGLMKKY](http://crdd.osdd.net/raghava/ifnepitope/pep_design.php?sequence=TKKAMACLVGLMKKY&method=hybrid&model=main) | MERCI | NEGATIVE | 1 |
| 383-398 | [KKAMACLVGLMKKYV](http://crdd.osdd.net/raghava/ifnepitope/pep_design.php?sequence=KKAMACLVGLMKKYV&method=hybrid&model=main) | MERCI | NEGATIVE | 1 |
| 390-405 | [VGLMKKYVYSRVKNL](http://crdd.osdd.net/raghava/ifnepitope/pep_design.php?sequence=VGLMKKYVYSRVKNL&method=hybrid&model=main) | MERCI | NEGATIVE | 1 |
| 391-406 | [GLMKKYVYSRVKNLN](http://crdd.osdd.net/raghava/ifnepitope/pep_design.php?sequence=GLMKKYVYSRVKNLN&method=hybrid&model=main) | MERCI | NEGATIVE | 1 |
| 392-407 | [LMKKYVYSRVKNLNS](http://crdd.osdd.net/raghava/ifnepitope/pep_design.php?sequence=LMKKYVYSRVKNLNS&method=hybrid&model=main) | MERCI | NEGATIVE | 1 |
| 393-408 | [MKKYVYSRVKNLNSS](http://crdd.osdd.net/raghava/ifnepitope/pep_design.php?sequence=MKKYVYSRVKNLNSS&method=hybrid&model=main) | MERCI | NEGATIVE | 1 |
| 394-409 | [KKYVYSRVKNLNSSR](http://crdd.osdd.net/raghava/ifnepitope/pep_design.php?sequence=KKYVYSRVKNLNSSR&method=hybrid&model=main) | MERCI | NEGATIVE | 1 |
| 395-410 | [KYVYSRVKNLNSSRV](http://crdd.osdd.net/raghava/ifnepitope/pep_design.php?sequence=KYVYSRVKNLNSSRV&method=hybrid&model=main) | MERCI | NEGATIVE | 1 |
| 419-434 | [SRVKNLNSSRVPKKL](http://crdd.osdd.net/raghava/ifnepitope/pep_design.php?sequence=SRVKNLNSSRVPKKL&method=hybrid&model=main) | MERCI | NEGATIVE | 1 |
| 420-435 | [RVKNLNSSRVPKKLC](http://crdd.osdd.net/raghava/ifnepitope/pep_design.php?sequence=RVKNLNSSRVPKKLC&method=hybrid&model=main) | MERCI | NEGATIVE | 1 |
| 421-436 | [VKNLNSSRVPKKLCA](http://crdd.osdd.net/raghava/ifnepitope/pep_design.php?sequence=VKNLNSSRVPKKLCA&method=hybrid&model=main) | MERCI | NEGATIVE | 1 |
| 422-437 | [KNLNSSRVPKKLCAY](http://crdd.osdd.net/raghava/ifnepitope/pep_design.php?sequence=KNLNSSRVPKKLCAY&method=hybrid&model=main) | MERCI | NEGATIVE | 1 |
| 423-438 | [NLNSSRVPKKLCAYC](http://crdd.osdd.net/raghava/ifnepitope/pep_design.php?sequence=NLNSSRVPKKLCAYC&method=hybrid&model=main) | MERCI | NEGATIVE | 1 |
| 424-439 | [LNSSRVPKKLCAYCC](http://crdd.osdd.net/raghava/ifnepitope/pep_design.php?sequence=LNSSRVPKKLCAYCC&method=hybrid&model=main) | MERCI | NEGATIVE | 1 |
| 425-440 | [NSSRVPKKLCAYCCN](http://crdd.osdd.net/raghava/ifnepitope/pep_design.php?sequence=NSSRVPKKLCAYCCN&method=hybrid&model=main) | MERCI | NEGATIVE | 1 |
| 426-441 | [SSRVPKKLCAYCCNI](http://crdd.osdd.net/raghava/ifnepitope/pep_design.php?sequence=SSRVPKKLCAYCCNI&method=hybrid&model=main) | MERCI | NEGATIVE | 1 |
| 427-442 | [SRVPKKLCAYCCNIV](http://crdd.osdd.net/raghava/ifnepitope/pep_design.php?sequence=SRVPKKLCAYCCNIV&method=hybrid&model=main) | MERCI | NEGATIVE | 1 |
| 428-443 | [RVPKKLCAYCCNIVK](http://crdd.osdd.net/raghava/ifnepitope/pep_design.php?sequence=RVPKKLCAYCCNIVK&method=hybrid&model=main) | MERCI | NEGATIVE | 1 |
| 440-455 | [IVKKPGSSRGTSPAR](http://crdd.osdd.net/raghava/ifnepitope/pep_design.php?sequence=IVKKPGSSRGTSPAR&method=hybrid&model=main) | MERCI | NEGATIVE | 1 |
| 441-456 | [VKKPGSSRGTSPARM](http://crdd.osdd.net/raghava/ifnepitope/pep_design.php?sequence=VKKPGSSRGTSPARM&method=hybrid&model=main) | MERCI | NEGATIVE | 1 |
| 442-457 | [KKPGSSRGTSPARMA](http://crdd.osdd.net/raghava/ifnepitope/pep_design.php?sequence=KKPGSSRGTSPARMA&method=hybrid&model=main) | MERCI | NEGATIVE | 1 |
| 443-458 | [KPGSSRGTSPARMAG](http://crdd.osdd.net/raghava/ifnepitope/pep_design.php?sequence=KPGSSRGTSPARMAG&method=hybrid&model=main) | MERCI | NEGATIVE | 1 |
| 444-459 | [PGSSRGTSPARMAGG](http://crdd.osdd.net/raghava/ifnepitope/pep_design.php?sequence=PGSSRGTSPARMAGG&method=hybrid&model=main) | MERCI | NEGATIVE | 1 |
| 484-493 | [APSASAFKK](http://crdd.osdd.net/raghava/ifnepitope/pep_design.php?sequence=APSASAFKK&method=hybrid&model=main) | SVM | POSITIVE | 0.97618 |
| 207-222 | [GGGSFVVFLLVTLGG](http://crdd.osdd.net/raghava/ifnepitope/pep_design.php?sequence=GGGSFVVFLLVTLGG&method=hybrid&model=main) | SVM | POSITIVE | 0.94972 |
| 208-223 | [GGSFVVFLLVTLGGG](http://crdd.osdd.net/raghava/ifnepitope/pep_design.php?sequence=GGSFVVFLLVTLGGG&method=hybrid&model=main) | SVM | POSITIVE | 0.94972 |
| 203-218 | [VILRGGGSFVVFLLV](http://crdd.osdd.net/raghava/ifnepitope/pep_design.php?sequence=VILRGGGSFVVFLLV&method=hybrid&model=main) | SVM | POSITIVE | 0.92107 |
| 483-493 | [FAPSASAFKK](http://crdd.osdd.net/raghava/ifnepitope/pep_design.php?sequence=FAPSASAFKK&method=hybrid&model=main) | SVM | POSITIVE | 0.91602 |
| 140-155 | [LGTVASQTRAVGERA](http://crdd.osdd.net/raghava/ifnepitope/pep_design.php?sequence=LGTVASQTRAVGERA&method=hybrid&model=main) | SVM | POSITIVE | 0.88559 |
| 41-56 | [RAEETRAETRTRVEE](http://crdd.osdd.net/raghava/ifnepitope/pep_design.php?sequence=RAEETRAETRTRVEE&method=hybrid&model=main) | SVM | POSITIVE | 0.86835 |
| 148-163 | [RAVGERAAKLVGIEL](http://crdd.osdd.net/raghava/ifnepitope/pep_design.php?sequence=RAVGERAAKLVGIEL&method=hybrid&model=main) | SVM | POSITIVE | 0.86574 |
| 480-493 | [IAQFAPSASAFKK](http://crdd.osdd.net/raghava/ifnepitope/pep_design.php?sequence=IAQFAPSASAFKK&method=hybrid&model=main) | SVM | POSITIVE | 0.80752 |
| 167-182 | [KAKFVAAWTLKAAAG](http://crdd.osdd.net/raghava/ifnepitope/pep_design.php?sequence=KAKFVAAWTLKAAAG&method=hybrid&model=main) | SVM | POSITIVE | 0.79461 |
| 153-168 | [RAAKLVGIELEAAAK](http://crdd.osdd.net/raghava/ifnepitope/pep_design.php?sequence=RAAKLVGIELEAAAK&method=hybrid&model=main) | SVM | POSITIVE | 0.79004 |
| 209-224 | [GSFVVFLLVTLGGGS](http://crdd.osdd.net/raghava/ifnepitope/pep_design.php?sequence=GSFVVFLLVTLGGGS&method=hybrid&model=main) | SVM | POSITIVE | 0.78052 |
| 141-156 | [GTVASQTRAVGERAA](http://crdd.osdd.net/raghava/ifnepitope/pep_design.php?sequence=GTVASQTRAVGERAA&method=hybrid&model=main) | SVM | POSITIVE | 0.77149 |
| 481-493 | [AQFAPSASAFKK](http://crdd.osdd.net/raghava/ifnepitope/pep_design.php?sequence=AQFAPSASAFKK&method=hybrid&model=main) | SVM | POSITIVE | 0.75559 |
| 17-32 | [LLAALGAADLALATV](http://crdd.osdd.net/raghava/ifnepitope/pep_design.php?sequence=LLAALGAADLALATV&method=hybrid&model=main) | SVM | POSITIVE | 0.75125 |
| 149-164 | [AVGERAAKLVGIELE](http://crdd.osdd.net/raghava/ifnepitope/pep_design.php?sequence=AVGERAAKLVGIELE&method=hybrid&model=main) | SVM | POSITIVE | 0.72554 |
| 482-493 | [QFAPSASAFKK](http://crdd.osdd.net/raghava/ifnepitope/pep_design.php?sequence=QFAPSASAFKK&method=hybrid&model=main) | SVM | POSITIVE | 0.7198 |
| 40-55 | [ERAEETRAETRTRVE](http://crdd.osdd.net/raghava/ifnepitope/pep_design.php?sequence=ERAEETRAETRTRVE&method=hybrid&model=main) | SVM | POSITIVE | 0.71555 |
| 152-167 | [ERAAKLVGIELEAAA](http://crdd.osdd.net/raghava/ifnepitope/pep_design.php?sequence=ERAAKLVGIELEAAA&method=hybrid&model=main) | SVM | POSITIVE | 0.71058 |
| 38-53 | [LRERAEETRAETRTR](http://crdd.osdd.net/raghava/ifnepitope/pep_design.php?sequence=LRERAEETRAETRTR&method=hybrid&model=main) | SVM | POSITIVE | 0.68872 |
| 18-33 | [LAALGAADLALATVN](http://crdd.osdd.net/raghava/ifnepitope/pep_design.php?sequence=LAALGAADLALATVN&method=hybrid&model=main) | SVM | POSITIVE | 0.68127 |
| 142-157 | [TVASQTRAVGERAAK](http://crdd.osdd.net/raghava/ifnepitope/pep_design.php?sequence=TVASQTRAVGERAAK&method=hybrid&model=main) | SVM | POSITIVE | 0.67359 |
| 169-184 | [KFVAAWTLKAAAGGG](http://crdd.osdd.net/raghava/ifnepitope/pep_design.php?sequence=KFVAAWTLKAAAGGG&method=hybrid&model=main) | SVM | POSITIVE | 0.67221 |
| 212-227 | [VVFLLVTLGGGSAIK](http://crdd.osdd.net/raghava/ifnepitope/pep_design.php?sequence=VVFLLVTLGGGSAIK&method=hybrid&model=main) | SVM | POSITIVE | 0.66971 |
| 39-54 | [RERAEETRAETRTRV](http://crdd.osdd.net/raghava/ifnepitope/pep_design.php?sequence=RERAEETRAETRTRV&method=hybrid&model=main) | SVM | POSITIVE | 0.66965 |
| 210-225 | [SFVVFLLVTLGGGSA](http://crdd.osdd.net/raghava/ifnepitope/pep_design.php?sequence=SFVVFLLVTLGGGSA&method=hybrid&model=main) | SVM | POSITIVE | 0.66581 |
| 211-226 | [FVVFLLVTLGGGSAI](http://crdd.osdd.net/raghava/ifnepitope/pep_design.php?sequence=FVVFLLVTLGGGSAI&method=hybrid&model=main) | SVM | POSITIVE | 0.64505 |
| 44-59 | [ETRAETRTRVEERRA](http://crdd.osdd.net/raghava/ifnepitope/pep_design.php?sequence=ETRAETRTRVEERRA&method=hybrid&model=main) | SVM | POSITIVE | 0.63285 |
| 168-183 | [AKFVAAWTLKAAAGG](http://crdd.osdd.net/raghava/ifnepitope/pep_design.php?sequence=AKFVAAWTLKAAAGG&method=hybrid&model=main) | SVM | POSITIVE | 0.61624 |
| 80-95 | [TEELRKAAEGYLEAA](http://crdd.osdd.net/raghava/ifnepitope/pep_design.php?sequence=TEELRKAAEGYLEAA&method=hybrid&model=main) | SVM | POSITIVE | 0.58031 |
| 202-217 | [AVILRGGGSFVVFLL](http://crdd.osdd.net/raghava/ifnepitope/pep_design.php?sequence=AVILRGGGSFVVFLL&method=hybrid&model=main) | SVM | POSITIVE | 0.578 |
| 45-60 | [TRAETRTRVEERRAR](http://crdd.osdd.net/raghava/ifnepitope/pep_design.php?sequence=TRAETRTRVEERRAR&method=hybrid&model=main) | SVM | POSITIVE | 0.56408 |
| 42-57 | [AEETRAETRTRVEER](http://crdd.osdd.net/raghava/ifnepitope/pep_design.php?sequence=AEETRAETRTRVEER&method=hybrid&model=main) | SVM | POSITIVE | 0.5592 |
| 204-219 | [ILRGGGSFVVFLLVT](http://crdd.osdd.net/raghava/ifnepitope/pep_design.php?sequence=ILRGGGSFVVFLLVT&method=hybrid&model=main) | SVM | POSITIVE | 0.55594 |
| 81-96 | [EELRKAAEGYLEAAT](http://crdd.osdd.net/raghava/ifnepitope/pep_design.php?sequence=EELRKAAEGYLEAAT&method=hybrid&model=main) | SVM | POSITIVE | 0.55007 |
| 457-472 | [GGGKKTEPKKDKKKK](http://crdd.osdd.net/raghava/ifnepitope/pep_design.php?sequence=GGGKKTEPKKDKKKK&method=hybrid&model=main) | SVM | POSITIVE | 0.5409 |
| 201-216 | [GAVILRGGGSFVVFL](http://crdd.osdd.net/raghava/ifnepitope/pep_design.php?sequence=GAVILRGGGSFVVFL&method=hybrid&model=main) | SVM | POSITIVE | 0.5344 |
| 139-154 | [ALGTVASQTRAVGER](http://crdd.osdd.net/raghava/ifnepitope/pep_design.php?sequence=ALGTVASQTRAVGER&method=hybrid&model=main) | SVM | POSITIVE | 0.52937 |
| 170-185 | [FVAAWTLKAAAGGGS](http://crdd.osdd.net/raghava/ifnepitope/pep_design.php?sequence=FVAAWTLKAAAGGGS&method=hybrid&model=main) | SVM | POSITIVE | 0.52759 |
| 464-479 | [PKKDKKKKADKKHWP](http://crdd.osdd.net/raghava/ifnepitope/pep_design.php?sequence=PKKDKKKKADKKHWP&method=hybrid&model=main) | SVM | POSITIVE | 0.51258 |
| 150-165 | [VGERAAKLVGIELEA](http://crdd.osdd.net/raghava/ifnepitope/pep_design.php?sequence=VGERAAKLVGIELEA&method=hybrid&model=main) | SVM | POSITIVE | 0.51028 |
| 16-31 | [PLLAALGAADLALAT](http://crdd.osdd.net/raghava/ifnepitope/pep_design.php?sequence=PLLAALGAADLALAT&method=hybrid&model=main) | SVM | POSITIVE | 0.49927 |
| 465-480 | [KKDKKKKADKKHWPQ](http://crdd.osdd.net/raghava/ifnepitope/pep_design.php?sequence=KKDKKKKADKKHWPQ&method=hybrid&model=main) | SVM | POSITIVE | 0.49512 |
| 462-477 | [TEPKKDKKKKADKKH](http://crdd.osdd.net/raghava/ifnepitope/pep_design.php?sequence=TEPKKDKKKKADKKH&method=hybrid&model=main) | SVM | POSITIVE | 0.49293 |
| 463-478 | [EPKKDKKKKADKKHW](http://crdd.osdd.net/raghava/ifnepitope/pep_design.php?sequence=EPKKDKKKKADKKHW&method=hybrid&model=main) | SVM | POSITIVE | 0.48031 |
| 138-153 | [EALGTVASQTRAVGE](http://crdd.osdd.net/raghava/ifnepitope/pep_design.php?sequence=EALGTVASQTRAVGE&method=hybrid&model=main) | SVM | POSITIVE | 0.4792 |
| 343-358 | [VRQKKVRQIAPGKKM](http://crdd.osdd.net/raghava/ifnepitope/pep_design.php?sequence=VRQKKVRQIAPGKKM&method=hybrid&model=main) | SVM | POSITIVE | 0.47896 |
| 151-166 | [GERAAKLVGIELEAA](http://crdd.osdd.net/raghava/ifnepitope/pep_design.php?sequence=GERAAKLVGIELEAA&method=hybrid&model=main) | SVM | POSITIVE | 0.472 |
| 34-49 | [LIANLRERAEETRAE](http://crdd.osdd.net/raghava/ifnepitope/pep_design.php?sequence=LIANLRERAEETRAE&method=hybrid&model=main) | SVM | POSITIVE | 0.45548 |
| 43-58 | [EETRAETRTRVEERR](http://crdd.osdd.net/raghava/ifnepitope/pep_design.php?sequence=EETRAETRTRVEERR&method=hybrid&model=main) | SVM | POSITIVE | 0.43696 |
| 458-473 | [GGKKTEPKKDKKKKA](http://crdd.osdd.net/raghava/ifnepitope/pep_design.php?sequence=GGKKTEPKKDKKKKA&method=hybrid&model=main) | SVM | POSITIVE | 0.43274 |
| 82-97 | [ELRKAAEGYLEAATN](http://crdd.osdd.net/raghava/ifnepitope/pep_design.php?sequence=ELRKAAEGYLEAATN&method=hybrid&model=main) | SVM | POSITIVE | 0.42302 |
| 37-52 | [NLRERAEETRAETRT](http://crdd.osdd.net/raghava/ifnepitope/pep_design.php?sequence=NLRERAEETRAETRT&method=hybrid&model=main) | SVM | POSITIVE | 0.41405 |
| 251-266 | [PGGPGPGLACFVLAA](http://crdd.osdd.net/raghava/ifnepitope/pep_design.php?sequence=PGGPGPGLACFVLAA&method=hybrid&model=main) | SVM | POSITIVE | 0.41113 |
| 137-152 | [QEALGTVASQTRAVG](http://crdd.osdd.net/raghava/ifnepitope/pep_design.php?sequence=QEALGTVASQTRAVG&method=hybrid&model=main) | SVM | POSITIVE | 0.40993 |
| 86-101 | [AAEGYLEAATNRYNE](http://crdd.osdd.net/raghava/ifnepitope/pep_design.php?sequence=AAEGYLEAATNRYNE&method=hybrid&model=main) | SVM | POSITIVE | 0.40596 |
| 452-467 | [PARMAGGGKKTEPKK](http://crdd.osdd.net/raghava/ifnepitope/pep_design.php?sequence=PARMAGGGKKTEPKK&method=hybrid&model=main) | SVM | POSITIVE | 0.40478 |
| 459-474 | [GKKTEPKKDKKKKAD](http://crdd.osdd.net/raghava/ifnepitope/pep_design.php?sequence=GKKTEPKKDKKKKAD&method=hybrid&model=main) | SVM | POSITIVE | 0.40304 |
| 250-265 | [APGGPGPGLACFVLA](http://crdd.osdd.net/raghava/ifnepitope/pep_design.php?sequence=APGGPGPGLACFVLA&method=hybrid&model=main) | SVM | POSITIVE | 0.40011 |
| 249-264 | [IAPGGPGPGLACFVL](http://crdd.osdd.net/raghava/ifnepitope/pep_design.php?sequence=IAPGGPGPGLACFVL&method=hybrid&model=main) | SVM | POSITIVE | 0.37564 |
| 468-483 | [KKKKADKKHWPQIAQ](http://crdd.osdd.net/raghava/ifnepitope/pep_design.php?sequence=KKKKADKKHWPQIAQ&method=hybrid&model=main) | SVM | POSITIVE | 0.36521 |
| 85-100 | [KAAEGYLEAATNRYN](http://crdd.osdd.net/raghava/ifnepitope/pep_design.php?sequence=KAAEGYLEAATNRYN&method=hybrid&model=main) | SVM | POSITIVE | 0.3645 |
| 460-475 | [KKTEPKKDKKKKADK](http://crdd.osdd.net/raghava/ifnepitope/pep_design.php?sequence=KKTEPKKDKKKKADK&method=hybrid&model=main) | SVM | POSITIVE | 0.35791 |
| 461-476 | [KTEPKKDKKKKADKK](http://crdd.osdd.net/raghava/ifnepitope/pep_design.php?sequence=KTEPKKDKKKKADKK&method=hybrid&model=main) | SVM | POSITIVE | 0.35791 |
| 46-61 | [RAETRTRVEERRARL](http://crdd.osdd.net/raghava/ifnepitope/pep_design.php?sequence=RAETRTRVEERRARL&method=hybrid&model=main) | SVM | POSITIVE | 0.34997 |
| 467-482 | [DKKKKADKKHWPQIA](http://crdd.osdd.net/raghava/ifnepitope/pep_design.php?sequence=DKKKKADKKHWPQIA&method=hybrid&model=main) | SVM | POSITIVE | 0.34908 |
| 83-98 | [LRKAAEGYLEAATNR](http://crdd.osdd.net/raghava/ifnepitope/pep_design.php?sequence=LRKAAEGYLEAATNR&method=hybrid&model=main) | SVM | POSITIVE | 0.34433 |
| 84-99 | [RKAAEGYLEAATNRY](http://crdd.osdd.net/raghava/ifnepitope/pep_design.php?sequence=RKAAEGYLEAATNRY&method=hybrid&model=main) | SVM | POSITIVE | 0.33923 |
| 254-269 | [PGPGLACFVLAAVYR](http://crdd.osdd.net/raghava/ifnepitope/pep_design.php?sequence=PGPGLACFVLAAVYR&method=hybrid&model=main) | SVM | POSITIVE | 0.32136 |
| 256-271 | [PGLACFVLAAVYRIN](http://crdd.osdd.net/raghava/ifnepitope/pep_design.php?sequence=PGLACFVLAAVYRIN&method=hybrid&model=main) | SVM | POSITIVE | 0.31675 |
| 36-51 | [ANLRERAEETRAETR](http://crdd.osdd.net/raghava/ifnepitope/pep_design.php?sequence=ANLRERAEETRAETR&method=hybrid&model=main) | SVM | POSITIVE | 0.30829 |
| 35-50 | [IANLRERAEETRAET](http://crdd.osdd.net/raghava/ifnepitope/pep_design.php?sequence=IANLRERAEETRAET&method=hybrid&model=main) | SVM | POSITIVE | 0.30682 |
| 135-150 | [LTQEALGTVASQTRA](http://crdd.osdd.net/raghava/ifnepitope/pep_design.php?sequence=LTQEALGTVASQTRA&method=hybrid&model=main) | SVM | POSITIVE | 0.30595 |
| 121-136 | [ASARAEGYVDQAVEL](http://crdd.osdd.net/raghava/ifnepitope/pep_design.php?sequence=ASARAEGYVDQAVEL&method=hybrid&model=main) | SVM | POSITIVE | 0.30587 |
| 346-361 | [KKVRQIAPGKKMWSF](http://crdd.osdd.net/raghava/ifnepitope/pep_design.php?sequence=KKVRQIAPGKKMWSF&method=hybrid&model=main) | SVM | POSITIVE | 0.28719 |
| 136-151 | [TQEALGTVASQTRAV](http://crdd.osdd.net/raghava/ifnepitope/pep_design.php?sequence=TQEALGTVASQTRAV&method=hybrid&model=main) | SVM | POSITIVE | 0.28301 |
| 342-357 | [EVRQKKVRQIAPGKK](http://crdd.osdd.net/raghava/ifnepitope/pep_design.php?sequence=EVRQKKVRQIAPGKK&method=hybrid&model=main) | SVM | POSITIVE | 0.28258 |
| 120-135 | [DASARAEGYVDQAVE](http://crdd.osdd.net/raghava/ifnepitope/pep_design.php?sequence=DASARAEGYVDQAVE&method=hybrid&model=main) | SVM | POSITIVE | 0.2742 |
| 87-102 | [AEGYLEAATNRYNEL](http://crdd.osdd.net/raghava/ifnepitope/pep_design.php?sequence=AEGYLEAATNRYNEL&method=hybrid&model=main) | SVM | POSITIVE | 0.2637 |
| 124-139 | [RAEGYVDQAVELTQE](http://crdd.osdd.net/raghava/ifnepitope/pep_design.php?sequence=RAEGYVDQAVELTQE&method=hybrid&model=main) | SVM | POSITIVE | 0.25935 |
| 15-30 | [APLLAALGAADLALA](http://crdd.osdd.net/raghava/ifnepitope/pep_design.php?sequence=APLLAALGAADLALA&method=hybrid&model=main) | SVM | POSITIVE | 0.25404 |
| 448-463 | [RGTSPARMAGGGKKT](http://crdd.osdd.net/raghava/ifnepitope/pep_design.php?sequence=RGTSPARMAGGGKKT&method=hybrid&model=main) | SVM | POSITIVE | 0.25282 |
| 449-464 | [GTSPARMAGGGKKTE](http://crdd.osdd.net/raghava/ifnepitope/pep_design.php?sequence=GTSPARMAGGGKKTE&method=hybrid&model=main) | SVM | POSITIVE | 0.24922 |
| 258-273 | [LACFVLAAVYRINWI](http://crdd.osdd.net/raghava/ifnepitope/pep_design.php?sequence=LACFVLAAVYRINWI&method=hybrid&model=main) | SVM | POSITIVE | 0.24829 |
| 248-263 | [QIAPGGPGPGLACFV](http://crdd.osdd.net/raghava/ifnepitope/pep_design.php?sequence=QIAPGGPGPGLACFV&method=hybrid&model=main) | SVM | POSITIVE | 0.24754 |
| 255-270 | [GPGLACFVLAAVYRI](http://crdd.osdd.net/raghava/ifnepitope/pep_design.php?sequence=GPGLACFVLAAVYRI&method=hybrid&model=main) | SVM | POSITIVE | 0.24462 |
| 456-471 | [AGGGKKTEPKKDKKK](http://crdd.osdd.net/raghava/ifnepitope/pep_design.php?sequence=AGGGKKTEPKKDKKK&method=hybrid&model=main) | SVM | POSITIVE | 0.24328 |
| 79-94 | [TTEELRKAAEGYLEA](http://crdd.osdd.net/raghava/ifnepitope/pep_design.php?sequence=TTEELRKAAEGYLEA&method=hybrid&model=main) | SVM | POSITIVE | 0.2308 |
| 450-465 | [TSPARMAGGGKKTEP](http://crdd.osdd.net/raghava/ifnepitope/pep_design.php?sequence=TSPARMAGGGKKTEP&method=hybrid&model=main) | SVM | POSITIVE | 0.22636 |
| 466-481 | [KDKKKKADKKHWPQI](http://crdd.osdd.net/raghava/ifnepitope/pep_design.php?sequence=KDKKKKADKKHWPQI&method=hybrid&model=main) | SVM | POSITIVE | 0.21323 |
| 447-462 | [SRGTSPARMAGGGKK](http://crdd.osdd.net/raghava/ifnepitope/pep_design.php?sequence=SRGTSPARMAGGGKK&method=hybrid&model=main) | SVM | POSITIVE | 0.20811 |
| 446-461 | [SSRGTSPARMAGGGK](http://crdd.osdd.net/raghava/ifnepitope/pep_design.php?sequence=SSRGTSPARMAGGGK&method=hybrid&model=main) | SVM | POSITIVE | 0.2059 |
| 445-460 | [GSSRGTSPARMAGGG](http://crdd.osdd.net/raghava/ifnepitope/pep_design.php?sequence=GSSRGTSPARMAGGG&method=hybrid&model=main) | SVM | POSITIVE | 0.20093 |
| 253-268 | [GPGPGLACFVLAAVY](http://crdd.osdd.net/raghava/ifnepitope/pep_design.php?sequence=GPGPGLACFVLAAVY&method=hybrid&model=main) | SVM | POSITIVE | 0.19967 |
| 122-137 | [SARAEGYVDQAVELT](http://crdd.osdd.net/raghava/ifnepitope/pep_design.php?sequence=SARAEGYVDQAVELT&method=hybrid&model=main) | SVM | POSITIVE | 0.19607 |
| 88-103 | [EGYLEAATNRYNELV](http://crdd.osdd.net/raghava/ifnepitope/pep_design.php?sequence=EGYLEAATNRYNELV&method=hybrid&model=main) | SVM | POSITIVE | 0.18595 |
| 344-359 | [RQKKVRQIAPGKKMW](http://crdd.osdd.net/raghava/ifnepitope/pep_design.php?sequence=RQKKVRQIAPGKKMW&method=hybrid&model=main) | SVM | POSITIVE | 0.18054 |
| 92-107 | [EAATNRYNELVERGE](http://crdd.osdd.net/raghava/ifnepitope/pep_design.php?sequence=EAATNRYNELVERGE&method=hybrid&model=main) | SVM | POSITIVE | 0.17927 |
| 93-108 | [AATNRYNELVERGEA](http://crdd.osdd.net/raghava/ifnepitope/pep_design.php?sequence=AATNRYNELVERGEA&method=hybrid&model=main) | SVM | POSITIVE | 0.17927 |
| 94-109 | [ATNRYNELVERGEAA](http://crdd.osdd.net/raghava/ifnepitope/pep_design.php?sequence=ATNRYNELVERGEAA&method=hybrid&model=main) | SVM | POSITIVE | 0.17927 |
| 246-261 | [VRQIAPGGPGPGLAC](http://crdd.osdd.net/raghava/ifnepitope/pep_design.php?sequence=VRQIAPGGPGPGLAC&method=hybrid&model=main) | SVM | POSITIVE | 0.16354 |
| 451-466 | [SPARMAGGGKKTEPK](http://crdd.osdd.net/raghava/ifnepitope/pep_design.php?sequence=SPARMAGGGKKTEPK&method=hybrid&model=main) | SVM | POSITIVE | 0.15601 |
| 218-233 | [TLGGGSAIKLDDKDP](http://crdd.osdd.net/raghava/ifnepitope/pep_design.php?sequence=TLGGGSAIKLDDKDP&method=hybrid&model=main) | SVM | POSITIVE | 0.14325 |
| 91-106 | [LEAATNRYNELVERG](http://crdd.osdd.net/raghava/ifnepitope/pep_design.php?sequence=LEAATNRYNELVERG&method=hybrid&model=main) | SVM | POSITIVE | 0.1428 |
| 469-484 | [KKKADKKHWPQIAQF](http://crdd.osdd.net/raghava/ifnepitope/pep_design.php?sequence=KKKADKKHWPQIAQF&method=hybrid&model=main) | SVM | POSITIVE | 0.13712 |
| 337-352 | [KKKGDEVRQKKVRQI](http://crdd.osdd.net/raghava/ifnepitope/pep_design.php?sequence=KKKGDEVRQKKVRQI&method=hybrid&model=main) | SVM | POSITIVE | 0.13705 |
| 341-356 | [DEVRQKKVRQIAPGK](http://crdd.osdd.net/raghava/ifnepitope/pep_design.php?sequence=DEVRQKKVRQIAPGK&method=hybrid&model=main) | SVM | POSITIVE | 0.13109 |
| 473-488 | [DKKHWPQIAQFAPSA](http://crdd.osdd.net/raghava/ifnepitope/pep_design.php?sequence=DKKHWPQIAQFAPSA&method=hybrid&model=main) | SVM | POSITIVE | 0.12379 |
| 455-470 | [MAGGGKKTEPKKDKK](http://crdd.osdd.net/raghava/ifnepitope/pep_design.php?sequence=MAGGGKKTEPKKDKK&method=hybrid&model=main) | SVM | POSITIVE | 0.12277 |
| 33-48 | [DLIANLRERAEETRA](http://crdd.osdd.net/raghava/ifnepitope/pep_design.php?sequence=DLIANLRERAEETRA&method=hybrid&model=main) | SVM | POSITIVE | 0.12149 |
| 345-360 | [QKKVRQIAPGKKMWS](http://crdd.osdd.net/raghava/ifnepitope/pep_design.php?sequence=QKKVRQIAPGKKMWS&method=hybrid&model=main) | SVM | POSITIVE | 0.12133 |
| 257-272 | [GLACFVLAAVYRINW](http://crdd.osdd.net/raghava/ifnepitope/pep_design.php?sequence=GLACFVLAAVYRINW&method=hybrid&model=main) | SVM | POSITIVE | 0.11887 |
| 336-351 | [YKKKGDEVRQKKVRQ](http://crdd.osdd.net/raghava/ifnepitope/pep_design.php?sequence=YKKKGDEVRQKKVRQ&method=hybrid&model=main) | SVM | POSITIVE | 0.11152 |
| 223-238 | [SAIKLDDKDPGPGPG](http://crdd.osdd.net/raghava/ifnepitope/pep_design.php?sequence=SAIKLDDKDPGPGPG&method=hybrid&model=main) | SVM | POSITIVE | 0.11009 |
| 244-259 | [DEVRQIAPGGPGPGL](http://crdd.osdd.net/raghava/ifnepitope/pep_design.php?sequence=DEVRQIAPGGPGPGL&method=hybrid&model=main) | SVM | POSITIVE | 0.11 |
| 247-262 | [RQIAPGGPGPGLACF](http://crdd.osdd.net/raghava/ifnepitope/pep_design.php?sequence=RQIAPGGPGPGLACF&method=hybrid&model=main) | SVM | POSITIVE | 0.10972 |
| 245-260 | [EVRQIAPGGPGPGLA](http://crdd.osdd.net/raghava/ifnepitope/pep_design.php?sequence=EVRQIAPGGPGPGLA&method=hybrid&model=main) | SVM | POSITIVE | 0.10675 |
| 95-110 | [TNRYNELVERGEAAL](http://crdd.osdd.net/raghava/ifnepitope/pep_design.php?sequence=TNRYNELVERGEAAL&method=hybrid&model=main) | SVM | POSITIVE | 0.10441 |
| 97-112 | [RYNELVERGEAALQR](http://crdd.osdd.net/raghava/ifnepitope/pep_design.php?sequence=RYNELVERGEAALQR&method=hybrid&model=main) | SVM | POSITIVE | 0.09905 |
| 252-267 | [GGPGPGLACFVLAAV](http://crdd.osdd.net/raghava/ifnepitope/pep_design.php?sequence=GGPGPGLACFVLAAV&method=hybrid&model=main) | SVM | POSITIVE | 0.09839 |
| 384-399 | [KAMACLVGLMKKYVY](http://crdd.osdd.net/raghava/ifnepitope/pep_design.php?sequence=KAMACLVGLMKKYVY&method=hybrid&model=main) | SVM | POSITIVE | 0.09751 |
| 219-234 | [LGGGSAIKLDDKDPG](http://crdd.osdd.net/raghava/ifnepitope/pep_design.php?sequence=LGGGSAIKLDDKDPG&method=hybrid&model=main) | SVM | POSITIVE | 0.09084 |
| 123-138 | [ARAEGYVDQAVELTQ](http://crdd.osdd.net/raghava/ifnepitope/pep_design.php?sequence=ARAEGYVDQAVELTQ&method=hybrid&model=main) | SVM | POSITIVE | 0.08553 |
| 29-44 | [ATVNDLIANLRERAE](http://crdd.osdd.net/raghava/ifnepitope/pep_design.php?sequence=ATVNDLIANLRERAE&method=hybrid&model=main) | SVM | POSITIVE | 0.07625 |
| 266-281 | [VYRINWIGPGPGVTL](http://crdd.osdd.net/raghava/ifnepitope/pep_design.php?sequence=VYRINWIGPGPGVTL&method=hybrid&model=main) | SVM | POSITIVE | 0.06146 |
| 104-119 | [RGEAALQRLRSQTAF](http://crdd.osdd.net/raghava/ifnepitope/pep_design.php?sequence=RGEAALQRLRSQTAF&method=hybrid&model=main) | SVM | POSITIVE | 0.06045 |
| 119-134 | [EDASARAEGYVDQAV](http://crdd.osdd.net/raghava/ifnepitope/pep_design.php?sequence=EDASARAEGYVDQAV&method=hybrid&model=main) | SVM | POSITIVE | 0.05152 |
| 105-120 | [GEAALQRLRSQTAFE](http://crdd.osdd.net/raghava/ifnepitope/pep_design.php?sequence=GEAALQRLRSQTAFE&method=hybrid&model=main) | SVM | POSITIVE | 0.04796 |
| 96-111 | [NRYNELVERGEAALQ](http://crdd.osdd.net/raghava/ifnepitope/pep_design.php?sequence=NRYNELVERGEAALQ&method=hybrid&model=main) | SVM | POSITIVE | 0.04684 |
| 114-129 | [SQTAFEDASARAEGY](http://crdd.osdd.net/raghava/ifnepitope/pep_design.php?sequence=SQTAFEDASARAEGY&method=hybrid&model=main) | SVM | POSITIVE | 0.0441 |
| 265-280 | [AVYRINWIGPGPGVT](http://crdd.osdd.net/raghava/ifnepitope/pep_design.php?sequence=AVYRINWIGPGPGVT&method=hybrid&model=main) | SVM | POSITIVE | 0.03759 |
| 338-353 | [KKGDEVRQKKVRQIA](http://crdd.osdd.net/raghava/ifnepitope/pep_design.php?sequence=KKGDEVRQKKVRQIA&method=hybrid&model=main) | SVM | POSITIVE | 0.03544 |
| 339-354 | [KGDEVRQKKVRQIAP](http://crdd.osdd.net/raghava/ifnepitope/pep_design.php?sequence=KGDEVRQKKVRQIAP&method=hybrid&model=main) | SVM | POSITIVE | 0.02761 |
| 453-468 | [ARMAGGGKKTEPKKD](http://crdd.osdd.net/raghava/ifnepitope/pep_design.php?sequence=ARMAGGGKKTEPKKD&method=hybrid&model=main) | SVM | POSITIVE | 0.02188 |
| 129-144 | [VDQAVELTQEALGTV](http://crdd.osdd.net/raghava/ifnepitope/pep_design.php?sequence=VDQAVELTQEALGTV&method=hybrid&model=main) | SVM | POSITIVE | 0.02102 |
| 98-113 | [YNELVERGEAALQRL](http://crdd.osdd.net/raghava/ifnepitope/pep_design.php?sequence=YNELVERGEAALQRL&method=hybrid&model=main) | SVM | POSITIVE | 0.01551 |
| 126-141 | [EGYVDQAVELTQEAL](http://crdd.osdd.net/raghava/ifnepitope/pep_design.php?sequence=EGYVDQAVELTQEAL&method=hybrid&model=main) | SVM | POSITIVE | 0.01382 |
| 89-104 | [GYLEAATNRYNELVE](http://crdd.osdd.net/raghava/ifnepitope/pep_design.php?sequence=GYLEAATNRYNELVE&method=hybrid&model=main) | SVM | POSITIVE | 0.01 |
| 264-279 | [AAVYRINWIGPGPGV](http://crdd.osdd.net/raghava/ifnepitope/pep_design.php?sequence=AAVYRINWIGPGPGV&method=hybrid&model=main) | SVM | POSITIVE | 0.00871 |
| 134-149 | [ELTQEALGTVASQTR](http://crdd.osdd.net/raghava/ifnepitope/pep_design.php?sequence=ELTQEALGTVASQTR&method=hybrid&model=main) | SVM | POSITIVE | 0.00535 |
| 335-350 | [NYKKKGDEVRQKKVR](http://crdd.osdd.net/raghava/ifnepitope/pep_design.php?sequence=NYKKKGDEVRQKKVR&method=hybrid&model=main) | SVM | NEGATIVE | -0.0003 |
| 259-274 | [ACFVLAAVYRINWIG](http://crdd.osdd.net/raghava/ifnepitope/pep_design.php?sequence=ACFVLAAVYRINWIG&method=hybrid&model=main) | SVM | NEGATIVE | -0.0016 |
| 318-333 | [DEVRQIAPGQTGKIA](http://crdd.osdd.net/raghava/ifnepitope/pep_design.php?sequence=DEVRQIAPGQTGKIA&method=hybrid&model=main) | SVM | NEGATIVE | -0.0023 |
| 243-258 | [GDEVRQIAPGGPGPG](http://crdd.osdd.net/raghava/ifnepitope/pep_design.php?sequence=GDEVRQIAPGGPGPG&method=hybrid&model=main) | SVM | NEGATIVE | -0.0043 |
| 340-355 | [GDEVRQKKVRQIAPG](http://crdd.osdd.net/raghava/ifnepitope/pep_design.php?sequence=GDEVRQKKVRQIAPG&method=hybrid&model=main) | SVM | NEGATIVE | -0.0119 |
| 454-469 | [RMAGGGKKTEPKKDK](http://crdd.osdd.net/raghava/ifnepitope/pep_design.php?sequence=RMAGGGKKTEPKKDK&method=hybrid&model=main) | SVM | NEGATIVE | -0.0143 |
| 78-93 | [FTTEELRKAAEGYLE](http://crdd.osdd.net/raghava/ifnepitope/pep_design.php?sequence=FTTEELRKAAEGYLE&method=hybrid&model=main) | SVM | NEGATIVE | -0.0175 |
| 347-362 | [KVRQIAPGKKMWSFN](http://crdd.osdd.net/raghava/ifnepitope/pep_design.php?sequence=KVRQIAPGKKMWSFN&method=hybrid&model=main) | SVM | NEGATIVE | -0.0254 |
| 470-485 | [KKADKKHWPQIAQFA](http://crdd.osdd.net/raghava/ifnepitope/pep_design.php?sequence=KKADKKHWPQIAQFA&method=hybrid&model=main) | SVM | NEGATIVE | -0.0293 |
| 26-41 | [LALATVNDLIANLRE](http://crdd.osdd.net/raghava/ifnepitope/pep_design.php?sequence=LALATVNDLIANLRE&method=hybrid&model=main) | SVM | NEGATIVE | -0.0332 |
| 125-140 | [AEGYVDQAVELTQEA](http://crdd.osdd.net/raghava/ifnepitope/pep_design.php?sequence=AEGYVDQAVELTQEA&method=hybrid&model=main) | SVM | NEGATIVE | -0.0391 |
| 76-91 | [DKFTTEELRKAAEGY](http://crdd.osdd.net/raghava/ifnepitope/pep_design.php?sequence=DKFTTEELRKAAEGY&method=hybrid&model=main) | SVM | NEGATIVE | -0.0403 |
| 115-130 | [QTAFEDASARAEGYV](http://crdd.osdd.net/raghava/ifnepitope/pep_design.php?sequence=QTAFEDASARAEGYV&method=hybrid&model=main) | SVM | NEGATIVE | -0.0453 |
| 28-43 | [LATVNDLIANLRERA](http://crdd.osdd.net/raghava/ifnepitope/pep_design.php?sequence=LATVNDLIANLRERA&method=hybrid&model=main) | SVM | NEGATIVE | -0.0475 |
| 113-128 | [RSQTAFEDASARAEG](http://crdd.osdd.net/raghava/ifnepitope/pep_design.php?sequence=RSQTAFEDASARAEG&method=hybrid&model=main) | SVM | NEGATIVE | -0.0484 |
| 116-131 | [TAFEDASARAEGYVD](http://crdd.osdd.net/raghava/ifnepitope/pep_design.php?sequence=TAFEDASARAEGYVD&method=hybrid&model=main) | SVM | NEGATIVE | -0.0549 |
| 90-105 | [YLEAATNRYNELVER](http://crdd.osdd.net/raghava/ifnepitope/pep_design.php?sequence=YLEAATNRYNELVER&method=hybrid&model=main) | SVM | NEGATIVE | -0.0552 |
| 133-148 | [VELTQEALGTVASQT](http://crdd.osdd.net/raghava/ifnepitope/pep_design.php?sequence=VELTQEALGTVASQT&method=hybrid&model=main) | SVM | NEGATIVE | -0.0596 |
| 118-133 | [FEDASARAEGYVDQA](http://crdd.osdd.net/raghava/ifnepitope/pep_design.php?sequence=FEDASARAEGYVDQA&method=hybrid&model=main) | SVM | NEGATIVE | -0.0615 |
| 0-15 | [EAAKMAENPNIDDLP](http://crdd.osdd.net/raghava/ifnepitope/pep_design.php?sequence=EAAKMAENPNIDDLP&method=hybrid&model=main) | SVM | NEGATIVE | -0.0628 |
| 99-114 | [NELVERGEAALQRLR](http://crdd.osdd.net/raghava/ifnepitope/pep_design.php?sequence=NELVERGEAALQRLR&method=hybrid&model=main) | SVM | NEGATIVE | -0.0641 |
| 47-62 | [AETRTRVEERRARLT](http://crdd.osdd.net/raghava/ifnepitope/pep_design.php?sequence=AETRTRVEERRARLT&method=hybrid&model=main) | SVM | NEGATIVE | -0.0704 |
| 319-334 | [EVRQIAPGQTGKIAD](http://crdd.osdd.net/raghava/ifnepitope/pep_design.php?sequence=EVRQIAPGQTGKIAD&method=hybrid&model=main) | SVM | NEGATIVE | -0.0711 |
| 103-118 | [ERGEAALQRLRSQTA](http://crdd.osdd.net/raghava/ifnepitope/pep_design.php?sequence=ERGEAALQRLRSQTA&method=hybrid&model=main) | SVM | NEGATIVE | -0.0764 |
| 408-423 | [RVPKKPSFYVYSRVK](http://crdd.osdd.net/raghava/ifnepitope/pep_design.php?sequence=RVPKKPSFYVYSRVK&method=hybrid&model=main) | SVM | NEGATIVE | -0.0776 |
| 75-90 | [RDKFTTEELRKAAEG](http://crdd.osdd.net/raghava/ifnepitope/pep_design.php?sequence=RDKFTTEELRKAAEG&method=hybrid&model=main) | SVM | NEGATIVE | -0.0823 |
| 130-145 | [DQAVELTQEALGTVA](http://crdd.osdd.net/raghava/ifnepitope/pep_design.php?sequence=DQAVELTQEALGTVA&method=hybrid&model=main) | SVM | NEGATIVE | -0.084 |
| 132-147 | [AVELTQEALGTVASQ](http://crdd.osdd.net/raghava/ifnepitope/pep_design.php?sequence=AVELTQEALGTVASQ&method=hybrid&model=main) | SVM | NEGATIVE | -0.092 |
| 77-92 | [KFTTEELRKAAEGYL](http://crdd.osdd.net/raghava/ifnepitope/pep_design.php?sequence=KFTTEELRKAAEGYL&method=hybrid&model=main) | SVM | NEGATIVE | -0.0943 |
| 74-89 | [LRDKFTTEELRKAAE](http://crdd.osdd.net/raghava/ifnepitope/pep_design.php?sequence=LRDKFTTEELRKAAE&method=hybrid&model=main) | SVM | NEGATIVE | -0.0973 |
| 102-117 | [VERGEAALQRLRSQT](http://crdd.osdd.net/raghava/ifnepitope/pep_design.php?sequence=VERGEAALQRLRSQT&method=hybrid&model=main) | SVM | NEGATIVE | -0.103 |
| 188-203 | [NYKLPDGGGSLVIGA](http://crdd.osdd.net/raghava/ifnepitope/pep_design.php?sequence=NYKLPDGGGSLVIGA&method=hybrid&model=main) | SVM | NEGATIVE | -0.1056 |
| 128-143 | [YVDQAVELTQEALGT](http://crdd.osdd.net/raghava/ifnepitope/pep_design.php?sequence=YVDQAVELTQEALGT&method=hybrid&model=main) | SVM | NEGATIVE | -0.1144 |
| 186-201 | [DYNYKLPDGGGSLVI](http://crdd.osdd.net/raghava/ifnepitope/pep_design.php?sequence=DYNYKLPDGGGSLVI&method=hybrid&model=main) | SVM | NEGATIVE | -0.1156 |
| 127-142 | [GYVDQAVELTQEALG](http://crdd.osdd.net/raghava/ifnepitope/pep_design.php?sequence=GYVDQAVELTQEALG&method=hybrid&model=main) | SVM | NEGATIVE | -0.1196 |
| 389-404 | [LVGLMKKYVYSRVKN](http://crdd.osdd.net/raghava/ifnepitope/pep_design.php?sequence=LVGLMKKYVYSRVKN&method=hybrid&model=main) | SVM | NEGATIVE | -0.1263 |
| 224-239 | [AIKLDDKDPGPGPGS](http://crdd.osdd.net/raghava/ifnepitope/pep_design.php?sequence=AIKLDDKDPGPGPGS&method=hybrid&model=main) | SVM | NEGATIVE | -0.1322 |
| 399-414 | [SRVKNLNSSRVPKKP](http://crdd.osdd.net/raghava/ifnepitope/pep_design.php?sequence=SRVKNLNSSRVPKKP&method=hybrid&model=main) | SVM | NEGATIVE | -0.147 |
| 117-132 | [AFEDASARAEGYVDQ](http://crdd.osdd.net/raghava/ifnepitope/pep_design.php?sequence=AFEDASARAEGYVDQ&method=hybrid&model=main) | SVM | NEGATIVE | -0.1546 |
| 131-146 | [QAVELTQEALGTVAS](http://crdd.osdd.net/raghava/ifnepitope/pep_design.php?sequence=QAVELTQEALGTVAS&method=hybrid&model=main) | SVM | NEGATIVE | -0.1563 |
| 407-422 | [SRVPKKPSFYVYSRV](http://crdd.osdd.net/raghava/ifnepitope/pep_design.php?sequence=SRVPKKPSFYVYSRV&method=hybrid&model=main) | SVM | NEGATIVE | -0.1637 |
| 30-45 | [TVNDLIANLRERAEE](http://crdd.osdd.net/raghava/ifnepitope/pep_design.php?sequence=TVNDLIANLRERAEE&method=hybrid&model=main) | SVM | NEGATIVE | -0.1653 |
| 267-282 | [YRINWIGPGPGVTLA](http://crdd.osdd.net/raghava/ifnepitope/pep_design.php?sequence=YRINWIGPGPGVTLA&method=hybrid&model=main) | SVM | NEGATIVE | -0.1671 |
| 187-202 | [YNYKLPDGGGSLVIG](http://crdd.osdd.net/raghava/ifnepitope/pep_design.php?sequence=YNYKLPDGGGSLVIG&method=hybrid&model=main) | SVM | NEGATIVE | -0.1674 |
| 225-240 | [IKLDDKDPGPGPGSF](http://crdd.osdd.net/raghava/ifnepitope/pep_design.php?sequence=IKLDDKDPGPGPGSF&method=hybrid&model=main) | SVM | NEGATIVE | -0.1696 |
| 49-64 | [TRTRVEERRARLTKF](http://crdd.osdd.net/raghava/ifnepitope/pep_design.php?sequence=TRTRVEERRARLTKF&method=hybrid&model=main) | SVM | NEGATIVE | -0.1708 |
| 268-283 | [RINWIGPGPGVTLAI](http://crdd.osdd.net/raghava/ifnepitope/pep_design.php?sequence=RINWIGPGPGVTLAI&method=hybrid&model=main) | SVM | NEGATIVE | -0.1882 |
| 400-415 | [RVKNLNSSRVPKKPS](http://crdd.osdd.net/raghava/ifnepitope/pep_design.php?sequence=RVKNLNSSRVPKKPS&method=hybrid&model=main) | SVM | NEGATIVE | -0.1891 |
| 471-486 | [KADKKHWPQIAQFAP](http://crdd.osdd.net/raghava/ifnepitope/pep_design.php?sequence=KADKKHWPQIAQFAP&method=hybrid&model=main) | SVM | NEGATIVE | -0.195 |
| 14-29 | [PAPLLAALGAADLAL](http://crdd.osdd.net/raghava/ifnepitope/pep_design.php?sequence=PAPLLAALGAADLAL&method=hybrid&model=main) | SVM | NEGATIVE | -0.1958 |
| 261-276 | [FVLAAVYRINWIGPG](http://crdd.osdd.net/raghava/ifnepitope/pep_design.php?sequence=FVLAAVYRINWIGPG&method=hybrid&model=main) | SVM | NEGATIVE | -0.2007 |
| 101-116 | [LVERGEAALQRLRSQ](http://crdd.osdd.net/raghava/ifnepitope/pep_design.php?sequence=LVERGEAALQRLRSQ&method=hybrid&model=main) | SVM | NEGATIVE | -0.2085 |
| 228-243 | [DDKDPGPGPGSFVIR](http://crdd.osdd.net/raghava/ifnepitope/pep_design.php?sequence=DDKDPGPGPGSFVIR&method=hybrid&model=main) | SVM | NEGATIVE | -0.2089 |
| 354-369 | [GKKMWSFNPETNKKL](http://crdd.osdd.net/raghava/ifnepitope/pep_design.php?sequence=GKKMWSFNPETNKKL&method=hybrid&model=main) | SVM | NEGATIVE | -0.2106 |
| 227-242 | [LDDKDPGPGPGSFVI](http://crdd.osdd.net/raghava/ifnepitope/pep_design.php?sequence=LDDKDPGPGPGSFVI&method=hybrid&model=main) | SVM | NEGATIVE | -0.2131 |
| 348-363 | [VRQIAPGKKMWSFNP](http://crdd.osdd.net/raghava/ifnepitope/pep_design.php?sequence=VRQIAPGKKMWSFNP&method=hybrid&model=main) | SVM | NEGATIVE | -0.215 |
| 260-275 | [CFVLAAVYRINWIGP](http://crdd.osdd.net/raghava/ifnepitope/pep_design.php?sequence=CFVLAAVYRINWIGP&method=hybrid&model=main) | SVM | NEGATIVE | -0.2185 |
| 396-411 | [YVYSRVKNLNSSRVP](http://crdd.osdd.net/raghava/ifnepitope/pep_design.php?sequence=YVYSRVKNLNSSRVP&method=hybrid&model=main) | SVM | NEGATIVE | -0.2246 |
| 416-431 | [YVYSRVKNLNSSRVP](http://crdd.osdd.net/raghava/ifnepitope/pep_design.php?sequence=YVYSRVKNLNSSRVP&method=hybrid&model=main) | SVM | NEGATIVE | -0.2246 |
| 333-348 | [DYNYKKKGDEVRQKK](http://crdd.osdd.net/raghava/ifnepitope/pep_design.php?sequence=DYNYKKKGDEVRQKK&method=hybrid&model=main) | SVM | NEGATIVE | -0.2258 |
| 221-236 | [GGSAIKLDDKDPGPG](http://crdd.osdd.net/raghava/ifnepitope/pep_design.php?sequence=GGSAIKLDDKDPGPG&method=hybrid&model=main) | SVM | NEGATIVE | -0.2266 |
| 27-42 | [ALATVNDLIANLRER](http://crdd.osdd.net/raghava/ifnepitope/pep_design.php?sequence=ALATVNDLIANLRER&method=hybrid&model=main) | SVM | NEGATIVE | -0.2275 |
| 294-309 | [PGPGDLSPRWYFYYL](http://crdd.osdd.net/raghava/ifnepitope/pep_design.php?sequence=PGPGDLSPRWYFYYL&method=hybrid&model=main) | SVM | NEGATIVE | -0.2283 |
| 472-487 | [ADKKHWPQIAQFAPS](http://crdd.osdd.net/raghava/ifnepitope/pep_design.php?sequence=ADKKHWPQIAQFAPS&method=hybrid&model=main) | SVM | NEGATIVE | -0.2381 |
| 263-278 | [LAAVYRINWIGPGPG](http://crdd.osdd.net/raghava/ifnepitope/pep_design.php?sequence=LAAVYRINWIGPGPG&method=hybrid&model=main) | SVM | NEGATIVE | -0.2408 |
| 229-244 | [DKDPGPGPGSFVIRG](http://crdd.osdd.net/raghava/ifnepitope/pep_design.php?sequence=DKDPGPGPGSFVIRG&method=hybrid&model=main) | SVM | NEGATIVE | -0.2411 |
| 329-344 | [GKIADYNYKKKGDEV](http://crdd.osdd.net/raghava/ifnepitope/pep_design.php?sequence=GKIADYNYKKKGDEV&method=hybrid&model=main) | SVM | NEGATIVE | -0.2476 |
| 226-241 | [KLDDKDPGPGPGSFV](http://crdd.osdd.net/raghava/ifnepitope/pep_design.php?sequence=KLDDKDPGPGPGSFV&method=hybrid&model=main) | SVM | NEGATIVE | -0.2487 |
| 398-413 | [YSRVKNLNSSRVPKK](http://crdd.osdd.net/raghava/ifnepitope/pep_design.php?sequence=YSRVKNLNSSRVPKK&method=hybrid&model=main) | SVM | NEGATIVE | -0.2521 |
| 418-433 | [YSRVKNLNSSRVPKK](http://crdd.osdd.net/raghava/ifnepitope/pep_design.php?sequence=YSRVKNLNSSRVPKK&method=hybrid&model=main) | SVM | NEGATIVE | -0.2521 |
| 220-235 | [GGGSAIKLDDKDPGP](http://crdd.osdd.net/raghava/ifnepitope/pep_design.php?sequence=GGGSAIKLDDKDPGP&method=hybrid&model=main) | SVM | NEGATIVE | -0.2566 |
| 353-368 | [PGKKMWSFNPETNKK](http://crdd.osdd.net/raghava/ifnepitope/pep_design.php?sequence=PGKKMWSFNPETNKK&method=hybrid&model=main) | SVM | NEGATIVE | -0.2574 |
| 48-63 | [ETRTRVEERRARLTK](http://crdd.osdd.net/raghava/ifnepitope/pep_design.php?sequence=ETRTRVEERRARLTK&method=hybrid&model=main) | SVM | NEGATIVE | -0.2687 |
| 334-349 | [YNYKKKGDEVRQKKV](http://crdd.osdd.net/raghava/ifnepitope/pep_design.php?sequence=YNYKKKGDEVRQKKV&method=hybrid&model=main) | SVM | NEGATIVE | -0.2771 |
| 13-28 | [LPAPLLAALGAADLA](http://crdd.osdd.net/raghava/ifnepitope/pep_design.php?sequence=LPAPLLAALGAADLA&method=hybrid&model=main) | SVM | NEGATIVE | -0.2993 |
| 283-298 | [LTALRLCAYCGPGPG](http://crdd.osdd.net/raghava/ifnepitope/pep_design.php?sequence=LTALRLCAYCGPGPG&method=hybrid&model=main) | SVM | NEGATIVE | -0.3036 |
| 397-412 | [VYSRVKNLNSSRVPK](http://crdd.osdd.net/raghava/ifnepitope/pep_design.php?sequence=VYSRVKNLNSSRVPK&method=hybrid&model=main) | SVM | NEGATIVE | -0.3066 |
| 417-432 | [VYSRVKNLNSSRVPK](http://crdd.osdd.net/raghava/ifnepitope/pep_design.php?sequence=VYSRVKNLNSSRVPK&method=hybrid&model=main) | SVM | NEGATIVE | -0.3066 |
| 184-199 | [SADYNYKLPDGGGSL](http://crdd.osdd.net/raghava/ifnepitope/pep_design.php?sequence=SADYNYKLPDGGGSL&method=hybrid&model=main) | SVM | NEGATIVE | -0.3079 |
| 100-115 | [ELVERGEAALQRLRS](http://crdd.osdd.net/raghava/ifnepitope/pep_design.php?sequence=ELVERGEAALQRLRS&method=hybrid&model=main) | SVM | NEGATIVE | -0.3159 |
| 331-346 | [IADYNYKKKGDEVRQ](http://crdd.osdd.net/raghava/ifnepitope/pep_design.php?sequence=IADYNYKKKGDEVRQ&method=hybrid&model=main) | SVM | NEGATIVE | -0.3175 |
| 281-296 | [AILTALRLCAYCGPG](http://crdd.osdd.net/raghava/ifnepitope/pep_design.php?sequence=AILTALRLCAYCGPG&method=hybrid&model=main) | SVM | NEGATIVE | -0.3226 |
| 240-255 | [VIRGDEVRQIAPGGP](http://crdd.osdd.net/raghava/ifnepitope/pep_design.php?sequence=VIRGDEVRQIAPGGP&method=hybrid&model=main) | SVM | NEGATIVE | -0.3281 |
| 415-430 | [FYVYSRVKNLNSSRV](http://crdd.osdd.net/raghava/ifnepitope/pep_design.php?sequence=FYVYSRVKNLNSSRV&method=hybrid&model=main) | SVM | NEGATIVE | -0.3302 |
| Jan-16 | [AAKMAENPNIDDLPA](http://crdd.osdd.net/raghava/ifnepitope/pep_design.php?sequence=AAKMAENPNIDDLPA&method=hybrid&model=main) | SVM | NEGATIVE | -0.3337 |
| 317-332 | [GDEVRQIAPGQTGKI](http://crdd.osdd.net/raghava/ifnepitope/pep_design.php?sequence=GDEVRQIAPGQTGKI&method=hybrid&model=main) | SVM | NEGATIVE | -0.3349 |
| 50-65 | [RTRVEERRARLTKFQ](http://crdd.osdd.net/raghava/ifnepitope/pep_design.php?sequence=RTRVEERRARLTKFQ&method=hybrid&model=main) | SVM | NEGATIVE | -0.3371 |
| 316-331 | [RGDEVRQIAPGQTGK](http://crdd.osdd.net/raghava/ifnepitope/pep_design.php?sequence=RGDEVRQIAPGQTGK&method=hybrid&model=main) | SVM | NEGATIVE | -0.3436 |
| 330-345 | [KIADYNYKKKGDEVR](http://crdd.osdd.net/raghava/ifnepitope/pep_design.php?sequence=KIADYNYKKKGDEVR&method=hybrid&model=main) | SVM | NEGATIVE | -0.3718 |
| 296-311 | [PGDLSPRWYFYYLGT](http://crdd.osdd.net/raghava/ifnepitope/pep_design.php?sequence=PGDLSPRWYFYYLGT&method=hybrid&model=main) | SVM | NEGATIVE | -0.3722 |
| 349-364 | [RQIAPGKKMWSFNPE](http://crdd.osdd.net/raghava/ifnepitope/pep_design.php?sequence=RQIAPGKKMWSFNPE&method=hybrid&model=main) | SVM | NEGATIVE | -0.3752 |
| 404-419 | [LNSSRVPKKPSFYVY](http://crdd.osdd.net/raghava/ifnepitope/pep_design.php?sequence=LNSSRVPKKPSFYVY&method=hybrid&model=main) | SVM | NEGATIVE | -0.3837 |
| 314-329 | [KIRGDEVRQIAPGQT](http://crdd.osdd.net/raghava/ifnepitope/pep_design.php?sequence=KIRGDEVRQIAPGQT&method=hybrid&model=main) | SVM | NEGATIVE | -0.3846 |
| 67-82 | [LPEQFIELRDKFTTE](http://crdd.osdd.net/raghava/ifnepitope/pep_design.php?sequence=LPEQFIELRDKFTTE&method=hybrid&model=main) | SVM | NEGATIVE | -0.397 |
| 66-81 | [DLPEQFIELRDKFTT](http://crdd.osdd.net/raghava/ifnepitope/pep_design.php?sequence=DLPEQFIELRDKFTT&method=hybrid&model=main) | SVM | NEGATIVE | -0.3983 |
| 328-343 | [TGKIADYNYKKKGDE](http://crdd.osdd.net/raghava/ifnepitope/pep_design.php?sequence=TGKIADYNYKKKGDE&method=hybrid&model=main) | SVM | NEGATIVE | -0.4072 |
| 313-328 | [KKIRGDEVRQIAPGQ](http://crdd.osdd.net/raghava/ifnepitope/pep_design.php?sequence=KKIRGDEVRQIAPGQ&method=hybrid&model=main) | SVM | NEGATIVE | -0.4099 |
| Apr-19 | [MAENPNIDDLPAPLL](http://crdd.osdd.net/raghava/ifnepitope/pep_design.php?sequence=MAENPNIDDLPAPLL&method=hybrid&model=main) | SVM | NEGATIVE | -0.4134 |
| Dec-27 | [DLPAPLLAALGAADL](http://crdd.osdd.net/raghava/ifnepitope/pep_design.php?sequence=DLPAPLLAALGAADL&method=hybrid&model=main) | SVM | NEGATIVE | -0.4165 |
| 222-237 | [GSAIKLDDKDPGPGP](http://crdd.osdd.net/raghava/ifnepitope/pep_design.php?sequence=GSAIKLDDKDPGPGP&method=hybrid&model=main) | SVM | NEGATIVE | -0.4197 |
| 315-330 | [IRGDEVRQIAPGQTG](http://crdd.osdd.net/raghava/ifnepitope/pep_design.php?sequence=IRGDEVRQIAPGQTG&method=hybrid&model=main) | SVM | NEGATIVE | -0.4236 |
| 241-256 | [IRGDEVRQIAPGGPG](http://crdd.osdd.net/raghava/ifnepitope/pep_design.php?sequence=IRGDEVRQIAPGGPG&method=hybrid&model=main) | SVM | NEGATIVE | -0.4237 |
| 62-77 | [KFQEDLPEQFIELRD](http://crdd.osdd.net/raghava/ifnepitope/pep_design.php?sequence=KFQEDLPEQFIELRD&method=hybrid&model=main) | SVM | NEGATIVE | -0.4304 |
| 312-327 | [PKKIRGDEVRQIAPG](http://crdd.osdd.net/raghava/ifnepitope/pep_design.php?sequence=PKKIRGDEVRQIAPG&method=hybrid&model=main) | SVM | NEGATIVE | -0.4377 |
| Feb-17 | [AKMAENPNIDDLPAP](http://crdd.osdd.net/raghava/ifnepitope/pep_design.php?sequence=AKMAENPNIDDLPAP&method=hybrid&model=main) | SVM | NEGATIVE | -0.4527 |
| Jun-21 | [ENPNIDDLPAPLLAA](http://crdd.osdd.net/raghava/ifnepitope/pep_design.php?sequence=ENPNIDDLPAPLLAA&method=hybrid&model=main) | SVM | NEGATIVE | -0.4543 |
| 262-277 | [VLAAVYRINWIGPGP](http://crdd.osdd.net/raghava/ifnepitope/pep_design.php?sequence=VLAAVYRINWIGPGP&method=hybrid&model=main) | SVM | NEGATIVE | -0.4593 |
| 63-78 | [FQEDLPEQFIELRDK](http://crdd.osdd.net/raghava/ifnepitope/pep_design.php?sequence=FQEDLPEQFIELRDK&method=hybrid&model=main) | SVM | NEGATIVE | -0.4624 |
| 293-308 | [GPGPGDLSPRWYFYY](http://crdd.osdd.net/raghava/ifnepitope/pep_design.php?sequence=GPGPGDLSPRWYFYY&method=hybrid&model=main) | SVM | NEGATIVE | -0.4777 |
| Mar-18 | [KMAENPNIDDLPAPL](http://crdd.osdd.net/raghava/ifnepitope/pep_design.php?sequence=KMAENPNIDDLPAPL&method=hybrid&model=main) | SVM | NEGATIVE | -0.4841 |
| 332-347 | [ADYNYKKKGDEVRQK](http://crdd.osdd.net/raghava/ifnepitope/pep_design.php?sequence=ADYNYKKKGDEVRQK&method=hybrid&model=main) | SVM | NEGATIVE | -0.4948 |
| 242-257 | [RGDEVRQIAPGGPGP](http://crdd.osdd.net/raghava/ifnepitope/pep_design.php?sequence=RGDEVRQIAPGGPGP&method=hybrid&model=main) | SVM | NEGATIVE | -0.5048 |
| 185-200 | [ADYNYKLPDGGGSLV](http://crdd.osdd.net/raghava/ifnepitope/pep_design.php?sequence=ADYNYKLPDGGGSLV&method=hybrid&model=main) | SVM | NEGATIVE | -0.5056 |
| 61-76 | [TKFQEDLPEQFIELR](http://crdd.osdd.net/raghava/ifnepitope/pep_design.php?sequence=TKFQEDLPEQFIELR&method=hybrid&model=main) | SVM | NEGATIVE | -0.5099 |
| Aug-23 | [PNIDDLPAPLLAALG](http://crdd.osdd.net/raghava/ifnepitope/pep_design.php?sequence=PNIDDLPAPLLAALG&method=hybrid&model=main) | SVM | NEGATIVE | -0.5219 |
| 297-312 | [GDLSPRWYFYYLGTG](http://crdd.osdd.net/raghava/ifnepitope/pep_design.php?sequence=GDLSPRWYFYYLGTG&method=hybrid&model=main) | SVM | NEGATIVE | -0.5227 |
| 280-295 | [LAILTALRLCAYCGP](http://crdd.osdd.net/raghava/ifnepitope/pep_design.php?sequence=LAILTALRLCAYCGP&method=hybrid&model=main) | SVM | NEGATIVE | -0.5288 |
| 295-310 | [GPGDLSPRWYFYYLG](http://crdd.osdd.net/raghava/ifnepitope/pep_design.php?sequence=GPGDLSPRWYFYYLG&method=hybrid&model=main) | SVM | NEGATIVE | -0.5374 |
| 51-66 | [TRVEERRARLTKFQE](http://crdd.osdd.net/raghava/ifnepitope/pep_design.php?sequence=TRVEERRARLTKFQE&method=hybrid&model=main) | SVM | NEGATIVE | -0.5378 |
| 351-366 | [IAPGKKMWSFNPETN](http://crdd.osdd.net/raghava/ifnepitope/pep_design.php?sequence=IAPGKKMWSFNPETN&method=hybrid&model=main) | SVM | NEGATIVE | -0.5408 |
| 64-79 | [QEDLPEQFIELRDKF](http://crdd.osdd.net/raghava/ifnepitope/pep_design.php?sequence=QEDLPEQFIELRDKF&method=hybrid&model=main) | SVM | NEGATIVE | -0.5479 |
| 403-418 | [NLNSSRVPKKPSFYV](http://crdd.osdd.net/raghava/ifnepitope/pep_design.php?sequence=NLNSSRVPKKPSFYV&method=hybrid&model=main) | SVM | NEGATIVE | -0.5551 |
| 355-370 | [KKMWSFNPETNKKLF](http://crdd.osdd.net/raghava/ifnepitope/pep_design.php?sequence=KKMWSFNPETNKKLF&method=hybrid&model=main) | SVM | NEGATIVE | -0.5558 |
| 311-326 | [GPKKIRGDEVRQIAP](http://crdd.osdd.net/raghava/ifnepitope/pep_design.php?sequence=GPKKIRGDEVRQIAP&method=hybrid&model=main) | SVM | NEGATIVE | -0.5615 |
| 52-67 | [RVEERRARLTKFQED](http://crdd.osdd.net/raghava/ifnepitope/pep_design.php?sequence=RVEERRARLTKFQED&method=hybrid&model=main) | SVM | NEGATIVE | -0.5642 |
| 406-421 | [SSRVPKKPSFYVYSR](http://crdd.osdd.net/raghava/ifnepitope/pep_design.php?sequence=SSRVPKKPSFYVYSR&method=hybrid&model=main) | SVM | NEGATIVE | -0.5656 |
| Sep-24 | [NIDDLPAPLLAALGA](http://crdd.osdd.net/raghava/ifnepitope/pep_design.php?sequence=NIDDLPAPLLAALGA&method=hybrid&model=main) | SVM | NEGATIVE | -0.5793 |
| 352-367 | [APGKKMWSFNPETNK](http://crdd.osdd.net/raghava/ifnepitope/pep_design.php?sequence=APGKKMWSFNPETNK&method=hybrid&model=main) | SVM | NEGATIVE | -0.584 |
| 350-365 | [QIAPGKKMWSFNPET](http://crdd.osdd.net/raghava/ifnepitope/pep_design.php?sequence=QIAPGKKMWSFNPET&method=hybrid&model=main) | SVM | NEGATIVE | -0.5851 |
| 32-47 | [NDLIANLRERAEETR](http://crdd.osdd.net/raghava/ifnepitope/pep_design.php?sequence=NDLIANLRERAEETR&method=hybrid&model=main) | SVM | NEGATIVE | -0.5922 |
| 401-416 | [VKNLNSSRVPKKPSF](http://crdd.osdd.net/raghava/ifnepitope/pep_design.php?sequence=VKNLNSSRVPKKPSF&method=hybrid&model=main) | SVM | NEGATIVE | -0.6085 |
| 282-297 | [ILTALRLCAYCGPGP](http://crdd.osdd.net/raghava/ifnepitope/pep_design.php?sequence=ILTALRLCAYCGPGP&method=hybrid&model=main) | SVM | NEGATIVE | -0.616 |
| 60-75 | [LTKFQEDLPEQFIEL](http://crdd.osdd.net/raghava/ifnepitope/pep_design.php?sequence=LTKFQEDLPEQFIEL&method=hybrid&model=main) | SVM | NEGATIVE | -0.6227 |
| 439-454 | [NIVKKPGSSRGTSPA](http://crdd.osdd.net/raghava/ifnepitope/pep_design.php?sequence=NIVKKPGSSRGTSPA&method=hybrid&model=main) | SVM | NEGATIVE | -0.6288 |
| 413-428 | [PSFYVYSRVKNLNSS](http://crdd.osdd.net/raghava/ifnepitope/pep_design.php?sequence=PSFYVYSRVKNLNSS&method=hybrid&model=main) | SVM | NEGATIVE | -0.6307 |
| 409-424 | [VPKKPSFYVYSRVKN](http://crdd.osdd.net/raghava/ifnepitope/pep_design.php?sequence=VPKKPSFYVYSRVKN&method=hybrid&model=main) | SVM | NEGATIVE | -0.6323 |
| 65-80 | [EDLPEQFIELRDKFT](http://crdd.osdd.net/raghava/ifnepitope/pep_design.php?sequence=EDLPEQFIELRDKFT&method=hybrid&model=main) | SVM | NEGATIVE | -0.635 |
| 307-322 | [YLGTGPKKIRGDEVR](http://crdd.osdd.net/raghava/ifnepitope/pep_design.php?sequence=YLGTGPKKIRGDEVR&method=hybrid&model=main) | SVM | NEGATIVE | -0.6369 |
| 366-381 | [KKLFARTRSMWSFNP](http://crdd.osdd.net/raghava/ifnepitope/pep_design.php?sequence=KKLFARTRSMWSFNP&method=hybrid&model=main) | SVM | NEGATIVE | -0.6463 |
| 284-299 | [TALRLCAYCGPGPGD](http://crdd.osdd.net/raghava/ifnepitope/pep_design.php?sequence=TALRLCAYCGPGPGD&method=hybrid&model=main) | SVM | NEGATIVE | -0.6535 |
| 31-46 | [VNDLIANLRERAEET](http://crdd.osdd.net/raghava/ifnepitope/pep_design.php?sequence=VNDLIANLRERAEET&method=hybrid&model=main) | SVM | NEGATIVE | -0.6571 |
| Oct-25 | [IDDLPAPLLAALGAA](http://crdd.osdd.net/raghava/ifnepitope/pep_design.php?sequence=IDDLPAPLLAALGAA&method=hybrid&model=main) | SVM | NEGATIVE | -0.6699 |
| 365-380 | [NKKLFARTRSMWSFN](http://crdd.osdd.net/raghava/ifnepitope/pep_design.php?sequence=NKKLFARTRSMWSFN&method=hybrid&model=main) | SVM | NEGATIVE | -0.6852 |
| 301-316 | [PRWYFYYLGTGPKKI](http://crdd.osdd.net/raghava/ifnepitope/pep_design.php?sequence=PRWYFYYLGTGPKKI&method=hybrid&model=main) | SVM | NEGATIVE | -0.6873 |
| 53-68 | [VEERRARLTKFQEDL](http://crdd.osdd.net/raghava/ifnepitope/pep_design.php?sequence=VEERRARLTKFQEDL&method=hybrid&model=main) | SVM | NEGATIVE | -0.6916 |
| 414-429 | [SFYVYSRVKNLNSSR](http://crdd.osdd.net/raghava/ifnepitope/pep_design.php?sequence=SFYVYSRVKNLNSSR&method=hybrid&model=main) | SVM | NEGATIVE | -0.6931 |
| 411-426 | [KKPSFYVYSRVKNLN](http://crdd.osdd.net/raghava/ifnepitope/pep_design.php?sequence=KKPSFYVYSRVKNLN&method=hybrid&model=main) | SVM | NEGATIVE | -0.6968 |
| 405-420 | [NSSRVPKKPSFYVYS](http://crdd.osdd.net/raghava/ifnepitope/pep_design.php?sequence=NSSRVPKKPSFYVYS&method=hybrid&model=main) | SVM | NEGATIVE | -0.6971 |
| 56-71 | [RRARLTKFQEDLPEQ](http://crdd.osdd.net/raghava/ifnepitope/pep_design.php?sequence=RRARLTKFQEDLPEQ&method=hybrid&model=main) | SVM | NEGATIVE | -0.7003 |
| 298-313 | [DLSPRWYFYYLGTGP](http://crdd.osdd.net/raghava/ifnepitope/pep_design.php?sequence=DLSPRWYFYYLGTGP&method=hybrid&model=main) | SVM | NEGATIVE | -0.7072 |
| May-20 | [AENPNIDDLPAPLLA](http://crdd.osdd.net/raghava/ifnepitope/pep_design.php?sequence=AENPNIDDLPAPLLA&method=hybrid&model=main) | SVM | NEGATIVE | -0.7122 |
| 292-307 | [CGPGPGDLSPRWYFY](http://crdd.osdd.net/raghava/ifnepitope/pep_design.php?sequence=CGPGPGDLSPRWYFY&method=hybrid&model=main) | SVM | NEGATIVE | -0.7148 |
| 300-315 | [SPRWYFYYLGTGPKK](http://crdd.osdd.net/raghava/ifnepitope/pep_design.php?sequence=SPRWYFYYLGTGPKK&method=hybrid&model=main) | SVM | NEGATIVE | -0.7174 |
| 306-321 | [YYLGTGPKKIRGDEV](http://crdd.osdd.net/raghava/ifnepitope/pep_design.php?sequence=YYLGTGPKKIRGDEV&method=hybrid&model=main) | SVM | NEGATIVE | -0.7196 |
| Nov-26 | [DDLPAPLLAALGAAD](http://crdd.osdd.net/raghava/ifnepitope/pep_design.php?sequence=DDLPAPLLAALGAAD&method=hybrid&model=main) | SVM | NEGATIVE | -0.7235 |
| 285-300 | [ALRLCAYCGPGPGDL](http://crdd.osdd.net/raghava/ifnepitope/pep_design.php?sequence=ALRLCAYCGPGPGDL&method=hybrid&model=main) | SVM | NEGATIVE | -0.7288 |
| 302-317 | [RWYFYYLGTGPKKIR](http://crdd.osdd.net/raghava/ifnepitope/pep_design.php?sequence=RWYFYYLGTGPKKIR&method=hybrid&model=main) | SVM | NEGATIVE | -0.7312 |
| 303-318 | [WYFYYLGTGPKKIRG](http://crdd.osdd.net/raghava/ifnepitope/pep_design.php?sequence=WYFYYLGTGPKKIRG&method=hybrid&model=main) | SVM | NEGATIVE | -0.7384 |
| 308-323 | [LGTGPKKIRGDEVRQ](http://crdd.osdd.net/raghava/ifnepitope/pep_design.php?sequence=LGTGPKKIRGDEVRQ&method=hybrid&model=main) | SVM | NEGATIVE | -0.7398 |
| Jul-22 | [NPNIDDLPAPLLAAL](http://crdd.osdd.net/raghava/ifnepitope/pep_design.php?sequence=NPNIDDLPAPLLAAL&method=hybrid&model=main) | SVM | NEGATIVE | -0.7489 |
| 291-306 | [YCGPGPGDLSPRWYF](http://crdd.osdd.net/raghava/ifnepitope/pep_design.php?sequence=YCGPGPGDLSPRWYF&method=hybrid&model=main) | SVM | NEGATIVE | -0.7501 |
| 437-452 | [CCNIVKKPGSSRGTS](http://crdd.osdd.net/raghava/ifnepitope/pep_design.php?sequence=CCNIVKKPGSSRGTS&method=hybrid&model=main) | SVM | NEGATIVE | -0.7538 |
| 57-72 | [RARLTKFQEDLPEQF](http://crdd.osdd.net/raghava/ifnepitope/pep_design.php?sequence=RARLTKFQEDLPEQF&method=hybrid&model=main) | SVM | NEGATIVE | -0.7579 |
| 58-73 | [ARLTKFQEDLPEQFI](http://crdd.osdd.net/raghava/ifnepitope/pep_design.php?sequence=ARLTKFQEDLPEQFI&method=hybrid&model=main) | SVM | NEGATIVE | -0.7605 |
| 402-417 | [KNLNSSRVPKKPSFY](http://crdd.osdd.net/raghava/ifnepitope/pep_design.php?sequence=KNLNSSRVPKKPSFY&method=hybrid&model=main) | SVM | NEGATIVE | -0.769 |
| 412-427 | [KPSFYVYSRVKNLNS](http://crdd.osdd.net/raghava/ifnepitope/pep_design.php?sequence=KPSFYVYSRVKNLNS&method=hybrid&model=main) | SVM | NEGATIVE | -0.7751 |
| 310-325 | [TGPKKIRGDEVRQIA](http://crdd.osdd.net/raghava/ifnepitope/pep_design.php?sequence=TGPKKIRGDEVRQIA&method=hybrid&model=main) | SVM | NEGATIVE | -0.7823 |
| 410-425 | [PKKPSFYVYSRVKNL](http://crdd.osdd.net/raghava/ifnepitope/pep_design.php?sequence=PKKPSFYVYSRVKNL&method=hybrid&model=main) | SVM | NEGATIVE | -0.7857 |
| 59-74 | [RLTKFQEDLPEQFIE](http://crdd.osdd.net/raghava/ifnepitope/pep_design.php?sequence=RLTKFQEDLPEQFIE&method=hybrid&model=main) | SVM | NEGATIVE | -0.7924 |
| 290-305 | [AYCGPGPGDLSPRWY](http://crdd.osdd.net/raghava/ifnepitope/pep_design.php?sequence=AYCGPGPGDLSPRWY&method=hybrid&model=main) | SVM | NEGATIVE | -0.8031 |
| 378-393 | [FNPETKKAMACLVGL](http://crdd.osdd.net/raghava/ifnepitope/pep_design.php?sequence=FNPETKKAMACLVGL&method=hybrid&model=main) | SVM | NEGATIVE | -0.8055 |
| 286-301 | [LRLCAYCGPGPGDLS](http://crdd.osdd.net/raghava/ifnepitope/pep_design.php?sequence=LRLCAYCGPGPGDLS&method=hybrid&model=main) | SVM | NEGATIVE | -0.8066 |
| 367-382 | [KLFARTRSMWSFNPE](http://crdd.osdd.net/raghava/ifnepitope/pep_design.php?sequence=KLFARTRSMWSFNPE&method=hybrid&model=main) | SVM | NEGATIVE | -0.8085 |
| 55-70 | [ERRARLTKFQEDLPE](http://crdd.osdd.net/raghava/ifnepitope/pep_design.php?sequence=ERRARLTKFQEDLPE&method=hybrid&model=main) | SVM | NEGATIVE | -0.8175 |
| 299-314 | [LSPRWYFYYLGTGPK](http://crdd.osdd.net/raghava/ifnepitope/pep_design.php?sequence=LSPRWYFYYLGTGPK&method=hybrid&model=main) | SVM | NEGATIVE | -0.8349 |
| 54-69 | [EERRARLTKFQEDLP](http://crdd.osdd.net/raghava/ifnepitope/pep_design.php?sequence=EERRARLTKFQEDLP&method=hybrid&model=main) | SVM | NEGATIVE | -0.8635 |
| 309-324 | [GTGPKKIRGDEVRQI](http://crdd.osdd.net/raghava/ifnepitope/pep_design.php?sequence=GTGPKKIRGDEVRQI&method=hybrid&model=main) | SVM | NEGATIVE | -0.8688 |
| 289-304 | [CAYCGPGPGDLSPRW](http://crdd.osdd.net/raghava/ifnepitope/pep_design.php?sequence=CAYCGPGPGDLSPRW&method=hybrid&model=main) | SVM | NEGATIVE | -0.8734 |
| 438-453 | [CNIVKKPGSSRGTSP](http://crdd.osdd.net/raghava/ifnepitope/pep_design.php?sequence=CNIVKKPGSSRGTSP&method=hybrid&model=main) | SVM | NEGATIVE | -0.8876 |
| 305-320 | [FYYLGTGPKKIRGDE](http://crdd.osdd.net/raghava/ifnepitope/pep_design.php?sequence=FYYLGTGPKKIRGDE&method=hybrid&model=main) | SVM | NEGATIVE | -0.9188 |
| 287-302 | [RLCAYCGPGPGDLSP](http://crdd.osdd.net/raghava/ifnepitope/pep_design.php?sequence=RLCAYCGPGPGDLSP&method=hybrid&model=main) | SVM | NEGATIVE | -0.957 |
| 304-319 | [YFYYLGTGPKKIRGD](http://crdd.osdd.net/raghava/ifnepitope/pep_design.php?sequence=YFYYLGTGPKKIRGD&method=hybrid&model=main) | SVM | NEGATIVE | -0.9702 |
| 377-392 | [SFNPETKKAMACLVG](http://crdd.osdd.net/raghava/ifnepitope/pep_design.php?sequence=SFNPETKKAMACLVG&method=hybrid&model=main) | SVM | NEGATIVE | -1.0471 |
| 288-303 | [LCAYCGPGPGDLSPR](http://crdd.osdd.net/raghava/ifnepitope/pep_design.php?sequence=LCAYCGPGPGDLSPR&method=hybrid&model=main) | SVM | NEGATIVE | -1.0641 |
| 371-386 | [RTRSMWSFNPETKKA](http://crdd.osdd.net/raghava/ifnepitope/pep_design.php?sequence=RTRSMWSFNPETKKA&method=hybrid&model=main) | SVM | NEGATIVE | -1.1062 |
| 368-383 | [LFARTRSMWSFNPET](http://crdd.osdd.net/raghava/ifnepitope/pep_design.php?sequence=LFARTRSMWSFNPET&method=hybrid&model=main) | SVM | NEGATIVE | -1.1275 |
| 375-390 | [MWSFNPETKKAMACL](http://crdd.osdd.net/raghava/ifnepitope/pep_design.php?sequence=MWSFNPETKKAMACL&method=hybrid&model=main) | SVM | NEGATIVE | -1.1624 |
| 370-385 | [ARTRSMWSFNPETKK](http://crdd.osdd.net/raghava/ifnepitope/pep_design.php?sequence=ARTRSMWSFNPETKK&method=hybrid&model=main) | SVM | NEGATIVE | -1.1847 |
| 376-391 | [WSFNPETKKAMACLV](http://crdd.osdd.net/raghava/ifnepitope/pep_design.php?sequence=WSFNPETKKAMACLV&method=hybrid&model=main) | SVM | NEGATIVE | -1.1925 |
| 369-384 | [FARTRSMWSFNPETK](http://crdd.osdd.net/raghava/ifnepitope/pep_design.php?sequence=FARTRSMWSFNPETK&method=hybrid&model=main) | SVM | NEGATIVE | -1.1972 |
| 374-389 | [SMWSFNPETKKAMAC](http://crdd.osdd.net/raghava/ifnepitope/pep_design.php?sequence=SMWSFNPETKKAMAC&method=hybrid&model=main) | SVM | NEGATIVE | -1.1998 |
| 372-387 | [TRSMWSFNPETKKAM](http://crdd.osdd.net/raghava/ifnepitope/pep_design.php?sequence=TRSMWSFNPETKKAM&method=hybrid&model=main) | SVM | NEGATIVE | -1.2596 |
| 373-388 | [RSMWSFNPETKKAMA](http://crdd.osdd.net/raghava/ifnepitope/pep_design.php?sequence=RSMWSFNPETKKAMA&method=hybrid&model=main) | SVM | NEGATIVE | -1.3469 |
