## Supplementary File 1 for "Genome based Evolutionary study of SARS-CoV-2 towards the Prediction of Epitope Based Chimeric Vaccine"

**Supplementary Table 1:** NCBI IDs of the complete genome, spike glycoprotein, envelope protein, membrane protein and nucleocapsid protein of coronavirus

**Table 1.1:** NCBI IDs of the complete genome of coronavirus

| NCBI_IDs | Short Description | Genera;Sub-genera |
| --- | --- | --- |
| NC_010646 | NC 010646 Beluga Whale coronavirus SW1 | Gammacoronavirus; Cegacovirus |
| NC_001451 | NC 001451 Avian infectious bronchitis virus | Gammacoronavirus; Igacovirus |
| NC_010800 | NC 010800 Turkey coronavirus | Gammacoronavirus; Igacovirus |
| NC_035191 | NC 035191 Wencheng Sm shrew coronavirus isolate Xingguo01 | Alphacoronavirus; unclassified |
| NC_034972 | NC 034972 Coronavirus AcCoV.JC34 | Alphacoronavirus; unclassified |
| NC_032730 | NC 032730 Lucheng Rn rat coronavirus isolate Lucheng9 | Alphacoronavirus; Luchacovirus. |
| NC_009988 | NC 009988 Bat coronavirus HKU2 | Alphacoronavirus; Rhinacovirus |
| NC_028824 | NC 028824 BtRf.AlphaCoV-YN2012 | Alphacoronavirus; unclassified |
| NC_023760 | NC 023760 Mink coronavirus strain WD1127 | Alphacoronavirus; Minacovirus |
| NC_030292 | NC 030292 Ferret coronavirus isolate FRCoV.NL010 | Alphacoronavirus; Minacovirus |
| LC119077 | LC119077 Ferret coronavirus genomic RNA strain FRCoV4370 | Alphacoronavirus; Minacovirus |
| NC_002306 | NC 002306 Feline infectious peritonitis virus | Alphacoronavirus; Minacovirus |
| NC_038861 | NC 038861 Transmissible gastroenteritis virus complete genome | Alphacoronavirus; Tegacovirus |
| NC_028806 | NC 028806 Swine enteric coronavirus strain Italy-213306-2009 | Alphacoronavirus; Tegacovirus |
| LT545990 | LT545990 Swine enteric coronavirus isolate SeCoV GER L00930 2012 genome assembly monopartite | Alphacoronavirus; Tegacovirus |
| NC_002645 | NC 002645 Human coronavirus 229E | Alphacoronavirus; Duvinacovirus |
| NC_028752 | NC 028752 Camel alphacoronavirus isolate camel-Riyadh-Ry141-2015 | Alphacoronavirus; Duvinacovirus |
| KY073745 | KY073745 NL63.related bat coronavirus strain BtKYNL63.9b | Alphacoronavirus; Setracovirus |
| NC_032107 | NC 032107 NL63.related bat coronavirus strain BtKYNL63.9a | Alphacoronavirus; Setracovirus |
| NC_005831 | NC 005831 Human Coronavirus NL63 | Alphacoronavirus; Setracovirus |
| NC_028811 | NC 028811 BtMr.AlphaCoV-SAX2011 | Alphacoronavirus; Myotacovirus |
| NC_028833 | NC 028833 BtNv.AlphaCoV-SC2013 | Alphacoronavirus; Nyctacovirus |
| NC_022103 | NC 022103 Bat coronavirus CDPHE15-USA-2006 | Alphacoronavirus; Colacovirus |
| NC_009657 | NC 009657 Scotophilus bat coronavirus 512 | Alphacoronavirus; Pedacovirus |
| NC_010437 | NC 010437 Bat coronavirus 1A | Alphacoronavirus; unclassified |
| NC_010438 | NC 010438 Bat coronavirus HKU8 | Alphacoronavirus; Minunacovirus |
| NC_018871 | NC 018871 Rousettus bat coronavirus HKU10 | Alphacoronavirus; Decacovirus |
| NC_028814 | NC 028814 BtRf.AlphaCoV-HuB2013 | Alphacoronavirus; Decacovirus |
| NC_006577 | NC 006577 Human coronavirus HKU1 | Betacoronavirus; Embecovirus |
| NC_012936 | NC 012936 Rat coronavirus Parker | Betacoronavirus; Embecovirus |
| NC_001846 | NC 001846 Mouse hepatitis virus strain MHV.A59 C12 mutant | Betacoronavirus; Embecovirus |
| AY700211 | AY700211 Murine hepatitis virus strain A59 | Betacoronavirus; Embecovirus |
| NC_026011 | NC 026011 Betacoronavirus HKU24 strain HKU24.R05005I | Betacoronavirus; Embecovirus |
| NC_017083 | NC 017083 Rabbit coronavirus HKU14 | Betacoronavirus; unclassified |
| NC_006213 | NC 006213 Human coronavirus OC43 strain ATCC VR.759 | Betacoronavirus; Embecovirus |
| NC_003045 | NC 003045 Bovine coronavirus | Betacoronavirus; Embecovirus |
| KC545386 | KC545386 Betacoronavirus Erinaceus-VMC-DEU-2012 isolate ErinaceusCoV-201216-GER-2012 | Betacoronavirus; Merbecovirus |
| NC_039207 | NC 039207 Betacoronavirus Erinaceus-VMC-DEU-2012 isolate ErinaceusCoV-2012-174-GER-2012 | Betacoronavirus; Merbecovirus |
| NC_009020 | NC 009020 Bat coronavirus HKU5 | Betacoronavirus; Merbecovirus |
| NC_009019 | NC 009019 Bat coronavirus HKU4 | Betacoronavirus; Merbecovirus |
| NC_038294 | NC 038294 Betacoronavirus England 1 | Betacoronavirus; Merbecovirus |
| NC_019843 | NC 019843 Middle East respiratory syndrome coronavirus | Betacoronavirus; Merbecovirus |
| NC_009021 | NC 009021 Bat coronavirus HKU9 | Betacoronavirus; Nobecovirus |
| NC_030886 | NC 030886 Rousettus bat coronavirus isolate GCCDC1 356 | Betacoronavirus; Nobecovirus |
| NC_025217 | NC 025217 Bat Hp.betacoronavirus-Zhejiang2013 | Betacoronavirus; Hibecovirus |
| NC_004718 | NC 004718 SARS coronavirus | Betacoronavirus; Sarbecovirus |
| MN997409 | MN997409 Wuhan seafood market pneumonia virus isolate 2019.nCoV-USA.AZ1-2020 | Betacoronavirus; Sarbecovirus |
| MN938384 | MN938384 Wuhan seafood market pneumonia virus isolate 2019.nCoV HKU.SZ.002a 2020 | Betacoronavirus; Sarbecovirus |
| MN975262 | MN975262 Wuhan seafood market pneumonia virus isolate 2019.nCoV HKU.SZ.005b 2020 | Betacoronavirus; Sarbecovirus |
| MN985325 | MN985325 Wuhan seafood market pneumonia virus isolate 2019.nCoV-USA.WA1-2020 | Betacoronavirus; Sarbecovirus |
| MN994467 | MN994467 Wuhan seafood market pneumonia virus isolate 2019.nCoV-USA.CA1-2020 | Betacoronavirus; Sarbecovirus |
| MN988713 | MN988713 Wuhan seafood market pneumonia virus isolate 2019.nCoV-USA.IL1-2020 | Betacoronavirus; Sarbecovirus |
| MN994468 | MN994468 Severe acute respiratory syndrome coronavirus 2 isolate 2019-nCoV-USA-CA2-2020 | Betacoronavirus; Sarbecovirus |
| MN996528 | MN996528 Wuhan seafood market pneumonia virus isolate WIV04 | Betacoronavirus; Sarbecovirus |
| NC_045512 | NC 045512 Wuhan seafood market pneumonia virus isolate Wuhan.Hu | Betacoronavirus; Sarbecovirus |
| MN988669 | MN988669 Wuhan seafood market pneumonia virus isolate 2019.nCoV WHU02 | Betacoronavirus; Sarbecovirus |
| MN988668 | MN988668 Wuhan seafood market pneumonia virus isolate 2019.nCoV WHU01 | Betacoronavirus; Sarbecovirus |
| MN996531 | MN996531 Wuhan seafood market pneumonia virus isolate WIV07 | Betacoronavirus; Sarbecovirus |
| MN996530 | MN996530 Wuhan seafood market pneumonia virus isolate WIV06 | Betacoronavirus; Sarbecovirus |
| MN996529 | MN996529 Wuhan seafood market pneumonia virus isolate WIV05 | Betacoronavirus; Sarbecovirus |
| MN996527 | MN996527 Wuhan seafood market pneumonia virus isolate WIV02 | Betacoronavirus; Sarbecovirus |

**Table 1.2:** Spike Proteins

| NCBI ID | Type | Genera, Sub-Genera | Length |
| --- | --- | --- | --- |
| YP_009273005.1 | Rousettus bat | Betacoronavirus; Nobecovirus | 1290 |
| YP_001039971.1 | Rousettus bat | Betacoronavirus; Nobecovirus | 1247 |
| YP_009072440.1 | Bat Hp-betacoronavirus | Betacoronavirus; Hibecovirus | 1317 |
| NP_828851.1 | *SARS-CoV* | Betacoronavirus; Sarbecovirus | 1255 |
| QIC53213.1 | SARS-CoV-2 | Betacoronavirus; Sarbecovirus | 1273 |
| QHR63280.2 | SARS-CoV-2 | Betacoronavirus; Sarbecovirus | 1273 |
| QHR63290.2 | SARS-CoV-2 | Betacoronavirus; Sarbecovirus | 1273 |
| QHR63260.2 | SARS-CoV-2 | Betacoronavirus; Sarbecovirus | 1273 |
| QHR63250.2 | SARS-CoV-2 | Betacoronavirus; Sarbecovirus | 1273 |
| YP_009724390.1 | SARS-CoV-2 | Betacoronavirus; Sarbecovirus | 1273 |
| YP_003767.1 | *HCoV-NL63* | Alphacoronavirus; Setracovirus | 1356 |
| YP_001941166.1 | Turkey coronavirus | Gammacoronavirus; Igacovirus | 1226 |
| YP_001039962.1 | Pipistrellus bat coronavirus | Betacoronavirus; Merbecovirus | 1352 |
| YP_001039953.1 | Tylonycteris bat coronavirus HKU4 | Betacoronavirus; Merbecovirus | 1352 |
| YP_007188579.1 | Betacoronavirus England 1 | Betacoronavirus; Merbecovirus | 1253 |
| YP_009047204.1 | MERS | Betacoronavirus; Merbecovirus | 1353 |
| YP_173238.1 | HCoV-HKU1 | Betacoronavirus; Embecovirus | 1356 |
| YP_003029848.1 | *Rat coronavirus Parker* | Betacoronavirus; Embecovirus | 1360 |
| NP_045300.1 E2 | Murine hepatitis virus | Betacoronavirus; Embecovirus | 1324 |
| AAU06356.1 | Murine hepatitis virus | Betacoronavirus; Embecovirus | 1324 |
| YP_009113025.1 | Betacoronavirus HKU24 | Betacoronavirus; Embecovirus | 1358 |
| YP_005454245.1 | Rabbit coronavirus HKU14 | Betacoronavirus; unclassified | 1362 |
| YP_009555241.1 | Human coronavirus OC43 | Betacoronavirus; Embecovirus | 1353 |
| NP_150077.1 | Bovine coronavirus (strain 98TXSF-110-ENT | Betacoronavirus; Embecovirus | 1363 |
| NP_073551.1 | Human coronavirus 229E | Alphacoronavirus; Duvinacovirus | 1173 |

**Table 1.3:** Envelope Proteins

| NCBI ID | Types | Genera; Sub-Genera | Length |
| --- | --- | --- | --- |
| YP_009724392.1 | SARS-CoV-2 | BetaCoronavirus;Sarbecovirus | 75 |
| QHR63282.1 | SARS-CoV-2 | BetaCoronavirus;Sarbecovirus | 75 |
| QHR63292.1 | SARS-CoV-2 | BetaCoronavirus;Sarbecovirus | 75 |
| QHR63262.1 | SARS-CoV-2 | BetaCoronavirus;Sarbecovirus | 83 |
| QHR63252.1 | SARS-CoV-2 | BetaCoronavirus;Sarbecovirus | 75 |
| QHO62113.1 | SARS-CoV-2 | BetaCoronavirus;Sarbecovirus | 75 |
| QHQ71975.1 | SARS-CoV-2 | BetaCoronavirus;Sarbecovirus | 75 |
| QHQ71965.1 | SARS-CoV-2 | BetaCoronavirus;Sarbecovirus | 75 |
| QHO62879.1 | SARS-CoV-2 | BetaCoronavirus;Sarbecovirus | 75 |
| QHO60596.1 | SARS-CoV-2 | BetaCoronavirus;Sarbecovirus | 75 |
| QHN73812.1 | SARS-CoV-2 | BetaCoronavirus;Sarbecovirus | 75 |
| QHN73797.1 | SARS-CoV-2 | BetaCoronavirus;Sarbecovirus | 75 |
| QHR84451.1 | SARS-CoV-2 | BetaCoronavirus;Sarbecovirus | 75 |
| QIC53215.1 | SARS-CoV-2 | BetaCoronavirus;Sarbecovirus | 75 |
| QHQ82466.1 | SARS-CoV-2 | BetaCoronavirus;Sarbecovirus | 75 |
| QIC53206.1 | SARS-CoV-2 | BetaCoronavirus;Sarbecovirus | 75 |
| QHD43418.1 | SARS-CoV-2 | BetaCoronavirus;Sarbecovirus | 75 |
| YP_001941169.1 | Turkey coronavirus | GammaCoronavirus;Igacovirus | 99 |
| YP_003029850.1 | Rat coronavirus Parker | BetaCoronavirus;Embecovirus | 88 |
| AAU06359.1 | Murine hepatitis virus | BetaCoronavirus;Embecovirus | 83 |
| YP_009113028.1 | Betacoronavirus HKU24 | BetaCoronavirus;Embecovirus | 82 |
| YP_005454247.1 | Rabbit coronavirus HKU14 | BetaCoronavirus;unclassified | 89 |
| YP_009555243.1 | HCoV-OC43 | BetaCoronavirus;Embecovirus | 84 |
| YP_009047209.1 | MERS-CoV | BetaCoronavirus;Merbecovirus | 82 |
| YP_007188584.1 | Betacoronavirus England 1 | BetaCoronavirus;Merbecovirus | 82 |
| YP_009273007.1 | Rousettus bat coronavirus | BetaCoronavirus;Nobecovirus | 76 |
| NP_828854.1 | SARS-CoV-1 | BetaCoronavirus;Sarbecovirus | 76 |
| YP_009072442.1 | Bat Hp-betacoronavirus | BetaCoronavirus;Hibecovirus | 79 |
| YP_003769.1 | Human coronavirus NL63 | AlphaCoronavirus;Setracovirus | 77 |
| NP_073554.1 | Human coronavirus 229E | AlphaCoronavirus;Duvinacovirus | 77 |
| YP_009724392.1 | SARS-CoV-2 | BetaCoronavirus;Sarbecovirus | 75 |

**Table 1.4:** Membrane Proteins

| NCBI ID | Type | Genera, Sub-Genera | Length |
| --- | --- | --- | --- |
| NP_073555.1 | *HCoV-229E* | Alphacoronavirus; Duvinacovirus | 225 |
| YP_003770.1 | *HCoV-NL63* | Alphacoronavirus; Setracovirus | 226 |
| YP_009113029 | *HKU24* | Betacoronavirus; Embecovirus | 231 |
| YP_009555244 | HCoV-OC43 | Betacoronavirus; Embecovirus | 230 |
| YP_005454248 | *Rabbit coronavirus HKU14* | Betacoronavirus; unclassified | 230 |
| YP_173241.1 | *HCoV-HKU1* | Betacoronavirus; Embecovirus | 223 |
| AAU06360.1 | *Murine hepatitis virus* | Betacoronavirus; Embecovirus | 228 |
| YP_003029851 | *Rat coronavirus Parker* | Betacoronavirus; Embecovirus | 228 |
| YP_001941170 | *Turkey enteric coronavirus* | Gammacoronavirus; Igacovirus | 223 |
| YP_001039968 | *BtCoV/HKU5/2004* | Betacoronavirus; Merbecovirus | 220 |
| YP_001039959 | *BtCoV/HKU4/2004* | Betacoronavirus; Merbecovirus | 219 |
| YP_009047210 | *MERS* | Betacoronavirus; Merbecovirus | 219 |
| YP_007188585 | Betacoronavirus England 1 | Betacoronavirus; Merbecovirus | 219 |
| YP_001039958 | *BtCoV/HKU5/2004* | Betacoronavirus; Merbecovirus | 82 |
| YP_001039967 | *BtCoV/HKU5/2004* | Betacoronavirus; Merbecovirus | 82 |
| YP_001039973 | *BtCoV/HKU9* | Betacoronavirus; Nobecovirus | 79 |
| NP_150081.1 | Bovine coronavirus | Betacoronavirus; Embecovirus | 84 |
| YP_173240.1 | *HCoV-HKU1* | Betacoronavirus; Embecovirus | 82 |
| YP_001039974 | *BtCoV/HKU9* | Betacoronavirus; Nobecovirus | 222 |
| YP_009273008 | *Rousettus bat coronavirus* | Betacoronavirus; Nobecovirus | 221 |
| YP_009072443 | *Bat Hp-betacoronavirus/Zhejiang2013* | Betacoronavirus; Hibecovirus | 223 |
| YP_009724393 | COVID-19 | Betacoronavirus; Sarbecovirus | 222 |
| QHQ82467.1_m | COVID-19 | Betacoronavirus; Sarbecovirus | 222 |
| QHD43419.1_m | COVID-19 | Betacoronavirus; Sarbecovirus | 222 |
| QHR63263.1_m | COVID-19 | Betacoronavirus; Sarbecovirus | 222 |
| QHQ71976.1_m | COVID-19 | Betacoronavirus; Sarbecovirus | 222 |
| QHR63283.1_m | COVID-19 | Betacoronavirus; Sarbecovirus | 222 |
| QIC53216.1_m | COVID-19 | Betacoronavirus; Sarbecovirus | 222 |
| QHR63253.1_m | COVID-19 | Betacoronavirus; Sarbecovirus | 222 |
| QHN73798.1_m | COVID-19 | Betacoronavirus; Sarbecovirus | 222 |
| QHO60597.1_m | COVID-19 | Betacoronavirus; Sarbecovirus | 222 |
| QHR63293.1_m | COVID-19 | Betacoronavirus; Sarbecovirus | 222 |
| QHR84452.1_m | COVID-19 | Betacoronavirus; Sarbecovirus | 222 |
| QIC53207.1_m | COVID-19 | Betacoronavirus; Sarbecovirus | 222 |
| QHQ71966.1_m | COVID-19 | Betacoronavirus; Sarbecovirus | 222 |
| QHO62880.1_m | COVID-19 | Betacoronavirus; Sarbecovirus | 222 |
| QHN73813.1_m | COVID-19 | Betacoronavirus; Sarbecovirus | 222 |
| NP_828855.1 | SARS_CoV1 | Betacoronavirus; Sarbecovirus | 221 |

**Table 1.5:** Nucleocapsid Proteins

| NCBI ID | Type | Genera, Sub-Genera | Length |
| --- | --- | --- | --- |
| YP 173243.1 | HCoV-HKU1 | Betacoronavirus; Embecovirus | 205 |
| YP 009113031 | HCoV-HKU24 | Betacoronavirus; Embecovirus | 443 |
| YP 005454249 | Rabbit-HKU14 | Betacoronavirus; unclassified | 444 |
| YP 009555245 | HCoV-OC43 | Betacoronavirus; Embecovirus | 448 |
| NP 150083.1 | Bov-ENT | Betacoronavirus; Embecovirus | 448 |
| YP_173242.1 | HCoV-HKU1 | Betacoronavirus; Embecovirus | 441 |
| YP_003029852 | Rat-CoV | Betacoronavirus; Embecovirus | 454 |
| NP_045302.1 | Murin-Cov | Betacoronavirus; Embecovirus | 454 |
| AAU06361.1 | Murin-Cov | Betacoronavirus; Embecovirus | 454 |
| YP_001941174 | Tur_CoV | Gammacoronavirus; Igacovirus | 409 |
| NP_073556.1 | HCoV-229E | Alphacoronavirus; Duvinacovirus | 389 |
| YP_003771.1 | HCoV-NL63 | Alphacoronavirus; Setracovirus | 377 |
| YP_001039975 | Bov-CoV_HKU9 | Betacoronavirus; Nobecovirus | 468 |
| YP_009273009 | Bov-CoV | Betacoronavirus; Nobecovirus | 443 |
| YP_009072446 | Bov-HpCoV | Betacoronavirus; Hibecovirus | 418 |
| YP_001039960 | Bov-CoV_HKU4 | Betacoronavirus; Merbecovirus | 423 |
| YP_009047211 | MERS | Betacoronavirus; Merbecovirus | 413 |
| YP_007188586 | SARS-CoV!! | Betacoronavirus; Merbecovirus | 411 |
| NP 828858.1 | *SARS-CoV* | Betacoronavirus; Sarbecovirus | 422 |
| YP_009724397 | COVID-19 | Betacoronavirus; Sarbecovirus | 419 |
| QHR63258.1 | COVID-19 | Betacoronavirus; Sarbecovirus | 419 |
| QIC53221.1 | COVID-19 | Betacoronavirus; Sarbecovirus | 419 |
| QHO60601.1 | COVID-19 | Betacoronavirus; Sarbecovirus | 419 |
| QHN73817.1 | COVID-19 | Betacoronavirus; Sarbecovirus | 419 |
| QHQ71980.1 | COVID-19 | Betacoronavirus; Sarbecovirus | 419 |
| QHQ71970.1 | COVID-19 | Betacoronavirus; Sarbecovirus | 419 |
| QHO62884.1 | COVID-19 | Betacoronavirus; Sarbecovirus | 419 |
| QHR84456.1 | COVID-19 | Betacoronavirus; Sarbecovirus | 419 |
| QHQ82471.1 | COVID-19 | Betacoronavirus; Sarbecovirus | 419 |
| QHO62115.1 | COVID-19 | Betacoronavirus; Sarbecovirus | 419 |
| QHR63288.1 | COVID-19 | Betacoronavirus; Sarbecovirus | 419 |
| QHR63268.1 | COVID-19 | Betacoronavirus; Sarbecovirus | 419 |
| QIC53211.1 | COVID-19 | Betacoronavirus; Sarbecovirus | 419 |
| QHR63298.1 | COVID-19 | Betacoronavirus; Sarbecovirus | 419 |
| QHN73802.1 | COVID-19 | Betacoronavirus; Sarbecovirus | 419 |
