## Supplementary File 2 for "Genome based Evolutionary study of SARS-CoV-2 towards the Prediction of Epitope Based Chimeric Vaccine"

**Supplementary File 2:** Retrieved protein sequences of major structural proteins of COVID-19.

**31 protein sequences of surface glycoprotein**

>QHW06059.1 surface glycoprotein [Wuhan seafood market pneumonia virus]

MFVFLVLLPLVSSQCVNLTTRTQLPPAYTNSFTRGVYYPDKVFRSSVLYSTQDLFLPFFSNVTWFHAIHV

SGTNGTKRFDNPVLPFNDGVYFASTEKSNIIRGWIFGTTLDSKTQSLLIVNNATNVVIKVCEFQFCNDPF

LGVYYHKNNKSWMESEFRVYSSANNCTFEYVSQPFLMDLEGKQGNFKNLREFVFKNIDGYFKIYSKHTPI

NLVRDLPQGFSALEPLVDLPIGINITRFQTLLALHRSYLTPGDSSSGWTAGAAAYYVGYLQPRTFLLKYN

ENGTITDAVDCALDPLSETKCTLKSFTVEKGIYQTSNFRVQPTESIVRFPNITNLCPFGEVFNATRFASV

YAWNRKRISNCVADYSVLYNSASFSTFKCYGVSPTKLNDLCFTNVYADSFVIRGDEVRQIAPGQTGKIAD

YNYKLPDDFTGCVIAWNSNNLDSKVGGNYNYLYRLFRKSNLKPFERDISTEIYQAGSTPCNGVEGFNCYF

PLQSYGFQPTNGVGYQPYRVVVLSFELLHAPATVCGPKKSTNLVKNKCVNFNFNGLTGTGVLTESNKKFL

PFQQFGRDIADTTDAVRDPQTLEILDITPCSFGGVSVITPGTNTSNQVAVLYQDVNCTEVPVAIHADQLT

PTWRVYSTGSNVFQTRAGCLIGAEHVNNSYECDIPIGAGICASYQTQTNSPRRARSVASQSIIAYTMSLG

AENSVAYSNNSIAIPTNFTISVTTEILPVSMTKTSVDCTMYICGDSTECSNLLLQYGSFCTQLNRALTGI

AVEQDKNTQEVFAQVKQIYKTPPIKDFGGFNFSQILPDPSKPSKRSFIEDLLFNKVTLADAGFIKQYGDC

LGDIAARDLICAQKFNGLTVLPPLLTDEMIAQYTSALLAGTITSGWTFGAGAALQIPFAMQMAYRFNGIG

VTQNVLYENQKLIANQFNSAIGKIQDSLSSTASALGKLQDVVNQNAQALNTLVKQLSSNFGAISSVLNDI

LSRLDKVEAEVQIDRLITGRLQSLQTYVTQQLIRAAEIRASANLAATKMSECVLGQSKRVDFCGKGYHLM

SFPQSAPHGVVFLHVTYVPAQEKNFTTAPAICHDGKAHFPREGVFVSNGTHWFVTQRNFYEPQIITTDNT

FVSGNCDVVIGIVNNTVYDPLQPELDSFKEELDKYFKNHTSPDVDLGDISGINASVVNIQKEIDRLNEVA

KNLNESLIDLQELGKYEQYIKWPWYIWLGFIAGLIAIVMVTIMLCCMTSCCSCLKGCCSCGSCCKFDEDD

SEPVLKGVKLHYT

>QHW06049.1 surface glycoprotein [Wuhan seafood market pneumonia virus]

MFVFLVLLPLVSSQCVNLTTRTQLPPAYTNSFTRGVYYPDKVFRSSVLHSTQDLFLPFFSNVTWFHAIHV

SGTNGTKRFDNPVLPFNDGVYFASTEKSNIIRGWIFGTTLDSKTQSLLIVNNATNVVIKVCEFQFCNDPF

LGVYYHKNNKSWMESEFRVYSSANNCTFEYVSQPFLMDLEGKQGNFKNLREFVFKNIDGYFKIYSKHTPI

NLVRDLPQGFSALEPLVDLPIGINITRFQTLLALHRSYLTPGDSSSGWTAGAAAYYVGYLQPRTFLLKYN

ENGTITDAVDCALDPLSETKCTLKSFTVEKGIYQTSNFRVQPTESIVRFPNITNLCPFGEVFNATRFASV

YAWNRKRISNCVADYSVLYNSASFSTFKCYGVSPTKLNDLCFTNVYADSFVIRGDEVRQIAPGQTGKIAD

YNYKLPDDFTGCVIAWNSNNLDSKVGGNYNYLYRLFRKSNLKPFERDISTEIYQAGSTPCNGVEGFNCYF

PLQSYGFQPTNGVGYQPYRVVVLSFELLHAPATVCGPKKSTNLVKNKCVNFNFNGLTGTGVLTESNKKFL

PFQQFGRDIADTTDAVRDPQTLEILDITPCSFGGVSVITPGTNTSNQVAVLYQDVNCTEVPVAIHADQLT

PTWRVYSTGSNVFQTRAGCLIGAEHVNNSYECDIPIGAGICASYQTQTNSPRRARSVASQSIIAYTMSLG

AENSVAYSNNSIAIPTNFTISVTTEILPVSMTKTSVDCTMYICGDSTECSNLLLQYGSFCTQLNRALTGI

AVEQDKNTQEVFAQVKQIYKTPPIKDFGGFNFSQILPDPSKPSKRSFIEDLLFNKVTLADAGFIKQYGDC

LGDIAARDLICAQKFNGLTVLPPLLTDEMIAQYTSALLAGTITSGWTFGAGAALQIPFAMQMAYRFNGIG

VTQNVLYENQKLIANQFNSAIGKIQDSLSSTASALGKLQDVVNQNAQALNTLVKQLSSNFGAISSVLNDI

LSRLDKVEAEVQIDRLITGRLQSLQTYVTQQLIRAAEIRASANLAATKMSECVLGQSKRVDFCGKGYHLM

SFPQSAPHGVVFLHVTYVPAQEKNFTTAPAICHDGKAHFPREGVFVSNGTHWFVTQRNFYEPQIITTDNT

FVSGNCDVVIGIVNNTVYDPLQPELDSFKEELDKYFKNHTSPDVDLGDISGINASVVNIQKEIDRLNEVA

KNLNESLIDLQELGKYEQYIKWPWYIWLGFIAGLIAIVMVTIMLCCMTSCCSCLKGCCSCGSCCKFDEDD

SEPVLKGVKLHYT

>QHW06039.1 surface glycoprotein [Wuhan seafood market pneumonia virus]

MFVFLVLLPLVSSQCVNLTTRTQLPPAYTNSFTRGVYYPDKVFRSSVLHSTQDLFLPFFSNVTWFHAIHV

SGTNGTKRFDNPVLPFNDGVYFASTEKSNIIRGWIFGTTLDSKTQSLLIVNNATNVVIKVCEFQFCNDPF

LGVYYHKNNKSWMESEFRVYSSANNCTFEYVSQPFLMDLEGKQGNFKNLREFVFKNIDGYFKIYSKHTPI

NLVRDLPQGFSALEPLVDLPIGINITRFQTLLALHRSYLTPGDSSSGWTAGAAAYYVGYLQPRTFLLKYN

ENGTITDAVDCALDPLSETKCTLKSFTVEKGIYQTSNFRVQPTESIVRFPNITNLCPFGEVFNATRFASV

YAWNRKRISNCVADYSVLYNSASFSTFKCYGVSPTKLNDLCFTNVYADSFVIRGDEVRQIAPGQTGKIAD

YNYKLPDDFTGCVIAWNSNNLDSKVGGNYNYLYRLFRKSNLKPFERDISTEIYQAGSTPCNGVEGFNCYF

PLQSYGFQPTNGVGYQPYRVVVLSFELLHAPATVCGPKKSTNLVKNKCVNFNFNGLTGTGVLTESNKKFL

PFQQFGRDIADTTDAVRDPQTLEILDITPCSFGGVSVITPGTNTSNQVAVLYQDVNCTEVPVAIHADQLT

PTWRVYSTGSNVFQTRAGCLIGAEHVNNSYECDIPIGAGICASYQTQTNSPRRARSVASQSIIAYTMSLG

AENSVAYSNNSIAIPTNFTISVTTEILPVSMTKTSVDCTMYICGDSTECSNLLLQYGSFCTQLNRALTGI

AVEQDKNTQEVFAQVKQIYKTPPIKDFGGFNFSQILPDPSKPSKRSFIEDLLFNKVTLADAGFIKQYGDC

LGDIAARDLICAQKFNGLTVLPPLLTDEMIAQYTSALLAGTITSGWTFGAGAALQIPFAMQMAYRFNGIG

VTQNVLYENQKLIANQFNSAIGKIQDSLSSTASALGKLQDVVNQNAQALNTLVKQLSSNFGAISSVLNDI

LSRLDKVEAEVQIDRLITGRLQSLQTYVTQQLIRAAEIRASANLAATKMSECVLGQSKRVDFCGKGYHLM

SFPQSAPHGVVFLHVTYVPAQEKNFTTAPAICHDGKAHFPREGVFVSNGTHWFVTQRNFYEPQIITTDNT

FVSGNCDVVIGIVNNTVYDPLQPELDSFKEELDKYFKNHTSPDVDLGDISGINASVVNIQKEIDRLNEVA

KNLNESLIDLQELGKYEQYIKWPWYIWLGFIAGLIAIVMVTIMLCCMTSCCSCLKGCCSCGSCCKFDEDD

SEPVLKGVKLHYT

>BBW89517.1 surface glycoprotein [Wuhan seafood market pneumonia virus]

MFVFLVLLPLVSSQCVNLTTRTQLPPAYTNSFTRGVYYPDKVFRSSVLHSTQDLFLPFFSNVTWFHAIHV

SGTNGTKRFDNPVLPFNDGVYFASTEKSNIIRGWIFGTTLDSKTQSLLIVNNATNVVIKVCEFQFCNDPF

LGVYYHKNNKSWMESEFRVYSSANNCTFEYVSQPFLMDLEGKQGNFKNLREFVFKNIDGYFKIYSKHTPI

NLVRDLPQGFSALEPLVDLPIGINITRFQTLLALHRSYLTPGDSSSGWTAGAAAYYVGYLQPRTFLLKYN

ENGTITDAVDCALDPLSETKCTLKSFTVEKGIYQTSNFRVQPTESIVRFPNITNLCPFGEVFNATRFASV

YAWNRKRISNCVADYSVLYNSASFSTFKCYGVSPTKLNDLCFTNVYADSFVIRGDEVRQIAPGQTGKIAD

YNYKLPDDFTGCVIAWNSNNLDSKVGGNYNYLYRLFRKSNLKPFERDISTEIYQAGSTPCNGVEGFNCYF

PLQSYGFQPTNGVGYQPYRVVVLSFELLHAPATVCGPKKSTNLVKNKCVNFNFNGLTGTGVLTESNKKFL

PFQQFGRDIADTTDAVRDPQTLEILDITPCSFGGVSVITPGTNTSNQVAVLYQDVNCTEVPVAIHADQLT

PTWRVYSTGSNVFQTRAGCLIGAEHVNNSYECDIPIGAGICASYQTQTNSPRRARSVASQSIIAYTMSLG

AENSVAYSNNSIAIPTNFTISVTTEILPVSMTKTSVDCTMYICGDSTECSNLLLQYGSFCTQLNRALTGI

AVEQDKNTQEVFAQVKQIYKTPPIKDFGGFNFSQILPDPSKPSKRSFIEDLLFNKVTLADAGFIKQYGDC

LGDIAARDLICAQKFNGLTVLPPLLTDEMIAQYTSALLAGTITSGWTFGAGAALQIPFAMQMAYRFNGIG

VTQNVLYENQKLIANQFNSAIGKIQDSLSSTASALGKLQDVVNQNAQALNTLVKQLSSNFGAISSVLNDI

LSRLDKVEAEVQIDRLITGRLQSLQTYVTQQLIRAAEIRASANLAATKMSECVLGQSKRVDFCGKGYHLM

SFPQSAPHGVVFLHVTYVPAQEKNFTTAPAICHDGKAHFPREGVFVSNGTHWFVTQRNFYEPQIITTDNT

FVSGNCDVVIGIVNNTVYDPLQPELDSFKEELDKYFKNHTSPDVDLGDISGINASVVNIQKEIDRLNEVA

KNLNESLIDLQELGKYEQYIKWPWYIWLGFIAGLIAIVMVTIMLCCMTSCCSCLKGCCSCGSCCKFDEDD

SEPVLKGVKLHYT

>QHU79204.1 surface glycoprotein [Wuhan seafood market pneumonia virus]

MFVFLVLLPLVSSQCVNLTTRTQLPPAYTNSFTRGVYYPDKVFRSSVLHSTQDLFLPFFSNVTWFHAIHV

SGTNGTKRFDNPVLPFNDGVYFASTEKSNIIRGWIFGTTLDSKTQSLLIVNNATNVVIKVCEFQFCNDPF

LGVYYHKNNKSWMESEFRVYSSANNCTFEYVSQPFLMDLEGKQGNFKNLREFVFKNIDGYFKIYSKHTPI

NLVRDLPQGFSALEPLVDLPIGINITRFQTLLALHRSYLTPGDSSSGWTAGAAAYYVGYLQPRTFLLKYN

ENGTITDAVDCALDPLSETKCTLKSFTVEKGIYQTSNFRVQPTESIVRFPNITNLCPFGEVFNATRFASV

YAWNRKRISNCVADYSVLYNSASFSTFKCYGVSPTKLNDLCFTNVYADSFVIRGDEVRQIAPGQTGKIAD

YNYKLPDDFTGCVIAWNSNNLDSKVGGNYNYLYRLFRKSNLKPFERDISTEIYQAGSTPCNGVEGFNCYF

PLQSYGFQPTNGVGYQPYRVVVLSFELLHAPATVCGPKKSTNLVKNKCVNFNFNGLTGTGVLTESNKKFL

PFQQFGRDIADTTDAVRDPQTLEILDITPCSFGGVSVITPGTNTSNQVAVLYQDVNCTEVPVAIHADQLT

PTWRVYSTGSNVFQTRAGCLIGAEHVNNSYECDIPIGAGICASYQTQTNSPRRARSVASQSIIAYTMSLG

AENSVAYSNNSIAIPTNFTISVTTEILPVSMTKTSVDCTMYICGDSTECSNLLLQYGSFCTQLNRALTGI

AVEQDKNTQEVFAQVKQIYKTPPIKDFGGFNFSQILPDPSKPSKRSFIEDLLFNKVTLADAGFIKQYGDC

LGDIAARDLICAQKFNGLTVLPPLLTDEMIAQYTSALLAGTITSGWTFGAGAALQIPFAMQMAYRFNGIG

VTQNVLYENQKLIANQFNSAIGKIQDSLSSTASALGKLQDVVNQNAQALNTLVKQLSSNFGAISSVLNDI

LSRLDKVEAEVQIDRLITGRLQSLQTYVTQQLIRAAEIRASANLAATKMSECVLGQSKRVDFCGKGYHLM

SFPQSAPHGVVFLHVTYVPAQEKNFTTAPAICHDGKAHFPREGVFVSNGTHWFVTQRNFYEPQIITTDNT

FVSGNCDVVIGIVNNTVYDPLQPELDSFKEELDKYFKNHTSPDVDLGDISGINASVVNIQKEIDRLNEVA

KNLNESLIDLQELGKYEQYIKWPWYIWLGFIAGLIAIVMVTIMLCCMTSCCSCLKGCCSCGSCCKFDEDD

SEPVLKGVKLHYT

>QHU79194.1 surface glycoprotein [Wuhan seafood market pneumonia virus]

MFVFLVLLPLVSSQCVNLTTRTQLPPAYTNSFTRGVYYPDKVFRSSVLHSTQDLFLPFFSNVTWFHAIHV

SGTNGTKRFDNPVLPFNDGVYFASTEKSNIIRGWIFGTTLDSKTQSLLIVNNATNVVIKVCEFQFCNDPF

LGVYYHKNNKSWMESEFRVYSSANNCTFEYVSQPFLMDLEGKQGNFKNLREFVFKNIDGYFKIYSKHTPI

NLVRDLPQGFSALEPLVDLPIGINITRFQTLLALHRSYLTPGDSSSGWTAGAAAYYVGYLQPRTFLLKYN

ENGTITDAVDCALDPLSETKCTLKSFTVEKGIYQTSNFRVQPTESIVRFPNITNLCPFGEVFNATRFASV

YAWNRKRISNCVADYSVLYNSASFSTFKCYGVSPTKLNDLCFTNVYADSFVIRGDEVRQIAPGQTGKIAD

YNYKLPDDFTGCVIAWNSNNLDSKVGGNYNYLYRLFRKSNLKPFERDISTEIYQAGSTPCNGVEGFNCYF

PLQSYGFQPTNGVGYQPYRVVVLSFELLHAPATVCGPKKSTNLVKNKCVNFNFNGLTGTGVLTESNKKFL

PFQQFGRDIADTTDAVRDPQTLEILDITPCSFGGVSVITPGTNTSNQVAVLYQDVNCTEVPVAIHADQLT

PTWRVYSTGSNVFQTRAGCLIGAEHVNNSYECDIPIGAGICASYQTQTNSPRRARSVASQSIIAYTMSLG

AENSVAYSNNSIAIPTNFTISVTTEILPVSMTKTSVDCTMYICGDSTECSNLLLQYGSFCTQLNRALTGI

AVEQDKNTQEVFAQVKQIYKTPPIKDFGGFNFSQILPDPSKPSKRSFIEDLLFNKVTLADAGFIKQYGDC

LGDIAARDLICAQKFNGLTVLPPLLTDEMIAQYTSALLAGTITSGWTFGAGAALQIPFAMQMAYRFNGIG

VTQNVLYENQKLIANQFNSAIGKIQDSLSSTASALGKLQDVVNQNAQALNTLVKQLSSNFGAISSVLNDI

LSRLDKVEAEVQIDRLITGRLQSLQTYVTQQLIRAAEIRASANLAATKMSECVLGQSKRVDFCGKGYHLM

SFPQSAPHGVVFLHVTYVPAQEKNFTTAPAICHDGKAHFPREGVFVSNGTHWFVTQRNFYEPQIITTDNT

FVSGNCDVVIGIVNNTVYDPLQPELDSFKEELDKYFKNHTSPDVDLGDISGINASVVNIQKEIDRLNEVA

KNLNESLIDLQELGKYEQYIKWPWYIWLGFIAGLIAIVMVTIMLCCMTSCCSCLKGCCSCGSCCKFDEDD

SEPVLKGVKLHY

>QHU79173.1 surface glycoprotein [Wuhan seafood market pneumonia virus]

MFVFLVLLPLVSSQCVNLTTRXXXXXXXXXXXXXXXXXXXXXXXXXXXXSTQDLFLPFFSNVTWFHAIHV

SGTNGTKRFDNPVLPFNDGVYFASTEKSNIIRGWIFGTTLDSKTQSLLIVNNATNVVIKVCEFQFCNDPF

LGVYYHKNNKSWMESEFRVYSSANNCTFEYVSQPFLMDLEGKQGNFKNLREFVFKNIDGYFKIYSKHTPI

NLVRDLPQGFSALEPLVDLPIGINITRFQTLLALHRSYLTPGDSSSGWTAGAAAYYVGYLQPRTFLLKYN

ENGTITDAVDCALDPLSETKCTLKSFTVEKGIYQTSNFRVQPTESIVRFPNITNLCPFGEVFNATRFASV

YAWNRKRISNCVADYSVLYNSASFSTFKCYGVSPTKLNDLCFTNVYADSFVIRGDEVRQIAPGQTGKIAD

YNYKLPDDFTGCVIAWNSNNLDSKVGGNYNYLYRLFRKSNLKPFERDISTEIYQAGSTPCNGVEGFNCYF

PLQSYGFQPTNGVGYQPYRVVVLSFELLHAPATVCGPKKSTNLVKNKCVNFNFNGLTGTGVLTESNKKFL

PFQQFGRDIADTTDAVRDPQTLEILDITPCSFGGVSVITPGTNTSNQVAVLYQDVNCTEVPVAIHADQLT

PTWRVYSTGSNVFQTRAGCLIGAEHVNNSYECDIPIGAGICASYQTQTNSPRRARSVASQSIIAYTMSLG

AENSVAYSNNSIAIPTNFTISVTTEILPVSMTKTSVDCTMYICGDSTECSNLLLQYGSFCTQLNRALTGI

AVEQDKNTQEVFAQVKQIYKTPPIKDFGGFNFSQILPDPSKPSKRSFIEDLLFNKVTLADAGFIKQYGDC

LGDIAARDLICAQKFNGLTVLPPLLTDEMIAQYTSALLAGTITSGWTFGAGAALQIPFAMQMAYRFNGIG

VTQNVLYENQKLIANQFNSAIGKIQDSLSSTASALGKLQDVVNQNAQALNTLVKQLSSNFGAISSVLNDI

LSRLDKVEAEVQIDRLITGRLQSLQTYVTQQLIRAAEIRASANLAATKMSECVLGQSKRVDFCGKGYHLM

SFPQSAPHGVVFLHVTYVPAQEKNFTTAPAICHDGKAHFPREGVFVSNGTHWFVTQRNFYEPQIITTDNT

FVSGNCDVVIGIVNNTVYDPLQPELDSFKEELDKYFKNHTSPDVDLGDISGINASVVNIQKEIDRLNEVA

KNLNESLIDLQELGKYEQYIKWPWYIWLGFIAGLIAIVMVTIMLCCMTSCCSCLKGCCSCGSCCKFDEDD

SEPVLKGVKLHYT

>QHU36864.1 surface glycoprotein [Wuhan seafood market pneumonia virus]

MFVFLVLLPLVSSQCVNLTTRTQLPPAYTNSFTRGVYYPDKVFRSSVLHSTQDLFLPFFSNVTWFHAIHV

SGTNGTKRFDNPVLPFNDGVYFASTEKSNIIRGWIFGTTLDSKTQSLLIVNNATNVVIKVCEFQFCNDPF

LGVYYHKNNKSWMESEFRVYSSANNCTFEYVSQPFLMDLEGKQGNFKNLREFVFKNIDGYFKIYSKHTPI

NLVRDLPQGFSALEPLVDLPIGINITRFQTLLALHRSYLTPGDSSSGWTAGAAAYYVGYLQPRTFLLKYN

ENGTITDAVDCALDPLSETKCTLKSFTVEKGIYQTSNFRVQPTESIVRFPNITNLCPFGEVFNATRFASV

YAWNRKRISNCVADYSVLYNSASFSTFKCYGVSPTKLNDLCFTNVYADSFVIRGDEVRQIAPGQTGKIAD

YNYKLPDDFTGCVIAWNSNNLDSKVGGNYNYLYRLFRKSNLKPFERDISTEIYQAGSTPCNGVEGFNCYF

PLQSYGFQPTNGVGYQPYRVVVLSFELLHAPATVCGPKKSTNLVKNKCVNFNFNGLTGTGVLTESNKKFL

PFQQFGRDIADTTDAVRDPQTLEILDITPCSFGGVSVITPGTNTSNQVAVLYQDVNCTEVPVAIHADQLT

PTWRVYSTGSNVFQTRAGCLIGAEHVNNSYECDIPIGAGICASYQTQTNSPRRARSVASQSIIAYTMSLG

AENSVAYSNNSIAIPTNFTISVTTEILPVSMTKTSVDCTMYICGDSTECSNLLLQYGSFCTQLNRALTGI

AVEQDKNTQEVFAQVKQIYKTPPIKDFGGFNFSQILPDPSKPSKRSFIEDLLFNKVTLADAGFIKQYGDC

LGDIAARDLICAQKFNGLTVLPPLLTDEMIAQYTSALLAGTITSGWTFGAGAALQIPFAMQMAYRFNGIG

VTQNVLYENQKLIANQFNSAIGKIQDSLSSTASALGKLQDVVNQNAQALNTLVKQLSSNFGAISSVLNDI

LSRLDKVEAEVQIDRLITGRLQSLQTYVTQQLIRAAEIRASANLAATKMSECVLGQSKRVDFCGKGYHLM

SFPQSAPHGVVFLHVTYVPAQEKNFTTAPAICHDGKAHFPREGVFVSNGTHWFVTQRNFYEPQIITTDNT

FVSGNCDVVIGIVNNTVYDPLQPELDSFKEELDKYFKNHTSPDVDLGDISGINASVVNIQKEIDRLNEVA

KNLNESLIDLQELGKYEQYIKWPWYIWLGFIAGLIAIVMVTIMLCCMTSCCSCLKGCCSCGSCCKFDEDD

SEPVLKGVKLHYT

>QHU36854.1 surface glycoprotein [Wuhan seafood market pneumonia virus]

MFVFLVLLPLVSSQCVNLTTRTQLPPAYTNSFTRGVYYPDKVFRSSVLHSTQDLFLPFFSNVTWFHAIHV

SGTNGTKRFDNPVLPFNDGVYFASTEKSNIIRGWIFGTTLDSKTQSLLIVNNATNVVIKVCEFQFCNDPF

LGVYYHKNNKSWMESEFRVYSSANNCTFEYVSQPFLMDLEGKQGNFKNLREFVFKNIDGYFKIYSKHTPI

NLVRDLPQGFSALEPLVDLPIGINITRFQTLLALHRSYLTPGDSSSGWTAGAAAYYVGYLQPRTFLLKYN

ENGTITDAVDCALDPLSETKCTLKSFTVEKGIYQTSNFRVQPTESIVRFPNITNLCPFGEVFNATRFASV

YAWNRKRISNCVADYSVLYNSASFSTFKCYGVSPTKLNDLCFTNVYADSFVIRGDEVRQIAPGQTGKIAD

YNYKLPDDFTGCVIAWNSNNLDSKVGGNYNYLYRLFRKSNLKPFERDISTEIYQAGSTPCNGVEGFNCYF

PLQSYGFQPTNGVGYQPYRVVVLSFELLHAPATVCGPKKSTNLVKNKCVNFNFNGLTGTGVLTESNKKFL

PFQQFGRDIADTTDAVRDPQTLEILDITPCSFGGVSVITPGTNTSNQVAVLYQDVNCTEVPVAIHADQLT

PTWRVYSTGSNVFQTRAGCLIGAEHVNNSYECDIPIGAGICASYQTQTNSPRRARSVASQSIIAYTMSLG

AENSVAYSNNSIAIPTNFTISVTTEILPVSMTKTSVDCTMYICGDSTECSNLLLQYGSFCTQLNRALTGI

AVEQDKNTQEVFAQVKQIYKTPPIKDFGGFNFSQILPDPSKPSKRSFIEDLLFNKVTLADAGFIKQYGDC

LGDIAARDLICAQKFNGLTVLPPLLTDEMIAQYTSALLAGTITSGWTFGAGAALQIPFAMQMAYRFNGIG

VTQNVLYENQKLIANQFNSAIGKIQDSLSSTASALGKLQDVVNQNAQALNTLVKQLSSNFGAISSVLNDI

LSRLDKVEAEVQIDRLITGRLQSLQTYVTQQLIRAAEIRASANLAATKMSECVLGQSKRVDFCGKGYHLM

SFPQSAPHGVVFLHVTYVPAQEKNFTTAPAICHDGKAHFPREGVFVSNGTHWFVTQRNFYEPQIITTDNT

FVSGNCDVVIGIVNNTVYDPLQPELDSFKEELDKYFKNHTSPDVDLGDISGINASVVNIQKEIDRLNEVA

KNLNESLIDLQELGKYEQYIKWPWYIWLGFIAGLIAIVMVTIMLCCMTSCCSCLKGCCSCGSCCKFDEDD

SEPVLKGVKLHYT

>QHU36844.1 surface glycoprotein [Wuhan seafood market pneumonia virus]

MFVFLVLLPLVSSQCVNLTTRTQLPPAYTNSFTRGVYYPDKVFRSSVLHSTQDLFLPFFSNVTWFHAIHV

SGTNGTKRFDNPVLPFNDGVYFASTEKSNIIRGWIFGTTLDSKTQSLLIVNNATNVVIKVCEFQFCNDPF

LGVYYHKNNKSWMESEFRVYSSANNCTFEYVSQPFLMDLEGKQGNFKNLREFVFKNIDGYFKIYSKHTPI

NLVRDLPQGFSALEPLVDLPIGINITRFQTLLALHRSYLTPGDSSSGWTAGAAAYYVGYLQPRTFLLKYN

ENGTITDAVDCALDPLSETKCTLKSFTVEKGIYQTSNFRVQPTESIVRFPNITNLCPFGEVFNATRFASV

YAWNRKRISNCVADYSVLYNSASFSTFKCYGVSPTKLNDLCFTNVYADSFVIRGDEVRQIAPGQTGKIAD

YNYKLPDDFTGCVIAWNSNNLDSKVGGNYNYLYRLFRKSNLKPFERDISTEIYQAGSTPCNGVEGFNCYF

PLQSYGFQPTNGVGYQPYRVVVLSFELLHAPATVCGPKKSTNLVKNKCVNFNFNGLTGTGVLTESNKKFL

PFQQFGRDIADTTDAVRDPQTLEILDITPCSFGGVSVITPGTNTSNQVAVLYQDVNCTEVPVAIHADQLT

PTWRVYSTGSNVFQTRAGCLIGAEHVNNSYECDIPIGAGICASYQTQTNSPRRARSVASQSIIAYTMSLG

AENSVAYSNNSIAIPTNFTISVTTEILPVSMTKTSVDCTMYICGDSTECSNLLLQYGSFCTQLNRALTGI

AVEQDKNTQEVFAQVKQIYKTPPIKDFGGFNFSQILPDPSKPSKRSFIEDLLFNKVTLADAGFIKQYGDC

LGDIAARDLICAQKFNGLTVLPPLLTDEMIAQYTSALLAGTITSGWTFGAGAALQIPFAMQMAYRFNGIG

VTQNVLYENQKLIANQFNSAIGKIQDSLSSTASALGKLQDVVNQNAQALNTLVKQLSSNFGAISSVLNDI

LSRLDKVEAEVQIDRLITGRLQSLQTYVTQQLIRAAEIRASANLAATKMSECVLGQSKRVDFCGKGYHLM

SFPQSAPHGVVFLHVTYVPAQEKNFTTAPAICHDGKAHFPREGVFVSNGTHWFVTQRNFYEPQIITTDNT

FVSGNCDVVIGIVNNTVYDPLQPELDSFKEELDKYFKNHTSPDVDLGDISGINASVVNIQKEIDRLNEVA

KNLNESLIDLQELGKYEQYIKWPWYIWLGFIAGLIAIVMVTIMLCCMTSCCSCLKGCCSCGSCCKFDEDD

SEPVLKGVKLHYT

>QHU36834.1 surface glycoprotein [Wuhan seafood market pneumonia virus]

MFVFLVLLPLVSSQCVNLTTRTQLPPAYTNSFTRGVYYPDKVFRSSVLHSTQDLFLPFFSNVTWFHAIHV

SGTNGTKRFDNPVLPFNDGVYFASTEKSNIIRGWIFGTTLDSKTQSLLIVNNATNVVIKVCEFQFCNDPF

LGVYYHKNNKSWMESEFRVYSSANNCTFEYVSQPFLMDLEGKQGNFKNLREFVFKNIDGYFKIYSKHTPI

NLVRDLPQGFSALEPLVDLPIGINITRFQTLLALHRSYLTPGDSSSGWTAGAAAYYVGYLQPRTFLLKYN

ENGTITDAVDCALDPLSETKCTLKSFTVEKGIYQTSNFRVQPTESIVRFPNITNLCPFGEVFNATRFASV

YAWNRKRISNCVADYSVLYNSASFSTFKCYGVSPTKLNDLCFTNVYADSFVIRGDEVRQIAPGQTGKIAD

YNYKLPDDFTGCVIAWNSNNLDSKVGGNYNYLYRLFRKSNLKPFERDISTEIYQAGSTPCNGVEGFNCYF

PLQSYGFQPTNGVGYQPYRVVVLSFELLHAPATVCGPKKSTNLVKNKCVNFNFNGLTGTGVLTESNKKFL

PFQQFGRDIADTTDAVRDPQTLEILDITPCSFGGVSVITPGTNTSNQVAVLYQDVNCTEVPVAIHADQLT

PTWRVYSTGSNVFQTRAGCLIGAEHVNNSYECDIPIGAGICASYQTQTNSPRRARSVASQSIIAYTMSLG

AENSVAYSNNSIAIPTNFTISVTTEILPVSMTKTSVDCTMYICGDSTECSNLLLQYGSFCTQLNRALTGI

AVEQDKNTQEVFAQVKQIYKTPPIKDFGGFNFSQILPDPSKPSKRSFIEDLLFNKVTLADAGFIKQYGDC

LGDIAARDLICAQKFNGLTVLPPLLTDEMIAQYTSALLAGTITSGWTFGAGAALQIPFAMQMAYRFNGIG

VTQNVLYENQKLIANQFNSAIGKIQDSLSSTASALGKLQDVVNQNAQALNTLVKQLSSNFGAISSVLNDI

LSRLDKVEAEVQIDRLITGRLQSLQTYVTQQLIRAAEIRASANLAATKMSECVLGQSKRVDFCGKGYHLM

SFPQSAPHGVVFLHVTYVPAQEKNFTTAPAICHDGKAHFPREGVFVSNGTHWFVTQRNFYEPQIITTDNT

FVSGNCDVVIGIVNNTVYDPLQPELDSFKEELDKYFKNHTSPDVDLGDISGINASVVNIQKEIDRLNEVA

KNLNESLIDLQELGKYEQYIKWPWYIWLGFIAGLIAIVMVTIMLCCMTSCCSCLKGCCSCGSCCKFDEDD

SEPVLKGVKLHYT

>QHU36824.1 surface glycoprotein [Wuhan seafood market pneumonia virus]

MFVFLVLLPLVSSQCVNLTTRTQLPPAYTNSFTRGVYYPDKVFRSSVLHSTQDLFLPFFSNVTWFHAIHV

SGTNGTKRFDNPVLPFNDGVYFASTEKSNIIRGWIFGTTLDSKTQSLLIVNNATNVVIKVCEFQFCNDPF

LGVYYHKNNKSWMESEFRVYSSANNCTFEYVSQPFLMDLEGKQGNFKNLREFVFKNIDGYFKIYSKHTPI

NLVRDLPQGFSALEPLVDLPIGINITRFQTLLALHRSYLTPGDSSSGWTAGAAAYYVGYLQPRTFLLKYN

ENGTITDAVDCALDPLSETKCTLKSFTVEKGIYQTSNFRVQPTESIVRFPNITNLCPFGEVFNATRFASV

YAWNRKRISNCVADYSVLYNSASFSTFKCYGVSPTKLNDLCFTNVYADSFVIRGDEVRQIAPGQTGKIAD

YNYKLPDDFTGCVIAWNSNNLDSKVGGNYNYLYRLFRKSNLKPFERDISTEIYQAGSTPCNGVEGFNCYF

PLQSYGFQPTNGVGYQPYRVVVLSFELLHAPATVCGPKKSTNLVKNKCVNFNFNGLTGTGVLTESNKKFL

PFQQFGRDIADTTDAVRDPQTLEILDITPCSFGGVSVITPGTNTSNQVAVLYQDVNCTEVPVAIHADQLT

PTWRVYSTGSNVFQTRAGCLIGAEHVNNSYECDIPIGAGICASYQTQTNSPRRARSVASQSIIAYTMSLG

AENSVAYSNNSIAIPTNFTISVTTEILPVSMTKTSVDCTMYICGDSTECSNLLLQYGSFCTQLNRALTGI

AVEQDKNTQEVFAQVKQIYKTPPIKDFGGFNFSQILPDPSKPSKRSFIEDLLFNKVTLADAGFIKQYGDC

LGDIAARDLICAQKFNGLTVLPPLLTDEMIAQYTSALLAGTITSGWTFGAGAALQIPFAMQMAYRFNGIG

VTQNVLYENQKLIANQFNSAIGKIQDSLSSTASALGKLQDVVNQNAQALNTLVKQLSSNFGAISSVLNDI

LSRLDKVEAEVQIDRLITGRLQSLQTYVTQQLIRAAEIRASANLAATKMSECVLGQSKRVDFCGKGYHLM

SFPQSAPHGVVFLHVTYVPAQEKNFTTAPAICHDGKAHFPREGVFVSNGTHWFVTQRNFYEPQIITTDNT

FVSGNCDVVIGIVNNTVYDPLQPELDSFKEELDKYFKNHTSPDVDLGDISGINASVVNIQKEIDRLNEVA

KNLNESLIDLQELGKYEQYIKWPWYIWLGFIAGLIAIVMVTIMLCCMTSCCSCLKGCCSCGSCCKFDEDD

SEPVLKGVKLHYT

>QHR84449.1 surface glycoprotein [Wuhan seafood market pneumonia virus]

MFVFLVLLPLVSSQCVNLTTRTQLPPAYTNSFTRGVYYPDKVFRSSVLHSTQDLFLPFFSNVTWFHAIHV

SGTNGTKRFDNPVLPFNDGVYFASTEKSNIIRGWIFGTTLDSKTQSLLIVNNATNVVIKVCEFQFCNDPF

LGVYYHKNNKSWMESEFRVYSSANNCTFEYVSQPFLMDLEGKQGNFKNLREFVFKNIDGYFKIYSKHTPI

NLVRDLPQGFSALEPLVDLPIGINITRFQTLLALHRRYLTPGDSSSGWTAGAAAYYVGYLQPRTFLLKYN

ENGTITDAVDCALDPLSETKCTLKSFTVEKGIYQTSNFRVQPTESIVRFPNITNLCPFGEVFNATRFASV

YAWNRKRISNCVADYSVLYNSASFSTFKCYGVSPTKLNDLCFTNVYADSFVIRGDEVRQIAPGQTGKIAD

YNYKLPDDFTGCVIAWNSNNLDSKVGGNYNYLYRLFRKSNLKPFERDISTEIYQAGSTPCNGVEGFNCYF

PLQSYGFQPTNGVGYQPYRVVVLSFELLHAPATVCGPKKSTNLVKNKCVNFNFNGLTGTGVLTESNKKFL

PFQQFGRDIADTTDAVRDPQTLEILDITPCSFGGVSVITPGTNTSNQVAVLYQDVNCTEVPVAIHADQLT

PTWRVYSTGSNVFQTRAGCLIGAEHVNNSYECDIPIGAGICASYQTQTNSPRRARSVASQSIIAYTMSLG

AENSVAYSNNSIAIPTNFTISVTTEILPVSMTKTSVDCTMYICGDSTECSNLLLQYGSFCTQLNRALTGI

AVEQDKNTQEVFAQVKQIYKTPPIKDFGGFNFSQILPDPSKPSKRSFIEDLLFNKVTLADAGFIKQYGDC

LGDIAARDLICAQKFNGLTVLPPLLTDEMIAQYTSALLAGTITSGWTFGAGAALQIPFAMQMAYRFNGIG

VTQNVLYENQKLIANQFNSAIGKIQDSLSSTASALGKLQDVVNQNAQALNTLVKQLSSNFGAISSVLNDI

LSRLDKVEAEVQIDRLITGRLQSLQTYVTQQLIRAAEIRASANLAATKMSECVLGQSKRVDFCGKGYHLM

SFPQSAPHGVVFLHVTYVPAQEKNFTTAPAICHDGKAHFPREGVFVSNGTHWFVTQRNFYEPQIITTDNT

FVSGNCDVVIGIVNNTVYDPLQPELDSFKEELDKYFKNHTSPDVDLGDISGINASVVNIQKEIDRLNEVA

KNLNESLIDLQELGKYEQYIKWPWYIWLGFIAGLIAIVMVTIMLCCMTSCCSCLKGCCSCGSCCKFDEDD

SEPVLKGVKLHYT

>QHN73810.1 surface glycoprotein [Wuhan seafood market pneumonia virus]

MFVFLVLLPLVSSQCVNLTTRTQLPPAYTNSFTRGVYYPDKVFRSSVLHSTQDLFLPFFSNVTWFHAIHV

SGTNGTKRFDNPVLPFNDGVYFASTEKSNIIRGWIFGTTLDSKTQSLLIVNNATNVVIKVCEFQFCNDPF

LGVYYHKNNKSWMESEFRVYSSANNCTFEYVSQPFLMDLEGKQGNFKNLREFVFKNIDGYFKIYSKHTPI

NLVRDLPQGFSALEPLVDLPIGINITRFQTLLALHRSYLTPGDSSSGWTAGAAAYYVGYLQPRTFLLKYN

ENGTITDAVDCALDPLSETKCTLKSFTVEKGIYQTSNFRVQPTESIVRFPNITNLCPFGEVFNATRFASV

YAWNRKRISNCVADYSVLYNSASFSTFKCYGVSPTKLNDLCFTNVYADSFVIRGDEVRQIAPGQTGKIAD

YNYKLPDDFTGCVIAWNSNNLDSKVGGNYNYLYRLFRKSNLKPFERDISTEIYQAGSTPCNGVEGFNCYF

PLQSYGFQPTNGVGYQPYRVVVLSFELLHAPATVCGPKKSTNLVKNKCVNFNFNGLTGTGVLTESNKKFL

PFQQFGRDIADTTDAVRDPQTLEILDITPCSFGGVSVITPGTNTSNQVAVLYQDVNCTEVPVAIHADQLT

PTWRVYSTGSNVFQTRAGCLIGAEHVNNSYECDIPIGAGICASYQTQTNSPRRARSVASQSIIAYTMSLG

AENSVAYSNNSIAIPTNFTISVTTEILPVSMTKTSVDCTMYICGDSTECSNLLLQYGSFCTQLNRALTGI

AVEQDKNTQEVFAQVKQIYKTPPIKDFGGFNFSQILPDPSKPSKRSFIEDLLFNKVTLADAGFIKQYGDC

LGDIAARDLICAQKFNGLTVLPPLLTDEMIAQYTSALLAGTITSGWTFGAGAALQIPFAMQMAYRFNGIG

VTQNVLYENQKLIANQFNSAIGKIQDSLSSTASALGKLQDVVNQNAQALNTLVKQLSSNFGAISSVLNDI

LSRLDKVEAEVQIDRLITGRLQSLQTYVTQQLIRAAEIRASANLAATKMSECVLGQSKRVDFCGKGYHLM

SFPQSAPHGVVFLHVTYVPAQEKNFTTAPAICHDGKAHFPREGVFVSNGTHWFVTQRNFYEPQIITTDNT

FVSGNCDVVIGIVNNTVYDPLQPELDSFKEELDKYFKNHTSPDVDLGDISGINASVVNIQKEIDRLNEVA

KNLNESLIDLQELGKYEQYIKWPWYIWLGFIAGLIAIVMVTIMLCCMTSCCSCLKGCCSCGSCCKFDEDD

SEPVLKGVKLHYT

>QHN73795.1 surface glycoprotein [Wuhan seafood market pneumonia virus]

MFVFLVLLPLVSSQCVNLTTRTQLPPAYTNSFTRGVYYPDKVFRSSVLHSTQDLFLPFFSNVTWFHAIHV

SGTNGTKRFDNPVLPFNDGVYFASTEKSNIIRGWIFGTTLDSKTQSLLIVNNATNVVIKVCEFQFCNDPF

LGVYYHKNNKSWMESEFRVYSSANNCTFEYVSQPFLMDLEGKQGNFKNLREFVFKNIDGYFKIYSKHTPI

NLVRDLPQGFSALEPLVDLPIGINITRFQTLLALHRSYLTPGDSSSGWTAGAAAYYVGYLQPRTFLLKYN

ENGTITDAVDCALDPLSETKCTLKSFTVEKGIYQTSNFRVQPTESIVRFPNITNLCPFGEVFNATRFASV

YAWNRKRISNCVADYSVLYNSASFSTFKCYGVSPTKLNDLCFTNVYADSFVIRGDEVRQIAPGQTGKIAD

YNYKLPDDFTGCVIAWNSNNLDSKVGGNYNYLYRLFRKSNLKPFERDISTEIYQAGSTPCNGVEGFNCYF

PLQSYGFQPTNGVGYQPYRVVVLSFELLHAPATVCGPKKSTNLVKNKCVNFNFNGLTGTGVLTESNKKFL

PFQQFGRDIADTTDAVRDPQTLEILDITPCSFGGVSVITPGTNTSNQVAVLYQDVNCTEVPVAIHADQLT

PTWRVYSTGSNVFQTRAGCLIGAEHVNNSYECDIPIGAGICASYQTQTNSPRRARSVASQSIIAYTMSLG

AENSVAYSNNSIAIPTNFTISVTTEILPVSMTKTSVDCTMYICGDSTECSNLLLQYGSFCTQLNRALTGI

AVEQDKNTQEVFAQVKQIYKTPPIKDFGGFNFSQILPDPSKPSKRSFIEDLLFNKVTLADAGFIKQYGDC

LGDIAARDLICAQKFNGLTVLPPLLTDEMIAQYTSALLAGTITSGWTFGAGAALQIPFAMQMAYRFNGIG

VTQNVLYENQKLIANQFNSAIGKIQDSLSSTASALGKLQDVVNQNAQALNTLVKQLSSNFGAISSVLNDI

LSRLDKVEAEVQIDRLITGRLQSLQTYVTQQLIRAAEIRASANLAATKMSECVLGQSKRVDFCGKGYHLM

SFPQSAPHGVVFLHVTYVPAQEKNFTTAPAICHDGKAHFPREGVFVSNGTHWFVTQRNFYEPQIITTDNT

FVSGNCDVVIGIVNNTVYDPLQPELDSFKEELDKYFKNHTSPDVDLGDISGINASVVNIQKEIDRLNEVA

KNLNESLIDLQELGKYEQYIKWPWYIWLGFIAGLIAIVMVTIMLCCMTSCCSCLKGCCSCGSCCKFDEDD

SEPVLKGVKLHYT

>QHO60594.1 surface glycoprotein [Wuhan seafood market pneumonia virus]

MFVFLVLLPLVSSQCVNLTTRTQLPPAYTNSFTRGVYYPDKVFRSSVLHSTQDLFLPFFSNVTWFHAIHV

SGTNGTKRFDNPVLPFNDGVYFASTEKSNIIRGWIFGTTLDSKTQSLLIVNNATNVVIKVCEFQFCNDPF

LGVYYHKNNKSWMESEFRVYSSANNCTFEYVSQPFLMDLEGKQGNFKNLREFVFKNIDGYFKIYSKHTPI

NLVRDLPQGFSALEPLVDLPIGINITRFQTLLALHRSYLTPGDSSSGWTAGAAAYYVGYLQPRTFLLKYN

ENGTITDAVDCALDPLSETKCTLKSFTVEKGIYQTSNFRVQPTESIVRFPNITNLCPFGEVFNATRFASV

YAWNRKRISNCVADYSVLYNSASFSTFKCYGVSPTKLNDLCFTNVYADSFVIRGDEVRQIAPGQTGKIAD

YNYKLPDDFTGCVIAWNSNNLDSKVGGNYNYLYRLFRKSNLKPFERDISTEIYQAGSTPCNGVEGFNCYF

PLQSYGFQPTNGVGYQPYRVVVLSFELLHAPATVCGPKKSTNLVKNKCVNFNFNGLTGTGVLTESNKKFL

PFQQFGRDIADTTDAVRDPQTLEILDITPCSFGGVSVITPGTNTSNQVAVLYQDVNCTEVPVAIHADQLT

PTWRVYSTGSNVFQTRAGCLIGAEHVNNSYECDIPIGAGICASYQTQTNSPRRARSVASQSIIAYTMSLG

AENSVAYSNNSIAIPTNFTISVTTEILPVSMTKTSVDCTMYICGDSTECSNLLLQYGSFCTQLNRALTGI

AVEQDKNTQEVFAQVKQIYKTPPIKDFGGFNFSQILPDPSKPSKRSFIEDLLFNKVTLADAGFIKQYGDC

LGDIAARDLICAQKFNGLTVLPPLLTDEMIAQYTSALLAGTITSGWTFGAGAALQIPFAMQMAYRFNGIG

VTQNVLYENQKLIANQFNSAIGKIQDSLSSTASALGKLQDVVNQNAQALNTLVKQLSSNFGAISSVLNDI

LSRLDKVEAEVQIDRLITGRLQSLQTYVTQQLIRAAEIRASANLAATKMSECVLGQSKRVDFCGKGYHLM

SFPQSAPHGVVFLHVTYVPAQEKNFTTAPAICHDGKAHFPREGVFVSNGTHWFVTQRNFYEPQIITTDNT

FVSGNCDVVIGIVNNTVYDPLQPELDSFKEELDKYFKNHTSPDVDLGDISGINASVVNIQKEIDRLNEVA

KNLNESLIDLQELGKYEQYIKWPWYIWLGFIAGLIAIVMVTIMLCCMTSCCSCLKGCCSCGSCCKFDEDD

SEPVLKGVKLHYT

>QHQ82464.1 surface glycoprotein [Wuhan seafood market pneumonia virus]

MFVFLVLLPLVSSQCVNLTTRTQLPPAYTNSFTRGVYYPDKVFRSSVLHSTQDLFLPFFSNVTWFHAIHV

SGTNGTKRFDNPVLPFNDGVYFASTEKSNIIRGWIFGTTLDSKTQSLLIVNNATNVVIKVCEFQFCNDPF

LGVYYHKNNKSWMESEFRVYSSANNCTFEYVSQPFLMDLEGKQGNFKNLREFVFKNIDGYFKIYSKHTPI

NLVRDLPQGFSALEPLVDLPIGINITRFQTLLALHRSYLTPGDSSSGWTAGAAAYYVGYLQPRTFLLKYN

ENGTITDAVDCALDPLSETKCTLKSFTVEKGIYQTSNFRVQPTESIVRFPNITNLCPFGEVFNATRFASV

YAWNRKRISNCVADYSVLYNSASFSTFKCYGVSPTKLNDLCFTNVYADSFVIRGDEVRQIAPGQTGKIAD

YNYKLPDDFTGCVIAWNSNNLDSKVGGNYNYLYRLFRKSNLKPFERDISTEIYQAGSTPCNGVEGFNCYF

PLQSYGFQPTNGVGYQPYRVVVLSFELLHAPATVCGPKKSTNLVKNKCVNFNFNGLTGTGVLTESNKKFL

PFQQFGRDIADTTDAVRDPQTLEILDITPCSFGGVSVITPGTNTSNQVAVLYQDVNCTEVPVAIHADQLT

PTWRVYSTGSNVFQTRAGCLIGAEHVNNSYECDIPIGAGICASYQTQTNSPRRARSVASQSIIAYTMSLG

AENSVAYSNNSIAIPTNFTISVTTEILPVSMTKTSVDCTMYICGDSTECSNLLLQYGSFCTQLNRALTGI

AVEQDKNTQEVFAQVKQIYKTPPIKDFGGFNFSQILPDPSKPSKRSFIEDLLFNKVTLADAGFIKQYGDC

LGDIAARDLICAQKFNGLTVLPPLLTDEMIAQYTSALLAGTITSGWTFGAGAALQIPFAMQMAYRFNGIG

VTQNVLYENQKLIANQFNSAIGKIQDSLSSTASALGKLQDVVNQNAQALNTLVKQLSSNFGAISSVLNDI

LSRLDKVEAEVQIDRLITGRLQSLQTYVTQQLIRAAEIRASANLAATKMSECVLGQSKRVDFCGKGYHLM

SFPQSAPHGVVFLHVTYVPAQEKNFTTAPAICHDGKAHFPREGVFVSNGTHWFVTQRNFYEPQIITTDNT

FVSGNCDVVIGIVNNTVYDPLQPELDSFKEELDKYFKNHTSPDVDLGDISGINASVVNIQKEIDRLNEVA

KNLNESLIDLQELGKYEQYIKWPWYIWLGFIAGLIAIVMVTIMLCCMTSCCSCLKGCCSCGSCCKFDEDD

SEPVLKGVKLHYT

>QHQ71973.1 surface glycoprotein [Wuhan seafood market pneumonia virus]

MFVFLVLLPLVSSQCVNLTTRTQLPPAYTNSFTRGVYYPDKVFRSSVLHSTQDLFLPFFSNVTWFHAIHV

SGTNGTKRFDNPVLPFNDGVYFASTEKSNIIRGWIFGTTLDSKTQSLLIVNNATNVVIKVCEFQFCNDPF

LGVYYHKNNKSWMESEFRVYSSANNCTFEYVSQPFLMDLEGKQGNFKNLREFVFKNIDGYFKIYSKHTPI

NLVRDLPQGFSALEPLVDLPIGINITRFQTLLALHRSYLTPGDSSSGWTAGAAAYYVGYLQPRTFLLKYN

ENGTITDAVDCALDPLSETKCTLKSFTVEKGIYQTSNFRVQPTESIVRFPNITNLCPFGEVFNATRFASV

YAWNRKRISNCVADYSVLYNSASFSTFKCYGVSPTKLNDLCFTNVYADSFVIRGDEVRQIAPGQTGKIAD

YNYKLPDDFTGCVIAWNSNNLDSKVGGNYNYLYRLFRKSNLKPFERDISTEIYQAGSTPCNGVEGFNCYF

PLQSYGFQPTNGVGYQPYRVVVLSFELLHAPATVCGPKKSTNLVKNKCVNFNFNGLTGTGVLTESNKKFL

PFQQFGRDIADTTDAVRDPQTLEILDITPCSFGGVSVITPGTNTSNQVAVLYQDVNCTEVPVAIHADQLT

PTWRVYSTGSNVFQTRAGCLIGAEHVNNSYECDIPIGAGICASYQTQTNSPRRARSVASQSIIAYTMSLG

AENSVAYSNNSIAIPTNFTISVTTEILPVSMTKTSVDCTMYICGDSTECSNLLLQYGSFCTQLNRALTGI

AVEQDKNTQEVFAQVKQIYKTPPIKDFGGFNFSQILPDPSKPSKRSFIEDLLFNKVTLADAGFIKQYGDC

LGDIAARDLICAQKFNGLTVLPPLLTDEMIAQYTSALLAGTITSGWTFGAGAALQIPFAMQMAYRFNGIG

VTQNVLYENQKLIANQFNSAIGKIQDSLSSTASALGKLQDVVNQNAQALNTLVKQLSSNFGAISSVLNDI

LSRLDKVEAEVQIDRLITGRLQSLQTYVTQQLIRAAEIRASANLAATKMSECVLGQSKRVDFCGKGYHLM

SFPQSAPHGVVFLHVTYVPAQEKNFTTAPAICHDGKAHFPREGVFVSNGTHWFVTQRNFYEPQIITTDNT

FVSGNCDVVIGIVNNTVYDPLQPELDSFKEELDKYFKNHTSPDVDLGDISGINASVVNIQKEIDRLNEVA

KNLNESLIDLQELGKYEQYIKWPWYIWLGFIAGLIAIVMVTIMLCCMTSCCSCLKGCCSCGSCCKFDEDD

SEPVLKGVKLHYT

>QHQ71963.1 surface glycoprotein [Wuhan seafood market pneumonia virus]

MFVFLVLLPLVSSQCVNLTTRTQLPPAYTNSFTRGVYYPDKVFRSSVLHSTQDLFLPFFSNVTWFHAIHV

SGTNGTKRFDNPVLPFNDGVYFASTEKSNIIRGWIFGTTLDSKTQSLLIVNNATNVVIKVCEFQFCNDPF

LGVYYHKNNKSWMESEFRVYSSANNCTFEYVSQPFLMDLEGKQGNFKNLREFVFKNIDGYFKIYSKHTPI

NLVRDLPQGFSALEPLVDLPIGINITRFQTLLALHRSYLTPGDSSSGWTAGAAAYYVGYLQPRTFLLKYN

ENGTITDAVDCALDPLSETKCTLKSFTVEKGIYQTSNFRVQPTESIVRFPNITNLCPFGEVFNATRFASV

YAWNRKRISNCVADYSVLYNSASFSTFKCYGVSPTKLNDLCFTNVYADSFVIRGDEVRQIAPGQTGKIAD

YNYKLPDDFTGCVIAWNSNNLDSKVGGNYNYLYRLFRKSNLKPFERDISTEIYQAGSTPCNGVEGFNCYF

PLQSYGFQPTNGVGYQPYRVVVLSFELLHAPATVCGPKKSTNLVKNKCVNFNFNGLTGTGVLTESNKKFL

PFQQFGRDIADTTDAVRDPQTLEILDITPCSFGGVSVITPGTNTSNQVAVLYQDVNCTEVPVAIHADQLT

PTWRVYSTGSNVFQTRAGCLIGAEHVNNSYECDIPIGAGICASYQTQTNSPRRARSVASQSIIAYTMSLG

AENSVAYSNNSIAIPTNFTISVTTEILPVSMTKTSVDCTMYICGDSTECSNLLLQYGSFCTQLNRALTGI

AVEQDKNTQEVFAQVKQIYKTPPIKDFGGFNFSQILPDPSKPSKRSFIEDLLFNKVTLADAGFIKQYGDC

LGDIAARDLICAQKFNGLTVLPPLLTDEMIAQYTSALLAGTITSGWTFGAGAALQIPFAMQMAYRFNGIG

VTQNVLYENQKLIANQFNSAIGKIQDSLSSTASALGKLQDVVNQNAQALNTLVKQLSSNFGAISSVLNDI

LSRLDKVEAEVQIDRLITGRLQSLQTYVTQQLIRAAEIRASANLAATKMSECVLGQSKRVDFCGKGYHLM

SFPQSAPHGVVFLHVTYVPAQEKNFTTAPAICHDGKAHFPREGVFVSNGTHWFVTQRNFYEPQIITTDNT

FVSGNCDVVIGIVNNTVYDPLQPELDSFKEELDKYFKNHTSPDVDLGDISGINASVVNIQKEIDRLNEVA

KNLNESLIDLQELGKYEQYIKWPWYIWLGFIAGLIAIVMVTIMLCCMTSCCSCLKGCCSCGSCCKFDEDD

SEPVLKGVKLHYT

>QHO62112.1 surface glycoprotein [Wuhan seafood market pneumonia virus]

MFVFLVLLPLVSSQCVNLTTRTQLPPAYTNSFTRGVYYPDKVFRSSVLHSTQDLFLPFFSNVTWFHAIHV

SGTNGTKRFDNPVLPFNDGVYFASTEKSNIIRGWIFGTTLDSKTQSLLIVNNATNVVIKVCEFQFCNDPF

LGVYYHKNNKSWMESEFRVYSSANNCTFEYVSQPFLMDLEGKQGNFKNLREFVFKNIDGYFKIYSKHTPI

NLVRDLPQGFSALEPLVDLPIGINITRFQTLLALHRSYLTPGDSSSGWTAGAAAYYVGYLQPRTFLLKYN

ENGTITDAVDCALDPLSETKCTLKSFTVEKGIYQTSNFRVQPTESIVRFPNITNLCPFGEVFNATRFASV

YAWNRKRISNCVADYSVLYNSASFSTFKCYGVSPTKLNDLCFTNVYADSFVIRGDEVRQIAPGQTGKIAD

YNYKLPDDFTGCVIAWNSNNLDSKVGGNYNYLYRLFRKSNLKPFERDISTEIYQAGSTPCNGVEGFNCYF

PLQSYGFQPTNGVGYQPYRVVVLSFELLHAPATVCGPKKSTNLVKNKCVNFNFNGLTGTGVLTESNKKFL

PFQQFGRDIADTTDAVRDPQTLEILDITPCSFGGVSVITPGTNTSNQVAVLYQDVNCTEVPVAIHADQLT

PTWRVYSTGSNVFQTRAGCLIGAEHVNNSYECDIPIGAGICASYQTQTNSPRRARSVASQSIIAYTMSLG

AENSVAYSNNSIAIPTNFTISVTTEILPVSMTKTSVDCTMYICGDSTECSNLLLQYGSFCTQLNRALTGI

AVEQDKNTQEVFAQVKQIYKTPPIKDFGGFNFSQILPDPSKPSKRSFIEDLLFNKVTLADAGFIKQYGDC

LGDIAARDLICAQKFNGLTVLPPLLTDEMIAQYTSALLAGTITSGWTFGAGAALQIPFAMQMAYRFNGIG

VTQNVLYENQKLIANQFNSAIGKIQDSLSSTASALGKLQDVVNQNAQALNTLVKQLSSNFGAISSVLNDI

LSRLDKVEAEVQIDRLITGRLQSLQTYVTQQLIRAAEIRASANLAATKMSECVLGQSKRVDFCGKGYHLM

SFPQSAPHGVVFLHVTYVPAQEKNFTTAPAICHDGKAHFPREGVFVSNGTHWFVTQRNFYEPQIITTDNT

FVSGNCDVVIGIVNNTVYDPLQPELDSFKEELDKYFKNHTSPDVDLGDISGINASVVNIQKEIDRLNEVA

KNLNESLIDLQELGKYEQYIKWPWYIWLGFIAGLIAIVMVTIMLCCMTSCCSCLKGCCSCGSCCKFDEDD

SEPVLKGVKLHYT

>QHO62107.1 surface glycoprotein [Wuhan seafood market pneumonia virus]

MFVFLVLLPLVSSQCVNLTTRTQLPPAYTNSFTRGVYYPDKVFRSSVLHSTQDLFLPFFSNVTWFHAIHV

SGTNGTKRFDNPVLPFNDGVYFASTEKSNIIRGWIFGTTLDSKTQSLLIVNNATNVVIKVCEFQFCNDPF

LGVYYHKNNKSWMESEFRVYSSANNCTFEYVSQPFLMDLEGKQGNFKNLREFVFKNIDGYFKIYSKHTPI

NLVRDLPQGFSALEPLVDLPIGINITRFQTLLALHRSYLTPGDSSSGWTAGAAAYYVGYLQPRTFLLKYN

ENGTITDAVDCALDPLSETKCTLKSFTVEKGIYQTSNFRVQPTESIVRFPNITNLCPFGEVFNATRFASV

YAWNRKRISNCVADYSVLYNSASFSTFKCYGVSPTKLNDLCFTNVYADSFVIRGDEVRQIAPGQTGKIAD

YNYKLPDDFTGCVIAWNSNNLDSKVGGNYNYLYRLFRKSNLKPFERDISTEIYQAGSTPCNGVEGFNCYF

PLQSYGFQPTNGVGYQPYRVVVLSFELLHAPATVCGPKKSTNLVKNKCVNFNFNGLTGTGVLTESNKKFL

PFQQFGRDIADTTDAVRDPQTLEILDITPCSFGGVSVITPGTNTSNQVAVLYQDVNCTEVPVAIHADQLT

PTWRVYSTGSNVFQTRAGCLIGAEHVNNSYECDIPIGAGICASYQTQTNSPRRARSVASQSIIAYTMSLG

AENSVAYSNNSIAIPTNFTISVTTEILPVSMTKTSVDCTMYICGDSTECSNLLLQYGSFCTQLNRALTGI

AVEQDKNTQEVFAQVKQIYKTPPIKDFGGFNFSQILPDPSKPSKRSFIEDLLFNKVTLADAGFIKQYGDC

LGDIAARDLICAQKFNGLTVLPPLLTDEMIAQYTSALLAGTITSGWTFGAGAALQIPFAMQMAYRFNGIG

VTQNVLYENQKLIANQFNSAIGKIQDSLSSTASALGKLQDVVNQNAQALNTLVKQLSSNFGAISSVLNDI

LSRLDKVEAEVQIDRLITGRLQSLQTYVTQQLIRAAEIRASANLAATKMSECVLGQSKRVDFCGKGYHLM

SFPQSAPHGVVFLHVTYVPAQEKNFTTAPAICHDGKAHFPREGVFVSNGTHWFVTQRNFYEPQIITTDNT

FVSGNCDVVIGIVNNTVYDPLQPELDSFKEELDKYFKNHTSPDVDLGDISGINASVVNIQKEIDRLNEVA

KNLNESLIDLQELGKYEQYIKWPWYIWLGFIAGLIAIVMVTIMLCCMTSCCSCLKGCCSCGSCCKFDEDD

SEPVLKGVKLHYT

>YP_009724390.1 surface glycoprotein [Wuhan seafood market pneumonia virus]

MFVFLVLLPLVSSQCVNLTTRTQLPPAYTNSFTRGVYYPDKVFRSSVLHSTQDLFLPFFSNVTWFHAIHV

SGTNGTKRFDNPVLPFNDGVYFASTEKSNIIRGWIFGTTLDSKTQSLLIVNNATNVVIKVCEFQFCNDPF

LGVYYHKNNKSWMESEFRVYSSANNCTFEYVSQPFLMDLEGKQGNFKNLREFVFKNIDGYFKIYSKHTPI

NLVRDLPQGFSALEPLVDLPIGINITRFQTLLALHRSYLTPGDSSSGWTAGAAAYYVGYLQPRTFLLKYN

ENGTITDAVDCALDPLSETKCTLKSFTVEKGIYQTSNFRVQPTESIVRFPNITNLCPFGEVFNATRFASV

YAWNRKRISNCVADYSVLYNSASFSTFKCYGVSPTKLNDLCFTNVYADSFVIRGDEVRQIAPGQTGKIAD

YNYKLPDDFTGCVIAWNSNNLDSKVGGNYNYLYRLFRKSNLKPFERDISTEIYQAGSTPCNGVEGFNCYF

PLQSYGFQPTNGVGYQPYRVVVLSFELLHAPATVCGPKKSTNLVKNKCVNFNFNGLTGTGVLTESNKKFL

PFQQFGRDIADTTDAVRDPQTLEILDITPCSFGGVSVITPGTNTSNQVAVLYQDVNCTEVPVAIHADQLT

PTWRVYSTGSNVFQTRAGCLIGAEHVNNSYECDIPIGAGICASYQTQTNSPRRARSVASQSIIAYTMSLG

AENSVAYSNNSIAIPTNFTISVTTEILPVSMTKTSVDCTMYICGDSTECSNLLLQYGSFCTQLNRALTGI

AVEQDKNTQEVFAQVKQIYKTPPIKDFGGFNFSQILPDPSKPSKRSFIEDLLFNKVTLADAGFIKQYGDC

LGDIAARDLICAQKFNGLTVLPPLLTDEMIAQYTSALLAGTITSGWTFGAGAALQIPFAMQMAYRFNGIG

VTQNVLYENQKLIANQFNSAIGKIQDSLSSTASALGKLQDVVNQNAQALNTLVKQLSSNFGAISSVLNDI

LSRLDKVEAEVQIDRLITGRLQSLQTYVTQQLIRAAEIRASANLAATKMSECVLGQSKRVDFCGKGYHLM

SFPQSAPHGVVFLHVTYVPAQEKNFTTAPAICHDGKAHFPREGVFVSNGTHWFVTQRNFYEPQIITTDNT

FVSGNCDVVIGIVNNTVYDPLQPELDSFKEELDKYFKNHTSPDVDLGDISGINASVVNIQKEIDRLNEVA

KNLNESLIDLQELGKYEQYIKWPWYIWLGFIAGLIAIVMVTIMLCCMTSCCSCLKGCCSCGSCCKFDEDD

SEPVLKGVKLHYT

>QHO62877.1 surface glycoprotein [Wuhan seafood market pneumonia virus]

MFVFLVLLPLVSSQCVNLTTRTQLPPAYTNSFTRGVYYPDKVFRSSVLHSTQDLFLPFFSNVTWFHAIHV

SGTNGTKRFDNPVLPFNDGVYFASTEKSNIIRGWIFGTTLDSKTQSLLIVNNATNVVIKVCEFQFCNDPF

LGVYYHKNNKSWMESEFRVYSSANNCTFEYVSQPFLMDLEGKQGNFKNLREFVFKNIDGYFKIYSKHTPI

NLVRDLPQGFSALEPLVDLPIGINITRFQTLLALHRSYLTPGDSSSGWTAGAAAYYVGYLQPRTFLLKYN

ENGTITDAVDCALDPLSETKCTLKSFTVEKGIYQTSNFRVQPTESIVRFPNITNLCPFGEVFNATRFASV

YAWNRKRISNCVADYSVLYNSASFSTFKCYGVSPTKLNDLCFTNVYADSFVIRGDEVRQIAPGQTGKIAD

YNYKLPDDFTGCVIAWNSNNLDSKVGGNYNYLYRLFRKSNLKPFERDISTEIYQAGSTPCNGVEGFNCYF

PLQSYGFQPTNGVGYQPYRVVVLSFELLHAPATVCGPKKSTNLVKNKCVNFNFNGLTGTGVLTESNKKFL

PFQQFGRDIADTTDAVRDPQTLEILDITPCSFGGVSVITPGTNTSNQVAVLYQDVNCTEVPVAIHADQLT

PTWRVYSTGSNVFQTRAGCLIGAEHVNNSYECDIPIGAGICASYQTQTNSPRRARSVASQSIIAYTMSLG

AENSVAYSNNSIAIPTNFTISVTTEILPVSMTKTSVDCTMYICGDSTECSNLLLQYGSFCTQLNRALTGI

AVEQDKNTQEVFAQVKQIYKTPPIKDFGGFNFSQILPDPSKPSKRSFIEDLLFNKVTLADAGFIKQYGDC

LGDIAARDLICAQKFNGLTVLPPLLTDEMIAQYTSALLAGTITSGWTFGAGAALQIPFAMQMAYRFNGIG

VTQNVLYENQKLIANQFNSAIGKIQDSLSSTASALGKLQDVVNQNAQALNTLVKQLSSNFGAISSVLNDI

LSRLDKVEAEVQIDRLITGRLQSLQTYVTQQLIRAAEIRASANLAATKMSECVLGQSKRVDFCGKGYHLM

SFPQSAPHGVVFLHVTYVPAQEKNFTTAPAICHDGKAHFPREGVFVSNGTHWFVTQRNFYEPQIITTDNT

FVSGNCDVVIGIVNNTVYDPLQPELDSFKEELDKYFKNHTSPDVDLGDISGINASVVNIQKEIDRLNEVA

KNLNESLIDLQELGKYEQYIKWPWYIWLGFIAGLIAIVMVTIMLCCMTSCCSCLKGCCSCGSCCKFDEDD

SEPVLKGVKLHYT

>QHN73824.1 surface glycoprotein, partial [Wuhan seafood market pneumonia virus]

NVYADSFVIRGDEVRQIAPGQTGKIADYNYKLPDD

>QHN73823.1 surface glycoprotein, partial [Wuhan seafood market pneumonia virus]

NVYADSFVIRGDEVRQIAPGQTGKIADYNYKLPDD

>QHN73822.1 surface glycoprotein, partial [Wuhan seafood market pneumonia virus]

NVYADSFVIRGDEVRQIAPGQTGKIADYNYKLPDD

>QHN73808.1 surface glycoprotein, partial [Wuhan seafood market pneumonia virus]

NVYADSFVIRGDEVRQIAPGQTGKIADYNYKLPDD

>QHN73807.1 surface glycoprotein, partial [Wuhan seafood market pneumonia virus]

NVYADSFVIRGDEVRQIAPGQTGKIADYNYKLPDD

>QHN73806.1 surface glycoprotein, partial [Wuhan seafood market pneumonia virus]

NVYADSFVIRGDEVRQIAPGQTGKIADYNYKLPDD

>QHN73805.1 surface glycoprotein, partial [Wuhan seafood market pneumonia virus]

NVYADSFVIRGDEVRQIAPGQTGKIADYNYKLPDD

>QHD43416.1 surface glycoprotein [Wuhan seafood market pneumonia virus]

MFVFLVLLPLVSSQCVNLTTRTQLPPAYTNSFTRGVYYPDKVFRSSVLHSTQDLFLPFFSNVTWFHAIHV

SGTNGTKRFDNPVLPFNDGVYFASTEKSNIIRGWIFGTTLDSKTQSLLIVNNATNVVIKVCEFQFCNDPF

LGVYYHKNNKSWMESEFRVYSSANNCTFEYVSQPFLMDLEGKQGNFKNLREFVFKNIDGYFKIYSKHTPI

NLVRDLPQGFSALEPLVDLPIGINITRFQTLLALHRSYLTPGDSSSGWTAGAAAYYVGYLQPRTFLLKYN

ENGTITDAVDCALDPLSETKCTLKSFTVEKGIYQTSNFRVQPTESIVRFPNITNLCPFGEVFNATRFASV

YAWNRKRISNCVADYSVLYNSASFSTFKCYGVSPTKLNDLCFTNVYADSFVIRGDEVRQIAPGQTGKIAD

YNYKLPDDFTGCVIAWNSNNLDSKVGGNYNYLYRLFRKSNLKPFERDISTEIYQAGSTPCNGVEGFNCYF

PLQSYGFQPTNGVGYQPYRVVVLSFELLHAPATVCGPKKSTNLVKNKCVNFNFNGLTGTGVLTESNKKFL

PFQQFGRDIADTTDAVRDPQTLEILDITPCSFGGVSVITPGTNTSNQVAVLYQDVNCTEVPVAIHADQLT

PTWRVYSTGSNVFQTRAGCLIGAEHVNNSYECDIPIGAGICASYQTQTNSPRRARSVASQSIIAYTMSLG

AENSVAYSNNSIAIPTNFTISVTTEILPVSMTKTSVDCTMYICGDSTECSNLLLQYGSFCTQLNRALTGI

AVEQDKNTQEVFAQVKQIYKTPPIKDFGGFNFSQILPDPSKPSKRSFIEDLLFNKVTLADAGFIKQYGDC

LGDIAARDLICAQKFNGLTVLPPLLTDEMIAQYTSALLAGTITSGWTFGAGAALQIPFAMQMAYRFNGIG

VTQNVLYENQKLIANQFNSAIGKIQDSLSSTASALGKLQDVVNQNAQALNTLVKQLSSNFGAISSVLNDI

LSRLDKVEAEVQIDRLITGRLQSLQTYVTQQLIRAAEIRASANLAATKMSECVLGQSKRVDFCGKGYHLM

SFPQSAPHGVVFLHVTYVPAQEKNFTTAPAICHDGKAHFPREGVFVSNGTHWFVTQRNFYEPQIITTDNT

FVSGNCDVVIGIVNNTVYDPLQPELDSFKEELDKYFKNHTSPDVDLGDISGINASVVNIQKEIDRLNEVA

KNLNESLIDLQELGKYEQYIKWPWYIWLGFIAGLIAIVMVTIMLCCMTSCCSCLKGCCSCGSCCKFDEDD

SEPVLKGVKLHYT

**24 protein sequences of membrane glycoprotein for COVID-19**

>QHW06062.1 membrane glycoprotein [Wuhan seafood market pneumonia virus]

MADSNGTITVEELKKLLEQWNLVIGFLFLTWICLLQFAYANRNRFLYIIKLIFLWLLWPVTLACFVLAAV

YRINWITGGIAIAMACLVGLMWLSYFIASFRLFARTRSMWSFNPETNILLNVPLHGTILTRPLLESELVI

GAVILRGHLRIAGHHLGRCDIKDLPKEITVATSRTLSYYKLGASQRVAGDSGFAAYSRYRIGNYKLNTDH

SSSSDNIALLVQ

>QHW06052.1 membrane glycoprotein [Wuhan seafood market pneumonia virus]

MADSNGTITVEELKKLLEQWNLVIGFLFLTWICLLQFAYANRNRFLYIIKLIFLWLLWPVTLACFVLAAV

YRINWITGGIAIAMACLVGLMWLSYFIASFRLFARTRSMWSFNPETNILLNVPLHGTILTRPLLESELVI

GAVILRGHLRIAGHHLGRCDIKDLPKEITVATSRTLSYYKLGASQRVAGDSGFAAYSRYRIGNYKLNTDH

SSSSDNIALLVQ

>QHW06042.1 membrane glycoprotein [Wuhan seafood market pneumonia virus]

MADSNGTITVEELKKLLEQWNLVIGFLFLTWICLLQFAYANRNRFLYIIKLIFLWLLWPVTLACFVLAAV

YRINWITGGIAIAMACLVGLMWLSYFIASFRLFARTRSMWSFNPETNILLNVPLHGTILTRPLLESELVI

GAVILRGHLRIAGHHLGRCDIKDLPKEITVATSRTLSYYKLGASQRVAGDSGFAAYSRYRIGNYKLNTDH

SSSSDNIALLVQ

>BBW89520.1 membrane glycoprotein [Wuhan seafood market pneumonia virus]

MADSNGTITVEELKKLLEQWNLVIGFLFLTWICLLQFAYANRNRFLYIIKLIFLWLLWPVTLACFVLAAV

YRINWITGGIAIAMACLVGLMWLSYFIASFRLFARTRSMWSFNPETNILLNVPLHGTILTRPLLESELVI

GAVILRGHLRIAGHHLGRCDIKDLPKEITVATSRTLSYYKLGASQRVAGDSGFAAYSRYRIGNYKLNTDH

SSSSDNIALLVQ

>QHU79207.1 membrane glycoprotein [Wuhan seafood market pneumonia virus]

MADSNGTITVEELKKLLEQWNLVIGFLFLTWICLLQFAYANRNRFLYIIKLIFLWLLWPVTLACFVLAAV

YRINWITGGIAIAMACLVGLMWLSYFIASFRLFARTRSMWSFNPETNILLNVPLHGTILTRPLLESELVI

GAVILRGHLRIAGHHLGRCDIKDLPKEITVATSRTLSYYKLGASQRVAGDSGFAAYSRYRIGNYKLNTDH

SSSSDNIALLVQ

>QHU79197.1 membrane glycoprotein [Wuhan seafood market pneumonia virus]

MADSNGTITVEELKKLLEQWNLVIGFLFLTWICLLQFAYANRNRFLYIIKLIFLWLLWPVTLACFVLAAV

YRINWITGGIAIAMACLVGLMWLSYFIASFRLFARTRSMWSFNPETNILLNVPLHGTILTRPLLESELVI

GAVILRGHLRIAGHHLGRCDIKDLPKEITVATSRTLSYYKLGASQRVAGDSGFAAYSRYRIGNYKLNTDH

SSSSDNIALLVQ

>QHU79176.1 membrane glycoprotein [Wuhan seafood market pneumonia virus]

MADSNGTITVEELKKLLEQWNLVIGFLFLTWICLLQFAYANRNRFLYIIKLIFLWLLWPVTLACFVLAAV

YRINWITGGIAIAMACLVGLMWLSYFIASFRLFARTRSMWSFNPETNILLNVPLHGTILTRPLLESELVI

GAVILRGHLRIAGHHLGRCDIKDLPKEITVATSRTLSYYKLGASQRVAGDSGFAAYSRYRIGNYKLNTDH

SSSSDNIALLVQ

>QHU36867.1 membrane glycoprotein [Wuhan seafood market pneumonia virus]

MADSNGTITVEELKKLLEQWNLVIGFLFLTWICLLQFAYANRNRFLYIIKLIFLWLLWPVTLACFVLAAV

YRINWITGGIAIAMACLVGLMWLSYFIASFRLFARTRSMWSFNPETNILLNVPLHGTILTRPLLESELVI

GAVILRGHLRIAGHHLGRCDIKDLPKEITVATSRTLSYYKLGASQRVAGDSGFAAYSRYRIGNYKLNTDH

SSSSDNIALLVQ

>QHU36857.1 membrane glycoprotein [Wuhan seafood market pneumonia virus]

MADSNGTITVEELKKLLEQWNLVIGFLFLTWICLLQFAYANRNRFLYIIKLIFLWLLWPVTLACFVLAAV

YRINWITGGIAIAMACLVGLMWLSYFIASFRLFARTRSMWSFNPETNILLNVPLHGTILTRPLLESELVI

GAVILRGHLRIAGHHLGRCDIKDLPKEITVATSRTLSYYKLGASQRVAGDSGFAAYSRYRIGNYKLNTDH

SSSSDNIALLVQ

>QHU36847.1 membrane glycoprotein [Wuhan seafood market pneumonia virus]

MADSNGTITVEELKKLLEQWNLVIGFLFLTWICLLQFAYANRNRFLYIIKLIFLWLLWPVTLACFVLAAV

YRINWITGGIAIAMACLVGLMWLSYFIASFRLFARTRSMWSFNPETNILLNVPLHGTILTRPLLESELVI

GAVILRGHLRIAGHHLGRCDIKDLPKEITVATSRTLSYYKLGASQRVAGDSGFAAYSRYRIGNYKLNTDH

SSSSDNIALLVQ

>QHU36837.1 membrane glycoprotein [Wuhan seafood market pneumonia virus]

MADSNGTITVEELKKLLEQWNLVIGFLFLTWICLLQFAYANRNRFLYIIKLIFLWLLWPVTLACFVLAAV

YRINWITGGIAIAMACLVGLMWLSYFIASFRLFARTRSMWSFNPETNILLNVPLHGTILTRPLLESELVI

GAVILRGHLRIAGHHLGRCDIKDLPKEITVATSRTLSYYKLGASQRVAGDSGFAAYSRYRIGNYKLNTDH

SSSSDNIALLVQ

>QHU36827.1 membrane glycoprotein [Wuhan seafood market pneumonia virus]

MADSNGTITVEELKKLLEQWNLVIGFLFLTWICLLQFAYANRNRFLYIIKLIFLWLLWPVTLACFVLAAV

YRINWITGGIAIAMACLVGLMWLSYFIASFRLFARTRSMWSFNPETNILLNVPLHGTILTRPLLESELVI

GAVILRGHLRIAGHHLGRCDIKDLPKEITVATSRTLSYYKLGASQRVAGDSGFAAYSRYRIGNYKLNTDH

SSSSDNIALLVQ

>QHR93622.1 membrane glycoprotein, partial [Wuhan seafood market pneumonia virus]

LACFVLAAVYRINWITGGIAIAMACLVGLMWLSYFIASFRLFARTRSMWSFNPETNILLNVPLHGTILTR

PLLESELVIGAVILRGHLRIAGHHLGRCDIKDLPKEI

>QHR93621.1 membrane glycoprotein, partial [Wuhan seafood market pneumonia virus]

LACFVLAAVYRINWITGGIAIAMACLVGLMWLSYFIASFRLFARTRSMWSFNPETNILLNVPLHGTILTR

PLLESELVIGAVILRGHLRIAGHHLGRCDIKDLPKEI

>QHR84452.1 membrane glycoprotein [Wuhan seafood market pneumonia virus]

MADSNGTITVEELKKLLEQWNLVIGFLFLTWICLLQFAYANRNRFLYIIKLIFLWLLWPVTLACFVLAAV

YRINWITGGIAIAMACLVGLMWLSYFIASFRLFARTRSMWSFNPETNILLNVPLHGTILTRPLLESELVI

GAVILRGHLRIAGHHLGRCDIKDLPKEITVATSRTLSYYKLGASQRVAGDSGFAAYSRYRIGNYKLNTDH

SSSSDNIALLVQ

>QHN73813.1 membrane glycoprotein [Wuhan seafood market pneumonia virus]

MADSNGTITVEELKKLLEQWNLVIGFLFLTWICLLQFAYANRNRFLYIIKLIFLWLLWPVTLACFVLAAV

YRINWITGGIAIAMACLVGLMWLSYFIASFRLFARTRSMWSFNPETNILLNVPLHGTILTRPLLESELVI

GAVILRGHLRIAGHHLGRCDIKDLPKEITVATSRTLSYYKLGASQRVAGDSGFAAYSRYRIGNYKLNTDH

SSSSDNIALLVQ

>QHN73798.1 membrane glycoprotein [Wuhan seafood market pneumonia virus]

MADSNGTITVEELKKLLEQWNLVIGFLFLTWICLLQFAYANRNRFLYIIKLIFLWLLWPVTLACFVLAAV

YRINWITGGIAIAMACLVGLMWLSYFIASFRLFARTRSMWSFNPETNILLNVPLHGTILTRPLLESELVI

GAVILRGHLRIAGHHLGRCDIKDLPKEITVATSRTLSYYKLGASQRVAGDSGFAAYSRYRIGNYKLNTDH

SSSSDNIALLVQ

>QHO60597.1 membrane glycoprotein [Wuhan seafood market pneumonia virus]

MADSNGTITVEELKKLLEQWNLVIGFLFLTWICLLQFAYANRNRFLYIIKLIFLWLLWPVTLACFVLAAV

YRINWITGGIAIAMACLVGLMWLSYFIASFRLFARTRSMWSFNPETNILLNVPLHGTILTRPLLESELVI

GAVILRGHLRIAGHHLGRCDIKDLPKEITVATSRTLSYYKLGASQRVAGDSGFAAYSRYRIGNYKLNTDH

SSSSDNIALLVQ

>QHQ82467.1 membrane glycoprotein [Wuhan seafood market pneumonia virus]

MADSNGTITVEELKKLLEQWNLVIGFLFLTWICLLQFAYANRNRFLYIIKLIFLWLLWPVTLACFVLAAV

YRINWITGGIAIAMACLVGLMWLSYFIASFRLFARTRSMWSFNPETNILLNVPLHGTILTRPLLESELVI

GAVILRGHLRIAGHHLGRCDIKDLPKEITVATSRTLSYYKLGASQRVAGDSGFAAYSRYRIGNYKLNTDH

SSSSDNIALLVQ

>QHQ71976.1 membrane glycoprotein [Wuhan seafood market pneumonia virus]

MADSNGTITVEELKKLLEQWNLVIGFLFLTWICLLQFAYANRNRFLYIIKLIFLWLLWPVTLACFVLAAV

YRINWITGGIAIAMACLVGLMWLSYFIASFRLFARTRSMWSFNPETNILLNVPLHGTILTRPLLESELVI

GAVILRGHLRIAGHHLGRCDIKDLPKEITVATSRTLSYYKLGASQRVAGDSGFAAYSRYRIGNYKLNTDH

SSSSDNIALLVQ

>QHQ71966.1 membrane glycoprotein [Wuhan seafood market pneumonia virus]

MADSNGTITVEELKKLLEQWNLVIGFLFLTWICLLQFAYANRNRFLYIIKLIFLWLLWPVTLACFVLAAV

YRINWITGGIAIAMACLVGLMWLSYFIASFRLFARTRSMWSFNPETNILLNVPLHGTILTRPLLESELVI

GAVILRGHLRIAGHHLGRCDIKDLPKEITVATSRTLSYYKLGASQRVAGDSGFAAYSRYRIGNYKLNTDH

SSSSDNIALLVQ

>YP_009724393.1 membrane glycoprotein [Wuhan seafood market pneumonia virus]

MADSNGTITVEELKKLLEQWNLVIGFLFLTWICLLQFAYANRNRFLYIIKLIFLWLLWPVTLACFVLAAV

YRINWITGGIAIAMACLVGLMWLSYFIASFRLFARTRSMWSFNPETNILLNVPLHGTILTRPLLESELVI

GAVILRGHLRIAGHHLGRCDIKDLPKEITVATSRTLSYYKLGASQRVAGDSGFAAYSRYRIGNYKLNTDH

SSSSDNIALLVQ

>QHO62880.1 membrane glycoprotein [Wuhan seafood market pneumonia virus]

MADSNGTITVEELKKLLEQWNLVIGFLFLTWICLLQFAYANRNRFLYIIKLIFLWLLWPVTLACFVLAAV

YRINWITGGIAIAMACLVGLMWLSYFIASFRLFARTRSMWSFNPETNILLNVPLHGTILTRPLLESELVI

GAVILRGHLRIAGHHLGRCDIKDLPKEITVATSRTLSYYKLGASQRVAGDSGFAAYSRYRIGNYKLNTDH

SSSSDNIALLVQ

>QHD43419.1 membrane glycoprotein [Wuhan seafood market pneumonia virus]

MADSNGTITVEELKKLLEQWNLVIGFLFLTWICLLQFAYANRNRFLYIIKLIFLWLLWPVTLACFVLAAV

YRINWITGGIAIAMACLVGLMWLSYFIASFRLFARTRSMWSFNPETNILLNVPLHGTILTRPLLESELVI

GAVILRGHLRIAGHHLGRCDIKDLPKEITVATSRTLSYYKLGASQRVAGDSGFAAYSRYRIGNYKLNTDH

SSSSDNIALLVQ

**29 protein sequences of envelope protein for COVID-19**

>BBW89519.1 envelope protein [Wuhan seafood market pneumonia virus]

MYSFVSEETGTLIVNSVLLFLAFVVFLLVTLAILTALRLCAYCCNIVNVSLVKPSFYVYSRVKNLNSSRV

PDLLV

>QHU36866.1 envelope protein [Wuhan seafood market pneumonia virus]

MYSFVSEETGTLIVNSVLLFLAFVVFLLVTLAILTALRLCAYCCNIVNVSLVKPSFYVYSRVKNLNSSRV

PDLLV

>QHU36856.1 envelope protein [Wuhan seafood market pneumonia virus]

MYSFVSEETGTLIVNSVLLFLAFVVFLLVTLAILTALRLCAYCCNIVNVSLVKPSFYVYSRVKNLNSSRV

PDLLV

>QHU36846.1 envelope protein [Wuhan seafood market pneumonia virus]

MYSFVSEETGTLIVNSVLLFLAFVVFLLVTLAILTALRLCAYCCNIVNVSLVKPSFYVYSRVKNLNSSRV

PDLLV

>QHU36836.1 envelope protein [Wuhan seafood market pneumonia virus]

MYSFVSEETGTLIVNSVLLFLAFVVFLLVTLAILTALRLCAYCCNIVNVSLVKPSFYVYSRVKNLNSSRV

PDLLV

>QHU36826.1 envelope protein [Wuhan seafood market pneumonia virus]

MYSFVSEETGTLIVNSVLLFLAFVVFLLVTLAILTALRLCAYCCNIVNVSLVKPSFYVYSRVKNLNSSRV

PDLLV

>QHR84451.1 envelope protein [Wuhan seafood market pneumonia virus]

MYSFVSEETGTLIVNSVLLFLAFVVFLLVTLAILTALRLCAYCCNIVNVSLVKPSFYVYSRVKNLNSSRV

PDLLV

>QHN73812.1 envelope protein [Wuhan seafood market pneumonia virus]

MYSFVSEETGTLIVNSVLLFLAFVVFLLVTLAILTALRLCAYCCNIVNVSLVKPSFYVYSRVKNLNSSRV

PDLLV

>QHN73797.1 envelope protein [Wuhan seafood market pneumonia virus]

MYSFVSEETGTLIVNSVLLFLAFVVFLLVTLAILTALRLCAYCCNIVNVSLVKPSFYVYSRVKNLNSSRV

PDLLV

>YP_009724392.1 envelope protein [Wuhan seafood market pneumonia virus]

MYSFVSEETGTLIVNSVLLFLAFVVFLLVTLAILTALRLCAYCCNIVNVSLVKPSFYVYSRVKNLNSSRV

PDLLV

>QHD43418.1 envelope protein [Wuhan seafood market pneumonia virus]

MYSFVSEETGTLIVNSVLLFLAFVVFLLVTLAILTALRLCAYCCNIVNVSLVKPSFYVYSRVKNLNSSRV

PDLLV

>QHO62113.1 envelope protein [Wuhan seafood market pneumonia virus]

MYSFVSEETGTLIVNSVLLFLAFVVFLLVTLAILTALRLCAYCCNIVNVSLVKPSFYVYSRVKNLNSSRV

PDLLV

>QHO62108.1 envelope protein [Wuhan seafood market pneumonia virus]

MYSFVSEETGTLIVNSVLLFLAFVVFLLVTLAILTALRLCAYCCNIVNVSLVKPSFYVYSRVKNLNSSRV

PDLLV

>QHW06061.1 envelope protein [Wuhan seafood market pneumonia virus]

MYSFVSEETGTLIVNSVLLFLAFVVFLLVTLAILTALRLCAYCCNIVNVSLVKPSFYVYSRVKNLNSSRV

PDLLV

>QHW06051.1 envelope protein [Wuhan seafood market pneumonia virus]

MYSFVSEETGTLIVNSVLLFLAFVVFLLVTLAILTALRLCAYCCNIVNVSLVKPSFYVYSRVKNLNSSRV

PDLLV

>QHW06041.1 envelope protein [Wuhan seafood market pneumonia virus]

MYSFVSEETGTLIVNSVLLFLAFVVFLLVTLAILTALRLCAYCCNIVNVSLVKPSFYVYSRVKNLNSSRV

PDLLV

>QHU79206.1 envelope protein [Wuhan seafood market pneumonia virus]

MYSFVSEETGTLIVNSVLLFLAFVVFLLVTLAILTALRLCAYCCNIVNVSLVKPSFYVYSRVKNLNSSRV

PDLLV

>QHU79196.1 envelope protein [Wuhan seafood market pneumonia virus]

MYSFVSEETGTLIVNSVLLFLAFVVFLLVTLAILTALRLCAYCCNIVNVSLVKPSFYVYSRVKNLNSSRV

PDLLV

>QHU79175.1 envelope protein [Wuhan seafood market pneumonia virus]

MYSFVSEETGTLIVNSVLLFLAFVVFLLVTLAILTALRLCAYCCNIVNVSLVKPSFYVYSRVKNLNSSRV

PDLLV

>QHR63292.1 envelope protein [Wuhan seafood market pneumonia virus]

MYSFVSEETGTLIVNSVLLFLAFVVFLLVTLAILTALRLCAYCCNIVNVSLVKPSFYVYSRVKNLNSSRV

PDLLV

>QHR63282.1 envelope protein [Wuhan seafood market pneumonia virus]

MYSFVSEETGTLIVNSVLLFLAFVVFLLVTLAILTALRLCAYCCNIVNVSLVKPSFYVYSRVKNLNSSRV

PDLLV

>QHR63272.1 envelope protein [Wuhan seafood market pneumonia virus]

MYSFVSEETGTLIVNSVLLFLAFVVFLLVTLAILTALRLCAYCCNIVNVSLVKPSFYVYSRVKNLNSSRV

PDLLV

>QHR63262.1 envelope protein [Wuhan seafood market pneumonia virus]

MYSFVSEETGTLIVNSVLLFLAFVVFLLVTLAILTALRLCAYCCNIVNVSLVKPSFYVYSRVKNLNSSRV

PDLLV

>QHR63252.1 envelope protein [Wuhan seafood market pneumonia virus]

MYSFVSEETGTLIVNSVLLFLAFVVFLLVTLAILTALRLCAYCCNIVNVSLVKPSFYVYSRVKNLNSSRV

PDLLV

>QHO60596.1 envelope protein [Wuhan seafood market pneumonia virus]

MYSFVSEETGTLIVNSVLLFLAFVVFLLVTLAILTALRLCAYCCNIVNVSLVKPSFYVYSRVKNLNSSRV

PDLLV

>QHQ82466.1 envelope protein [Wuhan seafood market pneumonia virus]

MYSFVSEETGTLIVNSVLLFLAFVVFLLVTLAILTALRLCAYCCNIVNVSLVKPSFYVYSRVKNLNSSRV

PDLLV

>QHQ71975.1 envelope protein [Wuhan seafood market pneumonia virus]

MYSFVSEETGTLIVNSVLLFLAFVVFLLVTLAILTALRLCAYCCNIVNVSLVKPSFYVYSRVKNLNSSRV

PDLLV

>QHQ71965.1 envelope protein [Wuhan seafood market pneumonia virus]

MYSFVSEETGTLIVNSVLLFLAFVVFLLVTLAILTALRLCAYCCNIVNVSLVKPSFYVYSRVKNLNSSRV

PDLLV

>QHO62879.1 envelope protein [Wuhan seafood market pneumonia virus]

MYSFVSEETGTLIVNSVLLFLAFVVFLLVTLAILTALRLCAYCCNIVNVSLVKPSFYVYSRVKNLNSSRV

PDLLV

**29 protein sequences of nucleocapsid protein for COVID-19**

>QHW06066.1 nucleocapsid phosphoprotein [Wuhan seafood market pneumonia virus]

MSDNGPQNQRNAPRITFGGPSDSTGSNQNGERSGARSKQRRPQGLPNNTASWFTALTQHGKEDLKFPRGQ

GVPINTNSSPDDQIGYYRRATRRIRGGDGKMKDLSPRWYFYYLGTGPEAGLPYGANKDGIIWVATEGALN

TPKDHIGTRNPANNAAIVLQLPQGTTLPKGFYAEGSRGGSQASSRSSSRSRNSSRNSTPGSSRGTSPARM

AGNGGDAALALLLLDRLNQLESKMSGKGQQQQGQTVTKKSAAEASKKPRQKRTATKAYNVTQAFGRRGPE

QTQGNFGDQELIRQGTDYKHWPQIAQFAPSASAFFGMSRIGMEVTPSGTWLTYTGAIKLDDKDPNFKDQV

ILLNKHIDAYKTFPPTEPKKDKKKKADETQALPQRQKKQQTVTLLPAADLDDFSKQLQQSMSSADSTQA

>QHW06056.1 nucleocapsid phosphoprotein [Wuhan seafood market pneumonia virus]

MSDNGPQNQRNAPRITFGGPSDSTGSNQNGERSGARSKQRRPQGLPNNTASWFTALTQHGKEDLKFPRGQ

GVPINTNSSPDDQIGYYRRATRRIRGGDGKMKDLSPRWYFYYLGTGPEAGLPYGANKDGIIWVATEGALN

TPKDHIGTRNPANNAAIVLQLPQGTTLPKGFYAEGSRGGSQASSRSSSRSRNSLRNSTPGSSRGTSPARM

AGNGGDAALALLLLDRLNQLESKMSGKGQQQQGQTVTKKSAAEASKKPRQKRTATKAYNVTQAFGRRGPE

QTQGNFGDQELIRQGTDYKHWPQIAQFAPSASAFFGMSRIGMEVTPSGTWLTYTGAIKLDDKDPNFKDQV

ILLNKHIDAYKTFPPTEPKKDKKKKADETQALPQRQKKQQTVTLLPAADLDDFSKQLQQSMSSADSTQA

>QHW06046.1 nucleocapsid phosphoprotein [Wuhan seafood market pneumonia virus]

MSDNGPQNQRNAPRITFGGPSDSTGSNQNGERSGARSKQRRPQGLPNNTASWFTALTQHGKEDLKFPRGQ

GVPINTNSSPDDQIGYYRRATRRIRGGDGKMKDLSPRWYFYYLGTGPEAGLPYGANKDGIIWVATEGALN

TPKDHIGTRNPANNAAIVLQLPQGTTLPKGFYAEGSRGGSQASSRSSSRSRNSLRNSTPGSSRGTSPARM

AGNGGDAALALLLLDRLNQLESKMSGKGQQQQGQTVTKKSAAEASKKPRQKRTATKAYNVTQAFGRRGPE

QTQGNFGDQELIRQGTDYKHWPQIAQFAPSASAFFGMSRIGMEVTPSGTWLTYTGAIKLDDKDPNFKDQV

ILLNKHIDAYKTFPPTEPKKDKKKKADETQALPQRQKKQQTVTLLPAADLDDFSKQLQQSMSSADSTQA

>BBW89524.1 nucleocapsid phosphoprotein [Wuhan seafood market pneumonia virus]

MSDNGPQNQRNAPRITFGGPSDSTGSNQNGERSGARSKQRRPQGLPNNTASWFTALTQHGKEDLKFPRGQ

GVPINTNSSPDDQIGYYRRATRRIRGGDGKMKDLSPRWYFYYLGTGPEAGLPYGANKDGIIWVATEGALN

TPKDHIGTRNPANNAAIVLQLPQGTTLPKGFYAEGSRGGSQASSRSSSRSRNSSRNSTPGSSRGTSPARM

AGNGGDAALALLLLDRLNQLESKMSGKGQQQQGQTVTKKSAAEASKKPRQKRTATKAYNVTQAFGRRGPE

QTQGNFGDQELIRQGTDYKHWPQIAQFAPSASAFFGMSRIGMEVTPSGTWLTYTGAIKLDDKDPNFKDQV

ILLNKHIDAYKTFPPTEPKKDKKKKADETQALPQRQKKQQTVTLLPAADLDDFSKQLQQSMSSADSTQA

>QHU79211.1 nucleocapsid phosphoprotein [Wuhan seafood market pneumonia virus]

MSDNGPQNQRNAPRITFGGPSDSTGSNQNGERSGARSKQRRPQGLPNNTASWFTALTQHGKEDLKFPRGQ

GVPINTNSSPDDQIGYYRRATRRIRGGDGKMKDLSPRWYFYYLGTGPEAGLPYGANKDGIIWVATEGALN

TPKDHIGTRNPANNAAIVLQLPQGTTLPKGFYAEGSRGGSQASSRSSSRSRNSSRNSTPGSSRGTSPARM

AGNGGDAALALLLLDRLNQLESKMSGKGQQQQGQTVTKKSAAEASKKPRQKRTATKAYNVTQAFGRRGPE

QTQGNFGDQELIRQGTDYKHWPQIAQFAPSASAFFGMSRIGMEVTPSGTWLTYTGAIKLDDKDPNFKDQV

ILLNKHIDAYKTFPPTEPKKDKKKKADETQALPQRQKKQQTVTLLPAADLDDFSKQLQQSMSSADSTQA

>QHU79201.1 nucleocapsid phosphoprotein [Wuhan seafood market pneumonia virus]

MSDNGPQNQRNAPRITFGGPSDSTGSNQNGERSGARSKQRRPQGLPNNTASWFTALTQHGKEDLKFPRGQ

GVPINTNSSPDDQIGYYRRATRRIRGGDGKMKDLSPRWYFYYLGTGPEAGLPYGANKDGIIWVATEGALN

TPKDHIGTRNPANNAAIVLQLPQGTTLPKGFYAEGSRGGSQASSRSSSRSRNSSRNSTPGSSRGTSPARM

AGNGGDAALALLLLDRLNQLESKMSGKGQQQQGQTVTKKSAAEASKKPRQKRTATKAYNVTQAFGRRGPE

QTQGNFGDQELIRQGTDYKHWPQIAQFAPSASAFFGMSRIGMEVTPSGTWLTYTGAIKLDDKDPNFKDQV

ILLNKHIDAYKTFPPTEPKKDKKKKADETQALPQRQKKQQTVTLLPAADLDDFSKQLQQSMSSADSTQA

>QHU79181.1 nucleocapsid phosphoprotein [Wuhan seafood market pneumonia virus]

MSDNGPQNQRNAPRITFGGPSDSTGSNQNGERSGARSKQRRPQGLPNNTASWFTALTQHGKEDLKFPRGQ

GVPINTNSSPDDQIGYYRRATRRIRGGDGKMKDLSPRWYFYYLGTGPEAGLPYGANKDGIIWVATEGALN

TPKDHIGTRNPANNAAIVLQLPQGTTLPKGFYAEGSRGGSQASSRSSSRSRNSSRNSTPGSSRGTSPARM

AGNGGDAALALLLLDRLNQLESKMSGKGQQQQGQTVTKKSAAEASKKPRQKRTATKAYNVTQAFGRRGPE

QTQGNFGDQELIRQGTDYKHWPQIAQFAPSASAFFGMSRIGMEVTPSGTWLTYTGAIKLDDKDPNFKDQV

ILLNKHIDAYKTFPPTEPKKDKKKKADETQALPQRQKKQQTVTLLPAADLDDFSKQLQQSMSSADSTQA

>QHU36871.1 nucleocapsid phosphoprotein [Wuhan seafood market pneumonia virus]

MSDNGPQNQRNAPRITFGGPSDSTGSNQNGERSGARSKQRRPQGLPNNTASWFTALTQHGKEDLKFPRGQ

GVPINTNSSPDDQIGYYRRATRRIRGGDGKMKDLSPRWYFYYLGTGPEAGLPYGANKDGIIWVATEGALN

TPKDHIGTRNPANNAAIVLQLPQGTTLPKGFYAEGSRGGSQASSRSSSRSRNSSRNSTPGSSRGTSPARM

AGNGGDAALALLLLDRLNQLESKMSGKGQQQQGQTVTKKSAAEASKKPRQKRTATKAYNVTQAFGRRGPE

QTQGNFGDQELIRQGTDYKHWPQIAQFAPSASAFFGMSRIGMEVTPSGTWLTYTGAIKLDDKDPNFKDQV

ILLNKHIDAYKTFPPTEPKKDKKKKADETQALPQRQKKQQTVTLLPAADLDDFSKQLQQSMSSADSTQA

>QHU36861.1 nucleocapsid phosphoprotein [Wuhan seafood market pneumonia virus]

MSDNGPQNQRNAPRITFGGPSDSTGSNQNGERSGARSKQRRPQGLPNNTASWFTALTQHGKEDLKFPRGQ

GVPINTNSSPDDQIGYYRRATRRIRGGDGKMKDLSPRWYFYYLGTGPEAGLPYGANKDGIIWVATEGALN

TPKDHIGTRNPANNAAIVLQLPQGTTLPKGFYAEGSRGGSQASSRSSSRSRNSSRNSTPGSSRGTSPARM

AGNGGDAALALLLLDRLNQLESKMSGKGQQQQGQTVTKKSAAEASKKPRQKRTATKAYNVTQAFGRRGPE

QTQGNFGDQELIRQGTDYKHWPQIAQFAPSASAFFGMSRIGMEVTPSGTWLTYTGAIKLDDKDPNFKDQV

ILLNKHIDAYKTFPPTEPKKDKKKKADETQALPQRQKKQQTVTLLPAADLDDFSKQLQQSMSSADSTQA

>QHU36851.1 nucleocapsid phosphoprotein [Wuhan seafood market pneumonia virus]

MSDNGPQNQRNAPRITFGGPSDSTGSNQNGERSGARSKQRRPQGLPNNTASWFTALTQHGKEDLKFPRGQ

GVPINTNSSPDDQIGYYRRATRRIRGGDGKMKDLSPRWYFYYLGTGPEAGLPYGANKDGIIWVATEGALN

TPKDHIGTRNPANNAAIVLQLPQGTTLPKGFYAEGSRGGSQASSRSSSRSRNSSRNSTPGSSRGTSPARM

AGNGGDAALALLLLDRLNQLESKMSGKGQQQQGQTVTKKSAAEASKKPRQKRTATKAYNVTQAFGRRGPE

QTQGNFGDQELIRQGTDYKHWPQIAQFAPSASAFFGMSRIGMEVTPSGTWLTYTGAIKLDDKDPNFKDQV

ILLNKHIDAYKTFPPTEPKKDKKKKADETQALPQRQKKQQTVTLLPAADLDDFSKQLQQSMSSADSTQA

>QHU36841.1 nucleocapsid phosphoprotein [Wuhan seafood market pneumonia virus]

MSDNGPQNQRNAPRITFGGPSDSTGSNQNGERSGARSKQRRPQGLPNNTASWFTALTQHGKEDLKFPRGQ

GVPINTNSSPDDQIGYYRRATRRIRGGDGKMKDLSPRWYFYYLGTGPEAGLPYGANKDGIIWVATEGALN

TPKDHIGTRNPANNAAIVLQLPQGTTLPKGFYAEGSRGGSQASSRSSSRSRNSSRNSTPGSSRGTSPARM

AGNGGDAALALLLLDRLNQLESKMSGKGQQQQGQTVTKKSAAEASKKPRQKRTATKAYNVTQAFGRRGPE

QTQGNFGDQELIRQGTDYKHWPQIAQFAPSASAFFGMSRIGMEVTPSGTWLTYTGAIKLDDKDPNFKDQV

ILLNKHIDAYKTFPPTEPKKDKKKKADETQALPQRQKKQQTVTLLPAADLDDFSKQLQQSMSSADSTQA

>QHU36831.1 nucleocapsid phosphoprotein [Wuhan seafood market pneumonia virus]

MSDNGPQNQRNAPRITFGGPSDSTGSNQNGERSGARSKQRRPQGLPNNTASWFTALTQHGKEDLKFPRGQ

GVPINTNSSPDDQIGYYRRATRRIRGGDGKMKDLSPRWYFYYLGTGPEAGLPYGANKDGIIWVATEGALN

TPKDHIGTRNPANNAAIVLQLPQGTTLPKGFYAEGSRGGSQASSRSSSRSRNSSRNSTPGSSRGTSPARM

AGNGGDAALALLLLDRLNQLESKMSGKGQQQQGQTVTKKSAAEASKKPRQKRTATKAYNVTQAFGRRGPE

QTQGNFGDQELIRQGTDYKHWPQIAQFAPSASAFFGMSRIGMEVTPSGTWLTYTGAIKLDDKDPNFKDQV

ILLNKHIDAYKTFPPTEPKKDKKKKADETQALPQRQKKQQTVTLLPAADLDDFSKQLQQSMSSADSTQA

>QHR84456.1 nucleocapsid phosphoprotein [Wuhan seafood market pneumonia virus]

MSDNGPQNQRNAPRITFGGPSDSTGSNQNGERSGARSKQRRPQGLPNNTASWFTALTQHGKEDLKFPRGQ

GVPINTNSSPDDQIGYYRRATRRIRGGDGKMKDLSPRWYFYYLGTGPEAGLPYGANKDGIIWVATEGALN

TPKDHIGTRNPANNAAIVLQLPQGTTLPKGFYAEGSRGGSQASSRSSSRSRNSSRNSTPGSSRGTSPARM

AGNGGDAALALLLLDRLNQLESKMSGKGQQQQGQTVTKKSAAEASKKPRQKRTATKAYNVTQAFGRRGPE

QTQGNFGDQELIRQGTDYKHWPQIAQFAPSASAFFGMSRIGMEVTPSGTWLTYTGAIKLDDKDPNFKDQV

ILLNKHIDAYKTFPPTEPKKDKKKKADETQALPQRQKKQQTVTLLPAADLDDFSKQLQQSMSSADSTQA

>QHN73817.1 nucleocapsid phosphoprotein [Wuhan seafood market pneumonia virus]

MSDNGPQNQRNAPRITFGGPSDSTGSNQNGERSGARSKQRRPQGLPNNTASWFTALTQHGKEDLKFPRGQ

GVPINTNSSPDDQIGYYRRATRRIRGGDGKMKDLSPRWYFYYLGTGPEAGLPYGANKDGIIWVATEGALN

TPKDHIGTRNPANNAAIVLQLPQGTTLPKGFYAEGSRGGSQASSRSSSRSRNSSRNSTPGSSRGTSPARM

AGNGGDAALALLLLDRLNQLESKMSGKGQQQQGQTVTKKSAAEASKKPRQKRTATKAYNVTQAFGRRGPE

QTQGNFGDQELIRQGTDYKHWPQIAQFAPSASAFFGMSRIGMEVTPSGTWLTYTGAIKLDDKDPNFKDQV

ILLNKHIDAYKTFPPTEPKKDKKKKADETQALPQRQKKQQTVTLLPAADLDDFSKQLQQSMSSADSTQA

>QHN73802.1 nucleocapsid phosphoprotein [Wuhan seafood market pneumonia virus]

MSDNGPQNQRNAPRITFGGPSDSTGSNQNGERSGARSKQRRPQGLPNNTASWFTALTQHGKEDLKFPRGQ

GVPINTNSSPDDQIGYYRRATRRIRGGDGKMKDLSPRWYFYYLGTGPEAGLPYGANKDGIIWVATEGALN

TPKDHIGTRNPANNAAIVLQLPQGTTLPKGFYAEGSRGGSQASSRSSSRSRNSSRNSTPGSSRGTSPARM

AGNGGDAALALLLLDRLNQLESKMSGKGQQQQGQTVTKKSAAEASKKPRQKRTATKAYNVTQAFGRRGPE

QTQGNFGDQELIRQGTDYKHWPQIAQFAPSASAFFGMSRIGMEVTPSGTWLTYTGAIKLDDKDPNFKDQV

ILLNKHIDAYKTFPPTEPKKDKKKKADETQALPQRQKKQQTVTLLPAADLDDFSKQLQQSMSSADSTQA

>QHR63298.1 nucleocapsid protein [Wuhan seafood market pneumonia virus]

MSDNGPQNQRNAPRITFGGPSDSTGSNQNGERSGARSKQRRPQGLPNNTASWFTALTQHGKEDLKFPRGQ

GVPINTNSSPDDQIGYYRRATRRIRGGDGKMKDLSPRWYFYYLGTGPEAGLPYGANKDGIIWVATEGALN

TPKDHIGTRNPANNAAIVLQLPQGTTLPKGFYAEGSRGGSQASSRSSSRSRNSSRNSTPGSSRGTSPARM

AGNGGDAALALLLLDRLNQLESKMSGKGQQQQGQTVTKKSAAEASKKPRQKRTATKAYNVTQAFGRRGPE

QTQGNFGDQELIRQGTDYKHWPQIAQFAPSASAFFGMSRIGMEVTPSGTWLTYTGAIKLDDKDPNFKDQV

ILLNKHIDAYKTFPPTEPKKDKKKKADETQALPQRQKKQQTVTLLPAADLDDFSKQLQQSMSSADSTQA

>QHR63288.1 nucleocapsid protein [Wuhan seafood market pneumonia virus]

MSDNGPQNQRNAPRITFGGPSDSTGSNQNGERSGARSKQRRPQGLPNNTASWFTALTQHGKEDLKFPRGQ

GVPINTNSSPDDQIGYYRRATRRIRGGDGKMKDLSPRWYFYYLGTGPEAGLPYGANKDGIIWVATEGALN

TPKDHIGTRNPANNAAIVLQLPQGTTLPKGFYAEGSRGGSQASSRSSSRSRNSSRNSTPGSSRGTSPARM

AGNGGDAALALLLLDRLNQLESKMSGKGQQQQGQTVTKKSAAEASKKPRQKRTATKAYNVTQAFGRRGPE

QTQGNFGDQELIRQGTDYKHWPQIAQFAPSASAFFGMSRIGMEVTPSGTWLTYTGAIKLDDKDPNFKDQV

ILLNKHIDAYKTFPPTEPKKDKKKKADETQALPQRQKKQQTVTLLPAADLDDFSKQLQQSMSSADSTQA

>QHR63278.1 nucleocapsid protein [Wuhan seafood market pneumonia virus]

MSDNGPQNQRNAPRITFGGPSDSTGSNQNGERSGARSKQRRPQGLPNNTASWFTALTQHGKEDLKFPRGQ

GVPINTNSSPDDQIGYYRRATRRIRGGDGKMKDLSPRWYFYYLGTGPEAGLPYGANKDGIIWVATEGALN

TPKDHIGTRNPANNAAIVLQLPQGTTLPKGFYAEGSRGGSQASSRSSSRSRNSSRNSTPGSSRGTSPARM

AGNGGDAALALLLLDRLNQLESKMSGKGQQQQGQTVTKKSAAEASKKPRQKRTATKAYNVTQAFGRRGPE

QTQGNFGDQELIRQGTDYKHWPQIAQFAPSASAFFGMSRIGMEVTPSGTWLTYTGAIKLDDKDPNFKDQV

ILLNKHIDAYKTFPPTEPKKDKKKKADETQALPQRQKKQQTVTLLPAADLDDFSKQLQQSMSSADSTQA

>QHR63268.1 nucleocapsid protein [Wuhan seafood market pneumonia virus]

MSDNGPQNQRNAPRITFGGPSDSTGSNQNGERSGARSKQRRPQGLPNNTASWFTALTQHGKEDLKFPRGQ

GVPINTNSSPDDQIGYYRRATRRIRGGDGKMKDLSPRWYFYYLGTGPEAGLPYGANKDGIIWVATEGALN

TPKDHIGTRNPANNAAIVLQLPQGTTLPKGFYAEGSRGGSQASSRSSSRSRNSSRNSTPGSSRGTSPARM

AGNGGDAALALLLLDRLNQLESKMSGKGQQQQGQTVTKKSAAEASKKPRQKRTATKAYNVTQAFGRRGPE

QTQGNFGDQELIRQGTDYKHWPQIAQFAPSASAFFGMSRIGMEVTPSGTWLTYTGAIKLDDKDPNFKDQV

ILLNKHIDAYKTFPPTEPKKDKKKKADETQALPQRQKKQQTVTLLPAADLDDFSKQLQQSMSSADSTQA

>QHR63258.1 nucleocapsid protein [Wuhan seafood market pneumonia virus]

MSDNGPQNQRNAPRITFGGPSDSTGSNQNGERSGARSKQRRPQGLPNNTASWFTALTQHGKEDLKFPRGQ

GVPINTNSSPDDQIGYYRRATRRIRGGDGKMKDLSPRWYFYYLGTGPEAGLPYGANKDGIIWVATEGALN

TPKDHIGTRNPANNAAIVLQLPQGTTLPKGFYAEGSRGGSQASSRSSSRSRNSSRNSTPGSSRGTSPARM

AGNGGDAALALLLLDRLNQLESKMSGKGQQQQGQTVTKKSAAEASKKPRQKRTATKAYNVTQAFGRRGPE

QTQGNFGDQELIRQGTDYKHWPQIAQFAPSASAFFGMSRIGMEVTPSGTWLTYTGAIKLDDKDPNFKDQV

ILLNKHIDAYKTFPPTEPKKDKKKKADETQALPQRQKKQQTVTLLPAADLDDFSKQLQQSMSSADSTQA

>QHO60601.1 nucleocapsid phosphoprotein [Wuhan seafood market pneumonia virus]

MSDNGPQNQRNAPRITFGGPSDSTGSNQNGERSGARSKQRRPQGLPNNTASWFTALTQHGKEDLKFPRGQ

GVPINTNSSPDDQIGYYRRATRRIRGGDGKMKDLSPRWYFYYLGTGPEAGLPYGANKDGIIWVATEGALN

TPKDHIGTRNPANNAAIVLQLPQGTTLPKGFYAEGSRGGSQASSRSSSRSRNSSRNSTPGSSRGTSPARM

AGNGGDAALALLLLDRLNQLESKMSGKGQQQQGQTVTKKSAAEASKKPRQKRTATKAYNVTQAFGRRGPE

QTQGNFGDQELIRQGTDYKHWPQIAQFAPSASAFFGMSRIGMEVTPSGTWLTYTGAIKLDDKDPNFKDQV

ILLNKHIDAYKTFPPTEPKKDKKKKADETQALPQRQKKQQTVTLLPAADLDDFSKQLQQSMSSADSTQA

>QHQ82471.1 nucleocapsid phosphoprotein [Wuhan seafood market pneumonia virus]

MSDNGPQNQRNAPRITFGGPSDSTGSNQNGERSGARSKQRRPQGLPNNTASWFTALTQHGKEDLKFPRGQ

GVPINTNSSPDDQIGYYRRATRRIRGGDGKMKDLSPRWYFYYLGTGPEAGLPYGANKDGIIWVATEGALN

TPKDHIGTRNPANNAAIVLQLPQGTTLPKGFYAEGSRGGSQASSRSSSRSRNSSRNSTPGSSRGTSPARM

AGNGGDAALALLLLDRLNQLESKMSGKGQQQQGQTVTKKSAAEASKKPRQKRTATKAYNVTQAFGRRGPE

QTQGNFGDQELIRQGTDYKHWPQIAQFAPSASAFFGMSRIGMEVTPSGTWLTYTGAIKLDDKDPNFKDQV

ILLNKHIDAYKTFPPTEPKKDKKKKADETQALPQRQKKQQTVTLLPAADLDDFSKQLQQSMSSADSTQA

>QHQ71980.1 nucleocapsid phosphoprotein [Wuhan seafood market pneumonia virus]

MSDNGPQNQRNAPRITFGGPSDSTGSNQNGERSGARSKQRRPQGLPNNTASWFTALTQHGKEDLKFPRGQ

GVPINTNSSPDDQIGYYRRATRRIRGGDGKMKDLSPRWYFYYLGTGPEAGLPYGANKDGIIWVATEGALN

TPKDHIGTRNPANNAAIVLQLPQGTTLPKGFYAEGSRGGSQASSRSSSRSRNSSRNSTPGSSRGTSPARM

AGNGGDAALALLLLDRLNQLESKMSGKGQQQQGQTVTKKSAAEASKKPRQKRTATKAYNVTQAFGRRGPE

QTQGNFGDQELIRQGTDYKHWPQIAQFAPSASAFFGMSRIGMEVTPSGTWLTYTGAIKLDDKDPNFKDQV

ILLNKHIDAYKTFPPTEPKKDKKKKADETQALPQRQKKQQTVTLLPAADLDDFSKQLQQSMSSADSTQA

>QHQ71970.1 nucleocapsid phosphoprotein [Wuhan seafood market pneumonia virus]

MSDNGPQNQRNAPRITFGGPSDSTGSNQNGERSGARSKQRRPQGLPNNTASWFTALTQHGKEDLKFPRGQ

GVPINTNSSPDDQIGYYRRATRRIRGGDGKMKDLSPRWYFYYLGTGPEAGLPYGANKDGIIWVATEGALN

TPKDHIGTRNPANNAAIVLQLPQGTTLPKGFYAEGSRGGSQASSRSSSRSRNSSRNSTPGSSRGTSPARM

AGNGGDAALALLLLDRLNQLESKMSGKGQQQQGQTVTKKSAAEASKKPRQKRTATKAYNVTQAFGRRGPE

QTQGNFGDQELIRQGTDYKHWPQIAQFAPSASAFFGMSRIGMEVTPSGTWLTYTGAIKLDDKDPNFKDQV

ILLNKHIDAYKTFPPTEPKKDKKKKADETQALPQRQKKQQTVTLLPAADLDDFSKQLQQSMSSADSTQA

>QHO62115.1 nucleocapsid protein [Wuhan seafood market pneumonia virus]

MSDNGPQNQRNAPRITFGGPSDSTGSNQNGERSGARSKQRRPQGLPNNTASWFTALTQHGKEDLKFPRGQ

GVPINTNSSPDDQIGYYRRATRRIRGGDGKMKDLSPRWYFYYLGTGPEAGLPYGANKDGIIWVATEGALN

TPKDHIGTRNPANNAAIVLQLPQGTTLPKGFYAEGSRGGSQASSRSSSRSRNSSRNSTPGSSRGTSPARM

AGNGGDAALALLLLDRLNQLESKMSGKGQQQQGQTVTKKSAAEASKKPRQKRTATKAYNVTQAFGRRGPE

QTQGNFGDQELIRQGTDYKHWPQIAQFAPSASAFFGMSRIGMEVTPSGTWLTYTGAIKLDDKDPNFKDQV

ILLNKHIDAYKTFPPTEPKKDKKKKADETQALPQRQKKQQTVTLLPAADLDDFSKQLQQSMSSADSTQA

>QHO62110.1 nucleocapsid protein [Wuhan seafood market pneumonia virus]

MSDNGPQNQRNAPRITFGGPSDSTGSNQNGERSGARSKQRRPQGLPNNTASWFTALTQHGKEDLKFPRGQ

GVPINTNSSPDDQIGYYRRATRRIRGGDGKMKDLSPRWYFYYLGTGPEAGLPYGANKDGIIWVATEGALN

TPKDHIGTRNPANNAAIVLQLPQGTTLPKGFYAEGSRGGSQASSRSSSRSRNSSRNSTPGSSRGTSPARM

AGNGGDAALALLLLDRLNQLESKMSGKGQQQQGQTVTKKSAAEASKKPRQKRTATKAYNVTQAFGRRGPE

QTQGNFGDQELIRQGTDYKHWPQIAQFAPSASAFFGMSRIGMEVTPSGTWLTYTGAIKLDDKDPNFKDQV

ILLNKHIDAYKTFPPTEPKKDKKKKADETQALPQRQKKQQTVTLLPAADLDDFSKQLQQSMSSADSTQA

>YP_009724397.2 nucleocapsid phosphoprotein [Wuhan seafood market pneumonia virus]

MSDNGPQNQRNAPRITFGGPSDSTGSNQNGERSGARSKQRRPQGLPNNTASWFTALTQHGKEDLKFPRGQ

GVPINTNSSPDDQIGYYRRATRRIRGGDGKMKDLSPRWYFYYLGTGPEAGLPYGANKDGIIWVATEGALN

TPKDHIGTRNPANNAAIVLQLPQGTTLPKGFYAEGSRGGSQASSRSSSRSRNSSRNSTPGSSRGTSPARM

AGNGGDAALALLLLDRLNQLESKMSGKGQQQQGQTVTKKSAAEASKKPRQKRTATKAYNVTQAFGRRGPE

QTQGNFGDQELIRQGTDYKHWPQIAQFAPSASAFFGMSRIGMEVTPSGTWLTYTGAIKLDDKDPNFKDQV

ILLNKHIDAYKTFPPTEPKKDKKKKADETQALPQRQKKQQTVTLLPAADLDDFSKQLQQSMSSADSTQA

>QHO62884.1 nucleocapsid phosphoprotein [Wuhan seafood market pneumonia virus]

MSDNGPQNQRNAPRITFGGPSDSTGSNQNGERSGARSKQRRPQGLPNNTASWFTALTQHGKEDLKFPRGQ

GVPINTNSSPDDQIGYYRRATRRIRGGDGKMKDLSPRWYFYYLGTGPEAGLPYGANKDGIIWVATEGALN

TPKDHIGTRNPANNAAIVLQLPQGTTLPKGFYAEGSRGGSQASSRSSSRSRNSXRNSTPGSSRGTSPARM

AGNGGDAALALLLLDRLNQLESKMSGKGQQQQGQTVTKKSAAEASKKPRQKRTATKAYNVTQAFGRRGPE

QTQGNFGDQELIRQGTDYKHWPQIAQFAPSASAFFGMSRIGMEVTPSGTWLTYTGAIKLDDKDPNFKDQV

ILLNKHIDAYKTFPPTEPKKDKKKKADETQALPQRQKKQQTVTLLPAADLDDFSKQLQQSMSSADSTQA

>QHD43423.2 nucleocapsid phosphoprotein [Wuhan seafood market pneumonia virus]

MSDNGPQNQRNAPRITFGGPSDSTGSNQNGERSGARSKQRRPQGLPNNTASWFTALTQHGKEDLKFPRGQ

GVPINTNSSPDDQIGYYRRATRRIRGGDGKMKDLSPRWYFYYLGTGPEAGLPYGANKDGIIWVATEGALN

TPKDHIGTRNPANNAAIVLQLPQGTTLPKGFYAEGSRGGSQASSRSSSRSRNSSRNSTPGSSRGTSPARM

AGNGGDAALALLLLDRLNQLESKMSGKGQQQQGQTVTKKSAAEASKKPRQKRTATKAYNVTQAFGRRGPE

QTQGNFGDQELIRQGTDYKHWPQIAQFAPSASAFFGMSRIGMEVTPSGTWLTYTGAIKLDDKDPNFKDQV

ILLNKHIDAYKTFPPTEPKKDKKKKADETQALPQRQKKQQTVTLLPAADLDDFSKQLQQSMSSADSTQA
